# A versatile enzymatic pathway for modification of peptide C-termini

**DOI:** 10.1101/2025.07.11.664356

**Authors:** Shravan R. Dommaraju, Sanath K. Kandy, Hengqian Ren, Dominic P. Luciano, Shogo Fujiki, David Sarlah, Huimin Zhao, Jonathan R. Chekan, Douglas A. Mitchell

## Abstract

Advances in bioinformatics have enabled the discovery of unique enzymatic reactions, particularly for ribosomally synthesized and post-translationally modified peptides (RiPPs). The recently discovered daptides, peptides with their C-terminus replaced by an amine, represent one such case, but the diversity, requirements, and engineering potential of daptide biosynthesis remain to be established. Using the daptide biosynthetic gene clusters from *Thermobifida fusca* and *Streptomyces azureus*, we reconstituted daptide biosynthesis *in vitro*, revealing the enzymatic requirements for successive oxidative decarboxylation, transamination, and *N,N*-dimethylation. *In vitro* and *in vivo* studies showed a tailoring family of YcaO enzymes convert a secondary amine intermediate to a C-terminal imidazoline. We further demonstrated enzymatic activity toward shortened, leader peptide-free, and non-native core peptides, highlighting a broad substrate tolerance. Using these insights, we directed the daptide pathway to install new C-termini, including a bioconjugation-compatible aminoacetone, on various peptide and protein substrates.

---

Ribosomal peptides and proteins adopt diverse structures, driven by their constituent amino acid sequences. While genetic mutations readily alter the composition of the side chain, other methods are required to edit the polypeptide backbone. One such route involves post-translational modification (PTM), and numerous enzymes have evolved to act on the backbone.^1^ Of particular interest are enzymes that modify the C-terminus.^2^ For example, HRas, a cancer therapeutic target, receives modifications to enhance membrane targeting at a terminal Cys residue, adding a methyl ester to mask the charge at the terminus and an isoprene unit to enhance lipophilicity (**Figure 1**).^3^ Many peptide hormones receive critical C-terminal modifications to enhance their half-life or target engagement.^4–7^ The most frequent of these PTMs is C-terminal amidation as shown for the neuropeptide endomorphin-1. These modifications are also employed by antimicrobial peptides^5,8^ and FDA-approved hormone-analog drugs, such as leuprorelin, which contains a C-terminal *N*-alkyl group.^9^ Methods for enzymatic and selective installation of C-terminal modifications are actively being investigated with the current rise in development of biologic therapeutics.^10,11^

**Figure 1.**
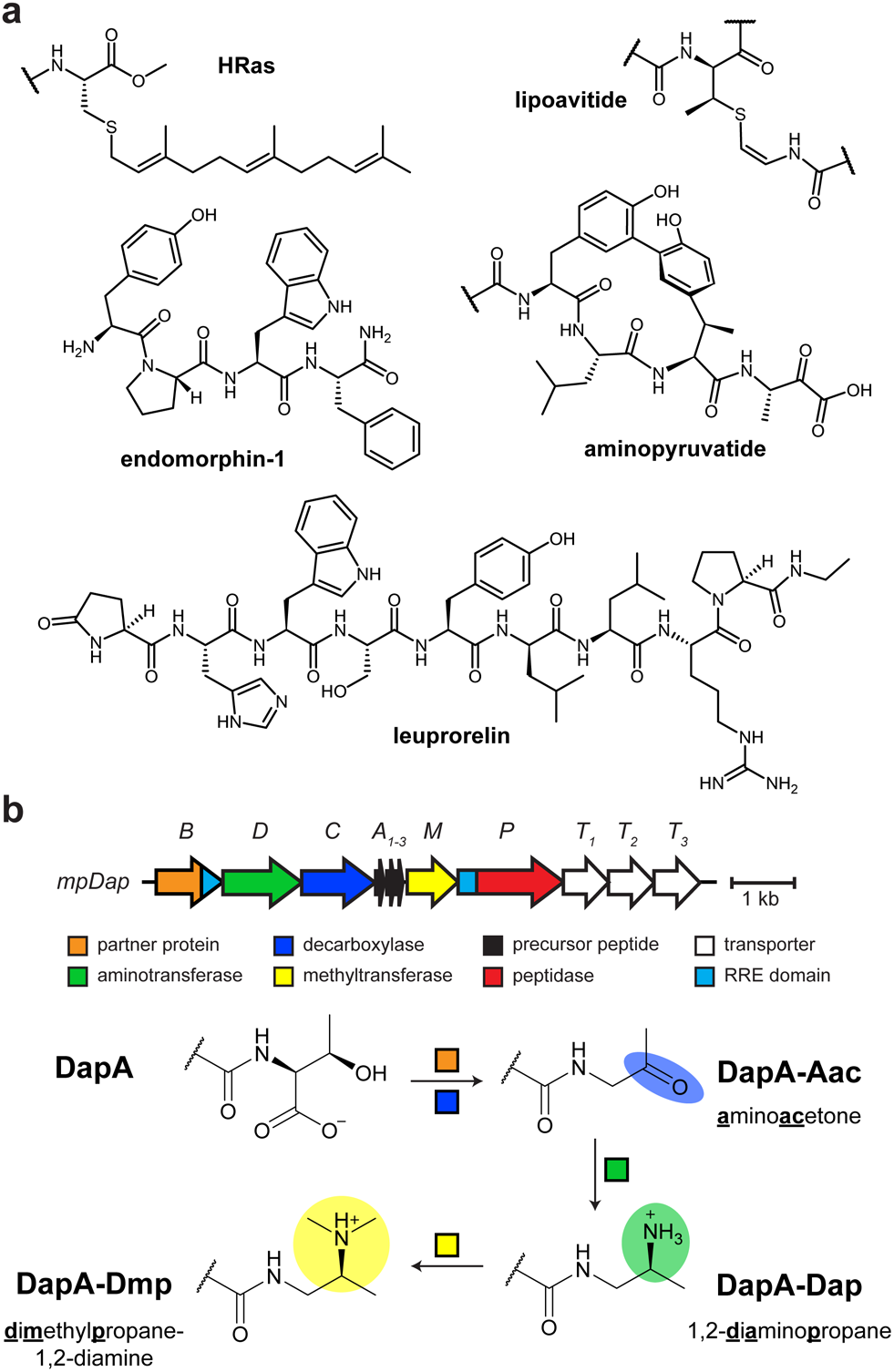
C-terminal modifications of peptides and proteins. **a**, Representative examples of C-terminal post-translational modifications. **b**, Previously reported *mpDap* biosynthetic gene cluster from *Microbacterium paraoxydans* and corresponding daptide biosynthetic scheme. Abbreviations: Aac, aminoacetone, Dap, 1,2-diaminopropane, Dmp, dimethylpropane-1,2-diamine; RRE, RiPP precursor peptide recognition element.

The study of ribosomally synthesized and post-translationally modified peptides (RiPPs) has expanded the availability of enzymes that modify the C-termini of polypeptides.^12,13^ Within RiPP biosynthesis, enzymes have been characterized that can macrocyclize,^14,15^ amidate,^16^ decarboxylate,^17,18^ or install other unusual C-terminal PTMs (**Figure 1**).^19,20^ Recently, we reported on daptides, a RiPP class featuring multi-step modification of a C-terminal Thr.^21^ Most characterized C-terminal PTMs result in neutralization of the negatively charged carboxylate, but daptide biosynthesis swaps the C-terminus for a positively-charged tertiary amine (*S*)-*N*_*2*_,*N*_*2*_-dimethylpropane-1,2-diamine (Dmp; **Figure 1**). Some daptides are hydrophobic α-helical peptides and act as hemolysins^21^ while hominicin is further modified and possesses antibacterial properties.^22^ In each case, the modified C-termini are likely integral to their bioactivity. Our initial report on daptides using the *mpa* (*i*.*e*., *mpDap*) biosynthetic gene cluster (BGC) suggested that daptide enzymes may tolerate diverse substrates.^21^ Thus, we hypothesized that daptide biosynthesis could be repurposed for modular functionalization of peptide C-termini.

In this work, we uncovered the enzymatic basis for daptide biosynthesis and explored the substrate scope and diversity of daptide PTMs. We completed a bioinformatic survey of daptide BGCs and identified representatives from *Thermobifida fusca* and *Streptomyces azureus* that indicated further elaboration of PTMs associated with the daptide class. Through *in vitro* reconstitution, we demonstrated NAD^+^-dependent oxidative decarboxylation, L-Lys-dependent transamination, and SAM-dependent *N,N*-dimethylation. Spectroscopic characterization of YcaO-catalyzed reaction products revealed conversion of the peptide C-terminus to a 3,4-dimethylimidazoline (Diz) moiety (formally, (*1S*)-1-amino-1-[(*4S*)-3,4-dimethyl-2-imidazolin-2-yl]isopentane). Heterologous expression of the *S. azureus* BGC further showed that daptide metabolism is branched, yielding different endpoint PTMs for precursor peptides in the same BGC. The characterized enzymes show broad substrate tolerance, processing shortened and leader-free substrates. Using these enzymes, we installed new C-termini, such as the bioconjugation-compatible aminoacetone, onto non-native substrates, including glucagon and green fluorescent protein (GFP). Altogether, we describe the enzymatic basis for daptide biosynthesis, expand their chemical diversity, and repurpose the pathway to functionalize non-native substrates.

## Results

### Selection of daptide biosynthetic gene clusters

To explore daptide biosynthesis, we first sought daptide BGCs that would allow *in vitro* reconstitution and discovery of expanded PTMs within the daptide RiPP class. Using PSIBLAST and BLAST searches of previously identified daptide proteins, we identified 1,034 putative daptide BGCs encoding 3,013 putative precursor peptides (**Dataset S1, Table S1, Figure S1**). We were intrigued by daptide biosynthetic genes encoded by *Thermobifida fusca*, a thermophilic organism from which other RiPP enzymes have been successfully reconstituted.^23–25^ As the BGC was encoded across multiple contigs, *T. fusca* DSM 43792 was obtained and sequenced, allowing assembly of a complete genome (hereafter, *tfDap* BGC; **Figure 2, Tables S2-S5**). The *tfDap* BGC encodes five precursor peptides (*Tf*DapA_1-5_; **Table S2**) as well as daptide biosynthetic proteins predicted to install the tertiary amine group (*Tf*DapBCDM), remove the leader peptide (*Tf*DapP), and export the mature daptides (*Tf*DapT_1_T_2_). The *tfDap* BGC also encodes homologs of a lanthipeptide dehydratase (*Tf*DapKC) and a luciferase-like monooxygenase (*Tf*DapJ), which have been shown to form D-Ala from L-Ser.^21,26^ Additionally, *tfDap* encodes a YcaO protein (*Tf*DapY).

**Figure 2.**
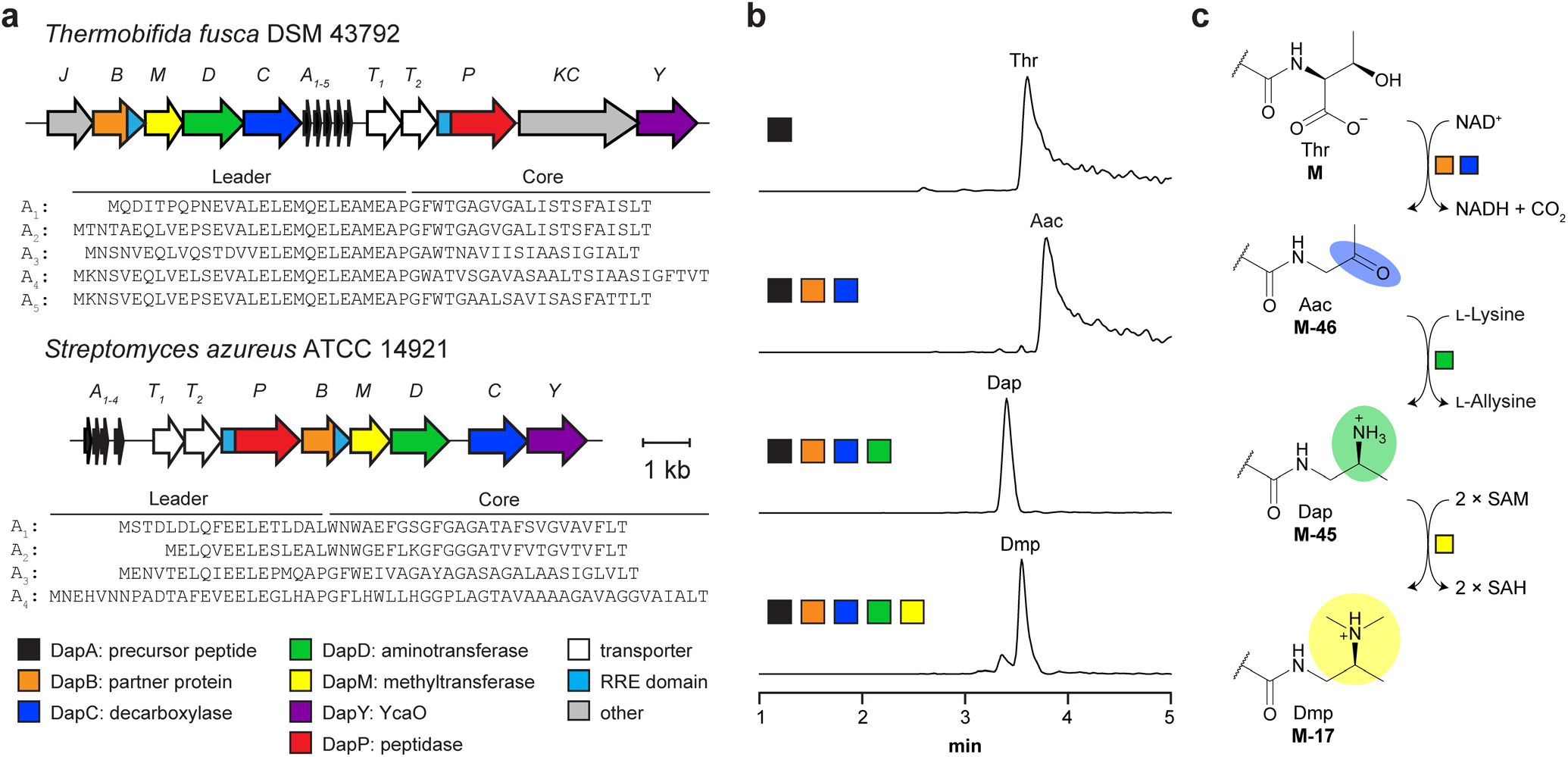
Reconstitution of daptide biosynthesis. **a**, *tfDap* and *saDap* BGC diagrams. **b**, Ion-count normalized extracted ion chromatograms showing only the expected reaction products for *Tf*DapBCDM. The five highest abundance masses in the expected isotopic envelope were extracted for each expected analyte. **c**, Reaction scheme for conversion of Thr to Dmp. Abbreviations: Aac, aminoacetone; Dap, 1,2-diaminopropane; Dmp, dimethylpropane-1,2-diamine; SAM, *S*-adenosylmethionine; SAH, *S*-adenosylhomocysteine; RRE, RiPP recognition element.

Given the frequency of YcaO occurrence in daptide BGCs,^21^ a sequence similarity network (SSN) for the YcaO protein family (Pfam ID: PF02624) was generated, and YcaOs occurring in daptide BGCs were annotated (**Figure S2**). These daptide-associated YcaOs were largely localized to a single group of the SSN, containing no previously characterized members. Daptide-associated YcaOs elsewhere in the SSN were annotated as azoline-forming YcaOs associated with thiopeptide biosynthesis (**Figure S3**). From the daptide-associated YcaO SSN group, we additionally selected the *Streptomyces azureus* ATCC 14921 BGC (hereafter, *saDap* BGC) for characterization, given the strain was already in our possession (**Figure 2, Tables S2-S5**). The *saDap* BGC encodes four precursor peptides (*Sa*DapA_1-4_), the central daptide proteins (*Sa*DapBCDMPT_1_T_2_), and a YcaO (*Sa*DapY), but it lacks the lanthipeptide synthetase and luciferase-like monooxygenase. Sequence analysis showed that *Tf*DapY and *Sa*DapY contain the canonical PxPxP sequence motif associated with azoline-forming YcaO cyclodehydratases^27,28^ (**Figure S4**); however, the only conserved residue among the precursor peptide core regions is the C-terminal Thr, which is putatively converted to Dmp in daptide biosynthesis.^21^ Additionally, no characterized YcaO enzymes have been shown to modify a C-terminal residue. The sequence divergence of *Tf*DapY and *Sa*DapY from characterized YcaO enzymes suggested they could expand diversity of daptide termini. Using these BGCs, we sought to fully reconstitute the daptide biosynthetic pathway *in vitro* and identify the function of the uncharacterized YcaO proteins.

### Requirements for aminoacetone biosynthesis

We first obtained His-tagged, *E. coli* codon-optimized *tfDapA*_*1*_*BCDMY* for expression in BL21(DE3) cells (**Dataset S2, Table S4-S7**). MBP-fusion constructs were also generated for *saDapA*_*1*_*BCDMY* (MBP, maltose-binding protein). While expression analysis showed that *saDap* proteins were of mixed quality, *tfDap* broadly gave high quality protein (**Figure S5**). This follows the trend of facile expression of biosynthetic enzymes from thermophilic organisms like *T. fusca*.^23–25^ MBP-fusion constructs were next prepared for *Sa*DapA1 and *Tf*DapA_1_ (**Table S3**). Full length *Sa*DapA_1_ and *Tf*DapA_1_ could both be produced, but significant proteolysis of the hydrophobic C-terminus was observed by MALDI-TOF-MS (matrix assisted laser desorption/ionization-time of flight-mass spectrometry; **Figure S6**). Therefore, solid phase peptide synthesis was leveraged to obtain *Tf*DapA_1_ for enzymatic reconstitution (**Table S8**).

Prior studies suggested that DapB and DapC proteins function as a two-component oxidative decarboxylase to generate the C-terminal aminoacetone (Aac).^21^ AlphaFold^29^ models further support *Tf*DapA_1_BC complex formation, and model analysis indicates that *Tf*DapB contains a C-terminal RRE (RiPP recognition element) domain,^30^ which putatively binds the leader region of *Tf*DapA_1_ (**Figure S7**). Actinobacterial DapC homologs (Pfam ID: PF00465) typically use NAD(P)^+^, and we predicted that *Tf*DapC would prefer NAD^+^ by the presence of Asp65 in a key binding loop.^31^ While members of PF00465 are additionally annotated as iron-dependent enzymes,^32^ the *Tf*DapC AlphaFold model showed substitutions to the canonical metal-coordinating residues of the family. The identities of these residues in *Tf*DapC suggested loss of metal coordination, and similar cases have been characterized in other PF00465 members that operate without metal-activation (**Figure S7**).^31^ To examine these predictions, we reacted *Tf*DapA_1_ with *Tf*DapBC and various co-substrates. LC-MS analysis demonstrated production of [M-46+3H]^3+^, corresponding to *Tf*DapA_1_-Aac (calc. *m/z*, 1665.8045; obs. *m/z*, 1665.8036; ppm error, −0.5). Aac formation was observed independent of metal supplementation, while replacement of NAD^+^ with NADP^+^ resulted in diminished turnover (**Figures 2, S8, S9**).

We next investigated *Tf*DapC dependence on *Tf*DapB. Analysis of reactions lacking *Tf*DapB showed no Aac production using *Tf*DapA_1_ as a substrate, and minimal Aac production when using (MBP)*Tf*DapA_1_ (**Figures S10, S11**). This suggested that optimal decarboxylase activity depended on both proteins, but *Tf*DapC was at least partially competent without *Tf*DapB. Analysis of the *Tf*DapA_1_BC AlphaFold model further suggested that Trp3 may engage *Tf*DapB during putative complex formation (**Figure S7**). To investigate this, we later generated *Tf*DapA_1_^(GS)6^, a solubility-optimized substrate with a shortened leader region and substitution of residues 5—16 by six GS repeats (*vide infra*). Following reaction with *Tf*DapBC, LC-MS analysis indicated production of the Aac product (**Figure S12**). We then synthesized the corresponding Trp3 variant, *Tf*DapA_1_^(GS)6,W3A^, which yielded minimal Aac product upon reaction with *Tf*DapBC. This supports the predicted role of Trp3 in substrate recognition for the oxidative decarboxylation step.

### Enzymatic generation of amino-modified C-termini

We subsequently focused on DapD reconstitution, which was predicted to convert Aac to 1,2-diaminopropane (Dap). *Tf*DapD is classified as a class-III PLP-dependent aminotransferase (Pfam ID: PF00202; PLP, pyridoxal phosphate), and activity of these proteins necessitates an external amine donor.^33^ To confirm activity, we reacted *Tf*DapA_1_BCD with NAD^+^ and *E. coli* BL21(DE3) lysate to provide amine donors. LC-MS analysis confirmed the formation of [M-45+3H]^3+^, corresponding to the *Tf*DapA_1_-Dap product (calc. *m/z*, 1666.1483; obs. *m/z*, 1666.1473; ppm error, −0.6; **Figure S13**). Having confirmed *Tf*DapD activity, we sought to identify the amine donor by screening reactions containing candidate amines. MS analyses demonstrated only L-Lys was a viable amine donor for transamination (**Figures 2, S14**). To elucidate which nitrogen was transferred to the peptide, we used [^15^N]-L-Lys selectively enriched at the α- or ε-amino positions. LC-MS analysis indicated ^15^N incorporation exclusively with [ε-^15^N]-L-Lys, demonstrating that the ε-amino group was transferred (**Figure S15**).

The final daptide biosynthetic step introduces the tertiary amine Dmp through two *N*-methylations catalyzed by DapM. To investigate this, we reacted *Tf*DapA_1_BCDM with NAD^+^, L-Lys, and *S*-adenosylmethionine (SAM) as the methyl donor. LC-MS analyses revealed a mixture of both doubly methylated Dmp (calc. *m/z*, 1675.4921; obs. *m/z*, 1675.4961; ppm error, 2.4) and a putative tri-methylated product (calc. *m/z*, 1680.1640; obs. *m/z*, 1680.1636; ppm error, −0.2; **Figures 2, S16**). We hypothesized that the trimethylated product may be an *in vitro* artifact, prompting us to evaluate *Sa*DapM activity. To generate *Sa*DapA_1_-Dap, we first co-expressed (MBP)*Sa*DapA_1_ with *Sa*DapBCD; however, MS analysis showed that the expression yielded only truncated products. Thus, we co-expressed (MBP)*Sa*DapA_1_ with *Tf*DapBCD instead, successfully yielding *Sa*DapA_1_-Dap (**Figure S17**). Reaction of *Sa*DapA_1_-Dap with purified *Sa*DapM yielded *Sa*DapA_1_-Dmp as the major product with no observable tri-methylated product (**Figure S18**).

### Reconstitution of daptide YcaO activity

Having characterized Dmp formation, we next reacted *Tf*DapY with *Tf*DapA_1_BCDM. The requisite co-substrates were supplied, as well as MgCl_2_ and ATP for the YcaO.^27^ LCMS analysis demonstrated a mixture of products, including [M-49+3H]^3+^ (**Figures 3, S19**); omission of *Tf*DapM from the reaction resulted in production of [M-63+3H]^3+^. Further omissions resulted in no new detectable analytes, indicating *Tf*DapY activity required Dap installation. Collectively, these data suggested generation of Miz (methylimidazoline; [M-63]; calc. *m/z*, 1660.1448; obs. *m/z*, 1660.1512; ppm error, 3.9) and Diz (dimethylimidazoline; [M-49]; calc. *m/z*, 1664.8167; obs. *m/z*, 1664.8188; ppm error, 1.3) by cyclodehydration of *Tf*DapA_1_-Dap with or without a single methylation. *Sa*DapA_1_-Dap was concurrently assayed, yielding only the Miz product when reacted with *Sa*DapY, and only the Diz product when reacted with *Sa*DapMY (**Figure S20**). MS/MS analyses further localized the mass changes to the C-terminus, including identification of a critical y_2_^+^ ion for *Sa*DapA_1_-Diz (**Figure S21, Tables S9-10**).

**Figure 3.**
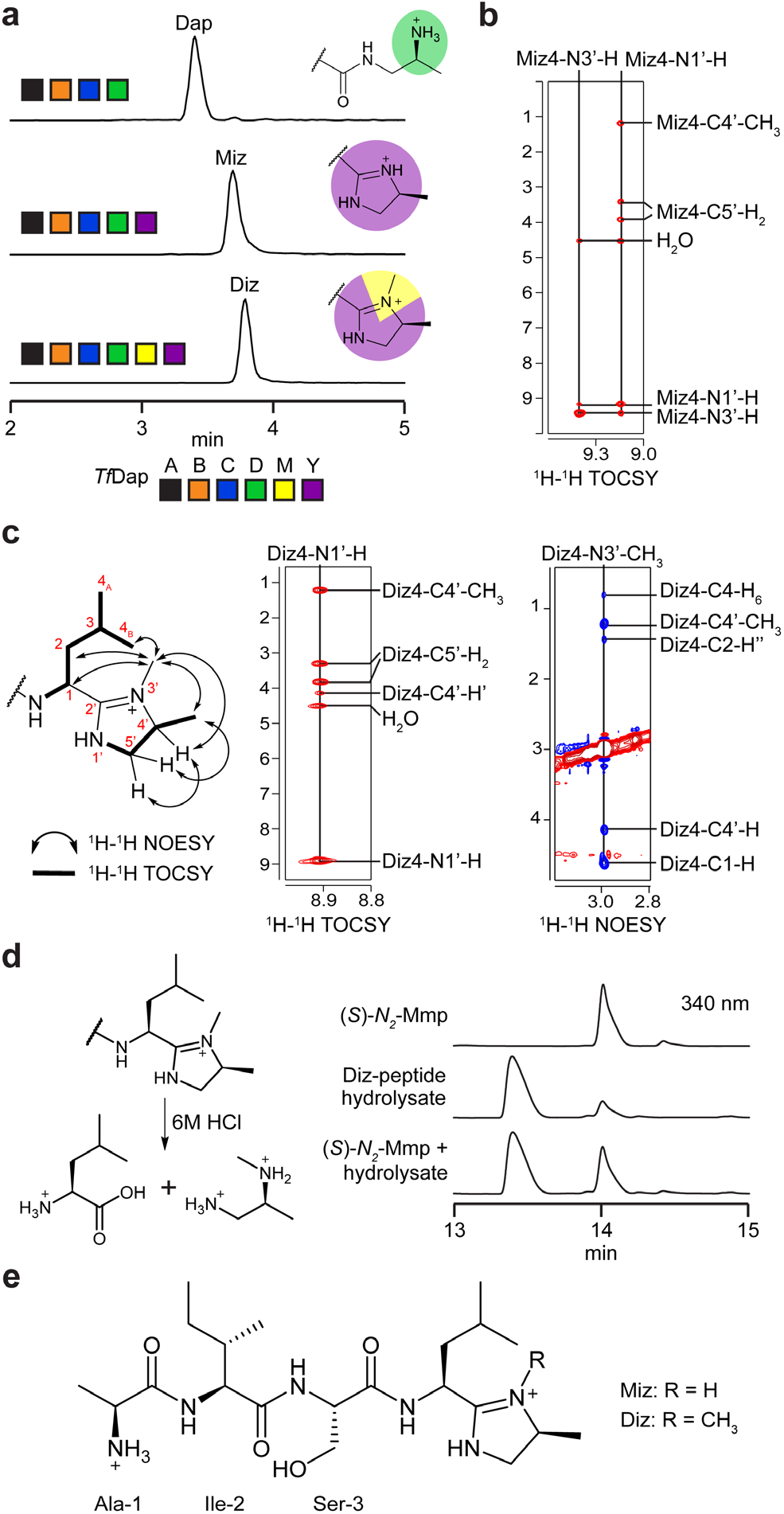
Characterization of DapY enzyme function. **a**, Ion-count normalized multiple ion chromatograms showing new products of *Tf*DapY. The five highest abundance masses in the expected isotopic envelope were extracted for each analyte. **b**, ^1^H-^1^H TOCSY correlations for amidine protons in the Miz spin system. **c**, ^1^H-^1^H TOCSY and ^1^H-^1^H NOESY correlations for assignment of Diz residue. **d**, Marfey’s analysis of Diz residue. Predicted hydrolysis products for Diz residue and HPLC chromatograms (340 nm) are shown. **e**, Full structure(s) for Miz- and Diz-modified peptide products of the daptide pathway. Abbreviations: Dap, 1,2-diaminopropane; Miz, 4-methylimidazoline; Diz, 3,4-dimethylimidazoline; Mmp, monomethyl-1,2-propanediamine.

### Structure elucidation of new daptide termini

To facilitate structure elucidation, we attempted to isolate smaller peptide fragments containing the modified termini. Single-site variants of *Sa*DapA_1_ were generated to introduce protease cleavage sites in the core region (**Figure S22**). While these variants still permitted installation of Diz, yields were poor owing to truncations. We next examined an engineered substrate, (MBP)DapA_eng_, to simplify purification (**Figure S23**). (MBP)DapA_eng_ contained shortened linker and leader regions with the central 12 amino acids of the core region replaced with a His_6_-tag and TEV protease site. Processing by TEV protease would then yield a pentapeptide ideal for structural characterization. Co-expression of (MBP)DapA_eng_ with *Tf*DapBCD resulted in a high titer of (MBP)DapA_eng_-Dap (75 mg protein/L culture). We reacted 100 mg of purified (MBP)DapA_eng_-Dap with *Tf*DapY overnight. After TEV protease cleavage and purification, fractions containing the Miz product were pooled (**Figure S24**). We repeated this process using both *Tf*DapY and *Sa*DapM *in vitro* to obtain the Diz product.

A series of NMR experiments (^1^H, ^1^H-^1^H COSY, ^1^H-^1^H TOCSY, ^1^H-^13^C HSQC, ^1^H-^1^H NOESY) were conducted on the putative Miz product (**Figure S25**). These spectral analyses assigned the modified Leu-Thr terminus as a 4-methylimidazoline (Miz, formally 1-amino-1-[4-methyl-2-imidazolin-2-yl]isopentane; **Table S11**). Characteristic TOCSY correlations permitted assignment of the Ala, Ile, and Ser residues (**Figure S26**). The aminoisopentane group displayed characteristic TOCSY/COSY correlations for Leu with amide and C1-H proton signals shifted further downfield than expected, suggesting modification. Two additional protons were observed between 9.0 and 9.5 ppm, which we assigned as amidine N-H protons (**Figures 3, S27**). The TOCSY data suggested these two protons were coupled as part of a unique spin system. We assigned the remaining signals of the spin system to the C5’-H’, C5’-H’’, C4’-H, and C4’-CH3 protons of the imidazoline by TOCSY and HSQC (**Figure S27**). Imidazoline formation also accounts for the observed loss of 63 Da and MS/MS fragmentation.

We next performed NMR experiments (^1^H, ^1^H-^1^H DQFCOSY, ^1^H-^1^H TOCSY, ^1^H-^13^C HSQC, ^1^H-^1^H NOESY) on the putative Diz product (**Figure S28, Table S12**). As before, TOCSY correlations readily assigned the Ala, Ile, and Ser residues along with the isopentane group (**Figure S29**). TOCSY correlations further supported the presence of an imidazoline (**Figures 3, S30**). In contrast to the Miz product, we observed the loss of one of the amidine protons and the gain of a new methyl group, suggesting a *C,N*-dimethylimidazoline. NOESY correlations were identified from the new methyl group to (i) C1-H, C2-Hα, and C4-H_6_ of the aminoisopentane group and (ii) C4’-H and C4’-CH3 of the imidazoline (**Figures 3, S31**). Correlations were not observed from the new methyl group to either C5’-H’ or C5’-H’’; however, NOEs were observed between the C5’ protons and the C4’-CH3 and C4’-H protons. These data support the formation of Diz with methylation of nitrogen N3’ (**Figures 3, S32**).

Marfey’s method was used to determine the stereochemistry of Diz. We anticipated that strong acid would catalyze hydrolysis at the 2’-position of the imidazoline to yield *N*-methylpropane-1,2-diamine (Mmp, monomethyl-1,2-propanediamine) and Leu (**Figure 3**). Following acid hydrolysis and derivatization with Marfey’s reagent. LC-MS retention time comparisons to amino acid standards allowed assignment of Ala, Ile, Ser, and Leu to their proteinogenic Lenantiomers (**Figure S33**). Standards for all four isomers of Mmp were then synthesized and derivatized with Marfey’s reagent. Comparison of retention times for the synthesized standards to the derivatized hydrolysate identified (*S*)-*N*_*2*_-Mmp in the hydrolysate (**Figures 3, S34**). Based on the expected route of hydrolysis, this suggested that the Leu-Thr terminus was converted to (*1S*)-1-amino-1-[(*4S*)-3,4-dime-thyl-2-imidazolin-2-yl]isopentane (Diz, hereafter 3,4-dimethylimidazoline).

### Biosynthetic bifurcation and order of events

Having reconstituted DapBCDMY, we next investigated the biosynthetic order of events. Given the identified PTMs, there remained multiple possible biosynthetic routes from Dap (**Figure S35**). Thus far, we knew that both DapY and DapM were required for Diz formation and that each enzyme could modify Dap individually. However, it remained unclear whether Diz formed from the cyclized Miz or through the singly methylated Mmp. To investigate this, we reacted *Tf*DapA_1_-Dap with *Tf*DapY and subsequently added *Tf*DapM to the reaction. LC-MS analysis showed production of *Tf*DapA_1_-Miz; *Tf*DapA_1_-Diz was not observed (**Figure S36**). We next reacted *Sa*DapA_1_-Dap with *Sa*DapY, followed by addition of *Sa*DapM. Analysis again only supported *Sa*DapA_1_-Miz formation (**Figure S37**). In both cases, these data show that DapM enzymes are unable to convert Miz to Diz; therefore, Diz formation must proceed through Mmp.

We next explored whether these *in vitro* data were recapitulated through *in vivo* studies. To begin, we purified genomic DNA from *S. azureus* and used the CAPTURE method to clone the *saDap* BGC into an integrative vector for heterologous overexpression (**Tables S6-7**).^21,34,35^ Following introduction of the plasmid into a *Streptomyces* host by interspecies conjugation, we extracted exconjugant colonies by methanol. MALDI-TOF-MS analysis of the cell extracts revealed production of new analytes corresponding to the four encoded precursor peptides (**Figures 4, S38**). Previously characterized daptides were proteolytically cleaved after the ELExxxxx motif,^21^ and each detected product was also cleaved following the same motif, despite variation in the P1 and P1’ sites (**Figure S39**). We propose naming these compounds azuritides 1-4, corresponding to the major product of each encoded precursor peptide.

**Figure 4.**
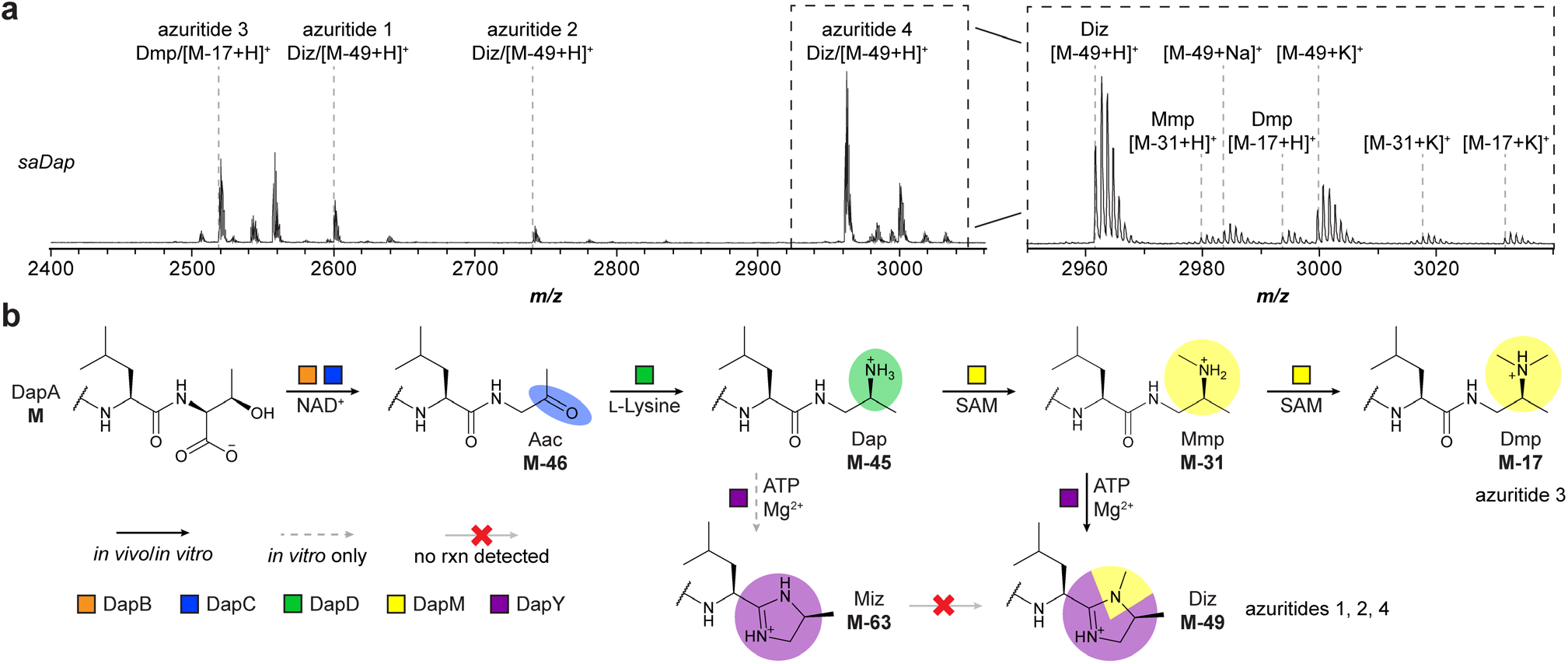
Biosynthetic route for YcaO-containing daptide BGCs. **a**, MALDI-TOF-MS for heterologous expression of *saDap* BGC in *S. albus* J1074. Relevant masses are listed as follows: azuritide 1 (Diz/[M-49+H]^+^), observed: 2600 Da, expected: 2600 Da; azuritide 2 (Diz/[M-49+H]^+^), observed: 2742 Da, expected: 2741 Da; azuritide 3 (Dmp/[M-17+H]^+^), observed: 2519 Da, expected: 2519 Da; azuritide 4 (Diz/[M-49+H]^+^), observed: 2962 Da, expected: 2962 Da. **b**, Updated scheme for daptide biosynthesis. Abbreviations: Aac, aminoacetone; Dap, 1,2-diaminopropane; Mmp, monomethylpropane-1,2-diamine; Miz, 4-methylimidazoline; Dmp, dimethylpropane-1,2-diamine; Diz, 3,4-dimethylimidazoline.

Assignment of the identified metabolites revealed that the C-termini of azuritides 1, 2, and 4 (corresponding to *Sa*DapA_1_, *Sa*DapA_2_, and *Sa*DapA_4_) were converted to the 3,4-dimethylimidazoline Diz, while the C-terminus of azuritide 3 (corresponding to *Sa*DapA_3_) was converted to the tertiary amine Dmp. We further identified secondary amine Mmp and tertiary amine Dmp products for *Sa*DapA_4_, while we did not observe production of the 4-methylimidazoline Miz for any of the encoded precursor peptides. These data suggest daptide biosynthesis proceeds to the Mmp intermediate, at which point the substrate can either undergo cyclodehydration to Diz (via DapY) or can undergo a second methylation (via DapM) to Dmp (**Figure 4**).

### Substrate tolerance of the *Tf*Dap enzymes

Thus far, investigation of the daptide pathway had yielded a route to modifying C-termini with a diverse set of functional groups. Daptide enzymes showed high tolerance for variation, with all enzymes tolerating minor substitutions in the core region. Major substitutions, such as DapA_eng_ and non-cognate precursor peptides were also tolerated (**Table 1**). We thus chose to “stress-test” the daptide enzymes using a panel of substrates and conditions.

**Table 1.**
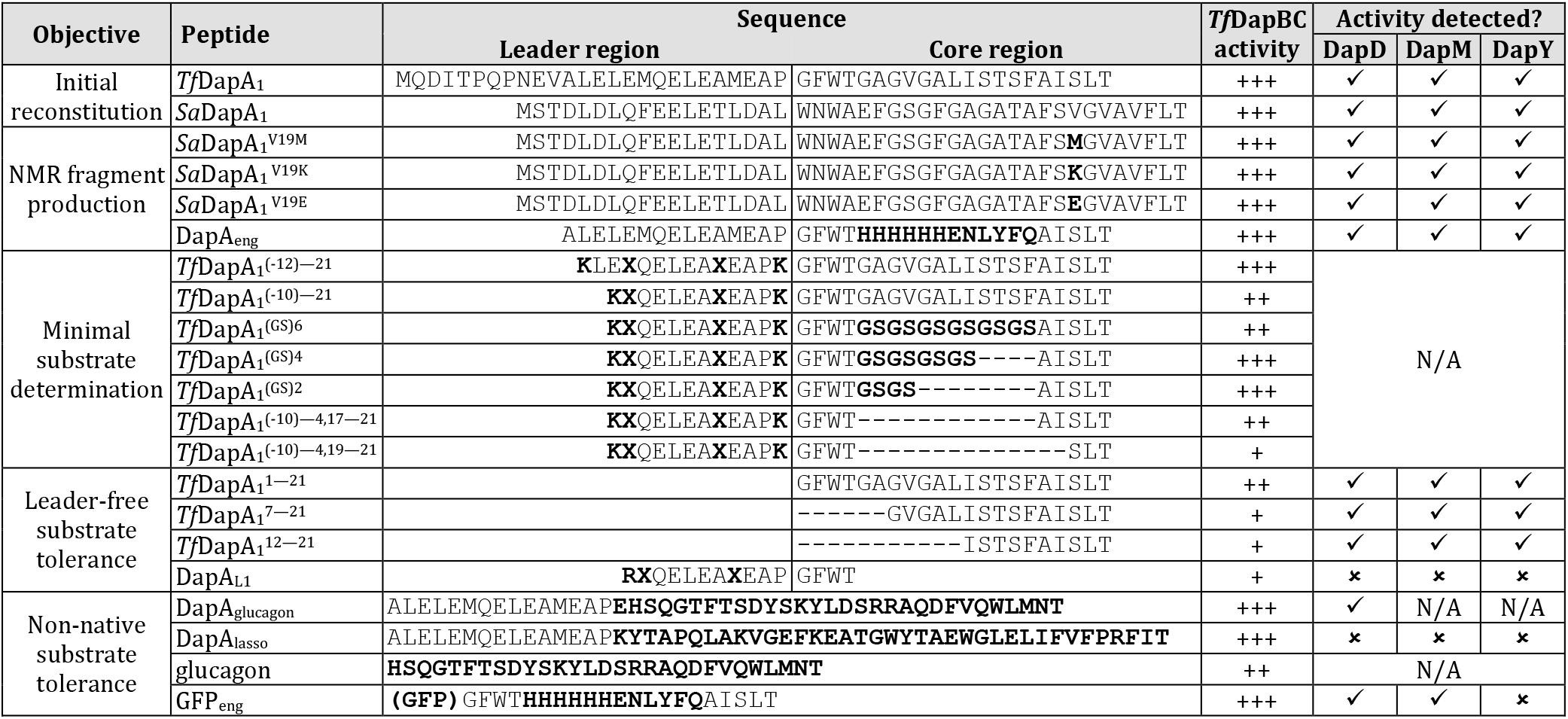
Summary of substrate scope data for daptide biosynthetic enzymes. Peptide sequences are split into leader region and core region where appropriate. Bold indicates substitutions from native daptide precursor peptide sequences. X = norleucine. *Tf*DapBC activity: +++, complete or nearly complete product formation; ++, moderate product formation; +, low but detectable product formation. DapDMY activity: ✓, Activity detected; ✗, No activity detected; N/A, Not assessed.

We first generated a series of leader region truncations of *Tf*DapA_1_. Analysis of *in vitro* reactions with *Tf*DapBC showed that removal of residues (−26)—(−13) from the leader region did not impede turnover, while removal of residues (−26)—(−11) slightly diminished turnover. (**Table 1, Figure S40**). Using this data, we generated an optimized substrate consisting of TfDapA_1_^(−10)—21^ with core residues 5—16 replaced by a (GS)_6_ repeat sequence. Notably, this peptide variant displayed a marked increase in solubility. After reaction with *Tf*DapBC, LC-MS analysis showed Aac production, consistent with previous findings showing DapBC tolerance to substitutions (**Table 1, Figure S12**). We subsequently synthesized a series of shortened substrates by removal of amino acids from the middle of the core region. Assays with *Tf*DapBC formed Aac in each case; however, the shortest substrate yielded only trace amounts of product (**Figure S41**). These results demonstrate that *Tf*DapBC can modify substantially truncated substrates.

We next determined whether the daptide enzymes could modify substrates lacking the leader peptide entirely. We first attempted a ConFusion strategy,^36^ which removes the leader region from the substrate and attaches it to a biosynthetic enzyme. We generated a chimeric expression construct for *Tf*DapB_conf_, encoding *Tf*DapB and *Tf*DapA_1_^(−26)—5^ inframe. We then generated three MBP-fusion constructs of the core peptide: *Tf*DapA_1_^1—21^, *Tf*DapA_1_^7—21^, and *Tf*DapA_1_^12— 21^. An *in vitro* reaction using *Tf*DapB_conf_CD resulted in partial conversion of all three substrates to mixtures of Aac and Dap (**Table 1, Figure S42**). As with the core truncation experiments, the length of each substrate correlated with the extent of modification, as *Tf*DapA_1_^1—21^ received the highest level while *Tf*DapA_1_^12—21^ received the least.

We further tested whether addition of the leader peptide *in trans* (i.e., as a separate peptide from the core peptide) would permit enzymatic modification.37 We obtained three variants of the *Tf*DapA_1_ leader region: DapA_L1_, DapA_L2_, and DapA_L3_ (**Table 1, Figure S43**). When assayed with the same three core variants as above, we detected Diz formation, showing that each daptide enzyme can modify the intended substrate with leader peptide supplied *in trans* (**Figure S43**). Notably, DapA_L1_ coincidentally contains a C-terminal Thr. Analysis of reactions containing *Tf*DapBC revealed decarboxylation of the DapA_L1_ peptide itself, providing an even shorter 15-mer minimal substrate (**Figure S43**).

In addition to examining substrate tolerance, we aimed to test the robustness of the daptide enzymes under different reaction conditions. Enzyme activity was retained after multiple freeze-thaw cycles, and reactions proceeded at neutral or basic pH, although buffer composition did affect product distribution (**Figure S44**). We next examined tolerance to desiccation stress by preparing a reaction containing *Tf*DapBCDMY and all required cofactors, but lacking substrate.^38,39^ We froze and lyophilized the reaction, followed by reconstitution in water and addition of DapA_eng_. Analysis by MALDI-TOF-MS detected only the final Diz product (**Figure S45**). Taken together, *Tf*DapBCDMY can tolerate a wide range of reaction conditions and modify diverse substrates.

### C-terminal functionalization of non-native substrates

Having investigated the substrate scope for the daptide enzymes, we next gauged the tolerance of the daptide enzymes towards non-native substrates. As C-terminal modification occurs frequently in peptide hormone maturation, we examined peptide hormone sequences for those naturally ending in Thr and identified glucagon as a candidate. We thus constructed a vector for expression of *Tf*DapBCD with (MBP)DapA_glucagon_, which replaced the *Tf*DapA_1_ core region with the native glucagon sequence. Following expression, (MBP)DapA_glucagon_ was purified and digested individually by LysC protease and GluC protease. MS analyses confirmed the conversion of full-length glucagon to glucagon-Dap (**Table 1, Figures 5, S46, S47**).

**Figure 5.**
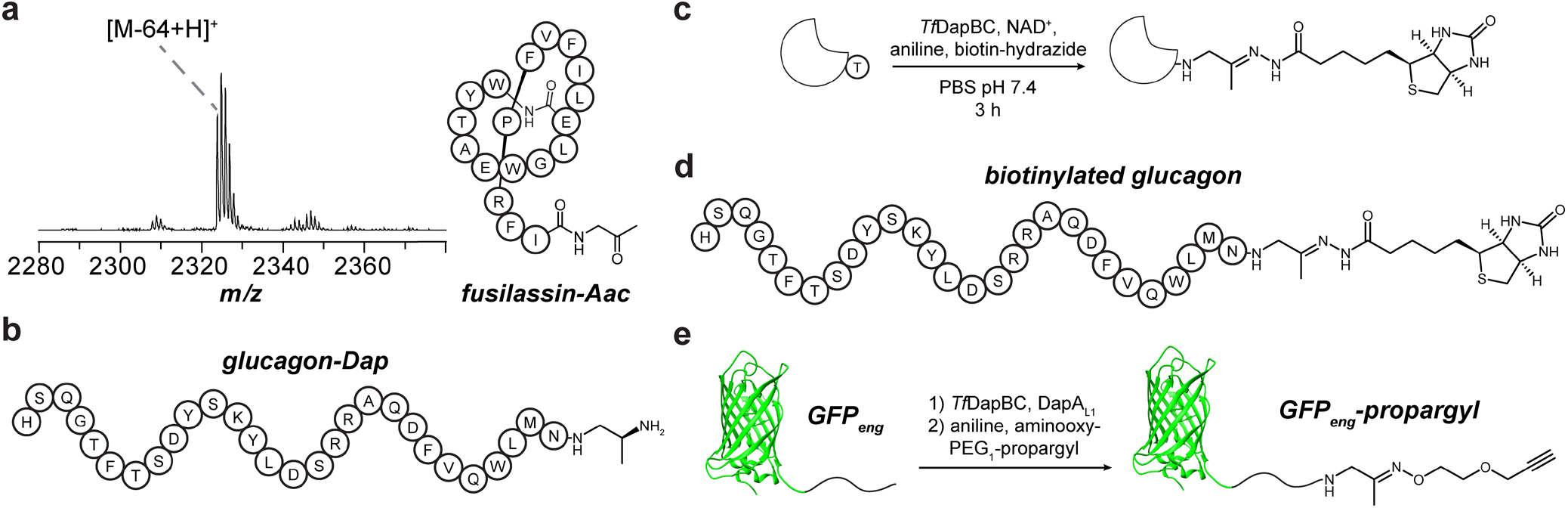
Aminoacetone installation on non-native substrates. **a**, MALDI-TOF-MS and bubble diagram of fusilassin-Aac. Expected *m/z*, 2324; observed *m/z*, 2324. Abbreviation: Aac, aminoacetone. **b**, Scheme for one-pot oxidative decarboxylation and biotinylation of substrates. **c**, Biotinylated glucagon generated using *Tf*DapB_conf_C. **d**, Bioconjugation reaction of GFP_eng_ to give the oxime product, GFP_eng_-propargyl.

Beyond peptide hormones, modification of macrocyclic peptides could enable new molecular diversity. In addition to *tfDap, T. fusca* encodes the BGC for fusilassin (*i*.*e*., fuscanodin), which has served as a scaffold for lasso peptide engineering.^40,41^ We examined whether the two *T. fusca* pathways could be combined to yield macrocyclic peptides with enzymatically modified C-termini. We generated a construct for (MBP)DapA_lasso_, which replaced *Tf*DapA_1_^1—20^ with residues (−18)—18 of the fusilassin precursor peptide (**Table 1, Figure S48**).^42^ Following expression, the peptide was treated with *Tf*DapBCDY, FusBCE (FusB, lasso leader peptidase; FusC, lasso cyclase; FusE, RRE), or both sets of enzymes in succession. We first showed that FusBCE can modify (MBP)DapA_lasso_ to produce a fusilassin variant with C-terminal Thr. *Tf*DapBC additionally showed full conversion of the linear peptide to the Aac product; however, *Tf*DapD did not detectably modify the chimeric substrate further (**Table 1, Figure S48**). Addition of FusBCE resulted in conversion of the Aac-modified peptide to the corresponding macrocyclic lasso peptide, fusilassin-Aac (**Figure 5**).

### Bioconjugation with aminoacetone-modified C-termini

One potential application for the daptide enzymes is the use of the Aac intermediate as a reactive handle for bioconjugation. We first examined whether *Tf*DapBC could be used in a one-pot bioconjugation reaction by testing reactions of (MBP)DapA_eng_, *Tf*DapBC, NAD^+^, aniline, and biotin-hydrazide (**Figure S49**).^43^ Following removal of small molecules and TEV proteolysis, the reactions were analyzed by LC-MS, demonstrating biotinylation of (MBP)DapA_eng_ at its C-terminus (**Figures 5, S49**). Given the substrate tolerance of *Tf*DapBC, we hypothesized that Aac installation would enable bioconjugation on the C-terminus of polypeptides while allowing variation in the residues preceding the C-terminal Thr. We thus treated glucagon with (i) *Tf*DapBC, (ii) *Tf*Dap-B_conf_C, or (iii) DapA_L1_ and *Tf*DapBC. Analysis by MALDI-TOF-MS confirmed glucagon-Aac product in the ConFusion (*Tf*DapB_conf_C) and *in trans* (DapA_L1_ and *Tf*DapBC) reactions, but not with *Tf*DapBC alone (**Figure S50**). We further added biotin-hydrazide and analyzed the products (**Figure S51**). Appearance of the [M-46+240+H]^+^ ion indicated production of biotinylated glucagon (**Figure 5**).

Our next goal was modification of protein substrates using the leader-free approach. We expressed and purified GFP_eng_, containing GFP and the core region of DapA_eng_ at its C-terminus (GFP, green fluorescent protein). GFP_eng_ was then reacted with *Tf*DapBCDMY and DapA_L1_ before digestion by GluC. MALDI-TOF-MS analyses demonstrated conversion to a mixture of GFP_eng_-Mmp and GFP_eng_-Dmp (**Table 1, Figure S52**). We further reacted GFP_eng_ with *Tf*DapBC and DapA_L1_ followed by addition of aniline and aminooxy-PEG_1_-propargyl. MALDI-TOF-MS analysis of the GluC proteolytic digest revealed near-complete conversion to the oxime product, GFP_eng_-propargyl (**Figures 5, S52**). These data show that *Tf*DapBC can generate a minimal Aac reactive handle onto non-native peptide and protein substrates. The examined conditions are mild and allow one-pot bioconjugation by site-selective C-terminal functionalization of diverse substrates.

## Discussion

We previously reported the genome-guided discovery of daptides, peptides bearing a tertiary amine in place of the C-terminal carboxylate.^21^ In this study, we reconstituted daptide biosynthesis *in vitro*, elucidating the required cofactors for this unique modification. By assessing the genomic context of daptide biosynthesis, we identified BGCs containing divergent YcaO enzymes expected to further elaborate daptide products. We showed that these DapY enzymes intercept a secondary amine intermediate and divert daptide biosynthesis away from the tertiary amine to a C-terminal 3,4-dimethylimidazoline (Diz) moiety.

Heterologous expression of the *saDap* BGC showed that the YcaO enzyme acts on only three of four precursor peptides. No obvious sequence trend was observed when comparing the precursor peptides to account for the difference in modification endpoint. Elucidation of the rules dictating this branch point in daptide metabolism will require additional investigation, and we suspect there are substrate-directed influences on binding or reaction kinetics that govern this behavior. Enzymatic competition for a particular amino acid is known in other RiPP pathways, such as with Ser in thiopeptide biosynthesis which can undergo oxazoline or dehydroalanine formation.^44^ Despite this, the determinants of these biosynthetic branches are not clear.

Amines have strong precedent as nucleophiles used by YcaO enzymes, including diaminopropionic acid-containing substrates installed by Flexizyme,^45^ macrolactamidine formation with the N-terminus,^46^ and formation of backbone amidines using exogenous ammonia.^47^ YcaO enzymes are known to use ribosomally installed nucleophiles and exogenous nucleophiles, but Diz biosynthesis represents an unconventional case of YcaO-catalyzed backbone modification using three successive PTMs to install the critical secondary amine nucleophile. YcaO enzymes have also now been shown to act at internal peptide bonds and at both termini. In other RiPPs with YcaO-catalyzed C-terminal heterocycles, cyclization precedes removal of a follower peptide to generate the terminal heterocycle, as in bottromycin^46^ and spyrimidone.^48^ Daptide biosynthesis is the first case to directly modify a C-terminal residue (**Figure S53**). The observation that this occurs after decarboxylation suggests that the C-terminal carboxylate of YcaO may be catalytic, as decarboxylation of the daptide substrate may alleviate charge-charge repulsion to facilitate cyclization.^49^

The Diz residue is ultimately generated through the actions of five biosynthetic proteins. This biosynthetic strategy modifies the Leu-Thr terminus to generate a nitrogenous heterocycle, mimicking alkaloid biosynthesis. To date, imidazolines have not been observed in RiPP natural products, but they are biosynthetically precedented; including alkaloids (*e*.*g*., spongotines,^50^ tulongicins,^51^ and penipanoid B^52^) and the siderophore, pseudochelin A (**Figure S54**).^53^ Perhaps the most well-known imidazoline compounds are the nutlins,^54^ anti-cancer compounds which inhibit the interaction of p53 and MDM2. A family of GPCRs, imidazoline receptors 1-3, have also been shown to bind imidazoline-containing molecules, such as clonidine.^55,56^ Imidazolines are further well-established head groups for detergents and surfactants,^57^ as linkage of the imidazoline to lipid chains generates a fatty imidazoline with cationic surfactant properties. These molecules are used as acid-stable detergents and anti-corrosion compounds. Given the propensity for daptides to interact with membranes, we speculate that Diz formation may allow the daptides to act in a similar surfactant role.

Although the functional role of daptides remains unclear, we have made significant progress in studying the enzymatic requirements for their production. Exploration of daptide substrate scope demonstrated C-terminal modifications of variant substrate core peptides. A common feature of RiPP biosynthesis is the separation of leader peptide recognition from catalysis, and we exploited this to modify leader-free, non-native, and protein substrates. In doing so, we installed new C-termini, including the reactive Aac moiety, on a hybrid macrocyclic lasso peptide, glucagon, and GFP. The broad tolerance of daptide biosynthesis for both non-native sequences and diverse reaction conditions offers a new tool for custom peptide engineering and protein bioconjugation applications. As the only apparent strict substrate requirement is a C-terminal Thr, daptide enzymes may be especially useful in cases where minimal modification of the substrate or mild reaction conditions are required. RiPP enzymes are increasingly being employed in various peptide and protein engineering projects.^58–64^ We expect that discoveries in natural product biosynthesis will continue to reveal promising catalysts for biotechnological development.

## ASSOCIATED CONTENT

## Supporting Information

Methods, Tables S1-S12, and Figures S1-S54 showing additional bioinformatic and spectral data.

Dataset S1: NCBI identifiers from bioinformatic analysis. Dataset S2: Plasmid sequences generated in this study.

## AUTHOR INFORMATION

## Author Contributions

J.R.C., and D.A.M. designed the research. S.R.D. performed bioinformatics, *in vitro* reconstitution, structure elucidation, substrate scope analysis, and bioconjugation experiments. S.K.K. performed solid phase peptide synthesis, *in vitro* reconstitution, co-substrate determination, and substrate scope analysis. H.R. generated the integrative vector for expression of *saDap*, with oversight from H.Z. D.P.L. collected NMR data for modified peptides.

S.F. and D.S. designed and performed synthesis of chemical standards. S.R.D. wrote the first draft of the manuscript. S.R.D., S.K.K., J.R.C., and D.A.M. wrote the manuscript with editorial input from the remaining authors.

## Notes

The authors declare the following competing financial interest(s): S.R.D. and D.A.M. are inventors on a provisional patent filed by Vanderbilt University pertaining to some aspects of this work. The other authors declare no competing financial interests.

## ACKNOWLEDGMENT

Access to instruments was provided by the following centers: Mass Spectrometry Research Center, Vanderbilt University; School of Chemical Sciences Mass Spectrometry Lab, University of Illinois at Urbana-Champaign; School of Chemical Sciences NMR Lab, University of Illinois at Urbana-Champaign; Roy J. Carver Biotechnology Center Proteomics Core, University of Illinois at Urbana-Champaign; Triad Mass Spectrometry Facility, University of North Carolina at Greensboro. Direct infusion HR-MS data was collected by Sangeetha Ramesh, Alex Battiste, and Austin Woodard. We thank Riley Carter, Dinh Nguyen, Hamada Saad, Yanqing Xue, Sharon Roth, and Chandrashekhar Padhi for helpful discussion.

## ABBREVIATIONS

RiPP: ribosomally synthesized and post-translationally modified peptide
PTM: post-translational modification
BGC: biosynthetic gene cluster
Aac: aminoacetone
Dap: 1,2-diaminopropane
Mmp: monomethylpropane-1,2-diamine
Dmp: dimethylpropane-1,2-diamine
Miz: 4-methylimidazoline
Diz: 3,4-dimethylimidazoline
GFP: green fluorescent protein
MBP: maltose-binding protein
SSN: sequence similarity network
MALDI-TOF-MS: matrix assisted laser desorption/ionization-time of flight-mass spectrometry
RRE: RiPP precursor peptide recognition element
LC-MS: liquid chromatography-mass spectrometry
NAD: nicotinamide adenine dinucleotide
PLP: pyridoxal phosphate
SAM: *S*-adenosylmethionine
ATP: adenosine triphosphate
TEV: tobacco etch virus
CAPTURE: Cas12a-assisted precise targeted cloning using *in vivo* Cre-Lox recombination
GPCR: G-protein coupled receptor.

## TOC figure

Synopsis (200 character limit): Daptide biosynthetic enzymes convert C-termini to aminoacetone, diaminopropane, dimethylimidazoline, etc. and can install these modifications onto a broad range of substrates.

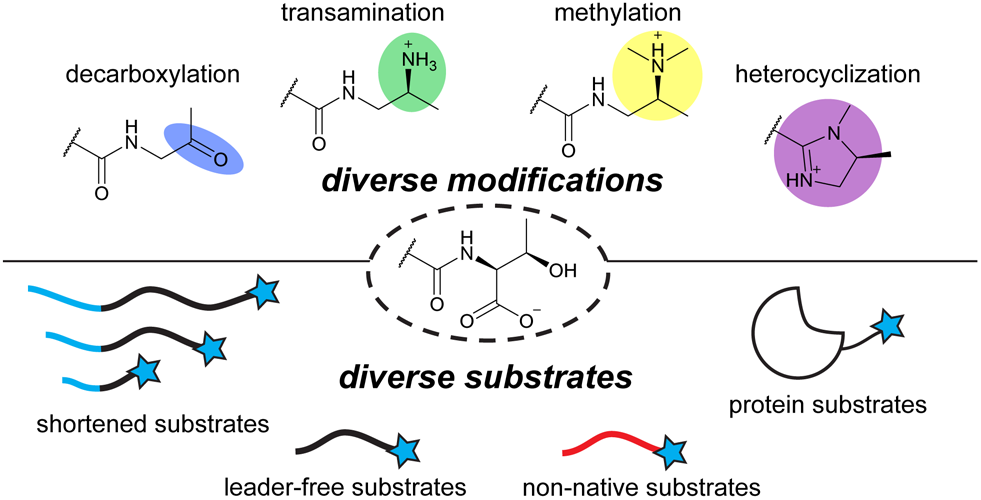

