## Supporting Information for "A versatile enzymatic pathway for modification of peptide C-termini"

### Materials and Methods

**Materials.** All chemicals were used without further purification unless stated otherwise. All antibiotics were purchased from ChemImpex or Goldbio. Antibiotics were used at the following concentrations in both solid and liquid cultures: kanamycin (Kan), 50 μg/mL; ampicillin (Amp), 100 μg/mL, chloramphenicol (Cam), 37 μg/mL. All DNA oligos were purchased from Integrated DNA Technologies, Inc. unless otherwise indicated. Enzymes were purchased from New England Biolabs (NEB) unless otherwise indicated. Sources for experiment-specific materials and reagents are provided in the relevant sections below.

**Gathering and annotation of daptide BGCs.** Identified DapB homologs from Ren et al.[1] were selected for regathering the set of daptide BGCs. A combination of BLASTP and Position Specific Iterative-BLAST (PSI-BLAST) searches were performed on the NCBI BLAST webtool (https://blast.ncbi.nlm.nih.gov/Blast.cgi).[2] Individual BLASTP results (max. 1,000 hits/protein query) and 2-round PSI-BLAST results (max. 3,000 hits/protein query) were gathered, and duplicate protein accession IDs were removed. The combined list of protein accessions was submitted to the RODEO webtool (https://webtool.ripp.rodeo/) for BGC analysis.[2] Results files were downloaded, and BGCs were assigned if a single locus contained hits for daptide-associated Pfams annotating aminotransferases, alcohol dehydrogenases, and methyltransferases. A final set of BGCs was generated containing 1,034 daptide BGCs. Short ORFs encoded in these BGCs were annotated as precursor peptides based on the presence of a C-terminal Thr and similarity to previously reported datasets. In total, 3,013 precursor peptides were identified (**Dataset S1**).

**Sequence similarity network analysis.** The sequence similarity network (SSN) was generated using the EFI-EST webpage (https://efi.igb.illinois.edu/efi-est/).[3] The YcaO Pfam (Pfam ID: PF02624)[4] was used as a "family" query to generate the SSN from all YcaO proteins identified in the UniProtKB database. Following initial edge calculation, the threshold for edge inclusion was set to alignment score 40 for generation of the SSN. For analysis, a RepNode 0.90 network was downloaded and opened in Cytoscape 3.7.3. The alignment score was further adjusted to 50 by removal of edges. The network layout was generated by the yFiles organic layout algorithm Cytoscape add-in. Annotation of daptide-associated YcaOs was performed by first gathering all YcaO proteins encoded in daptide BGCs as identified by RODEO analysis. The list of representative YcaO IDs in the RepNode 0.90 network were then mapped from UniProtKB AC/ID to EMBL/GenBank/DDBJ CDS and to RefSeq Protein using the Uniprot ID mapping tool (https://www.uniprot.org/id-mapping). The list of resulting IDs was used to search the RODEO-identified dataset of YcaO proteins, and hits were annotated on the corresponding node in the SSN.

**Protein modeling with AlphaFold 3.** AlphaFold 3[7] was accessed through the Alphafold server website (https://alphafoldserver.com/). Images of AlphaFold predictions were generated using Chimera (version 1.15).

**General molecular biology methods.** PCRs were performed using Q5 Polymerase (NEB) following manufacturer-recommended protocols. Where appropriate, DpnI (NEB) was added to PCR products to digest template plasmid. PCR products were then purified by silica spin-column and used for downstream cloning. Where necessary, sticky ends were generated using BamHI (NEB) and HindIII (NEB) restriction enzymes using manufacturer-recommended Blunt end ligation was performed at 10 uL scale using 1 uL each of DpnI, T4 Polynucleotide kinase (NEB), and T4 DNA Ligase (NEB) in T4 DNA ligase buffer. Gibson assembly reactions were performed using 2x HiFi DNA Assembly Master Mix (NEB) following manufacturer-recommended protocols. Blunt end ligation reactions, sticky end ligation reactions, Gibson assembly reactions, and Quikchange (Agilent) PCR products were all transformed into chemically competent *E. coli* DH5α or *E. coli* DH10β, and clones were selected by plating on LB-Miller medium (10 g/L tryptone, 5 g/L yeast extract, 10 g/L NaCl) supplemented with agar (20 g/L) and the appropriate antibiotic. Individual colonies were propagated in 5 mL LB-Miller medium containing the appropriate antibiotic, and plasmids were subsequently purified by alkaline lysis methods. Sequences were confirmed by Oxford nanopore sequencing (Plasmidsaurus) or Sanger sequencing (Roy J. Carver Biotechnology Center Sanger Sequencing Core, University of Illinois at Urbana-Champaign). Generated plasmids are provided in **Dataset S2**.

**Construction and mutagenesis of expression vectors for *saDap* genes.** The *saDapA₁*, *saDapC*, *saDapD*, *saDapY*, and *saDapM* genes were amplified by PCR from purified *Streptomyces azureus* ATCC 14921 genomic DNA. Restriction sites were incorporated by addition as 5' primer overhangs. Both pET-28-(MBP) empty vector and individual PCR amplicons were digested to make stick ends and then ligated by T4 DNA ligase (NEB).

**Construction of (MBP)DapA$_1$-DapBCD co-expression vectors.** Genes for *saDapA$_1$*, *saDapB*, *saDapC*, and *saDapD* were synthesized as *E. coli* codon-optimized sequences (Twist Biosciences) inserted into helper plasmids.[5] The *saDapB*, *saDapC*, and *saDapD* codon-optimized genes were amplified by PCR, with homology arms added by 5' primer overhangs. A pETDuet-(MBP)empty-empty vector was amplified by PCR, and the *saDapB*, *saDapC*, and *saDapD* genes were inserted into the second multiple cloning site by Gibson assembly to generate pETDuet-(MBP)empty-*saDapBCD*. Both the codon-optimized *saDapA$_1$* and pETDuet-empty-*saDapBCD* were then amplified by PCR with homology arms added by 5' primer overhangs. These amplicons were then assembled by Gibson assembly to give pETDuet-(MBP)*saDapA$_1$-saDapBCD*. The same protocol was repeated using primers designed for *tfDapB*, *tfDapC*, and *tfDapD* codon-optimized genes to give pETDuet-(MBP)*saDapA$_1$-tfDapBCD*. The plasmid pETDuet-(MBP)*tfDapA$_1$-tfDapBCD* was further generated by Gibson assembly using *tfDapA$_1$* gene as a template for insert generation by PCR and pETDuet-(MBP)*saDapA$_1$-tfDapBCD* as a template for backbone generation by PCR. *Sa*DapA$_1$ variants were generated by site-directed mutagenesis with the Quikchange protocol. Primers containing variant codons were used to generate the mutagenized linear DNA by PCR.

**Construction of pETDuet-(MBP)*dapA$_{eng}$-tfDapBCD*, pETDuet-(MBP)*dapA$_{eng}$*-empty and pETDuet-(MBP)*dapA$_{glucagon}$-tfDapBCD* plasmids.** To generate the DapA$_{eng}$ variant, we obtained three sets of overlap extension primers. The pETDuet-(MBP)*tfDapA$_1$-tfDapBCD* vector was used as template for the first round of overlap extension by a 5-cycle PCR. This reaction was subsequently used as the template for a second round of overlap extension by 5-cycle PCR. Finally, this reaction was used as template in a third round of overlap extension by 25-cycle PCR. The products of this reaction were assembled by Gibson assembly. The same strategy was applied using alternate primers to generate pETDuet-(MBP)*dapA$_{glucagon}$-tfDapBCD*. Using pETDuet-(MBP)*dapA$_{eng}$-tfDapBCD* as a template, we performed PCR to obtain a linearized version of pETDuet-(MBP)*dapA$_{eng}$*-empty. The plasmid was assembled by blunt end ligation.

**Construction of plasmids for leader-free substrate testing.** The pET28-*tfDapB* vector and a portion of $tfDapA_1$ corresponding to residues (-26)—5 were individually amplified by PCR. Homology arms, including a $(GS)_3$ linker, were inserted by 5' primer overhangs. The plasmid was then generated by Gibson assembly. Generation of pET-28a-(MBP)$tfDapA_1^{1-21}$, pET-28a-(MBP)$tfDapA_1^{7-21}$, pET-28a-(MBP)$tfDapA_1^{12-21}$ proceeded by amplification of regions of pET-28a-(MBP)-$tfDapA_1$. Linearized plasmids were then assembled by blunt end ligation.

**Construction of pET28-*fusC*-His$_6$, pETDuet-(MBP)$dapA_{lasso}$-empty, and pETDuet-$GFP_{eng}$-empty plasmids.** Linearized pET28a and *fusC* fragments were generated by PCR amplification with homology arms inserted by 5' primer overhangs for the amplification of *fusC*. The plasmid was then assembled by Gibson assembly. The same protocol was repeated using pETDuet-(MBP)$dapA_{eng}$-empty and *fusA* to give pETDuet-(MBP)$dapA_{lasso}$-empty. The same protocol was further repeated using pETDuet-(MBP)$dapA_{eng}$-empty and pEB1-mGFPmut2 (Addgene) to give pETDuet-$GFP_{eng}$-empty.

**Protein expression and purification for *Tf*Dap proteins.** *E. coli* BL-21 (DE3) cells containing the appropriate vector construct were grown overnight in 20 mL LB-Lennox (10 g/L tryptone, 5 g/L yeast extract, 5 g/L NaCl) with 50 µg/mL of kanamycin at 37 °C at 200 rpm. Overnight culture was used to inoculate 1 L of Terrific Broth media (12 g/L tryptone, 24 g/L yeast extract, 0.4% *v/v* glycerol, 17 mM $KH_2PO_4$, 72 mM $K_2HPO_4$) supplemented with 50 µg/mL of kanamycin in a 2.8 L Fernbach flask. The flasks were then placed in a shaking incubator set at 37 °C until the $OD_{600}$ reached approximately 0.6-0.8. The incubator temperature was further lowered to 18 °C, and the flasks were incubated for an additional hour. Isopropyl-β-D-thiogalactopyranoside (IPTG) was added to induce protein expression at a final concentration of 1 mM as the flasks were shaken for an additional 18 h. The cells were harvested by centrifugation at 3,000×g at 4 °C for 40 min and the pelleted cells were resuspended in a buffer containing 500 mM NaCl, 20 mM tris(hydroxymethyl)aminomethane (Tris, pH 8.0), and 10% *v/v* glycerol. The resuspended cells were then frozen and stored at -75 °C until purification.

Cells were thawed and lysed by sonication on ice using a Branson 150 Sonifier Cell Disruptor, microtip at 80% amplitude for 6 min of 15 s on and 45 s off. The lysate was clarified by centrifugation at 20,000×g for 20 min or until clear to remove insoluble cellular debris. For the protein purification, an ÄKTAGo FPLC system (Cytiva) at 4 °C was used, and all buffers were filtered through 0.22 µm polyvinylidene difluoride membrane filters. The supernatant was loaded onto a 5 mL HisTrap column (Cytiva) that was equilibrated with buffer A (1 M NaCl, 20 mM Tris, pH 7.5, and 30 mM imidazole). Upon sample loading the column was washed with buffer A. The His$_6$-tagged target protein was then eluted using a linear gradient from 0-100% with buffer B (1 M NaCl, 20 mM Tris pH 7.5, 250 mM imidazole) over a volume of 40 mL, with 5 mL fractions collected. The purity of the eluted fractions was assessed using SDS-PAGE (12% acrylamide), and fractions containing pure protein were concentrated to ~2 mL using Amicon Ultra-15 30 kDa molecular weight cut-off (MWCO) spin concentrators (Amicon; Millipore Sigma). The concentrated protein was loaded onto a preequilibrated HiLoad 16/60 Superdex 75 (GE Healthcare Life Sciences) or 200 (Cytiva) size exclusion column with GF Buffer (20 mM HEPES pH 7.5, 300 mM NaCl, and 10% glycerol), and 3 mL fraction were collected. The collected pure fractions were concentrated by 30 kDa MWCO Amicon to ~1 mL and the concentration was determined using Qubit Protein and Protein Broad Range (BR) Assay Kits on a Qubit 4 Fluorometer (Invitrogen).

**Expression and purification of MBP-tagged $DapA_1$ peptides and variants.** Chemically competent *E. coli* BL21(DE3) cells were transformed with the appropriate vector and plated on selective media. Individual colonies were picked and grown overnight at 37 °C and 250 rpm in 10 mL LB-Miller cultures supplemented with the appropriate antibiotic. Overnight cultures were used to inoculate expression cultures of either 250 mL or 1 L LB-Miller broth with antibiotic at a ratio of 1:100 *v/v*. Expression cultures were placed in a shaking incubator and grown at 37 °C and 220 rpm until reaching an $OD_{600}$ of 0.6-0.8, at which point IPTG was added to induce protein expression at a final concentration of 250 μM. Cultures were then allowed to continue growing at 37 °C for 4 h. Cells were harvested by centrifugation at 4,500×g for 45 min at 4 °C, and resuspended in phosphate-buffered saline (PBS). Cells were then transferred to 50 mL conical tubes, re-pelleted by centrifugation at 7,500×g for 10 min, flash-frozen in liquid $N_2$, and then stored at -80 °C until purification.

Cell pellets were thawed and resuspended in 5 mL cold lysis buffer (4 °C, pH 8.0, 50 mM Tris, 300 mM NaCl, 0.5 mM imidazole, and 2.5% glycerol) per 1 g cell pellet. Resuspended cells were supplemented with 4 mg/mL lysozyme and a 50× *v/v* protease inhibitor cocktail containing E-64 (40 μg/mL), leupeptin (50 μg/mL), benzamidine HCl (15 mg/mL), and PMSF (1.3 mg/mL). Resuspended cells were then nutated at 4 °C for 15-30 min prior to sonication. Sonication was performed using a Fisherbrand Model 120 Sonic cell disruptor set to 40% amplitude for 2-5 min of total sonication time depending on volume of lysate and degree of observed cell disruption. Cell lysate was then clarified by centrifugation at 20,000×g for 45 min. Clarified cell lysate was added to a gravity column containing pre-equilibrated HisPur Ni-NTA resin (Thermofisher). Loaded columns were allowed to nutate for 30 min at 4 °C before allowing lysate to flow through. The resin was subsequently washed with 10 column volumes (CV) of cold lysis buffer (4 °C, pH 8.0, 50 mM Tris, 300 mM NaCl, 0.5 mM imidazole, and 2.5% *v/v* glycerol, followed by 20 CV of cold wash buffer (4 °C, pH 8.0, 50 mM Tris, 300 mM NaCl, 30 mM imidazole, and 2.5% *v/v* glycerol). Columns were stopped, and 5 CV of cold elution buffer (4 °C, pH 8.0, 50 mM Tris, 300 mM NaCl, 200 mM imidazole, and 2.5% *v/v* glycerol) were added to the resin for 10 min. The eluent was then collected and buffer exchanged into a protein storage buffer (4 °C, pH 8.0, 50 mM HEPES, 300 mM NaCl, and 2.5% *v/v* glycerol) at least 100-fold using a 3 kDa or 30 kDa MWCO Amicon. Protein quantities were estimated by measurement of absorbance at 280 nm using a Nanodrop 2000, adjusted for the predicted molecular weight and extinction coefficient of each protein (determined using Expasy ProtParam software). Proteins were then concentrated as necessary using an Amicon. Sample purity was assessed using SDS-PAGE and/or mass spectrometry analysis of TEV protease digests. Samples were aliquoted into separate tubes, flash-frozen in liquid $N_2$, and then stored at -80 °C until use.

**Expression and purification of MBP-tagged *Sa*Dap enzymes.** Protocols for (MBP)*Sa*Dap enzymes proceeded as for MBP-tagged $DapA_1$ peptides with the following modifications. Upon reaching an $OD_{600}$ measurement of 0.6-0.8, cultures were cooled at 4 °C for 15 min, prior to the addition of 250 μM IPTG. Cultures were then grown overnight at 18 °C with 220 rpm shaking.

**Expression and purification of fusilassin biosynthetic enzymes.** (MBP)FusB and (MBP)FusE were prepared as previously described.[6] Preparation of FusC-$His_6$ began by co-transformation of pet28-*fusC*-$His_6$ with pGro7 (Takara Bio) into chemically competent *E. coli* Tuner (DE3) cells. Transformation outgrowth was plated onto an LB-agar plate supplemented with 30 μg/mL kanamycin and 25 µg/mL chloramphenicol. Colonies were picked and grown in 5 mL overnight LB cultures supplemented with 30 µg/mL kanamycin and 25 µg/mL chloramphenicol. Overnight cultures were used to inoculate 250 mL LB-Miller expression cultures with a ratio of 1:100 *v/v*. Cultures were further supplemented with 30 µg/mL kanamycin and 25 µg/mL chloramphenicol. Cultures were placed in a shaking incubator and grown at 37 °C and 220 rpm until reaching an $OD_{600}$ of 0.3-0.4, at which point arabinose (0.02% *w/v* or 0.1% *w/v*) was added to induce production of GroES and GroEL. Cultures were placed back into the shaking incubator and grown at 37 °C and 220 rpm until reaching an $OD_{600}$ of 0.8-1.0, at which point IPTG (100 µM or 500 µM) was added to induce protein expression. Protein expression continued at 37 °C and 220 rpm for 4 h. Purification of FusC-His6 was conducted as described for MBP-tagged $DapA_1$ peptides, except the lysis, wash, and elution buffers were prepared with 5% *v/v* glycerol, and the protein storage buffer was prepared with 10% *v/v* glycerol.

**Matrix-assisted laser desorption/ionization time-of-flight mass spectrometry (MALDI-TOF-MS).** MALDI-TOF-MS data were collected using a Bruker Ultraflextreme enabled for LIFT. Crystallization was performed using 50 mg/mL Super-DHB (SDHB; Thermo Scientific) prepared in 40:59.9:0.1 water:acetonitrile:formic acid (FA) solvent. Mass spectra were collected using positive-ion, reflector mode with methods appropriate for the mass range of interest. For samples acquired by LIFT mode (MALDI-LIFT-TOF/TOF-MS), the samples were first analyzed using the above description. Following identification and/or confirmation of the ion of interest, the LIFT method was loaded onto the FlexControl software, and target ions were selected for fragmentation. Data analysis was performed using FlexAnalysis version 3.3 (Bruker).

Samples were mixed 1:1 with the SDHB solution and spotted onto a ground steel 384 target plate (Bruker). Samples requiring desalting were purified by C18 ZipTip (Pierce). The ZipTip was conditioned in acetonitrile, 50% acetonitrile, and water in succession. Samples were loaded onto the ZipTip by pipetting, followed by 5×10 µL washes with water to remove salts. Following the wash steps, 2 µL of SDHB solution was loaded into the ZipTip and allowed to sit for 30 s. The solution was then ejected directly onto the ground steel target plate. All samples were allowed to dry in ambient air to allow co-crystallization of the matrix and analyte.

**Solid phase peptide synthesis (SPPS) and purification.** Peptides were synthesized using a Biotage Initiator+ Alstra Automated Microwave Peptide Synthesizer with fluorenylmethyl-oxycarbonyl (Fmoc) chemistry. The synthesis proceeded in a C-to-N terminus direction, incorporating Fmoc amino acids [5 equivalents, 0.5 M in *N*,*N*-dimethylformamide (DMF)] onto L-tyrosine-2-chlorotrityl resin (0.1 mmol, 0.64 mmol/g), all sourced from ChemImpex.

The process began with resin swelling in DMF for 20 min at 70 °C, followed by Fmoc deprotection at 25 °C for 10 min using 20% *v/v* 4-methylpiperidine in DMF. Coupling reactions were performed using diisopropylcarbodiimide (DIC) and Oxyma (0.5 M in DMF) for 5 min at 75 °C, followed by another round of Fmoc deprotection. For subsequent steps, each amino acid was introduced with programmed methods involving coupling with DIC/Oxyma at 75 °C for 5 min and deprotection with 20% *v/v* piperidine in DMF at 25 °C. Bulkier amino acids, such as tryptophan and phenylalanine, as well as consecutive hydrophobic residues, were double coupled at 50 °C for 10 min per cycle to ensure efficient incorporation. For the longer peptides *Tf*$DapA_1^{(-12)—21}$ and *Tf*$DapA_1^{(-10)—21}$, *N*-methylpyrrolidone (NMP) was used as a solvent instead of DMF.

Upon completion of the synthesis, the resin was washed sequentially with dichloromethane (DCM, 8 mL×3) and diethyl ether (8 mL×3). The crude peptide was then cleaved from the resin using a trifluoroacetic acid (TFA)/triisopropylsilane/water solution (95:2.5:2.5; 10 mL) for 4 hours. The peptide was precipitated with cold ether, followed by centrifugation (4,000×g, 5 min) and removal of the supernatant by decantation. The precipitated peptides were dried overnight using a SpeedVac (Savant SpeedVac SPD120 Vacuum Concentrator) connected to a vapor trap (Savant RVT5105 Refrigerated Vapor Trap) and vacuum pump (Thermo Scientific VLP120 Vacuum Pump). Peptides were confirmed using UPLC-HR-MS. The crude peptides were then stored at -20 °C until purification.

A portion of the crude peptide was dissolved in a 1:1 mixture of NMP and water and passed through a HyperSep C18 column (2,000 mg, Thermo Scientific) pre-washed with acetonitrile containing 0.1% *v/v* FA, followed by water with 0.1% *v/v* FA. The column was subsequently washed with 25:75, 50:50, 75:25, and 100:0 mixtures of acetonitrile (0.1% *v/v* FA):water (0.1% *v/v* FA). The fractions containing peptides were confirmed using UPLC-MS. The peptide fractions were then dried using a SpeedVac, and the concentrated peptides were resuspended in a 1:1:1 mixture of NMP/acetonitrile/water (600 µL) for further purification by HPLC.

**Purification of SPPS peptides with HPLC.** The resuspended peptides were purified on an Agilent 1200 HPLC system (with an Agilent HPLC G1312-90006 Binary Pump, Agilent 1260 Infinity Diode Array and an Agilent 1200 G1364-90010 Fraction collector) connected to a Luna Omega C18 (5 µm, 100 Å, 250×4.6 mm)/Kinetex Polar C18 (2.6 µm, 100 Å, 150×4.6 mm) fitted with a C18 guard cartridge; purification was monitored at 220 nm and 280 nm. The crude *Tf*DapA$_1$ peptide, obtained from GenScript, was dissolved in a 1:1:1 mixture of NMP/acetonitrile/water. A 100 µL aliquot of the crude sample was loaded onto the Luna Omega C18 column (5 µm, 100 Å, 250×4.6 mm) and purified using the following LC parameters: solvent A, water (10 mM ammonium bicarbonate); solvent B, acetonitrile; flow rate, 1 mL/min; gradient, 5% B from 0—5 min, 5—60% B from 5—35 min, 60—100% B from 35—40 min, 100% B from 40—45 min, 100—5% B from 45—50 min. Fractions were manually collected with *Tf*DapA$_1$ elution confirmed by LC-MS (*Tf*DapA$_1$ – 14.5 min). Fractions containing *Tf*DapA$_1$ were pooled, dried, and resuspended in water to achieve a final concentration of 1 mM of *Tf*DapA$_1$.

*Tf*DapA$_1^{(-12)—21}$ and *Tf*DapA$_1^{(GS)2}$ were each purified with a Luna Omega C18 column (5 µm, 100 Å, 250×4.6 mm). *Tf*DapA$_1^{(-12)—21}$ was purified using the following LC parameters: solvent A, water (10 mM ammonium bicarbonate); solvent B, acetonitrile; flow rate, 1 mL/min; gradient, 5% B from 0—5 min, 5—60% B from 5—35 min, 60—100% B from 35—40 min, 100% B from 40—45 min, 100—5% B from 45—50 min. Fractions were manually collected (*Tf*DapA$_1^{(-12)—21}$ – 15.2 min). *Tf*DapA$_1^{(GS)2}$ was purified using the following LC parameters: solvent A, water (10 mM ammonium bicarbonate); solvent B, acetonitrile; flow rate, 1 mL/min; gradient, 20% B from 0—5 min, 20—30% B from 5—35 min, 30—100% B from 35—40 min, 100% B from 40—45 min, 100—20% B from 45—50 min. Fractions were manually collected (*Tf*DapA$_1^{(GS)2}$ – 11.0 min).

The resuspended synthetic peptides (*Tf*DapA$_1^{(-10)—21}$, *Tf*DapA$_1^{(GS)6}$, *Tf*DapA$_1^{(GS)4}$, *Tf*DapA$_1^{(-10)—4,17—21}$) were loaded onto a Kinetex Polar C18 column (2.6 µm, 100 Å, 150×4.6 mm) and purified using the following LC parameters: solvent A, water (10 mM ammonium bicarbonate); solvent B, acetonitrile; flow rate, 1 mL/min; gradient, 5% B from 0—5 min, 5—60% B from 5—35 min, 60—100% B from 35—40 min, 100% B from 40—45 min, 100—5% B from 45—50 min. Fractions were manually collected (*Tf*DapA$_1^{(-10)—21}$ – 14.6 min, *Tf*DapA$_1^{(GS)6}$ – 10.6 min, *Tf*DapA$_1^{(GS)4}$ – 11 min, *Tf*DapA$_1^{(-10)—4,17—21}$ – 12.8 min).

***In vitro* enzymatic assays using chemically-synthesized peptides.** Enzyme assays were performed in GF Buffer (20 mM HEPES pH 7.5, 300 mM NaCl, 10% glycerol) using 100 µM of precursor peptide and 10 µM of purified enzyme. Where appropriate, co-substrates were added as follows: $NAD^+$, 250 µM; pyridoxal phosphate (PLP), 100 µM; L-Lys, 500 µM; (*S*)-adenosylmethionine (SAM), 100 µM; ATP, 100 µM; $MgCl_2$, 1 mM. In assays involving multiple enzymes, except *Tf*DapBC, each additional enzyme and its corresponding co-substrates were added following an overnight incubation with the preceding enzyme.

Total reaction volumes were brought to 50 µL using GF Buffer and incubated at room temperature overnight. The assay was quenched with equal volume of 100% acetonitrile, and precipitate was removed by centrifugation (21,000×g) for 5 min. The supernatant was collected and analyzed using UHPLC-HR-MS analysis. Controls containing no enzyme were performed in presence of 100 µM peptide and respective enzyme co-substrates in GF Buffer. Where appropriate, chymotrypsin (Sequencing grade, Promega, .01 µg/µL in 1 mM HCl) was added to the assays after the enzyme reaction, and samples were digested overnight.

**UHPLC-HR-MS methods.** General UHPLC-HR-MS analysis was conducted using a Dionex Ultimate 3000 UPLC system (Thermo Scientific) coupled with an LTQ Orbitrap Elite (Thermo Scientific) mass spectrometer in positive mode. For general methods the mass range was set to 400-2,000 *m/z* with dd-$MS^2$ (data-dependent MS/MS) at collision energies of 25 eV in CID. Chromatographic separation was achieved using an ACQUITY UPLC BEH C18 1.7 µm 2.1×50 mm column with the following LC parameters: solvent A, water (0.1% *v/v* FA); solvent B, acetonitrile (0.1% *v/v* FA); column temperature, 35 °C; flow rate, 0.3 mL/min; gradient, 5% B from 0—1 min (directed to waste), 5—95% B from 1—10 min, 95% B from 10—11 min, 95—5% B from 11-12 min. Specific chromatographic methods were developed for various peptide samples, and an internal standard of fluorescein was added to all samples prior to the LC-MS run to normalize retention times.

Assays with *Tf*DapA$_1$: BioZen 1.7 µm Peptide XB-C18 2.1×50 mm column; gradient, 30% B from 0—1 min (directed to waste), 30—85% B from 1—10 min, 85% B from 10—11 min, 85—30% B from 11-12 min; flow rate 0.3 mL/min; mass range 400-2,000.

Chymotrypsinized *Tf*DapA$_1$: BioZen 1.7 µm Peptide XB-C18 2.1×50 mm column; gradient, 5% B from 0—1 min (directed to waste), 5—95% B from 1—10 min, 95% B from 10—11 min, 95—5% B from 11-12 min; flow rate 0.3 mL/min; mass range 100-1,000.

Assays with *Tf*DapA$_1^{(-12)—21}$: ACQUITY UPLC BEH C18 1.7 µm 2.1×50 mm column; gradient, 20% B from 0—1 min (directed to waste), 20—55% B from 1—10 min, 55% B from 10—11 min, 55—20% B from 11-12 min; flow rate 0.3 mL/min; mass range 200-2,000.

Assays with *Tf*DapA$_1^{(-10)—21}$: ACQUITY UPLC BEH C18 1.7 µm 2.1×50 mm column; gradient, 30% B from 0—1 min (directed to waste), 30—85% B from 1—10 min, 85% B from 10—11 min, 85—30% B from 11-12 min; flow rate 0.3 mL/min; mass range 200-2,000.

Assays with *Tf*DapA$_1^{(GS)n}$: ACQUITY UPLC BEH C18 1.7 μm 2.1×50 mm column; gradient, 20% B from 0—1 min (directed to waste), 20—35% B from 1—10 min, 35% B from 10—11 min, 35—20% B from 11-12 min; flow rate 0.3 mL/min; mass range 200-2,000.

For MS/MS fragmentation, all ions were selected to generate the b and y ions for fragmentation tables using a mass tolerance of 5 ppm, mass precision of four decimals, and enabling curve smoothing (Gaussian, 9 points). We additionally note that for synthetic *Tf*DapA$_1$, Met residues were found to be oxidized in all cases during UHPLC-HR-MS. Masses reported for *Tf*DapA$_1$ assays include the mass of 3 additional O atoms to account for these oxidations. In some cases, the five highest abundance ions from the expected isotope envelope were calculated using ChemDraw (version 23.1.2.7) and extracted (5 ppm error tolerance). These *m/z* values are listed below, corresponding to the +3 charge state of each analyte (Aac, aminoacetone; Dap, 1,2-diaminopropane; Dmp, dimethylpropane-1,2-diamine; Miz, 4-methylimidazoline; Diz, 3,4-dimethylimidazoline).

*Tf*DapA$_1$-Thr - 1681.8085, 1681.4741, 1682.1430, 1682.4774, 1681.1396
*Tf*DapA$_1$-Aac - 1666.4734, 1666.1389, 1666.8078, 1667.1423, 1665.8045
*Tf*DapA$_1$-Dap - 1666.8172, 1666.4828, 1667.1517, 1667.4862, 1666.1483
*Tf*DapA$_1$-Dmp- 1676.1610, 1675.8266, 1676.4955, 1676.8299, 1675.4921
*Tf*DapA$_1$-Miz - 1660.8137, 1660.4793, 1661.1482, 1661.4826, 1660.1448
*Tf*DapA$_1$-Diz - 1665.4856, 1665.1512, 1665.8201, 1666.1545, 1664.8167

***In vitro* enzymatic assays using MBP-tagged substrates.** Enzyme assays were performed in a Tris-based reaction buffer (50 mM Tris-HCl pH 7.5, 125 mM NaCl, 10 mM DTT) using 100 μM of precursor peptide and 10 μM of purified enzyme. Where appropriate, the following supplementations were added to the enzyme reactions: $NAD^+$, 750 μM; L-Lys, 500 μM; SAM, 2 mM; ATP, 5 mM; $MgCl_2$, 20 mM. Reactions were typically performed at 50 μL scale for 3 h at 37 °C with all enzymes added immediately. TEV protease was added to the reactions 1:100 (*m/m*) relative to substrate after 3 h, and proteolysis was allowed to proceed for 1 h at room temperature. Samples were then analyzed by desalting and MALDI-TOF-MS. Variations in component concentrations are described in the table below.

| Assay | Variations of concentration |
|---|---|
| (MBP)*Tf*DapA$_1$BC | 50 μM precursor peptide, 5 μM enzymes |
| (MBP)*Tf*DapBCD amino acid scan assays | 50 μM precursor peptide, 5 μM enzymes |
| DapA$_{eng}$ scaleup | 2 μM enzymes (*Tf*DapY, *Tf*DapY/*Sa*DapM) |
| *Tf*DapB$_{conf}$ assays | 50 μM substrate peptide |
| Leader in trans assays | 50 μM substrate peptide, 10 μM leader peptide |

Reaction for evaluation of *Sa*DapY and *Sa*DapM order of events was conducted as above, but with *Sa*DapM added after 3 h of reaction. The reaction was then allowed to proceed overnight, and TEV protease was added the following day. Fusilassin reactions were conducted as above, except FusBCE enzymes were supplemented as necessary after 1 h rather than immediately. The protocol for determining enzymatic tolerance to lyophilization was performed by preparing a reaction as described above at 100 μL scale but lacking substrate. The total reaction was lyophilized overnight and then reconstituted in 100 μL $H_2O$ before proceeding by addition of substrate. For *Tf*DapBCDM buffer tolerance reactions with $DapA_{eng}$, five different buffers were examined: Tris – 20 mM Tris-HCl, 100 mM NaCl, pH 7.5; HEPES – 20 mM HEPES, 100 mM NaCl, pH 7.5; phosphate-buffered saline (PBS) – 137 mM NaCl, 2.7 mM KCl, 10 mM $Na_2HPO_4$, 1.8 mM $KH_2PO_4$, pH 7.4; MES – 50 mM MES, 100 mM NaCl, pH 6; CHES – 100 mM CHES, 100 mM NaCl, pH 9.

**Direct infusion HR-MS and HR-MS/MS methods.** Direct infusion HR-MS data was collected on a ThermoFisher Scientific Orbitrap Fusion ESI-MS. The Orbitrap Fusion was calibrated using Pierce LTQ Velos ESI Positive Ion Calibration Solution (ThermoFisher). Prior to data acquisition, samples were desalted by either HPLC or C18 ZipTip and dried. Samples were resuspended in 60:39.9:0.1 acetonitrile:water:acetic acid and subjected to centrifugation at 17,000×g to remove insoluble material. Samples were then directly infused into the Orbitrap Fusion ESI-MS using an Advion TriVersa Nanomate 100. The MS operated with the following parameters: 100,000 resolution, 1 *m/z* isolation width (MS/MS), 0.4 activation q value (MS/MS), and 30 ms activation time (MS/MS). Fragmentation of ions of interest was performed using collision-induced dissociation at 30 normalized collision energy. Data analysis was performed using the Qualbrowser application of Xcalibur software (version 4.1.31.9) and the Interactive Peptide Spectral Annotator Tool (http://www.interactivepeptidespectralannotator.com/PeptideAnnotator.html).

**Purification of modified AISLT pentapeptides.** *In vitro* reactions of $DapA_{eng}$-Dap were reacted with TEV protease (1:100 *m/m*). A 3 kDa MWCO Amicon was pre-washed by 3 rounds of centrifugation at 4,000×g with 10 mM aq. ammonium bicarbonate. After 4 h proteolysis reaction, the reaction mixture was loaded into the pre-washed Amicon. The sample was then subjected to centrifugation at 4,000×g, and flowthrough was collected containing the modified pentapeptide fragment. Once the total volume of retentate had reduced to <1 mL, an additional 9 mL of 10 mM aq. ammonium bicarbonate was added. Centrifugation was continued and additional flowthrough was collected. The flowthrough samples were combined, and the total volume was reduced by evaporation using a SpeedVac. Residual protein was precipitated by dropwise addition of trifluoroacetic acid. Insoluble material was removed by centrifugation at 17,000×g, and the supernatant was removed. The total volume was again reduced by evaporation in a SpeedVac, followed by dilution to a final concentration of 20% *v/v* acetonitrile.

Peptides were purified on a Thermo Vanquish Core HPLC equipped with an Accucore C18 analytical column (Thermo Scientific; 150 mm×4.6 mm, 2.6 μm particle size, 80 Å pore size) and photodiode array detector (190-800 nm). The products of reaction with *Tf*DapY were purified using the following LC parameters: solvent A, water (20 mM ammonium acetate); solvent B, acetonitrile; flow rate, 2 mL/min; gradient, 1% B from 0—2 min, 1—10% B from 2—3 min, 10—30% B from 3—9 min, 30—98% B from 9—10 min, 98% from 10—14 min, 98—1% B from 14—15 min, 1% B from 15—22 min. The products of reaction with *Tf*DapY and (MBP)*Sa*DapM were purified using the following LC parameters: solvent A, water (20 mM ammonium acetate); solvent B, acetonitrile; flow rate, 2 mL/min; gradient, 1% B from 0—2 min, 1—17% B from 2—4 min, 17—22% B from 4—14 min, 22—95% B from 14—15 min, 95% from 15—20 min, 98—1% B from 20—21 min, 1% B from 21—28 min. Fractions were collected and screened by MALDI-TOF-MS to identify analytes of interest, and the corresponding fractions were pooled. The combined fractions were evaporated to dryness in a SpeedVac prior to NMR analysis.

**NMR of AISLT pentapeptides.** Spectra were recorded on a Bruker Avance NEO 600 MHz spectrometer with a 5-mm Prodigy BBO probe using Bruker TopSpin software (Version 4.2.1, Bruker). Standard Bruker pulse sequences with Watergate solvent suppression were utilized for the following experiments: $^{1}$H, $^{1}$H-$^{1}$H COSY, $^{1}$H-$^{1}$H DQF-COSY, $^{1}$H-$^{1}$H TOCSY, $^{1}$H-$^{13}$C HSQC, and $^{1}$H-$^{1}$H NOESY. Samples (~100 µg) were dissolved in 79.9:20:0.1 $H_2O$:$CD_3CN$:$CDO_2D$ solvent. Spectral processing was performed in MestReNova (version 14.3.3, Mestrelab Research).

**Marfey's analysis.** Purified Diz-modified pentapeptide (~30 µg) was dried by evaporation in a Speedvac. The sample was resuspended in 500 µL 6M DCl in $D_2O$, transferred to an 8 mL borosilicate glass reaction vial, and the vial was sealed with a Teflon cap. A fitted aluminum reaction block was placed in an Optimag ST digital hotplate at ambient temperature, and the reaction vial was placed into the block. A stream of air was further directed to the top portion of the glass to allow condensation of vapors. The hotplate was turned on and set to 110 °C, and the hydrolysis reaction was allowed to proceed at 110 °C for 14 h. The hotplate was then turned off and allowed to cool to room temperature. The reaction mixture was subjected to centrifugation at 17,000×g for 5 min to remove insoluble material, and the soluble portion was transferred to a separate vial for evaporation to dryness in a Speedvac. Upon reaching dryness, the sample was resuspended in 250 µL $H_2O$ and re-dried in the Speedvac to aid evaporation of residual acid.

The hydrolysate was dissolved in 100 µL 1M aq. $NaHCO_3$. A stock solution of 1-fluoro-2,4-dinitrophenyl-5-L-alanine amide (L-FDAA) was prepared in acetone at a concentration of 12 mg/mL, and 100 µL was added to the dissolved hydrolysate. The contents were mixed and heated to 60 °C for 2 h without shaking using an Eppendorf Thermomixer C. A color change was observed from yellow to a dark orange/red. The reaction mixture was then quenched by dropwise addition of 6M HCl (~40 µL), resulting in return to a yellow color. The sample was then subjected to centrifugation at 17,000×g to remove insoluble material prior to LC or LC-MS analysis.

To prepare derivatized standards of L-amino acids, two aliquots of 50 µL L-amino acid solution (0.5 µmol/mL) were dried using a Speedvac concentrator. The dried amino acid stocks were resuspended in 50 µL 1M aq. $NaHCO_3$, followed by addition of 50 µL of 12 mg/mL L-FDAA in acetone or 50 µL of 12 mg/mL 1-fluoro-2,4-dinitrophenyl-5-D-alanine amide (D-FDAA) in acetone, respectively. The contents were mixed and the reaction proceeded at 60 °C as above. To prepare derivatized standards for isomers of *N*-methylpropane-1,2-diamine, each synthesized standard was diluted 50-fold in $H_2O$. To 49 µL 1M aq. $NaHCO_3$, 1 µL of the diluted standard was added. Subsequently, 50 µL of 12 mg/mL L-FDAA was added to each reaction, contents were mixed, and the reactions proceeded at 60 °C as above.

Analysis of L-amino acid contents was performed using a Shimadzu LCMS-2020 equipped with a Hypersil Gold C18 Selectivity HPLC Column (Thermo Scientific; 50 mm×4.6 mm; 3 µm particle size, 175 Å pore size). The following parameters were used: solvent A, water (0.1% *v/v* FA); solvent B, acetonitrile (0.1% *v/v* FA); flow rate, 0.4 mL/min; gradient, 15% B from 0—7 min (directed to waste), 15—80% B from 7—53 min, 80—95% B from 53—54 min, 95% B from 54—59 min, 95—15% B from 59—61 min, and 15% B from 61—66 min; MS detection, 7—66 min; mass range 100—1,000 *m/z*. Data analysis was performed using LabSolutions software (Shimadzu). Where necessary, samples were diluted further to prevent saturation of the MS detector. We consistently observed retention time drift across different samples, so the derivatized hydrolysate was mixed with each derivatized amino acid standard mix individually to allow confirmation of stereochemistry by co-elution. Analysis of methylpropane-1,2-diamine (Mmp) contents was performed using a Thermo Vanquish Flex HPLC equipped with an Accucore PFP HPLC column (Thermo Scientific; 150 mm×4.6 mm, 2.6 µm particle size, 80 Å pore size) and photodiode array detector (190-800 nm). The following LC parameters were used: solvent A, water (0.1% *v/v* FA); solvent B, acetonitrile (0.1% *v/v* FA); flow rate, 1.8 mL/min; gradient, 20% B from 0—min, 20—60% B from 5—25 min, 60—95% B from 25—27 min, 95% B from 27—35 min, 95—20% B from 35—38 min, 20% B from 38—47 min. Derivatized Mmp compounds were detected by monitoring UV absorbance at 340 nm. Data analysis was performed using Chromeleon software (Thermo Scientific). Following initial injections of derivatized standards and derivatized hydrolysate, the derivatized hydrolysate was further mixed with the suspected isomer of *N*-methylpropane-1,2-diamine and co-injected to allow secondary confirmation by co-elution.

**Production and analysis of azuritides 1-4.** Methods for generation of heterologous expression strain of *S. albus* J1074 containing the *saDap* BGC were followed as described in Ren et al.[1] The plasmid used for the conjugation protocol is provided in **Dataset S2**. Following appearance of exconjugant colonies, individual colonies were re-streaked onto fresh MS-agar (mannitol 20 g/L, soybean flour 20 g/L, agar 20 g/L) supplemented with apramycin (50 µg/mL) and allowed to grow for 5 d at 30 °C, at which point sporulation was apparent. Portions of cell mass and spores were scraped and resuspended in 50 µL MeOH for 1 h to allow metabolite extraction. The cell mass was removed by centrifugation at 17,000×g, and 1 µL of the methanolic fraction was analyzed by MALDI-TOF-MS. Following the identification of metabolites of interest, individual ions were further subjected to MALDI-LIFT-TOF/TOF-MS. Spores of daptide-producing exconjugant colonies were resuspended and used to inoculate MS-agar solid and MS liquid cultures both containing and lacking supplemented apramycin (50 µg/mL), but none of the tested conditions yielded observable azuritide production at larger scale.

**Aminoacetone bioconjugation experiments.** EZ-Link biotin hydrazide (Thermo Scientific) and aminooxy-$PEG_1$-propargyl (MedChemExpress) were each dissolved in DMSO to create 50 mM stock solutions. $GFP_{eng}$ was produced by expression of pETDuet-$GFP_{eng}$-empty plasmid as described above for MBP-tagged *Sa*Dap proteins. Biotinylation reactions were conducted in PBS (pH 7.4) supplemented with 750 µM $NAD^+$, 5 µM enzymes, and 50 µM substrate. Propargylation reactions were conducted in Tris-based reaction buffer (50 mM Tris-HCl pH 7.5, 125 mM NaCl) supplemented with 750 µM $NAD^+$, 2 µM enzymes, and 25 µM substrate. For one-pot biotinylation reactions, biotin-hydrazide was added to the reaction at a final concentration of 2 mM. For step-wise bioconjugation reactions, biotin-hydrazide or aminooxy-$PEG_1$-propargyl stock solutions were added to the reaction 1:25 (*v/v*). Aniline was added as a catalyst for hydrazone or oxime formation to a final concentration of 1 mM at the time of biotin hydrazide or aminooxy-$PEG_1$-propargyl addition. $DapA_{eng}$-biotin conjugates were subjected to 500-fold buffer exchange prior to TEV proteolysis and confirmation by LC-MS. $GFP_{eng}$-propargyl conjugates were digested by GluC, desalted by C18 ZipTip, and confirmed by MALDI-TOF-MS. Glucagon-biotin conjugates were desalted by C18 ZipTip and confirmed by MALDI-TOF-MS.

**Chemical synthesis of *N*-methyl-1,2-propanediamine standards.** Diethyl ether (ACS grade), dichloromethane (ACS grade), tetrahydrofuran (HPLC grade), acetonitrile (HPLC grade), and toluene (ACS grade) were dried for reactions using the MB-SPS solvent purification system containing activated alumina manufactured by MBRAUN. Reaction temperatures correspond to the external temperature of the reaction vessel. Analytical thin-layer chromatography (TLC) was performed on Merck silica gel 60 F254 aluminum sheets. Visualization was accomplished with UV light and/or potassium permanganate ($KMnO_4$). Silicycle SiliaFlash P60 ($SiO_2$, 40–63 μm particle size, 230–400 mesh) was used for flash column chromatography.

**(*R*)-*N₁*-methylpropane-1,2-diamine.** To a suspension of D-Ala-NHMe·HCl (600 mg, 4.33 mmol, 1.0 equiv.) in $Et_2O$ (15 mL) was slowly added $LiAlH_4$ (821 mg, 21.6 mmol, 5.0 equiv.) in $Et_2O$ (6.6 mL) at 0 °C. The reaction mixture was heated up to reflux and stirred for 24 hours. The reaction was cooled down and quenched with $H_2O$ (1 mL) and 1M aq. NaOH (0.2 mL) at 0 °C and stirred for 30 minutes. The reaction mixture was filtered through Celite and washed with $Et_2O$ (30 mL). The filtrate was carefully concentrated in *vacuo* to give (*R*)-$N_1$-methylpropane-1,2-diamine as a colorless oil (72.1 mg, 0.82 mmol, 19%). *Caution: This compound is volatile and co-evaporates with $Et_2O$ during evaporation.*

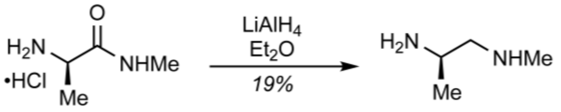

D-Ala-NHMe·HCl (*R*)-$N_1$-methylpropane-1,2-diamine

$^1$H NMR (600 MHz, $CDCl_3$) δ 2.98 (dqd, *J* = 8.4, 6.4, 4.3, 1H), 2.52 (dd, *J* = 11.7, 4.2, 1H), 2.42 (s, 3H), 2.34 (dd, *J* = 11.7, 8.4, 1H), 1.05 (d, *J* = 6.4, 3H)

$^1$H-$^{13}$C HSQC (2D, 600 MHz, $CDCl_3$) δ 60.6, 46.4, 36.7, 22.1

$[\alpha]_D^{20}$ +31.9 (*c* = 2.0 in $CHCl_3$)

**(*S*)-*N₁*-methylpropane-1,2-diamine.** To a suspension of D-Ala-NHMe·HCl (900 mg, 6.49 mmol, 1.0 equiv.) in $Et_2O$ (20 mL) was slowly added $LiAlH_4$ (1.23 g, 32.5 mmol, 5.0 equiv.) in $Et_2O$ (13 mL) at 0 °C. The reaction mixture was heated up to reflux and stirred for 48 hours. The reaction was quenched with $H_2O$ (1.5 mL) and 1 M NaOH aq. (0.3 mL) at 0 °C and stirred for 30 minutes. The reaction mixture was filtered through Celite and washed with $Et_2O$ (50 mL). The filtrate was carefully concentrated in *vacuo* to give (*S*)-$N_1$-methylpropane-1,2-diamine as a colorless oil (50.9 mg, 0.58 mmol, 9%). *Caution: This compound is volatile and co-evaporates with $Et_2O$ during evaporation.*

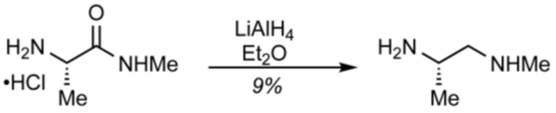

L-Ala-NHMe·HCl (*S*)-$N_1$-methylpropane-1,2-diamine

$^1$H NMR (600 MHz, $CDCl_3$) δ 2.96 (dqd, *J* = 8.4, 6.4, 4.2, 1H), 2.50 (dd, *J* = 11.7, 4.2, 1H), 2.39 (s, 3H), 2.32 (dd, *J* = 11.7, 8.4, 1H), 1.03 (d, *J* = 6.5, 3H)

$^1$H-$^{13}$C HSQC (2D, 600 MHz, $CDCl_3$) δ 60.6, 46.3, 36.6, 22.1

$[\alpha]_D^{20}$ –20.5 (*c* = 2.0 in $CHCl_3$)

**(*R*)-*N*$_2$-methylpropane-1,2-diamine.** To a suspension of (*R*)-2-(methylamino)propanamide hydrochloride (580 mg, 4.18 mmol, 1.0 equiv.) in $Et_2O$ (14 mL) was slowly added $LiAlH_4$ (794 mg, 21.6 mmol, 5.0 equiv.) in $Et_2O$ (7.0 mL) at 0 °C. The reaction mixture was heated up to reflux and stirred for 48 hours. The reaction was quenched with $H_2O$ (1 mL) and 1 M NaOH aq. (0.2 mL) at 0 °C and stirred for 30 minutes. The reaction mixture was filtered through Celite and washed with $Et_2O$ (30 mL). The filtrate was carefully concentrated in *vacuo* to give (*R*)-*N*$_2$-methylpropane-1,2-diamine as a colorless oil (50.6 mg, 0.57 mmol, 14%). *Caution: This compound is volatile and co-evaporates with Et$_2$O during evaporation.*

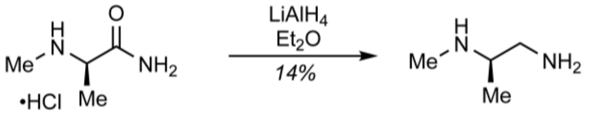

(*R*)-2-(methylamino)propanamide hydrochloride    (*R*)-*N*$_2$-methylpropane-1,2-diamine

| | |
|---|---|
| $^1$H NMR | (600 MHz, $CDCl_3$) δ 2.71 (dd, *J* = 12.4, 4.4, 1H), 2.54 (dd, *J* = 12.4, 6.7, 1H), 2.52–2.47 (m, 1H), 2.41 (s, 3H), 1.02 (d, *J* = 6.2, 3H) |
| $^1$H-$^{13}$C HSQC | (2D, 600 MHz, $CDCl_3$) δ 57.0, 47.1, 33.9, 17.7 |
| $[\alpha]_D^{20}$ | +22.7 (*c* = 2.0 in $CHCl_3$) |

**(*S*)-*N*$_2$-methylpropane-1,2-diamine.** To a suspension of (*S*)-2-(methylamino)propanamide hydrochloride (800 mg, 5.77 mmol, 1.0 equiv.) in $Et_2O$ (20 mL) was slowly added $LiAlH_4$ (1.10 g, 28.9 mmol, 5.0 equiv.) in $Et_2O$ (9.0 mL) at 0 °C. The reaction mixture was heated up to reflux and stirred for 48 hours. The reaction was quenched with $H_2O$ (1 mL) and 1 M NaOH aq. (0.2 mL) at 0 °C and stirred for 30 minutes. The reaction mixture was filtered through Celite and washed with $Et_2O$ (30 mL). The filtrate was carefully concentrated in *vacuo* to give (*S*)-*N*$_2$-methylpropane-1,2-diamine as a colorless oil (92.8 mg, 1.05 mmol, 18%). *Caution: This compound is volatile and co-evaporates with Et$_2$O during evaporation.*

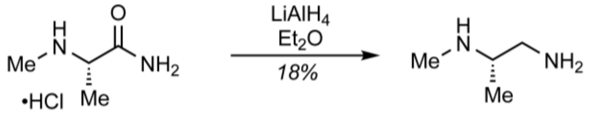

(*S*)-2-(methylamino)propanamide hydrochloride    (*S*)-*N*$_2$-methylpropane-1,2-diamine

| | |
|---|---|
| $^1$H NMR | (600 MHz, $CDCl_3$) δ 2.71 (dd, *J* = 12.4, 4.4, 1H), 2.54 (dd, *J* = 12.4, 6.7, 1H), 2.51–2.47 (m, 1H), 2.40 (s, 3H), 1.02 (d, *J* = 6.2, 3H) |
| $^1$H-$^{13}$C HSQC | (2D, 600 MHz, $CDCl_3$) δ 56.8, 46.9, 33.7, 17.6 |
| $[\alpha]_D^{20}$ | –20.5 (*c* = 2.0 in $CHCl_3$) |

**Table S1. Taxonomic distribution of daptide BGCs.**

| Phylum | # BGCs | Class | # BGCs | Order | # BGCs |
|---|---|---|---|---|---|
| Actinomycetota | 840 | Actinomycetia | 840 | Actinomycetales | 686 |
| | | | | Micromonosporales | 7 |
| | | | | Streptosporangiales | 2 |
| | | | | Propionibacteriales | 3 |
| | | | | Frankiales | 9 |
| | | | | Micrococcales | 123 |
| | | | | Pseudonocardiales | 1 |
| | | | | Corynebacteriales | 4 |
| | | | | Bifidobacteriales | 2 |
| | | | | Kineosporiales | 2 |
| | | | | Mycobacteriales | 1 |
| Bacillota | 194 | Bacilli | 192 | Bacillales | 86 |
| | | | | Lactobacillales | 106 |
| | | Clostridia | 2 | Clostridiales | 2 |

**Table S2. Proteins encoded in the *tfDap* and *saDap* BGCs.** The assembled *Thermobifida fusca* DSM 43792 genome was deposited to the NCBI nucleotide database under Bioproject accession ID PRJNA1271098. Abbreviations: Aac, aminoacetone; Dap, 1,2-diaminopropane; Mmp, mono-methylpropane-1,2-diamine.

| Name | Accession Code | PfamID | Pfam Annotation | Assigned function |
|---|---|---|---|---|
| *Tf*DapJ | WP_011290650.1 | PF00296 | Luciferase-like monooxygenase | Dehydroalanine reduction |
| *Tf*DapB | QOS60427.1 | - | N/A | Thr oxidative decarboxylase partner protein |
| *Tf*DapM | WP_011290652.1 | PF13649 | Methyltransferase domain | Dap methyltransferase |
| *Tf*DapD | WP_107491215.1 | PF00202 | Aminotransferase class-III | Aac aminotransferase |
| *Tf*DapC | WP_061783874.1 | PF00465 | Iron-containing alcohol dehydrogenase | Thr oxidative decarboxylase |
| *Tf*DapA$_1$ | WP_169803321.1 | - | N/A | Daptide precursor peptide |
| *Tf*DapA$_2$ | WP_169803320.1 | - | N/A | Daptide precursor peptide |
| *Tf*DapA$_3$ | WP_167368030.1 | - | N/A | Daptide precursor peptide |
| *Tf*DapA$_4$ | WP_169803318.1 | - | N/A | Daptide precursor peptide |
| *Tf*DapA$_5$ | WP_169803317.1 | - | N/A | Daptide precursor peptide |
| *Tf*DapT$_1$ | WP_061783872.1 | PF00005 | ABC transporter | Daptide export |
| *Tf*DapT$_2$ | WP_061783871.1 | PF12730 | ABC-2 family transporter protein | Daptide export |
| *Tf*DapP | WP_061783870.1 | - | N/A | Daptide leader peptidase |
| *Tf*DapK$_C$ | WP_082797714.1 | PF00069 | Protein kinase domain | Ser dehydration |
| *Tf*DapY | WP_061783869.1 | PF02624 | YcaO cyclodehydratase | Mmp cyclodehydratase |
| *Sa*DapA$_1$ | WP_269085462.1 | - | N/A | Daptide precursor peptide |
| *Sa*DapA$_2$ | WP_269085461.1 | - | N/A | Daptide precursor peptide |
| *Sa*DapA$_3$ | WP_167745747.1 | - | N/A | Daptide precursor peptide |
| *Sa*DapA$_4$ | WP_167745746.1 | - | N/A | Daptide precursor peptide |
| *Sa*DapT$_1$ | WP_236711747.1 | PF00005 | ABC transporter | Daptide export |
| *Sa*DapT$_2$ | WP_059419710.1 | PF12730 | ABC-2 family transporter protein | Daptide export |
| *Sa*DapP | WP_059419709.1 | - | N/A | Daptide leader peptidase |
| *Sa*DapB | WP_059419708.1 | - | N/A | Thr oxidative decarboxylase partner protein |
| *Sa*DapM | WP_059419706.1 | PF13649 | Methyltransferase domain | Dap methyltransferase |
| *Sa*DapD | WP_078945432.1 | PF00202 | Aminotransferase class-III | Aac aminotransferase |
| *Sa*DapC | WP_059419704.1 | PF00465 | Iron-containing alcohol dehydrogenase | Thr oxidative decarboxylase |
| *Sa*DapY | WP_059419702.1 | PF02624 | YcaO cyclodehydratase | Mmp cyclodehydratase |

### Table S3. Key gene sequences from the *tfDap* and *saDap* BGCs.

| Name | Sequence |
|---|---|
| *tfDapA*$_1$ (codon-optimized) | ATGCAGGATATCACGCCGCAACCGAATGAGGTAGCCCTTGAGCTGGAAATGCAAGAATTAGAGGCGATGGA<br>AGCTCCGGGCTTCTGGACTGGCGCAGGCGTCGGAGCATTGATATCAACATCCTTCGCGATAAGCTTGACC |
| *tfDapB* (codon-optimized) | ATGGATAAAACCGAGCTGCGCCGTATTAAGCGCCATTTGATGTCGTGGTCCACTGGGCAGTTCGAGGTGGG<br>TCCTGCGAATCAAGACAGCTGTTCTACTATCGTCGTAGCGGACCCCGCTCACTTGGCTGACTTACTGGACA<br>CCGACTTAGTGGGTGACAATACGGTGGTGTTAGCCCCTGAAGGCCACGGTGCAGATCCTTCCCGTGTGGTT<br>GGATTTGCCGGAACACTGGACGAGCCAAATGCAGAAATGTCTATTGGGTATGACTTCTTCCTGCAAACCCA<br>AGACTACGCTACAAGTCCCTTTATGAGCGTTCTGGGACCAACACTTGTTCGTGTCACCGGTCCTGACGACC<br>TTGAGTTGTTTCTCGCAGACGCTGACCGCGCACGTGAAGAGGGAGTTTTCCCAGACATCGCTGTTGTGCCT<br>GCAGTTCGCATCGCAGACCTTCCAGGTCTCGGAGCCGGGCCTGGTGTTGACGGCCCACAACTGCGCCTCTA<br>TGTCAACGCGGAAGGAGAGATCTCTACGTCCATTGGTGGCTGCTCATTAGGCCGCGTTGGCGACACTTTGG<br>CGTCGTTACGTGCCAATTGGGACCGTCGCAACGCGGCGTCTACAGCCCCTTGTGCCGTATGTCTGGGAGCA<br>GTTCTTCCGGAAGCGGATCGCACACGTGAGCTGCTGGCCCGTCCTTGGCTGGGTCGCTATTTAGCAGCAGT<br>GGATGGTTTACGTATGCTTGCCACACGTGAGATTGTCGGCTTACGTGTGTCAGGGTTCGGGCAACGTCTTA<br>ACCCCGCCCTTGATCACCACCCCGACAGCGCCGACGCCGCTACCCCGCTGTTATTTTGGAATGACGAGAGC<br>GGATACGTGTACGAGCCGGCGTCTGGTCGTACTTTCCAAGTAAATCTTGCCGCGGCAACCGCGGTTGAGGC<br>ATTACTTGTGACTGGTTCGGCGGAGCAGGCTAGTGAATACGCTTCGCCACGCAGCTTAGCAGCCGTGGATC<br>GTTTCTTCACTGAAAGCGGCGTGACACTGACCGGTATCGGGTCAGAGGTTGCGGTG |
| *tfDapC* (codon-optimized) | ATGACCCTTTTACCTACCGCGCCCCCACGCCCCATGTATGGTTGTGGCCATCTCACCGAGTGGTTGCGTCA<br>GAAGGGAGCCACTACGGTTGCCTTGGTCACGGATCAGACTTTAACAGATACTCCAATCCTTGCGACCGTCC<br>GTGCACGCATTGTTGCCGCGGGCGTTCAAGTTCGTTTCTTTGTACTGCCCGACGCGGGTAATCTGGATAGT<br>GTGCTCCGTCTCGCCGACCGCTTTGCGGACTGTGATCTGGTTGTCGGTGTTGGTGGTGGCAGCGTAATCGA<br>CCAAGTCAAATTAGCTGCGACCCTTTGTGGGAATCCGAAGGCTCGCGGCCGCTTGACAGTAGCTCAGCGTA<br>GTGGTACTATCATCCTGCCGGTCACCGTTGAGCGCCGCACGCCCCTTCTGGCCATTCCTACGACCCTCGGG<br>ACCGGTACAGAGCTGAGTGCGGTAGCCTGCTTAGTATATCCTGAGGGGAAGCGTATTATCTCAGGTCTCTG<br>TTTACGTCCTGATGCAGCTGTCCTCGACCCTCTTGCCACCGAGACCTTACCTGTTGAGCTTGTTGCCGAGG<br>GTACTTTAGAGGCGCTGTTCCGCACTGTGAGCCCTTACATTGGGGACCACGCCGACTTGCCAGAGCAAGAC<br>ACCTTAGTTGAGGCCGTTGCGGCAGAGCTGGTACGCTTGGGCGACGAGGTTAATCGCCTGCGTCGCCGCGG<br>CGAGCCAATTTCGCCCCAAATTCGTCTGCGTATCGCGGAATTGAGCGGACGTTCCCAACTTGGCGATCTTC<br>ATGCCGGGCGTAAGCCATACGCCGTCAAGGGCTGGGCGATTGCCAACGAGTTATCATGGGAGCTCTCACTG<br>CGTAAGATGCCTGCGATTGCTGCCCTGTTGCCGCCTTTGTGGCGTCGTATTGCAGACGGCGATCACCGTAT<br>TGGCAGTGCACCCCGTTTGGTCCGTATCTGGCAACGCTTGGCGGAGAACTCCCCTGGCTTACCGGATGACC<br>CGGCCGAGGGAATCGCGGCATTGATTGAGCGCTGGCAGATCGACCACCGTATCACTGCAGATCCGGCGCAA<br>CTCGATACTGTAGCCGCACGCGCCGTCCATGCGTGGGGAGCCGGGCTCCCAGTTTTAGGTGGTCTGACGCA<br>AGCTGAGATCCGTCAACTGTTGGCAGATGCTGTAGCAGATCGTGCCTTGGCACCTGGTACTCCCCAGACTT<br>CGGTCGCG |
| *tfDapD* (codon-optimized) | ATGTCCCACACAACTAGCACGGCACTGTGGCCGTATCTTCTTCCGCCGTCAGCCCATGGGGACGACTCTAT<br>CTGCGCCGTCTCCGCTCGCGGACACCGTGTACGTTTTGCAGATGGTCGTGAGCTCCTGTGTGGGGCCTCTG<br>GCCTTTGGAACGCCAATCTTGGTTACGGAAATCCGGCGATCGCCCAGGCCGTAGCCCAAGCACTCCATGAT<br>GCCTCATACTTAAGCGCGTTCCGTTACGAGAATGTGTATGCACGTCGCGCGGCGGCGGATCTGATCGAGGT<br>GTCGGGCCCAGAGCATTATTCGCGCGTTCTGTTCTCATCCAGCGGTGGGGCTGCCAACGATGCAGCCATGA<br>AGGTTGCACGCCACTACCATGCCCTGCTTGGGCGCACGCGCCGCTCACTTGTTGTTAGTTTACGTGGCTCG<br>TATCACGGGCTTACTTTCGGTGGCTTCGCACTCACCGGAGAGGACCTTGGGCAACGCTTATACGGGGTTGA<br>CCAGCGTTTAGTACGCCATGTTCGCCCCAATGACACGGCCGAGCTCAATGCGCTGGTGTCGCGCGCGGCGA<br>AACAAATCGCGGCAGTGGTTGTTGAGCCCGTGGTAGGCACTGGCACTATTCCGCTTACCGACGAATACGTG<br>GCGGAGTTGCTCCGTTTACGTGCAGAACACGGGTTCCTTTTAATTGCAGACGAGGTCGCTACGGGCTTTGG<br>CCGCACCGGAAGCTTCTTTGCGTCGCAGCGCTGGCCAGAGCAACCAGACCTGTTAATTACCAGCAAGGGTC<br>TCACCAATGGTACTTGTCCAGCAAGCGCCGTGATCGTGTCTCAACGTGTTGCCGATGCGTTTACCGAGCAC<br>GACGCCGTTCTCAGTCACGCGGAGACACAAGGTGCAACGCCTCTGACCTGCGCCGCAATCTCAGCAACCAT<br>TGCAGAAATGCGCCGCCTGGACGCCGTTGCAGCGGGCCAGGCACTTGGTGAAAAGTTAGGGGCCGGTATTG<br>CGGAGCTTATGGCGGAGATGCCGCAAATTATTGGTACTACCGGCGTAGGTTGTTTCCGCTCTTTGCGCATT<br>GCCGACGCTAGCGGGGCCCCGCTCCCGCAACATCGTGTACCAGCATTAGTCGCAGCGATCCGTGACGCTGG<br>TGCCATTGTACATCCGGGACCTAGCGGTGTCCAATTAGTGCCGGCACTCACTTACAGCGATGCCGAGCTTG<br>CTGAGTTGTTAGATTGTGTGCGTCGTGGTATCCTTGCACACCACGAGGCCGAGCAGGCAGCTACGCCTGCT<br>GAGGCCATCGCA |
| *tfDapM* (codon-optimized) | ATGGCGGCGACCACTTTACTTCCACCTGGCCGTGCTGGACAACTTGTGGCTGAACTTGGGGACCGTGCACG<br>TCTCTGCGATTTGTACGGTCCCCACGGGGCCCCAATATACCATGACATGTCTCTTCGAGACACAGGCGAGG<br>TAAGACACCTTGTAGGCCTCGTGCGCCCACACCCGGGGCCGGTACTTGATTTAGCCGCGGGGAGTGGCCGC<br>ATAACCCTTCCTCTGTTAGCGCTTGGACGTGAAGTTACGGCTCTCGACCTGTCGGCAGACATGCTTGCTCT<br>GTTACGCCAACAATTGGACAGAGCCCCGGCTCATCTGAGAGAACGGTGCACCGTTGTCCAGGCGGATATGG<br>CGGACTTCCGGCTGCCTAGACGTTATGCTGTCATTGTACTTGGTACTACCAGCATCAGCTTGTTAGACCGT |

| | |
|---|---|
| | GCAGGCCGCGCCGGTCTTTACCGCTGTGTGGCGGAGCATCTTGCTGATGGTGGACGATTCTTCCTTACAAC<br>GTTGGACCGAGGGGCTGACTCCCCGGAAGAAGAAGTGGAAATAACGGCAACAGGCGCGTCCGGCACAGCAT<br>ATCGACTTTATGAGCATTGGCCTGCAGGCGCGGACGTTCGCACAATCACCGTTCTGCCATCTGAATTGCCT<br>GACGGCCCGGTGCCGGTTTGTACAGGACACGTTCGTGTTTTACCCCCTGACAGATTGGCCGCTGAGCTGAC<br>TCAAGCAGGTTTTACAGTCCAACAACGCCGCACACCTCTTGAAGACGCTGGTCGCCATCGTGTTACCTTGA<br>TTGAGGCGGAAATTCAGAGA |
| *tfDapY* (codon-optimized) | ATTGATGTGACACGTACGCTTGGACCAGCCGGTGGAATTGCCCGCGTATTTGGGCTGCAACCCCCAACACC<br>TCGCGACCCATTGTGGACGGCTGGTGTTGAACTCCGTTCCCCCGGTCCAGACGATAACAGCCTGCCTTTGT<br>CTGCACGTATGGTAGGCGCCTGTGGCTTAGCGCGCAATGATGCGCTTGTCCGCGGGGCCGGCGAAGCCGTA<br>GAGCGTTTCGCGTTACATCCAGGCCGCGCCGCCGGTCCAGTACGCGGTCGCCGTACTGGCCTGCCTGCGCC<br>TGCCGTTGAATTTCACCGTCCAGAGGTCGCGCTCGGAGCGCCCCATGCCGCCGATCTTGAGCTTCACTGGT<br>ATCCCGCACGCCGCCTTCGTGATGGAGCTAAAGTTATGGTGCCGGCTCCGCTCGTGGACTGGCCCTGTGAT<br>CCTCGTGAGAGCGCCTATTTCGACCCCGGTCCTAGTGGCGCAGCGAGCGGTTTAGGGTGGGAGATGGCCTT<br>ACGTGCGGCCTTACTGGAAGTGGTAGAGCGTGACGCTGTGATGGTCGCCTGGCAACGCGGCCTGCGCGCGT<br>ATCGCGTGGCCGATCCAGCCGCGCTTGCGGCCCCTGGTGGTGATGGGGAGCGCAGCCGTCGCGCGCTGGGT<br>GAATTGTGGCAGCGCGCCCGTCGTGAGGGGATGACCCCCTTCTTAGTTGTTTTGCCTACGGCGCATCCGGC<br>CGTTTGGTGCTGTGTGGGCGGGTTAGATGACGCCGATGGTGACCTGGCGGCGATTGGGTGCAAGGCAAGCG<br>ACCGCCCTTGGGAAGCTGCCCTGGGCGCGTTCCAAGAGGCGTGGCAGGTTCGTAGTGTTTTGCTGCGTGCG<br>CGTGAATCGGGAGTGCGTCCAGTTGCTGCCGAGGAGATTGTGGATGAGGATGATCGTATCGCCTATTTGGC<br>GTCCCCGGCCGGAGCCGCGGCAGTTCGCGAGTGGCTTGCAGGGTGTGACGGGGACGCAGCGCAAGTAGCGT<br>GGACGCCCGGCGTTGGAACAGAGGAGCTCGTTGCTGCAGTATTAGCCGACGGTGGTGACCCATTAGTGGTC<br>GACTTAGCAGCGCGTCTCCCTGAACCACTGCGCGCGATGGGCTGGCACGCGTTAAAAGTAGTACCCGTAGG<br>CTACCAACAACTGCGCATGGACGAGCGCCACACCTGGAGTTGGAATCGCGCCCGTTTGGCCTCGGCCGTTG<br>AACGTACCGGATTAGCGGCTCGTCGTACCGCCGACGACGACGCCCCGCCACATCCGCTTCCA |
| *saDapA*$_1$ | ATGAGCACCGATCTGGATCTGCAGTTCGAGGAACTCGAGACGCTGGACGCTCTGTGGAACTGGGCCGAGTT<br>CGGCTCGGGCTTCGGGGCCGGCGCGACCGCCTTCTCCGTCGGCGTTGCCGTCTTCCTCACC |
| *saDapB* (codon-optimized) | ATGACAACTACCCCTCTTCGCCCGCAACCAGAGCGTAGCCCATTGCAAGCCACCGTTCACCTTGAGCGGTG<br>GCTGCGCGGTCACTACGATGCAGTCTCTGATACCGCCTTTGAAGTGGTTGCAGAGGCCGGCGCTGACTTAC<br>CGGCACTGGCGGCGAGTGGCTTGCTCGACGCGGATGGTGTCATCTTCGCCGAGCCTGGGGTAGCTGACTCC<br>CTGCCTGTCCCTGCTGTGGCATTAGAGGGATCTGTTCTCAACTGTGGGGACGACCTTGTTGTGGGTGGGGA<br>GTTCCATATACAGGTCTTTGACTACGTGGCATTAGGTTTCGTGGCTTTGGTAGGTCCGACCGTAGTCCGCA<br>TTACAGGAGAGGATGATTTGACCGCGTTTCTTGCTGACGCGGACCTGGCAGTCTCTGATGGAAGTCTGCCT<br>CAATGGTTACTGAACCCGGGAGTAGTTCTGGCAGATGCACCGGCCTTGGCCGGTATGGCCCCAACGGGAGT<br>GGCGCGTCTCTACGTCACTGCGGATGGTATGGTCCGTACAGCACCGGGCGGAGCTGACTTGGCTCCTCTCC<br>GGGATGGGGCAGCAGCCATACGTGCAGCGGTGGCCACGCATGCCACAGATCCCTCACTGGATGGTGTGCTG<br>CCAAGCAGAACCTTAGAGCGTGCGCGTGCGGAGCGTCCTTGGCTGCCCCGCTACTTACAAGCTCTGGACGC<br>GGTGCGGGCTCTCTCTCGCGTCGCTGGTGGTCCTGTGCGTATTAGTGGTTTCGGGATGAGACTCTGCCCCC<br>AAGCCCCGGCTGAACCTGTTGAGTCGGCGGCGCTCCCATTAATCGCCCGCGCCGACGATGGCACGTGTTTC<br>CTGCTCTACCCGAATGGTGGCCGTGCTTTCAAAGTCGGGCAGGATGTGGCTATCCTGGTCGAGGCTAAGAT<br>TGCATGTGGTGACCAGCGACAAGCTGACGCAGTGGCGGCCGCCGCTCTTGGAGTCGGAGCCGACGAAGTTC<br>CGGGACTGTACAGCCGGTTGCGCTTGCCTGAGATGCGCGCCGCT |
| *saDapC* | GTGACCACCGTGGTACCGCTCGCGGTTCGTGCCGAGCCCGCCACTGCCTTTCAGCCGGCCCGGTGGAGTCG<br>GCCCACAGAACTCCTGATCGGCCAGGGCGCCGCCCGCGAGGCCGTTCGCCAGGCCCGCTCCGATGGCACGA<br>CGGTGCTCGTGACCGACACCAGCCTTCCCGAGGGCCTCGTCGACCGTCTGGTCGGCCCAGCCGGGAAACGG<br>CTGCGGCGCCTTGGGCTGCACCCGGGGTCCATCACCCTCGCGAGCTCGGGGGAAATCGCCGACTCGCTGGG<br>TGATGTCCGGCACGTGGTGGCGATCGGCGGCGGCAGTGTGCTGGACGCCGCCGCCGTGGCCCGGGCGATCC<br>TCAGCCGCCCTGAGCTGTCCACTCTCATACGCCGCCCCACCATGCCTGGCCTGATCCTCTGTCCGCCCGGG<br>CGAAGGCCGGTGCCCCGGTTCACCGCCGTCCCCACCACCGTGGGAACCGCCGCCGAGGTCAGTTCCCTGGC<br>GACCGTCCTCGCCTGCGACCGGCGAAAGCTCGTCACGGGTGACGCGCTCACCCCGGGCACCGCCGCCCTCG<br>ACCCTGCGGCCACGGCGGGCCTGCCCCGGCGGCTGCTGCTGGAAGGGGTGCTTGAGGCGATGATGCGGCTG<br>CTCAACATCTGTGCCCTGCCGCCGGTCGGACAGTGCCCGGGTGCCTCCGATGCCGAACTGCTGACGCTGCT<br>GGCCCAGTTGGGCCGTTCGGGGGCTGAGGCGGCCAACGCGCGGTACGACGTGCCCGCCCCGCTGCGCATCG<br>ACATCGCCCTGCTCAGCGCACGCACCGTGCTTGGCTGGACGGCGCTGGGGCGTGACCCGTTCGGCGGGAAG<br>ATCTGGTACCTGTCCAACGAACTGTCCACGCTCGCCGGCGTACGCAAGATGACTGCCACCGTCTCGGTGGC<br>ACCCGTGGTGTGGTCCAGGCTGCTCGCTGGTGACACCCGGTTCGGCGACCCGGCCCGGCTGCACACCGCCT<br>GGCAGGCTCTGTGGCGCGCCCTGGGACCCATGCGGTTGCCCACCGACCCCGTCGCCGGATTCCGCGCCCTG<br>GTCGCCCGCGTGGGACGTACCGGCGCTGCGGAAGACCGAGGCCCCGCCGCCCCAGGAGCTGGCCTGGCGAAC<br>CGCACAAGCCTGGGGCGGGAGCCAGCCCATGCTGGGCGCCTTCAGCGCCGCCGAACTGACCGCCCTCTACC<br>GGGAGATCATCGACCAGCCGCGACCGCGGAGGAGCCAGCTG |
| *saDapC* (codon-optimized) | ATGACCACTGTAGTACCCTTGGCAGTTAGAGCGGAACCAGCCACTGCATTCCAACCAGCTCGTTGGAGCCG<br>CCCCACTGAATTGCTTATTGGGCAAGGAGCCGCCAGAGAGGCGGTTCGGCAAGCTCGATCAGACGGAACAA<br>CTGTCTTGGTAACTGATACGTCACTTCCTGAGGGCTTGGTTGACCGACTGGTTGGGCCAGCAGGAAAACGT<br>TTACGTCGGCTCGGATTACATCCAGGTTCTATCACCTTAGCATCGAGCGGCGAGATAGCTGACTCTCTGGG |

| | |
|---|---|
| | AGACGTACGACATGTTGTTGCGATCGGTGGTGGCTCTGTCCTGGACGCCGCGGCGGTGGCTCGAGCAATCT<br>TATCTCGGCCTGAACTGAGCACATTGATCCGCAGACCCACAATGCCGGGGCTGATTCTTTGCCCGCCAGGT<br>CGCCGTCCGGTCCCTCGATTCACAGCAGTGCCCACAACGGTTGGCACCGCAGCTGAAGTCTCGAGCTTGGC<br>CACGGTCTTAGCGTGTGATCGCCGGAAGCTGGTGACGGGCGACGCTTTAACCCCTGGAACCGCAGCGCTGG<br>ACCCGGCTGCAACAGCTGGGTTACCACGCCGTCTGTTACTGGAAGGAGTTCTTGAGGCTATGATGCGTTTG<br>CTTAACATTTGTGCTTTGCCCCCAGTTGGGCAATGCCCTGGCGCATCCGATGCCGAGCTCTTAACGCTGCT<br>GGCCCAATTGGGTCGCTCAGGAGCGGAAGCGGCAAACGCACGTTATGACGTTCCCGCGCCGCTCCGTATTG<br>ATATAGCCTTACTTAGCGCGCGAACCGTACTCGGATGGACTGCACTCGGGCGTGACCCTTTCGGCGGCAAG<br>ATTTGGTACCTGTCTAATGAGTTGAGTACGTTAGCTGGTGTGCGCAAGATGACCGCAACGGTTTCCGTGGC<br>ACCAGTGGTGTGGAGCAGATTACTCGCAGGCGACACTCGGTTTGGTGATCCGGCTCGTTTGCACACTGCTT<br>GGCAAGCCTTGTGGAGAGCGCTCGGTCCGATGCGTTTACCAACTGACCCCGTTGCGGGGTTTCGTGCTTTA<br>GTCGCCGCCTGGGATGTGCCAGCATTACGTAAGACCGAAGCGCCCCCACCCCAAGAGCTGGCGTGGCGTAC<br>CGCGCAAGCGTGGGGCGGATCCCAACCAATGTTGGGTGCATTCTCCGCTGCTGAGCTCACTGCATTATACC<br>GGGAGATAATCGATCAACCGCGTCCACGTCGTTCACAGTTG |
| *saDapD* | CTGTGGGAACTGCTCGTACCGCCCGGGGCGTACGCCAAGCCGGAGCGGCGGGCCGTGGCCGCGGCCGGCGC<br>CAGGCTCCGCTTCGACGACGGCTCCGAGGTCCTGGACGCCACCAGCGGCCTGTGGAACGTCAACCTCGGCT<br>ACGGCAACACGGCCATCGCCGACGCAGTGGACCGGGCCATGCGCGAGGCGTCGTACCTGACGCTCTTCCGC<br>TACTCGCACACCTACGCCCTGGAGGCCGCCCGCGCCCTGACAGAAGCCGCCGGGCCGGCGGTCTTCCGCCG<br>GGTGATCTTCTCCACCTCCGGCAGCTCAGCCAACGACCTGGTGATGAAGGTCGCCCGCCACCACGCCCTGC<br>TCACCGGCGAGCCCCAGCGCCGCCTGATCGTCGGCTTCAAGGGCAGCTACCACGGGCTGACCTATGGGGCG<br>TTCTCCCTGTCCGGTGAGGAACTGGGCCAGCAACTGTACGGAGTGGACACACGGCTCGTCCGCCATGTCGA<br>CGCCGGCCGCCCGGAGGACCTCGAACGGCTGATGCGGCGCGAGGGCCACCGGGTGGCGGCGGTGGTGTGCG<br>AGCCGGTGTTCGGCTCCGGCGCCCACGAACTGCCGCGCACGACGCTGGAGGCGCTGCTGGAACTCCGCCGG<br>GAGTACGGCTTCCTGCTCGTCGCCGACGAGGTGGCCACGGGATACGGCCGTACCGGGCCGCTGTTCGCCAG<br>CTCCGCCTGGGCGCAGGCCCCGGACCTGATGGTCACCTCGAAGGGGCTGACCAACGGCACCTGCGCCGCCT<br>CCGCGGTTCTGGCGTCGCACGCCGTCTGCGACGCCTTCGAGCGGGCCGACGCGCTGCTGGTGCACGGGGAG<br>ACACAGGCCGGGACACCGCCGACATGTGCCGCGATCCTGGCGACGCTGGCACAGTTCACGGAGTTGTCCGC<br>CCTGGAGTCCGGCGCCCGGGTGGCGCGCGCGCTCGACGCGCTGCTGGCGGAGCTCGTCGCCGAGTTGCCCT<br>CCGTCACCGCCGTCACCGGCCGGGGCTGCTTCCGCGGGATCCAGCTGGCCGAGCCCGACGGCACCCCGTAC<br>AGCGCCGAGCGGGTGCAGCAGGCCATCGCCGCGATCCGCGGCCATGGGACGCTGGCCCACCCGGGCCCGGG<br>GAGCGTACAGCTCGTCCCGCCGCTGACCCTCACCGACGAGGAGGCGCACGAACTGGGCACGGCCGTACACA<br>AGGGACTGGTGGAGGCGCGA |
| *saDapD* (codon-optimized) | ATGTGGGAACTCTTAGTGCCGCCAGGTGCCTATGCTAAACCGGAACGCCGCGCAGTAGCAGCAGCAGGTGC<br>GCGGCTTCGCTTTGACGACGGCAGCGAGGTTCTGGACGCTACGTCAGGGTTATGGAATGTTAACCTTGGAT<br>ACGGGAATACGGCTATTGCAGATGCTGTCGACCGCGCAATGCGTGAAGCGTCCTACCTGACTCTCTTTCGA<br>TACTCTCATACCTACGCACTTGAGGCAGCAAGAGCTCTGACTGAAGCTGCCGGTCCCGCAGTGTTCAGACG<br>AGTGATCTTTAGCACAAGTGGATCTAGCGCCAACGATCTTGTTATGAAGGTGGCGCGCCATCACGCTCTCT<br>TGACCGGTGAGCCGCAGCGACGACTGATTGTCGGCTTCAAGGGTTCGTATCATGGCTTGACTTATGGAGCC<br>TTCTCATTGTCCGGTGAGGAACTCGGGCAACAACTGTACGGAGTAGATACTCGGTTGGTTCGGCACGTGGA<br>CGCTGGTCGCCCCGAAGATCTCGAGCGCCTTATGAGACGCGAGGGGCACCGTGTTGCTGCGGTTGTTTGTG<br>AACCCGTGTTTGGAAGCGGGGCCCACGAGCTTCCGCGCACCACTCTGGAAGCGTTGCTTGAGCTTCGCCGC<br>GAGTATGGCTTCCTGCTCGTCGCTGACGAAGTTGCGACTGGTTACGGGAGAACCGGTCCGCTCTTCGCATC<br>TAGTGCGTGGGCCCAAGCACCAGATTTAATGGTAACCAGCAAGGGTTTAACAAACGGCACATGCGCAGCAT<br>CAGCAGTTCTGGCGTCACATGCCGTGTGCGATGCGTTTGAGCGCGCTGATGCCCTTCTTGTTCACGGTGAG<br>ACACAAGCGGGCACCCCACCGACATGCGCAGCTATACTGGCCACCCTTGCTCAGTTTACCGAGCTCTCGGC<br>ACTTGAGTCAGGTGCTCGCGTTGCTCGTGCTTTGGACGCACTTTTAGCGGAGTTGGTAGCCGAGCTCCCAT<br>CCGTAACTGCGGTTACCGGTAGAGGTTGTTTCCGCGGTATCCAATTAGCTGAGCCGGATGGACGCCCTAT<br>TCGGCGGAAAGAGTGCAGCAAGCTATTGCAGCGATCAGAGGGCACGGGACACTGGCCCATCCGGGTCCCGG<br>GTCAGTGCAACTCGTCCCACCCCTCACGTTAACAGACGAAGAAGCACACGAGTTAGGTACCGCTGTACATA<br>AAGGACTTGTGGAAGCACGT |
| *saDapM* | ATGAACGGCCGGACAGCGATCAGCACGCCGTCGGTGGCGCCGCGGACCGGTGCCCGGCACATCCCGCCGGG<br>CACCGCCACCCGAATCGCCGCGGCGCTGGGCGATGACCTCAGGATCGCCGACCTGTACGACGATCTCGGCG<br>GCCCGGTCTACCACCTGGTCACCAGCCATGACCGCAGCGAGATCACCGAGGTCCTCACCGCACTGCGCGGC<br>ACCGGGGGGCCCGTGCTGGAACTCGCCTGCGGTTCCGGAAGGCTGACCCTGCCGCTGCTGCTCGCCGGGCA<br>CCAGGTCACCGGGGTGGACACGTCCGACTCGCTGCTGGGCGAACTCACCGCCCAGGTCGGGCGGCTCCCTG<br>CGCACCGCGCCGCGGACCGTCTGACCGTGCTCCGCGAGGACATGACCGCGCTCGATCTGGACACGCGTTTC<br>GCGGCCGCCCTGCTCGGCATCACGACCATCGGGCTGGTGCCCCCCGGCGAACGGCCCGGCTTCCTGCGCAA<br>CGTTCACAGGCACCTGCTGCCCGGTGGCCGCTTCCTGGTCACCGTGGAGACGCCGCGGGTCCGACCCGGAG<br>AGGAGGAGATCCACACCTTCGTCGCCGACGACGCCGTGATCACCCTCATCTCGTACGTGGAGCCGGGACTC<br>GCCGCCCGCCACGTAGCCGCGCTGAGCGTCGCCCGAGGTGCCGGCCCTCTGCTGCTCACCTCGTTCGTGCA<br>CCTGGTGACCCCTGACCGGCTGAGCGCCGAACTCGTCGCGGCGGGCTTCGAGATCGAGTCCGTCAGCCCGG<br>TGCGAGATCCCCTCGCCGGCACCGCCGACGGGAACACCCTTGTCATCGCCAGGAGGCCGTCG |

| | |
|---|---|
| *saDapY* | GTGATCGACGTCCTCGCGACCCTCGCCCCTGAGGCACAGCTCACCCGCGGCTGGCAGTTGCACCCCCCGGG<br>CACCGGCGAGCCCCAGTGGCGGGGACTGGTTCATCTGGCCGGGCAGCAGGGAGAGCCGGTGTCCGGCGGCC<br>CGTCCCAGCCGATCTCCCACGAGGTGGTCGGCGCGTACGGGGTGAACCGTGCGGACGTGCTGGTCCGGGCC<br>ACGGGAGAAGCCGTCGAACGGTTCGCGCTCCAGCCGCTGCCCGGAGAGGGCCGCAGGGGCCCATCCACCGC<br>CCTGGACGGCGACGTACTCGGCTTCCACGAGGCCGGGCTGGGCGCCGAGGACTGCGTGGGACAGGTGCGGA<br>CCTGGTACCCGGCGCGCAGGCTGCTCGACGGCCACCGGATGTGGGTGCCGGCGGGGCTCGTCGACCACCCA<br>CCCCGGCCGGAGGACAGCGACGGATTCGACCCCACCCCGTCCGGGGCGGCCGCCGGCGCAGGGCCCGGGAT<br>GGCGCTGCGCTCGGCGCTGCTTGAGGTCATCGAACGGGACGCGCTGACCGTCGCCTGGGTTCGTGCGCTGA<br>CCCTGCGGCGCATCGACCCGGAGGTCTCGATCGCGGCAGCCCCCGGCTCTCCGTCCTGGCGGAGGTTCGGC<br>TCCGCCCTGCGCCACGCCCGGGCAGGCGGATGGGAACCGGTACTGGCCGAGGTGCCCACCGCGGTGGACGG<br>CGTGACGTGCGTGGTGGGCATCCTCCTGGGGGACCAGGGCGCGGCCGTCGGATGCAACGCCTCCACCGACC<br>CTGGCCAGAGCCTTGCGGGCGCGCTGCAGGAGGCCCTCCAGGTGATGTCGTGTCTGCGGCTGGTCGGTCCC<br>CGGTACGCGGACGAGCCCGTGCCCGGCATCGTCACCGAAGACATGGACCGGGCACGGTGGTTCGCATCGGC<br>TGAGGGGAGGGAGACGGTGAGGCGGTGGTGCGACGGGTTCGAGCCCGGCTCCGCGGCCCTGCGGCTCACCC<br>CCGGGCAGCCGTCGACGGAGGAAATCCTGGCGGGCCTGCATGCCCAGGGCGCACGTCCCCTGGCCGTCGAC<br>CTGAGCCACCGGTTGCCGCCGGCCGTGCGCGCCATGGGCTGGGCCGTGGTGAAGGTGATACCCGCCGGACT<br>GCAGCCGCTGCGCATCGACGAACGGTGCTCATTCGGCTGGAACCGCTTCCGGCTGGAGACCGCCGAGGCAC<br>GTACGGGGCTTCGGGCACGGTCCGGCCATTCGCCGCACCCCCACCCGCTGATA |

**Table S4. Key protein sequences from the *tfDap* and *saDap* BGCs.**

| Name | Sequence |
|---|---|
| *Tf*DapA$_1$ | MQDITPQPNEVALELEMQELEAMEAPGFWTGAGVGALISTSFAISLT |
| *Tf*DapB | MDKTELRRIKRHLMSWSTGQFEVGPANQDSCSTIVVADPAHLADLLDTDLVGDNTVVLAPEGHGADPSRVVGFA<br>GTLDEPNAEMSIGYDFFLQTQDYATSPFMSVLGPTLVRVTGPDDLELFLADADRAREEGVFPDIAVVPAVRIAD<br>LPGLGAGPGVDGPQLRLYVNAEGEISTSIGGCSLGRVGDTLASLRANWDRRNAASTAPCAVCLGAVLPEADRTR<br>ELLARPWLGRYLAAVDGLRMLATREIVGLRVSGFGQRLNPALDHHPDSADAATPLLFWNDESGYVYEPASGRTF<br>QVNLAAATAVEALLVTGSAEQASEYASPRSLAAVDRFFTESGVTLTGIGSEVAV |
| *Tf*DapC | MTLLPTAPPRPMYGCGHLTEWLRQKGATTVALVTDQTLTDTPILATVRARIVAAGVQVRFFVLPDAGNLDSVLR<br>LADRFADCDLVVGVGGGSVIDQVKLAATLCGNPKARGRLTVAQRSGTIILPVTVERRTPLLAIPTTLGTGTELS<br>AVACLVYPEGKRIISGLCLRPDAAVLDPLATETLPVELVAEGTLEALFRTVSPYIGDHADLPEQDTLVEAVAAE<br>LVRLGDEVNRLRRRGEPISPQIRLRIAELSGRSQLGDLHAGRKPYAVKGWAIANELSWELSLRKMPAIAALLPP<br>LWRRIADGDHRIGSAPRLVRIWQRLAENSPGLPDDPAEGIAALIERWQIDHRITADPAQLDTVAARAVHAWGAG<br>LPVLGGLTQAEIRQLLADAVADRALAPGTPQTSVA |
| *Tf*DapD | MSHTTSTALWPYLLPPSAHGDDSICAVSARGHRVRFADGRELLCGASGLWNANLGYGNPAIAQAVAQALHDASY<br>LSAFRYENVYARRAAADLIEVSGPEHYSRVLFSSSGGAANDAAMKVARHYHALLGRTRRSLVVSLRGSYHGLTF<br>GGFALTGEDLGQRLYGVDQRLVRHVRPNDTAELNALVSRAAKQIAAVVVEPVVGTGTIPLTDEYVAELLRLRAE<br>HGFLLIADEVATGFGRTGSFFASQRWPEQPDLLITSKGLTNGTCPASAVIVSQRVADAFTEHDAVLSHAETQGA<br>TPLTCAAISATIAEMRRLDAVAAGQALGEKLGAGIAELMAEMPQIIGTTGVGCFRSLRIADASGAPLPQHRVPA<br>LVAAIRDAGAIVHPGPSGVQLVPALTYSDAELAELLDCVRRGILAHHEAEQAATPAEAIA |
| *Tf*DapM | MAATTLLPPGRAGQLVAELGDRARLCDLYGPHGAPIYHDMSLRDTGEVRHLVGLVRPHPGPVLDLAAGSGRITL<br>PLLALGREVTALDLSADMLALLRQQLDRAPAHLRERCTVVQADMADFRLPRRYAVIVLGTTSISLLDRAGRAGL<br>YRCVAEHLADGGRFFLTTLDRGADSPEEEVEITATGASGTAYRLYEHWPAGADVRTITVLPSELPDGPVPVCTG<br>HVRVLPPDRLAAELTQAGFTVQQRRTPLEDAGRHRVTLIEAEIQR |
| *Tf*DapY | MIDVTRTLGPAGGIARVFGLQPPTPRDPLWTAGVELRSPGPDDNSLPLSARMVGACGLARNDALVRGAGEAVER<br>FALHPGRAAGPVRGRRTGLPAPAVEFHRPEVALGAPHAADLELHWYPARRLRDGAKVMVPAPLVDWPCDPRESA<br>YFDPGPSGAASGLGWEMALRAALLEVVERDAVMVAWQRGLRAYRVADPAALAAPGGDGERSRRALGELWQRARR<br>EGMTPFLVVLPTAHPAVWCCVGGLDDADGDLAAIGCKASDRPWEAALGAFQEAWQVRSVLLRARESGVRPVAAE<br>EIVDEDDRIAYLASPAGAAAVREWLAGCDGDAAQVAWTPGVGTEELVAAVLADGGDPLVVDLAARLPEPLRAMG<br>WHALKVVPVGYQQLRMDERHTWSWNRARLASAVERTGLAARRTADDDAPPHPLP |
| *Sa*DapA$_1$ | MSTDLDLQFEELETLDALWNWAEFGSGFGAGATAFSVGVAVFLT |
| *Sa*DapB | MTTTPLRPQPERSPLQATVHLERWLRGHYDAVSDTAFEVVAEAGADLPALAASGLLDADGVIFAEPGVADSLPV<br>PAVALEGSVLNCGDDLVVGGEFHIQVFDYVALGFVALVGPTVVRITGEDDLTAFLADADLAVSDGSLPQWLLNP<br>GVVLADAPALAGMAPTGVARLYVTADGMVRTAPGGADLAPLRDGAAAIRAAVATHATDPSLDGVLPSRTLERAR<br>AERPWLPRYLQALDAVRALSRVAGGPVRISGFGMRLCPQAPAEPVESAALPLIARADDGTCFLLYPNGGRAFKV<br>GQDVAILVEAKIACGDQRQADAVAAAALGVGADEVPGLYSRLRLPEMRAA |
| *Sa*DapC | MTTVVPLAVRAEPATAFQPARWSRPTELLIGQGAAREAVRQARSDGTTVLVTDTSLPEGLVDRLVGPAGKRLRR<br>LGLHPGSITLASSGEIADSLGDVRHVVAIGGGSVLDAAAVARAILSRPELSTLIRRPTMPGLILCPPGRRPVPR<br>FTAVPTTVGTAAEVSSLATVLACDRRKLVTGDALTPGTAALDPAATAGLPRRLLLEGVLEAMMRLLNICALPPV<br>GQCPGASDAELLTLLAQLGRSGAEAANARYDVPAPLRIDIALLSARTVLGWTALGRDPFGGKIWYLSNELSTLA<br>GVRKMTATVSVAPVVWSRLLAGDTRFGDPARLHTAWQALWRALGPMRLPTDPVAGFRALVAAWDVPALRKTEAP<br>PPQELAWRTAQAWGGSQPMLGAFSAAELTALYREIIDQPRPRRSQL |
| *Sa*DapD | MWELLVPPGAYAKPERRAVAAAGARLRFDDGSEVLDATSGLWNVNLGYGNTAIADAVDRAMREASYLTLFRYSH<br>TYALEAARALTEAAGPAVFRRVIFSTSGSSANDLVMKVARHHALLTGEPQRRLIVGFKGSYHGLTYGAFSLSGE<br>ELGQQLYGVDTRLVRHVDAGRPEDLERLMRREGHRVAAVVCEPVFGSGAHELPRTTLEALLELRREYGFLLVAD<br>EVATGYGRTGPLFASSAWAQAPDLMVTSKGLTNGTCAASAVLASHAVCDAFERADALLVHGETQAGTPPTCAAI<br>LATLAQFTELSALESGARVARALDALLAELVAELPSVTAVTGRGCFRGIQLAEPDGTPYSAERVQQAIAAIRGH<br>GTLAHPGPGSVQLVPPLTLTDEEAHELGTAVHKGLVEAR |
| *Sa*DapM | MNGRTAISTPSVAPRTGARHIPPGTATRIAAALGDDLRIADLYDDLGGPVYHLVTSHDRSEITEVLTALRGTGG<br>PVLELACGSGRLTLPLLLAGHQVTGVDTSDSLLGELTAQVGRLPAHRAADRLTVLREDMTALDLDTRFAAALLG<br>ITTIGLVPPGERPGFLRNVHRHLLPGGRFLVTVETPRVRPGEEEIHTFVADDAVITLISYVEPGLAARHVAALS<br>VARGAGPLLLTSFVHLVTPDRLSAELVAAGFEIESVSPVRDPLAGTADGNTLVIARRPS |
| *Sa*DapY | MIDVLATLAPEAQLTRGWQLHPPGTGEPQWRGLVHLAGQQGEPVSGGPSQPISHEVVGAYGVNRADVLVRATGE<br>AVERFALQPLPGEGRRGPSTALDGDVLGFHEAGLGAEDCVGQVRTWYPARRLLDGHRMWVPAGLVDHPPRPEDS<br>DGFDPTPSGAAAGAGPGMALRSALLEVIERDALTVAWVRALTLRRIDPEVSIAAAPGSPSWRRFGSALRHARAG<br>GWEPVLAEVPTAVDGVTCVVGILLGDQGAAVGCNASTDPGQSLAGALQEALQVMSCLRLVGPRYADEPVPGIVT<br>EDMDRARWFASAEGRETVRRWCDGFEPGSAALRLTPGQPSTEEILAGLHAQGARPLAVDLSHRLPPAVRAMGWA<br>VVKVIPAGLQPLRIDERCSFGWNRFRLETAEARTGLRARSGHSPHPHPLI |

**Table S5. Strains used in this study.**

| Organism and Strain Identifier | Phylum | Source / Reference |
|---|---|---|
| *E. coli* BL21(DE3) | Pseudomonadota | Novagen |
| *E. coli* DH10β | Pseudomonadota | Novagen |
| *Thermobifida fusca* DSM 43792 | Actinomycetota | DSMZ |
| *Streptomyces azureus* ATCC 14921 | Actinomycetota | USDA Agricultural Research Service Culture Collection |
| *E. coli* DH5α | Pseudomonadota | NEB |
| *E. coli* Tuner (DE3) | Pseudomonadota | Novagen |
| *E. coli* DH5α γpir | Pseudomonadota | William W. Metcalf |
| *E. coli* WM6026 | Pseudomonadota | William W. Metcalf |
| *Streptomyces albus* J1074 | Actinomycetota | USDA Agricultural Research Service Culture Collection |
| *E. coli* GB05-dir | Pseudomonadota | Youming Zhang |

**Table S6. Primers used in this study.**

| Name | Sequence 5'-3' |
|---|---|
| F_*saDapY*Rxn | GAACCTGTACTTCCAATCCGGATCCATGATCGACGTCCTCGCG |
| R_*saDapY*Rxn | AGTGGTGGTGGTGGTGGTGCTCGAGTCATATCAGCGGGTGGGG |
| F_*saDapD*Rxn | GAACCTGTACTTCCAATCCGGATCCATGAACCCTCGTGAGCCG |
| R_*saDapD*Rxn | AGTGGTGGTGGTGGTGGTGCTCGAGTCATCGCGCCTCCACC |
| F_*saDapC*Rxn | GAACCTGTACTTCCAATCCGGATCCATGACCACCGTGGTACCG |
| R_*saDapC*Rxn | AGTGGTGGTGGTGGTGGTGCTCGAGTCACAGCTGGCTCCTCC |
| F_*saDapM*Rxn | GAACCTGTACTTCCAATCCGGATCCATGAACGGCCGGACAGC |
| R_*saDapM*Rxn | AGTGGTGGTGGTGGTGGTGCTCGAGTCACGACGGCCTCCTG |
| F_*saDapA*$_1$Rxn | GAACCTGTACTTCCAATCCGGATCCATGAGCACCGATCTGGATCTG |
| R_*saDapA*$_1$Rxn | GGTGGTGGTGCTCGAGTGCGGCCGCTCAGGTGAGGAAGACGGC |
| F_*saDapA*$_1$_petduet | GTACTTCCAATCCATGAGCACCGATCTGGATC |
| R_*saDapA*$_1$_petduet | CATTATGCTGCGGCCGCTCAGGTGAGGAAG |
| F_petduet_*saDapA*$_1$ | CTTCCTCACCTGAGCGGCCGCAGCATAATG |
| R_petduet_*saDapA*$_1$ | GATCGGTGCTCATGGATTGGAAGTACAGGTTCTCAG |
| R_petduet_*tfDapB* | CATATGTATATCTCCTTCTTATACTTAACTAATATACTAAGATGGGG |
| F_petduet_*tfDapD* | CTCGAGTCTGGTAAAGAAACCG |
| F_*tfDapB*_petduet | GTATATTAGTTAAGTATAAGAAGGAGATATACATATGGATAAAACCGAGCTGCG |
| R_*tfDapB*_*tfDapC* | TCATGGTATATCTCCTTGAATCTCACACCGCAACCTCTG |
| F_*tfDapC*_*tfDapB* | GTGAGATTCAAGGAGATATACCATGACCCTTTTACCTACCGC |
| R_*tfDapC*_*tfDapD* | ACATTGTATAGCCTCCTCTTAGTCACGCGACCGAAGTC |
| F_*tfDapD*_*tfDapC* | GTGACTAAGAGGAGGCTATACAATGTCCCACACAACTAGCAC |
| R_*tfDapD*_petduet | GCAGCGGTTTCTTTACCAGACTCGAGTCATGCGATGGCCTCAG |
| F_*saDapB*_petduet | GTATATTAGTTAAGTATAAGAAGGAGATATACATATGACAACTACCCCTCTTCG |
| R_*saDapB*_*saDapC* | TCATGGTATATCTCCTAGACCGTCAAGCGGC |
| F_*saDapC*_*saDapB* | TTGACGGTCTAGGAGATATACCATGACCACTGTAGTACCCTTGG |
| R_*saDapC*_*saDapD* | ACATTGTATAGCCTCCTCTTAGTTACAACTGTGAACGACGTGG |
| F_*saDapD*_*saDapC* | GTAACTAAGAGGAGGCTATACAATGTGGGAACTCTTAGTGCC |
| R_*saDapD*_petduet | GCAGCGGTTTCTTTACCAGACTCGAGTTAACGTGCTTCCACAAGTCC |
| F_*saDapA*$_1$_V19M | CCGCCTTCTCCATGGGCGTTGCCGTCTTCCTCACCTGAGCGGC |
| R_*saDapA*$_1$_V19M | ACGGCAACGCCCATGGAGAAGGCGGTCGCGC |
| F_*saDapA*$_1$_V19K | CCGCCTTCTCCAAAGGCGTTGCCGTCTTCCTCACCTGAGCGGC |
| R_*saDapA*$_1$_V19K | ACGGCAACGCCTTTGGAGAAGGCGGTCGCGC |
| F_*saDapA*$_1$_V19E | CCGCCTTCTCCGAAGGCGTTGCCGTCTTCCTCACCTGAGCGGC |
| R_*saDapA*$_1$_V19E | ACGGCAACGCCTTCGGAGAAGGCGGTCGCGC |
| R_*tfDapA*_core | GGATCCGGATTGGAAGTACAGG |
| F_*tfDapA*_core | GGCTTCTGGACTGGCG |
| F_*tfDapA*_core_short | GGCGTCGGAGCATTGATATCAAC |
| R_*tfDapA*_full | CTTAAGCATTATGCTTTAGGTCAAGCTTATCGCGAAGGATG |
| F_*tfDapA*_full | CTGTACTTCCAATCCATGCAGGATATCACGCCGC |
| R_pETDuet_*tfDapA* | CGTGATATCCTGCATGGATTGGAAGTACAGGTTCTCAG |
| F_petDuet_*tfDapA* | ATAAGCTTGACCTAAAGCATAATGCTTAAGTCGAACAG |
| F_*tfDapA*_shorter | ATATCAACATCCTTCGCGATAAGCTTGAC |

| | |
|---|---|
| R_*tfDapB*_confusion | GCATGCTACCGCTACCCGATCCCACCGCAACCTCTGACC |
| F_*tfDapB*_confusion | CTTCTGGACTGGCTGAGATCCGGCTGCTAACAAAG |
| R_*tfDapA*_confusion | TAGCAGCCGGATCTCAGCCAGTCCAGAAGCC |
| F_*tfDapA*_confusion | GGTGGGATCGGGTAGCGGTAGCATGCAGGATATCACGCCGC |
| F_*DapA*$_{eng}$_overlap1 | TCACCACGAGAACCTGTACTTCCAAGCGATAAGCTTGACCTAAAGC |
| R_*DapA*$_{eng}$_overlap1 | CGGAGCTCGAATTCTTCGTGATACGAGTCTGCGCGTCTTTCAG |
| F_*DapA*$_{eng}$_overlap2 | GAAGCTCCGGGCTTCTGGACTCACCATCACCATCACCACGAGAACCTGTACTTC |
| R_*DapA*$_{eng}$_overlap2 | CATCGCCTCTAATTCTTGCATTTCCAGCTCAAGGGCGGAGCTCGAATTCTTCGTG |
| F_*DapA*-Glucagon_1 | CGTGCCCAAGACTTTGTTCAATGGTTGATGAATACCTAAAGCATAATGCTTAAGTCGAAC |
| F_*DapA*-Glucagon_2 | GTACCTTTACGTCGGATTACTCGAAATATTTGGATAGTCGTCGTGCCCAAGACTTTGTTC |
| F_*DapA*-Glucagon_3 | GGCGATGGAAGCTCCGGAACATTCACAAGGTACCTTTACGTCGGATTACTCG |
| R_*DapA*-Glucagon_3 | CCTTGTGAATGTTCCGGAGCTTCCATCGCCTCTAATTCTTGCATTTC |
| R_*DapA*$_{eng}$_overlap3 | AGCCCGGAGCTTCCATCGCCTCTAATTCTTGCATTTCC |
| F_*DapA*$_{eng}$_overlap3 | ATTAGAGGCGATGGAAGCTCCGGGCTTCTG |
| F_*fusC*_pet28 | CTTTAAGAAGGAGATATACCATGGTTGGCTGTATCAGCC |
| R_*fusC*_pet28 | TGGTGCTCGAGTGCGGCCGCGCTACGATTATCGCCACCATTC |
| R_pet28_startcodon | CATGGTATATCTCCTTCTTAAAGTTAAAC |
| F_pet28_NotI | GCGGCCGCACTCGAG |
| F_GFP_pETDuet | CATGGGCAGCAGCAAAGGAGAAGAACTTTTCACTGGAG |
| R_GFP_pETDuet | GAGTCCAGAAGCCTTTGTATAGTTCATCCATGCCATGTG |
| F_pETDuet_GFP | TGAACTATACAAAGGCTTCTGGACTCACCATC |
| R_pETDuet_GFP | GTTCTTCTCCTTTGCTGCTGCCCATGGTATATCTC |
| F_pBE44 | TTACCAATGCTTAATCAGTGAGGCACC |
| R_pBE45 | ATCTTTATAGTCCTGTCGGGTTTCG |
| R_pBE44-*saDap* | CTCGGCCTGGACTTCGTCGACCTGTACCTGATCCACTGGAACTTATATCGTATGGGGCTG |
| F_pBE45-*saDap* | GCGACGGGCAGCTTCTCGATCCGGATCGCCGACGCCTTGGACGCTCAGTGGAACGAAAAC |
| F_*saDap*-g1 | AATTAATACGACTCACTATAGGGAATTTCTACTGTTGTAGATGTCGACCTGTACCTGATC |
| R_*saDap*-g1 | GATCAGGTACAGGTCGACATCTACAACAGTAGAAATTCCCTATAGTGAGTCGTATTAATT |
| F_*saDap*-g2 | AATTAATACGACTCACTATAGGGAATTTCTACTGTTGTAGATTCGATCCGGATCGCCGAC |
| R_*saDap*-g2 | GTCGGCGATCCGGATCGAATCTACAACAGTAGAAATTCCCTATAGTGAGTCGTATTAATT |
| F_*saDap*_kasO | CGGTGCTCATTTCTCCACCGTCCTTTCGTTGGTTGGTATGGCGATCCCGCTGGCCACGACTT TACAACACCGCACAGCATGTTGTCAAAGCAGAGACCGTTCGAATGTGAACACAATTGTGGCG GGATCGTTGTATATTTCTTGACA |
| R_*saDap*_kasO | AAAGTGCGTCTTTCACCTGGCCGGCCGGCTTACGCACACCGGAGCGGTGTCAATTGTTACCA ATGCTTAATCAGTGAGGCACC |

**Table S7. Vectors used in this study.**

| Name | Source | Resistance Marker |
|---|---|---|
| pET-28a-*tfDapA*$_1$_Nhis | this work | Kanamycin |
| pET-28a-*tfDapB*_Nhis | this work | Kanamycin |
| pET-28a-*tfDapC*_Nhis | this work | Kanamycin |
| pET-28a-*tfDapD*_Nhis | this work | Kanamycin |
| pET-28a-*tfDapY*_Nhis | this work | Kanamycin |
| pET-28a-*tfDapC*$^{K286A}$_Nhis | this work | Kanamycin |
| pET-28a-*tfDapC*$^{K286E}$_Nhis | this work | Kanamycin |
| pET-28a-(MBP)*tfDapA*$_1$ | this work | Kanamycin |
| pET-28a-(MBP)*saDapA*$_1$ | this work | Kanamycin |
| pET-28a-(MBP)*saDapC* | this work | Kanamycin |
| pET-28a-(MBP)*saDapD* | this work | Kanamycin |
| pET-28a-(MBP)*saDapM* | this work | Kanamycin |
| pET-28a-(MBP)*saDapY* | this work | Kanamycin |
| pUC-*saDapB*(CO) | this work | Ampicillin |
| pUC-*saDapC*(CO) | this work | Ampicillin |
| pUC-*saDapD*(CO) | this work | Ampicillin |
| pETDuet-(MBP)*saDapA*$_1$-*sazBCD* | this work | Kanamycin |
| pETDuet-(MBP)*saDapA*$_1$-*tfDapBCD* | this work | Kanamycin |
| pETDuet-(MBP)*saDapA*$_1^{V19M}$-*tfDapBCD* | this work | Kanamycin |
| pETDuet-(MBP)*saDapA*$_1^{V19K}$-*tfDapBCD* | this work | Kanamycin |
| pETDuet-(MBP)*saDapA*$_1^{V19E}$-*tfDapBCD* | this work | Kanamycin |
| pETDuet-(MBP)*tfDapA*$_1$-*tfDapBCD* | this work | Kanamycin |
| pETDuet-(MBP)*dapA*$_{eng}$-*tfDapBCD* | this work | Kanamycin |
| pBE44 | Enghiad et al.[8] | Ampicillin |
| pBE45 | Enghiad et al.[8] | Apramycin |
| pBE14 | Enghiad et al.[8] | Tetracycline |
| pBE45-*saDap* | this work | Apramycin |
| pET-28a-*fusC_CHis* | this work | Kanamycin |
| pGro7 | Takara Bio | Chloramphenicol |
| pET-28a-(MBP)*tfDapA*$_1^{T21S}$ | this work | Kanamycin |
| pET-28a-*tfDapB*$_{conf}$ | this work | Kanamycin |
| pET-28a-(MBP)*tfDapA*$_1^{1—21}$ | this work | Kanamycin |
| pET-28a-(MBP)*tfDapA*$_1^{7—21}$ | this work | Kanamycin |
| pET-28a-(MBP)*tfDapA*$_1^{12—21}$ | this work | Kanamycin |
| pETDuet-(MBP)*dapA*$_{eng}$-empty | this work | Kanamycin |
| pETDuet-(MBP)*dapA*$_{glucagon}$ -*tfDapBCD* | this work | Kanamycin |
| pETDuet-(MBP)*dapA*$_{lasso}$-empty | this work | Kanamycin |
| pEB1-mGFPmut2 | Addgene | Chloramphenicol |
| pETDuet-GFP$_{eng}$-empty | this work | Kanamycin |

**Table S8. Amino acid sequences of key peptides generated by SPPS for *in vitro* assays.** Amino acid sequences are shown in single-letter code. Lys residues were added in some cases for enhanced solubility and ionizability for mass spectrometry. X = norleucine

| Name | Sequence |
|---|---|
| *Tf*DapA$_1$ | MQDITPQPNEVALELEMQELEAMEAPGFWTGAGVGALISTSFAISLT |
| *Tf*DapA$_1^{(-12)—21}$ | KLEXQELEAXEAPKGFWTGAGVGALISTSFAISLT |
| *Tf*DapA$_1^{(-10)—21}$ | KXQELEAXEAPKGFWTGAGVGALISTSFAISLT |
| *Tf*DapA$_1^{(GS)6}$ | KXQELEAXEAPKGFWTGSGSGSGSGSGSAISLT |
| *Tf*DapA$_1^{(GS)6,W3A}$ | KXQELEAXEAPKGFATGSGSGSGSGSGSAISLT |
| *Tf*DapA$_1^{(GS)4}$ | KXQELEAXEAPKGFWTGSGSGSGSAISLT |
| *Tf*DapA$_1^{(GS)2}$ | KXQELEAXEAPKGFWTGSGSAISLT |
| *Tf*DapA$_1^{(-10)—4,17—21}$ | KXQELEAXEAPKGFWTAISLT |
| *Tf*DapA$_1^{(-10)—4,19—21}$ | KXQELEAXEAPKGFWTSLT |

**Table S9. Table of daughter ion assignments for HR-MS/MS analysis of *Sa*DapA$_1$-Miz.**

| Fragment Type | Fragmented Bond Number | Fragment Charge | Experimental *m/z* | Theoretical *m/z* | Mass Error (ppm) |
|---|---|---|---|---|---|
| y | 6 | 1 | 586.4084 | 586.4075 | 1.5 |
| y | 7 | 1 | 643.4299 | 643.4290 | 1.4 |
| y | 8 | 1 | 742.4985 | 742.4974 | 1.5 |
| y | 9 | 1 | 829.5306 | 829.5294 | 1.4 |
| b | 8 | 1 | 863.3823 | 863.3815 | 0.9 |
| y | 10 | 1 | 976.5988 | 976.5979 | 1.0 |
| b | 9 | 1 | 991.4410 | 991.4401 | 0.9 |
| b | 18 | 2 | 1019.9626 | 1019.9617 | 0.9 |
| y | 11 | 1 | 1047.6360 | 1047.6350 | 1.0 |
| b | 19 | 2 | 1076.5049 | 1076.5037 | 1.1 |
| y | 12 | 1 | 1148.6836 | 1148.6826 | 0.8 |
| y | 14 | 1 | 1276.7429 | 1276.7412 | 1.3 |
| y | 26 | 2 | 1293.6515 | 1293.6501 | 1.1 |
| b | 22 | 2 | 1319.6061 | 1319.6045 | 1.2 |
| b | 14 | 1 | 1638.7214 | 1638.7203 | 0.7 |
| b | 16 | 1 | 1852.8530 | 1852.8521 | 0.5 |

**Table S10. Table of daughter ion assignments for HR-MS/MS analysis of *Sa*DapA$_1$-Diz.**

| Fragment Type | Fragmented Bond Number | Fragment Charge | Experimental *m/z* | Theoretical *m/z* | Mass Error (ppm) |
|---|---|---|---|---|---|
| b | 16 | 1 | 1852.8534 | 1852.8521 | 0.7 |
| b | 17 | 1 | 1967.8813 | 1967.879 | 1.2 |
| y | 7 | 1 | 657.4457 | 657.4447 | 1.5 |
| y | 8 | 1 | 756.5144 | 756.5131 | 1.7 |
| y | 9 | 1 | 843.5464 | 843.5451 | 1.5 |
| y | 10 | 1 | 990.6149 | 990.6136 | 1.4 |
| y | 11 | 1 | 1061.652 | 1061.6507 | 1.3 |
| y | 12 | 1 | 1162.6998 | 1162.6983 | 1.3 |
| y | 13 | 1 | 1233.7377 | 1233.7355 | 1.8 |
| y | 14 | 1 | 1290.759 | 1290.7569 | 1.6 |
| y | 15 | 1 | 1361.7963 | 1361.794 | 1.7 |
| y | 16 | 1 | 1418.8179 | 1418.8155 | 1.7 |
| y | 17 | 1 | 1565.8853 | 1565.8839 | 0.9 |
| y | 18 | 1 | 1622.9078 | 1622.9054 | 1.5 |
| y | 20 | 1 | 1766.9606 | 1766.9589 | 1.0 |
| y | 21 | 2 | 957.5185 | 957.5173 | 1.3 |
| y | 21 | 1 | 1914.0287 | 1914.0273 | 0.7 |
| y | 24 | 2 | 1150.5981 | 1150.5968 | 1.1 |
| y | 25 | 2 | 1207.6195 | 1207.6182 | 1.0 |
| y | 26 | 2 | 1300.6598 | 1300.6579 | 1.5 |
| y | 27 | 2 | 1357.2019 | 1357.1999 | 1.4 |
| y | 28 | 2 | 1392.7201 | 1392.7185 | 1.2 |
| y | 29 | 2 | 1450.2339 | 1450.232 | 1.3 |

**Table S11. NMR assignments for Miz-modified peptide product.**

| Residue | Position | $\delta_C$ | $\delta_H$, mult. (*J* in Hz) |
|---|---|---|---|
| Ala-1 | 1 | -----, C=O | - |
| | 2 | 49.8, CH | 4.00, q (7.3) |
|  | N-2 | -----, NH | - |
| | 3 | 17.5, $CH_3$ | 1.39, d (7.2) |
| Ile-2 | 1 | -----, C=O | - |
| | 2 | 59.2, CH | 4.12, m |
|  | N-2 | -----, NH | 8.27, d (7.5) |
| | 3 | 37.4, CH | 1.73, overlapped |
| | $4_A$ | 15.4, $CH_3$ | 0.81, d (7.0) |
| | $4_B$ | 25.3, $CH_2$ | 1.37, overlapped<br>1.08, m |
| | 5 | 11.2, $CH_3$ | 0.77, t (7.6) |
| Ser-3 | 1 | -----, C=O | - |
| | 2 | 55.6, CH | 4.39, overlapped |
|  | N-2 | -----, NH | 8.22, d (6.9) |
| | 3 | 61.9, $CH_2$ | 3.71, m |
|  | O-3 | -----, OH | - |
| Miz-4 - aminoisopentane | 1 | overlapped | 4.59, overlapped |
|  | N-1 | -----, NH | 8.58, d (6.6) |
| | 2 | 40.0, $CH_2$ | 1.50, m<br>1.73, overlapped |
| | 3 | 24.8, CH | 1.59, m |
| | $4_A$ | 22.8, $CH_3$ | 0.85, d (6.7) |
| | $4_B$ | 21.1, $CH_3$ | 0.79, d (6.7) |
| 4-methylimidazol-2-ine | N-1' | -----, NH | 9.14, s |
|  | 2' | -----, C | - |
|  | N-3' | -----, N | 9.41, s |
| | 4' | 54.3, CH | 4.29, m |
| | 4'-$CH_3$ | 20.6, $CH_3$ | 1.18, d (6.4) |
| | 5' | 52.2, $CH_2$ | 3.92, m<br>3.42, m |

**Table S12. NMR assignments for Diz-modified peptide product.**

| Residue | Position | $\delta_C$ | $\delta_H$, mult. (*J* in Hz) |
|---|---|---|---|
| Ala-1 | 1 | -----, C=O | - |
| | 2 | 49.0, CH | 3.97, q (7.3) |
|  | N-2 | -----, NH | - |
| | 3 | 16.7, $CH_3$ | 1.36, d (7.2) |
| Ile-2 | 1 | -----, C=O | - |
| | 2 | 58.4, CH | 4.09, m |
|  | N-2 | -----, NH | 8.25, d (7.5) |
| | 3 | 36.4, CH | 1.70, overlapped |
| | $4_A$ | 14.6, $CH_3$ | 0.78, overlapped |
| | $4_B$ | 24.5, $CH_2$ | 1.33, m<br>1.05, m |
| | 5 | 10.2, $CH_3$ | 0.74, t (7.6) |
| Ser-3 | 1 | -----, C=O | - |
| | 2 | 54.7, CH | 4.36, overlapped |
|  | N-2 | -----, NH | 8.19, d (7.3) |
| | 3 | 61.0, $CH_2$ | 3.67, m |
|  | O-3 | -----, OH | - |
| Diz-4 - aminoisopentane | 1 | 44.7, CH | 4.61, overlapped |
|  | N-1 | -----, NH | 8.61, d (6.2) |
| | 2 | 38.4, $CH_2$ | 1.44, m<br>1.68, overlapped |
| | 3 | 24.2, CH | 1.57, m |
| | $4_A$ | 21.9, $CH_3$ | 0.83, d (6.8) |
| | $4_B$ | 20.2, $CH_3$ | 0.79, overlapped |
| 3,4-dimethylimidazol-2-ine | N-1' | -----, NH | 8.91, s |
|  | 2' | -----, C | - |
|  | N-3' | -----, N | - |
| | N-3'-$CH_3$ | 30.4, $CH_3$ | 2.99, s |
| | 4' | 60.3, CH | 4.14, m |
| | 4'-$CH_3$ | 17.1, $CH_3$ | 1.22, d (6.4) |
| | 5' | 49.4, $CH_2$ | 3.83, m<br>3.3, dd (7.8, 11.7) |

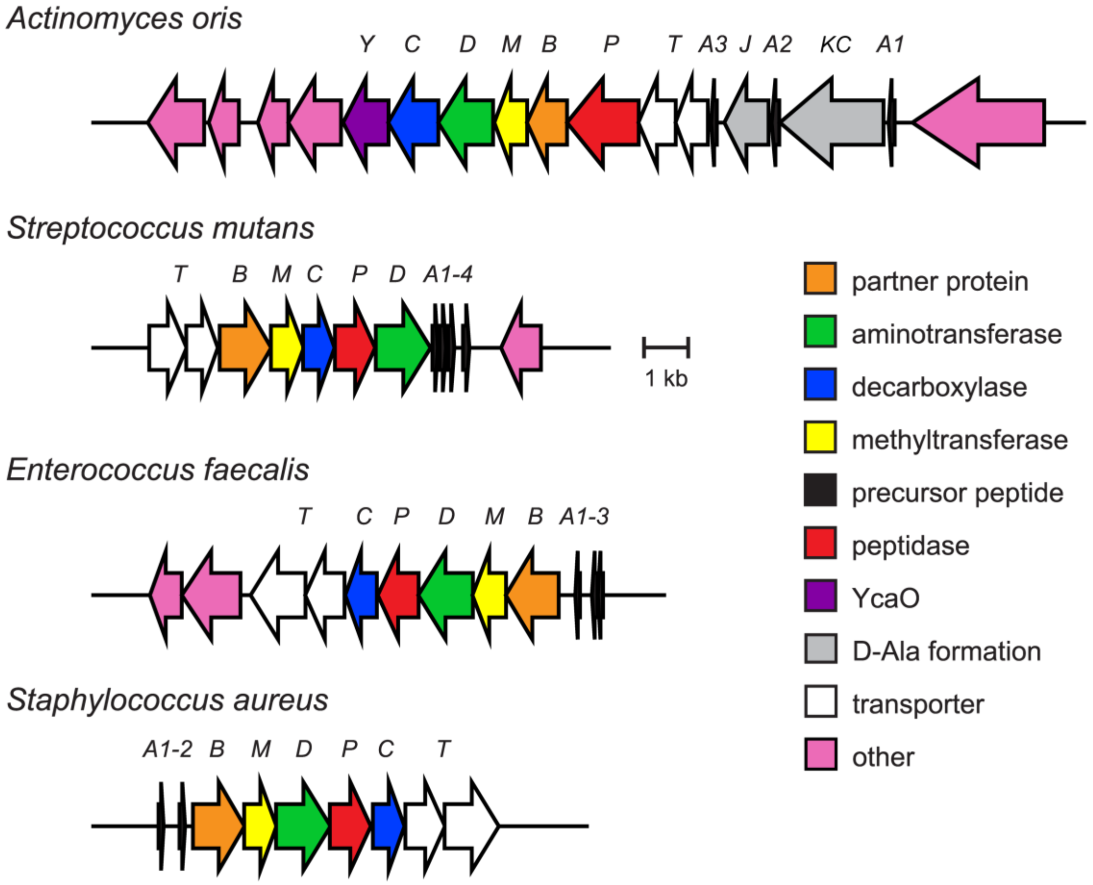

**Figure S1. Daptide BGCs from human-associated microbes.**

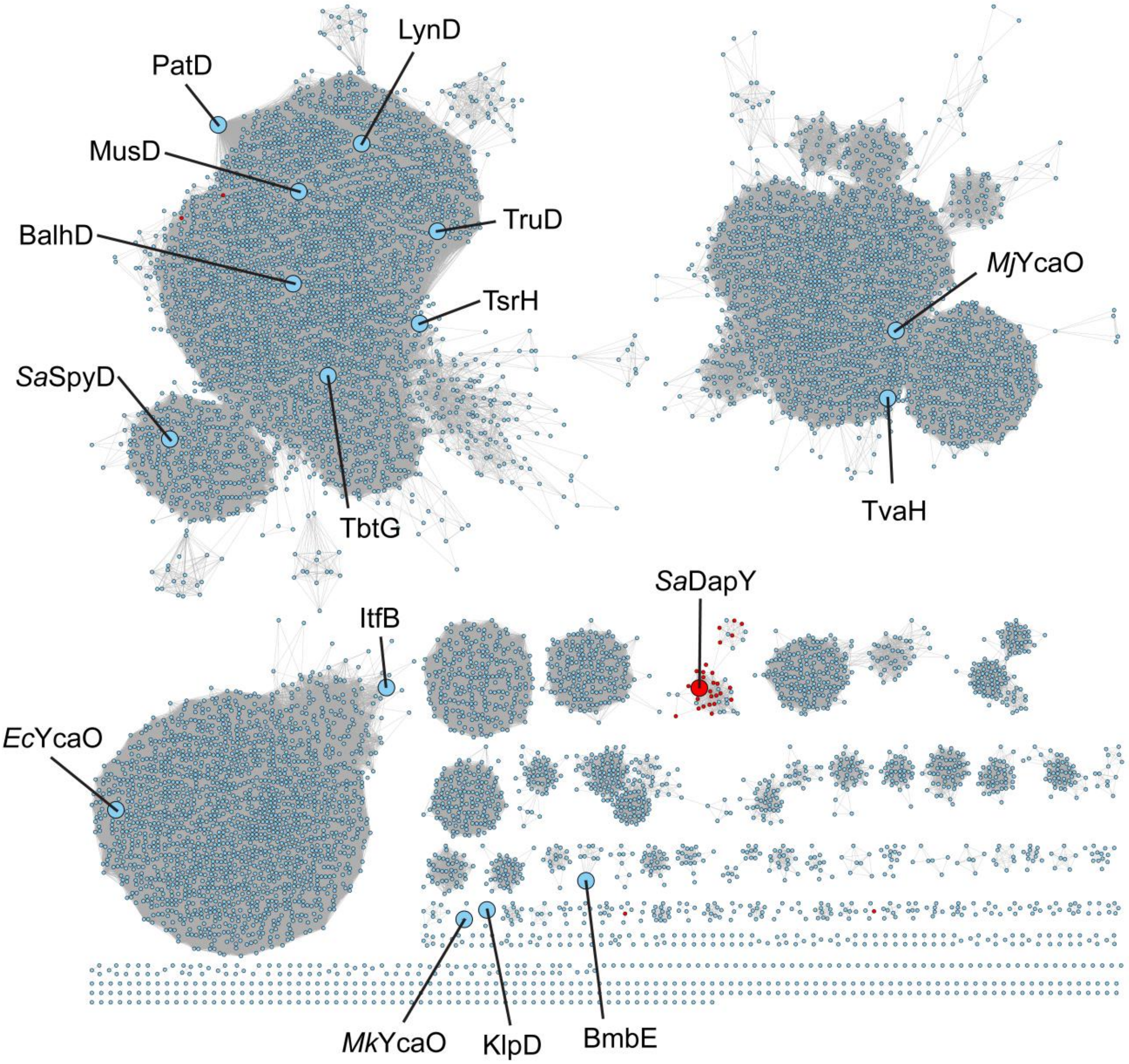

**Figure S2. Sequence similarity network for YcaO protein family.** The sequence similarity network (SSN) was generated using the Enzyme Function Initiative Enzyme Similarity Tool (EFI-EST)[3] and visualized using Cytoscape. All members of the YcaO protein family (Pfam ID: PF02624)[4] in UniprotKB were gathered by EFI-EST. Proteins with greater than 90% identity are conflated into a single node (RepNode 0.90). Edges with an alignment score less than 50 were removed. The layout was generated using the yFiles organic layout algorithm. Representative characterized YcaO proteins are indicated on the SSN by increased node size and labels.[9–15] YcaO proteins encoded within daptide BGCs are colored red.

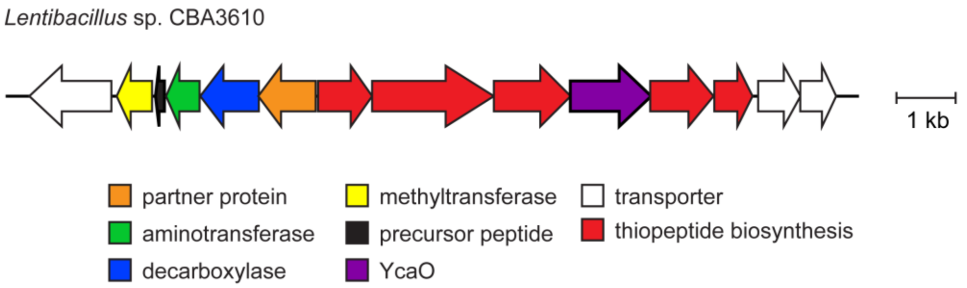

Precursor peptide: MTDSTNTVTESQIVDELFSDLEELEFVEIKEATALPETAATSGSATNGACSCCGSCSTCSSTT

**Figure S3. Representative hybrid thiopeptide-daptide BGC.**

```
heterocycle formation
TfDapY    DGDAAQVAWTPGVG---------TEELVAAVLADGGDPL-VVDLAARLPEPLRAMGWHALKVVPVGYQQLRM
SaDapY    EPGSAALRLTPGQPS--------TEEILAGLHAQGARPL-AVDLSHRLPPAVRAMGWAVVKVIPAGLQPLRI
TbtG      RESTVAFRSADLRDD--------LLEVIGRILAEGMDVI-VVDQTTPEH---RVHGFTCVKVLIPGMIPMTF
PatD      -NDYPKRWSDDIYTD--------VMACVEMAKVAGLETL-VLDQTRPD------IGLNVVKVMIPGMRTFWS
MusD      -QDYQQRWSDDIYED--------VMTCVEISKNAGLETL-VLDKTRPD------IGLNVAKVIVPEMPHYWL
SpSpyD    MAPYARRFTDPALGTVPVTAIAPTTPSALTSHLAGHRVI-TFDLTTEDI---RETPLRVARVLVSGLIPNQP
other reaction
TvaH      -QSLPTYLSADVMED--------TRRMIRNLQATGIEEVAVVDLSPKW------LPVSVVRVVVPGIESWAI
MjYcaO    -ADMPNNARYDLKKD--------LKFIKDKLSEFGFDKLIYVDLNK--------VGVDAVRVIIPKMEVYTI
ItfB      -TGENPTWGISNDSD--------LQYCFNIVEQLGFNIY-IRNNSI--------TDFPTYYIIIPGMNQVLF
BmbE      -HESTAFEADDGLPA-ATGPERLLRHLVDRLRESGVEAHHRVLTPP--------GPVTVVDVVAPGLEMFDK
                                            *                          ::

heterocycle formation
TfDapY    DERHTWSWNRA----------------RLASAVERTGLAARRTADDDAPPHPLP------------------
SaDapY    DERCSFGWNRF----------------RLETAEARTGLRAR----SGHSPHPHPLI----------------
TbtG      GHRNRRLDGLT----------------RPLVIPYRLGYRPAPLTPEALNPHPHPFP----------------
PatD      RYGPG----------------------RLYDVPVQLGWLKEPLAEAEMNPTNIPF-----------------
MusD      RMGAK----------------------RIYDVPVKMGWLSTPLTEEQMNPISVPI-----------------
SpSpyD    AAFAYLGCPRF-----------------AEQALSRRWRTSAPAAPTDFTLAPPPHM----------------
other reaction
TvaH      DRG-------------------------------RLGFRAAAVWEENLGLLRDALAEAAHRQ------EALR
MjYcaO    DRD-------------------------------RLSRRAFERVKKLYY-----------------------
ItfB      SRKAYEEKFRYGILGYMKSYPKLVNMSTAEVKELAQGIESRIQGGYTIDITHDYFLFNTHEGILNLDINIFL
BmbE      ARAGYPVLPTG-----------------------RLGERLRPRGDGGGARR---------------------
```

**Figure S4. Multiple sequence alignment of representative YcaO proteins.** Residues corresponding to the final 100 amino acids of *Tf*DapY in the YcaO domain are shown. The Pro-rich motif associated with YcaO cyclodehydratases is highlighted in yellow.[16]

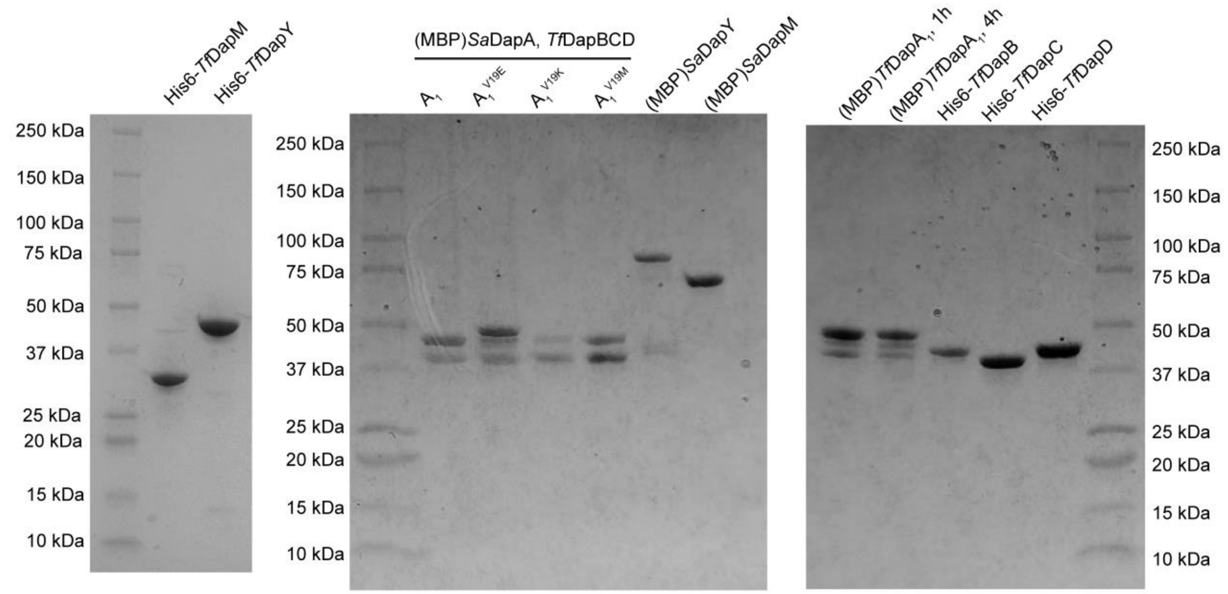

**Figure S5. SDS-PAGE gels for *Tf*Dap and *Sa*Dap proteins.**

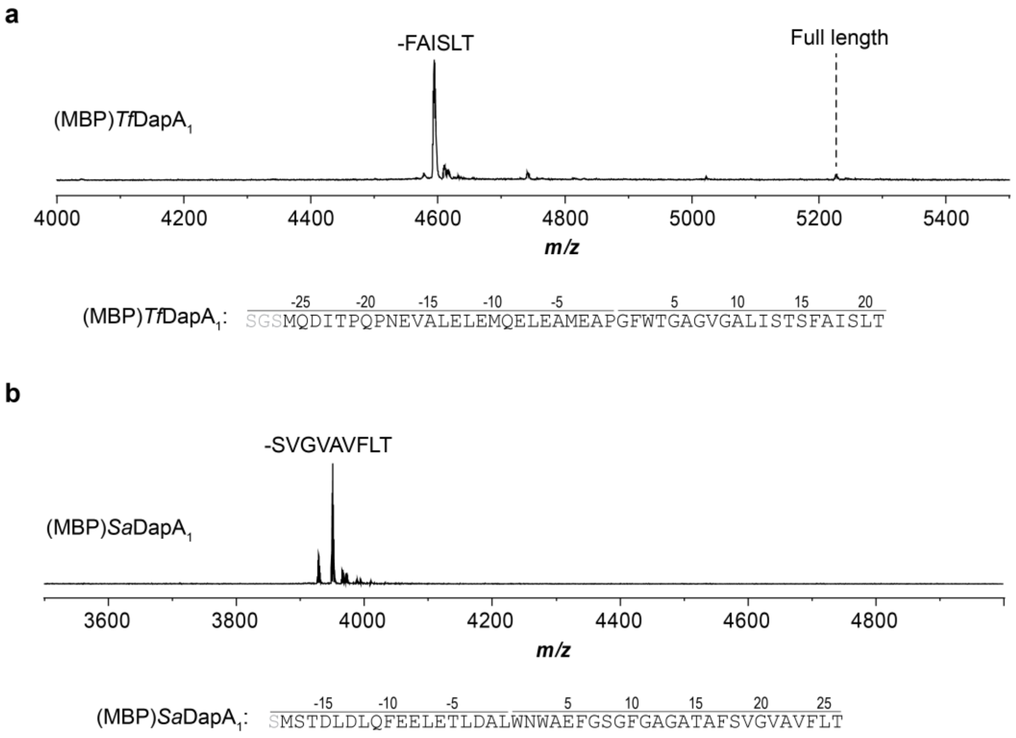

**Figure S6. MALDI-TOF-MS for MBP-tagged precursor peptides.** a, MALDI-TOF mass spectrum for expression of (MBP)*Tf*DapA1 following TEV proteolysis. b, MALDI-TOF mass spectrum for expression of (MBP)*Sa*DapA1 following TEV proteolysis. Spectra were collected after desalting by C18 ZipTip. Key expected masses: *Tf*DapA1, 5225 Da; *Sa*DapA1, 4800 Da.

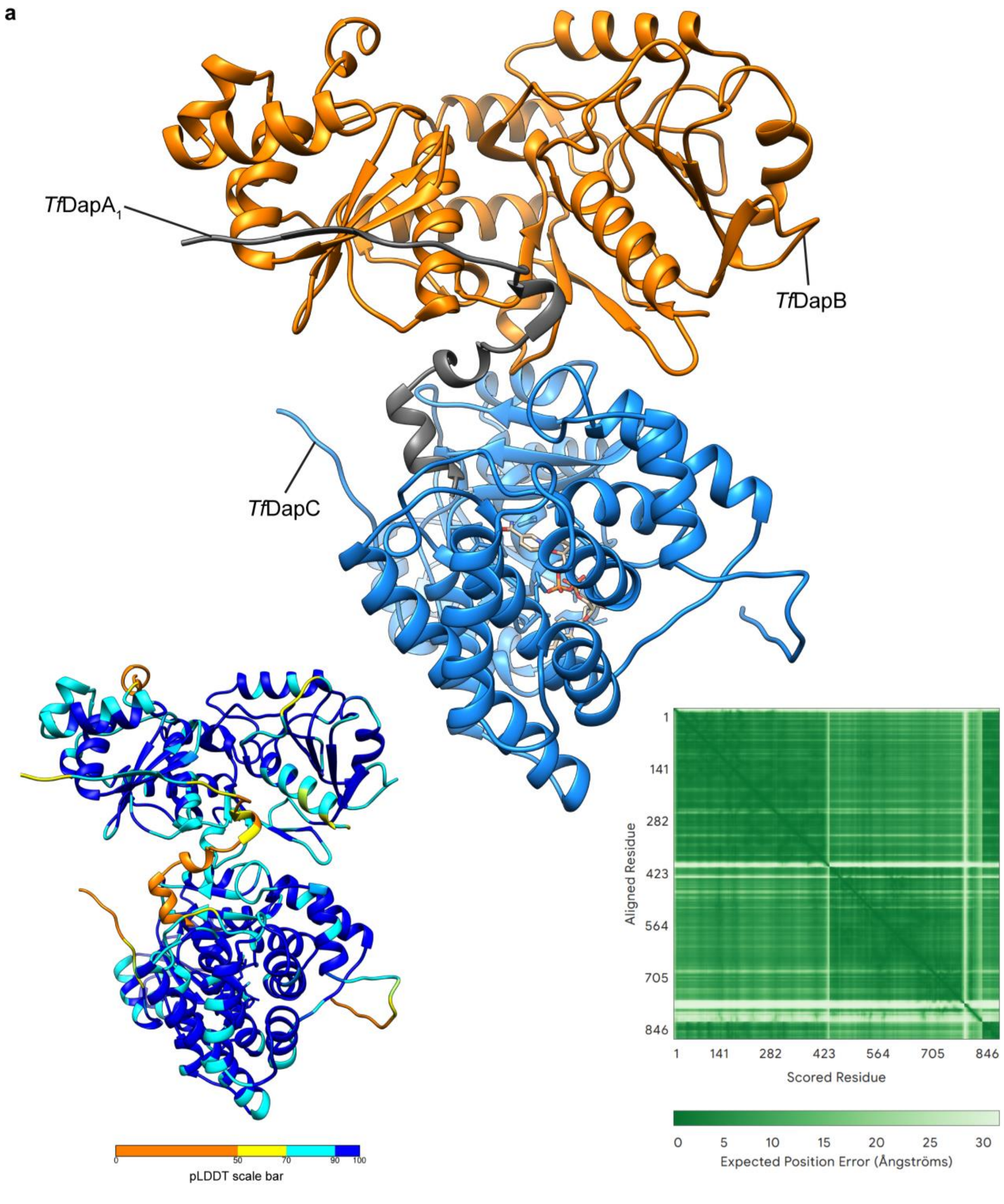

**Figure S7. AlphaFold predictions for *Tf*DapA$_1$BC with NAD$^+$.** a, AlphaFold[7] model of *Tf*DapA$_1$BC and NAD$^+$. pLDDT and pAE plots are also provided. b, Zoom-in of region surrounding Trp3 in *Tf*DapA$_1$ sequence. c, Zoom-in of *Tf*DapC active site with comparison to the active site of a crystallized Fe-ADH protein (PDB: 6C75).[17] The residues of *Tf*DapC which have putatively lost metal-binding capability are shown and colored green. The corresponding residues in 6C75 are also shown in green with the co-crystallized iron ligand.

b

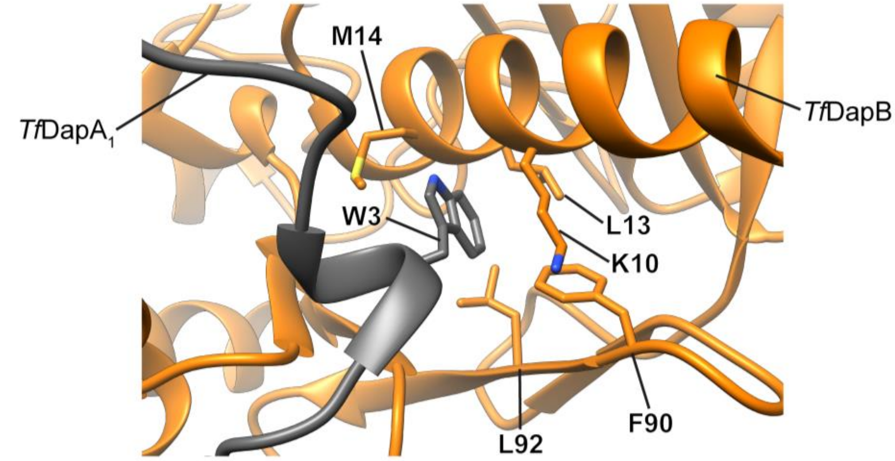

c

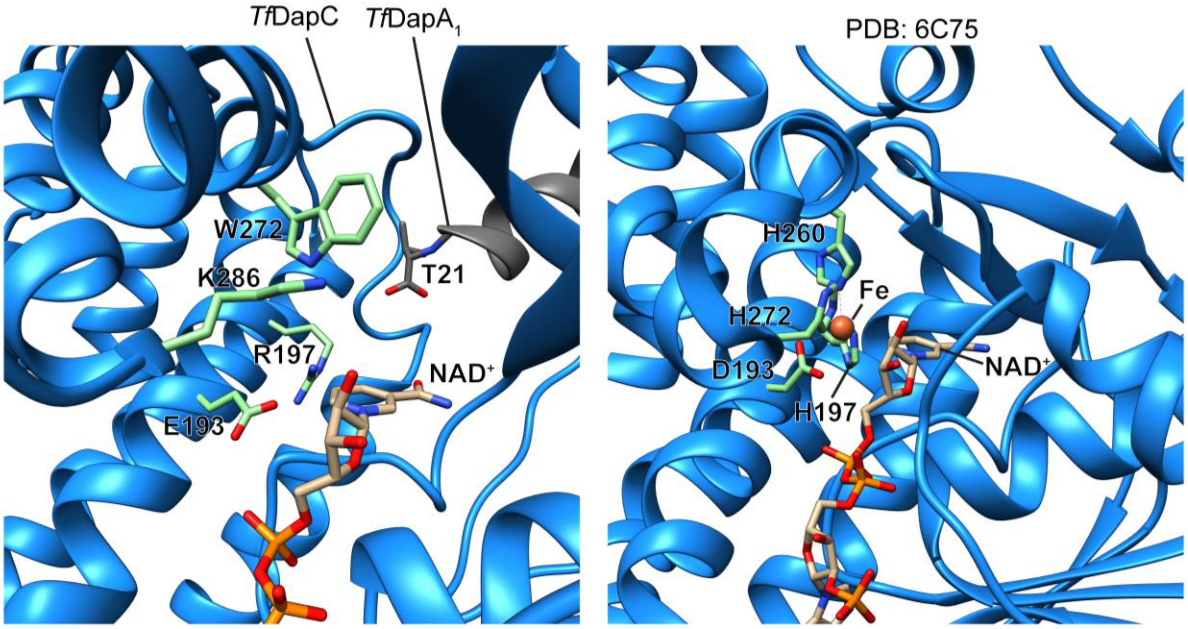

**Figure S7 (cont.). AlphaFold predictions for *Tf*DapA$_1$BC with NAD$^+$.**

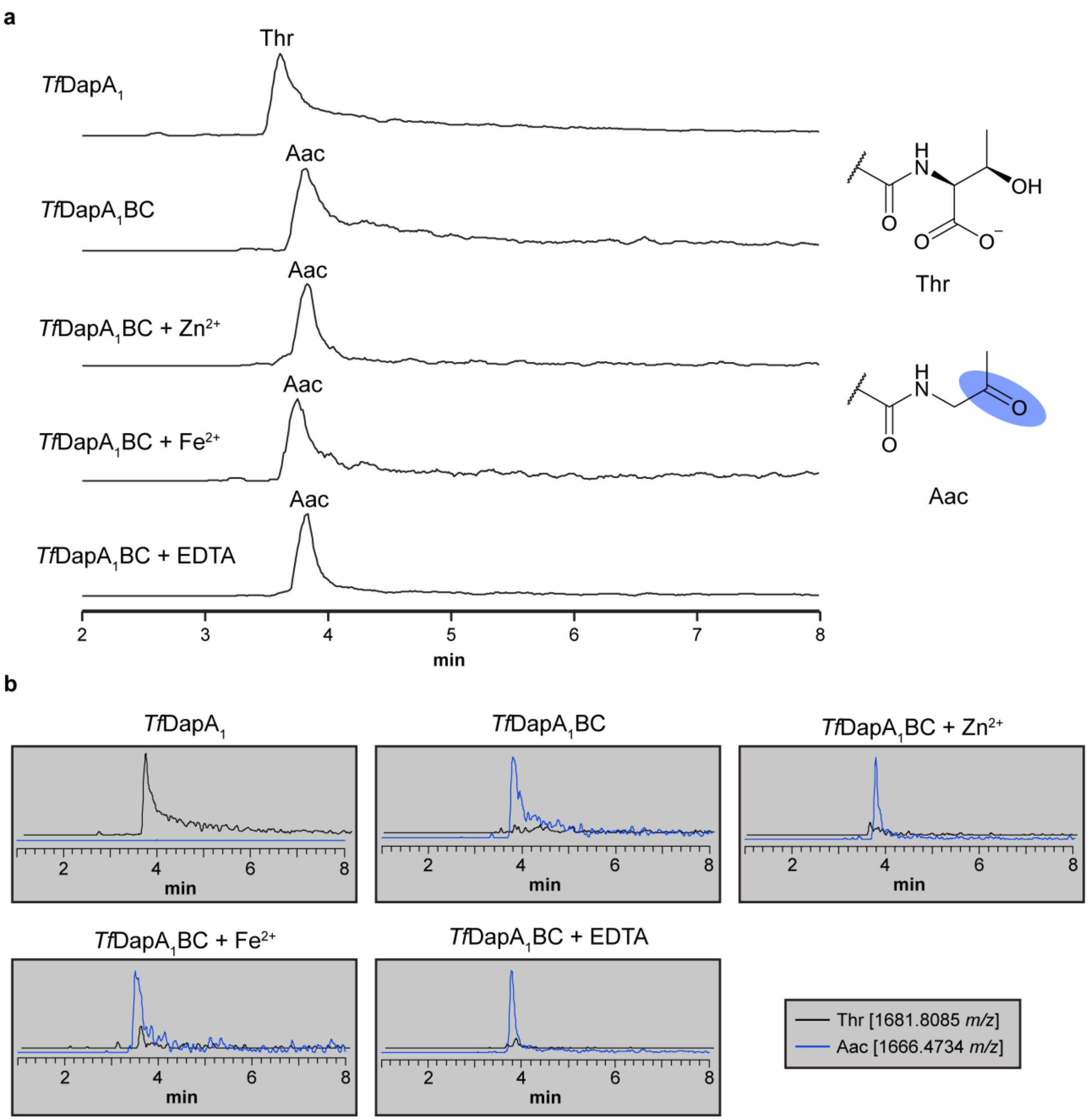

**Figure S8. LC-MS chromatograms for reconstitution of *Tf*DapBC.** a, Extracted ion chromatograms for the expected products of each reaction are shown. b, Extracted ion chromatograms corresponding to Thr and Aac (aminoacetone) are shown for each reaction. Extracted ions: Thr, 1681.8085 *m/z*; Aac, 1666.4734 *m/z*.

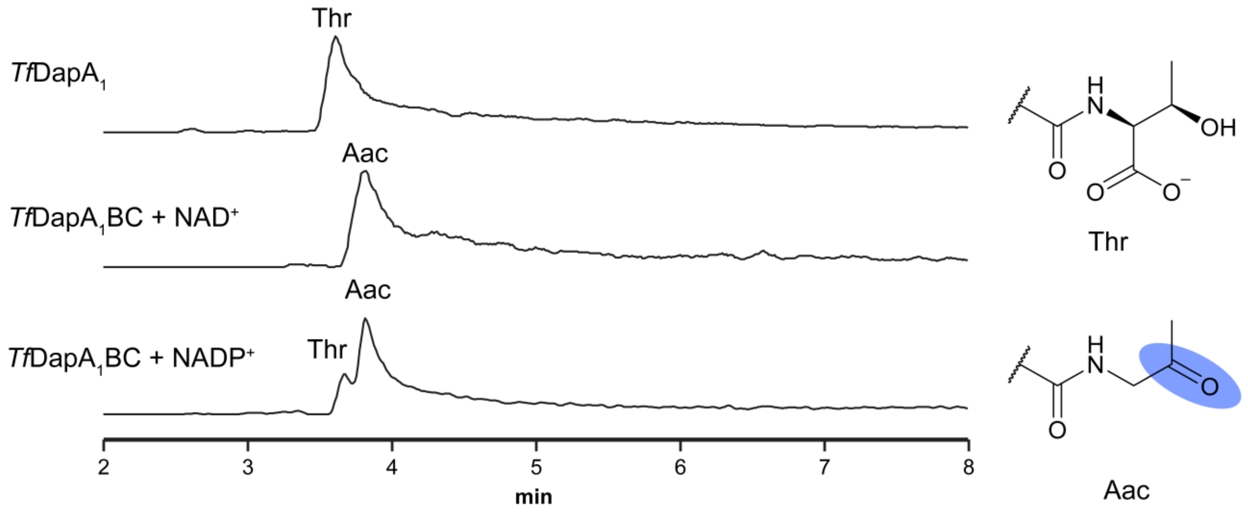

**Figure S9. LC-MS chromatograms for *Tf*DapBC reaction with NADP$^+$.** a, Extracted ion chromatograms for products of each reaction are shown. Extracted ions: Thr, 1681.8085 *m/z*; Aac, 1666.4734 *m/z*. Abbreviations: Aac, aminoacetone.

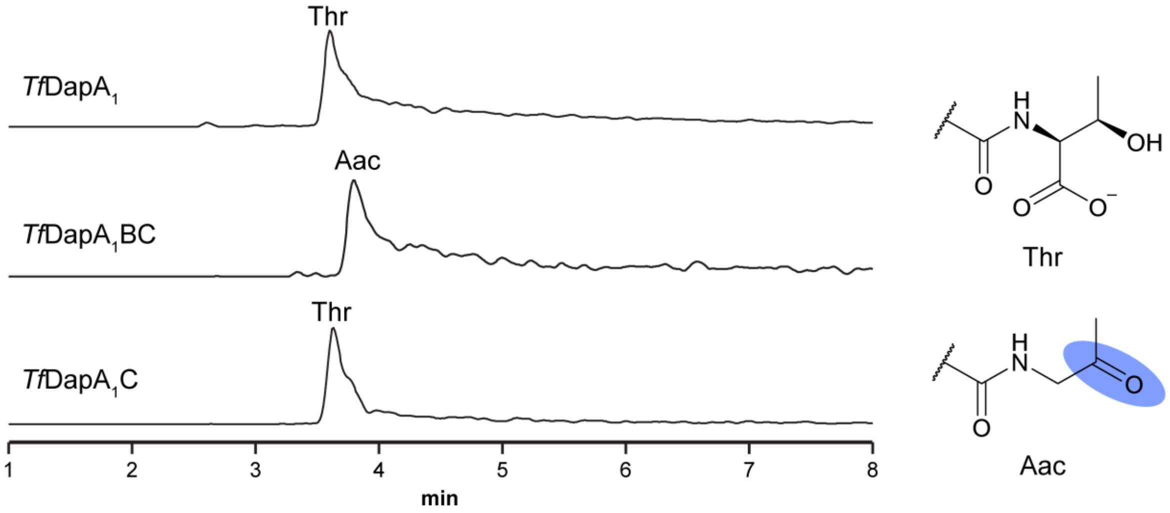

**Figure S10. LC-MS chromatograms evaluating *Tf*DapB dependency.** a, Extracted ion chromatograms for products of each reaction are shown. Extracted ions: Thr, 1681.8085 *m/z*; Aac, 1666.4734 *m/z*. Abbreviations: Aac, aminoacetone.

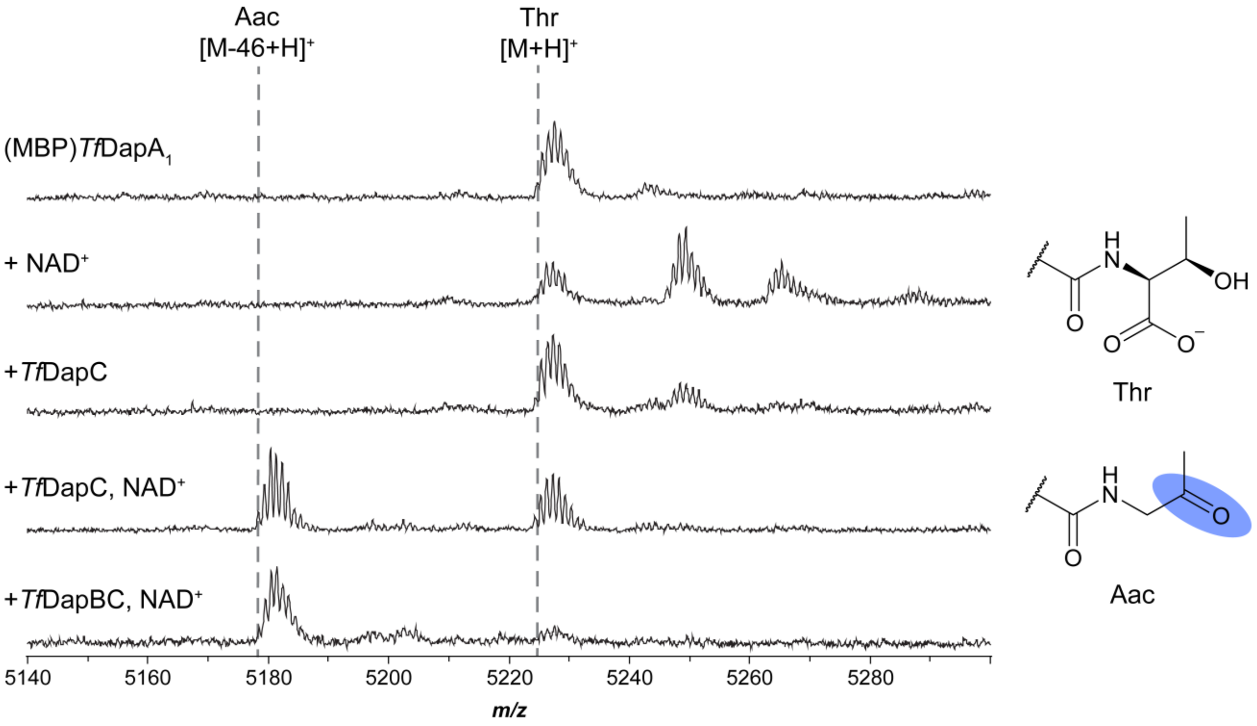

**Figure S11. MALDI-TOF mass spectra for activity of *Tf*DapBC on MBP-tagged *Tf*DapA$_1$.** Spectra were collected after TEV proteolysis and desalting by C18 ZipTip. Abbreviations: Aac, aminoacetone. Key expected masses: *Tf*DapA$_1$, 5225 Da; *Tf*DapA$_1$-Aac, 5179 Da.

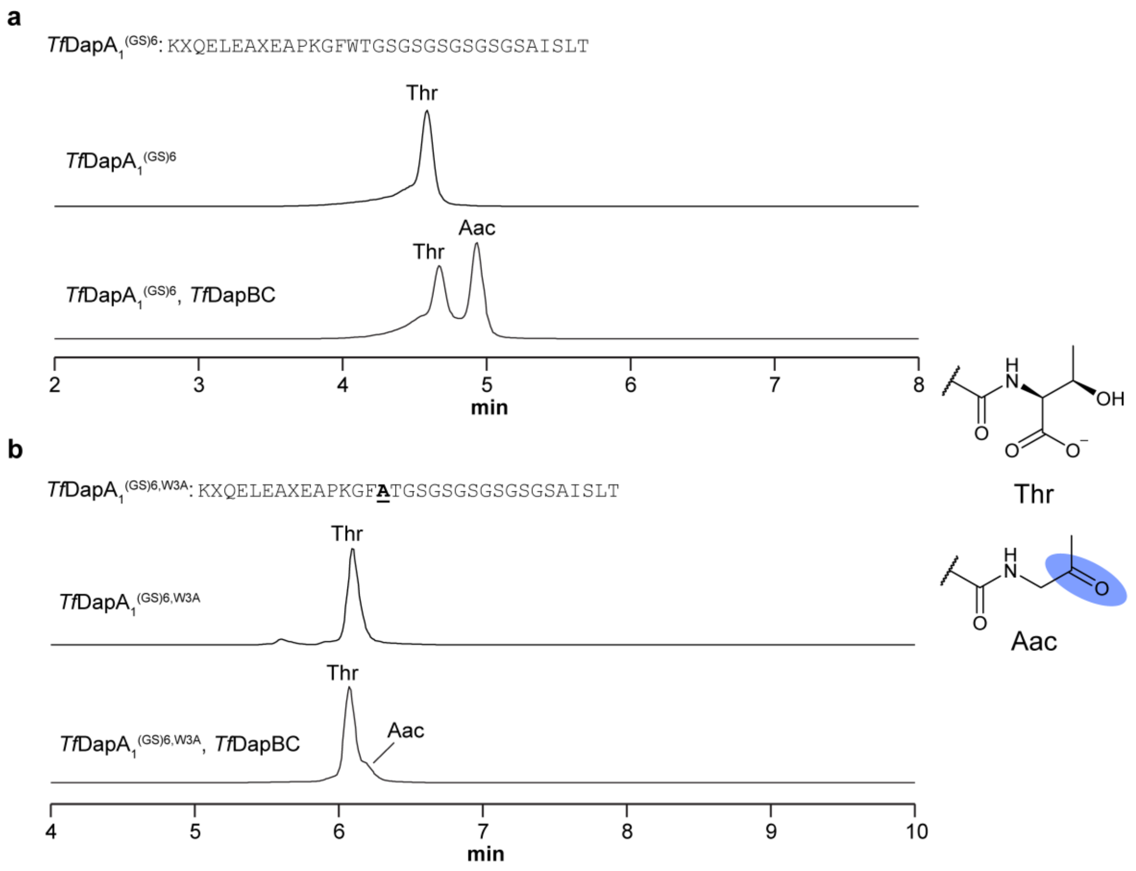

**Figure S12. Evaluation of *Tf*DapA$_1$-Trp3 importance for decarboxylase activity.** a, Extracted ion chromatograms for reactions of *Tf*DapA$_1^{(GS)6}$ are shown. b, Extracted ion chromatograms for *Tf*DapA$_1^{(GS)6,W3A}$ are shown. Extracted ions: *Tf*DapA$_1^{(GS)6}$, 1070.8713 *m/z*; *Tf*DapA$_1^{(GS)6}$-Aac, 1055.5362 *m/z*; *Tf*DapA$_1^{(GS)6,W3A}$, 1032.5239 *m/z*; *Tf*DapA$_1^{(GS)6,W3A}$-Aac, 1017.1888 *m/z*. Abbreviations: Aac, aminoacetone.

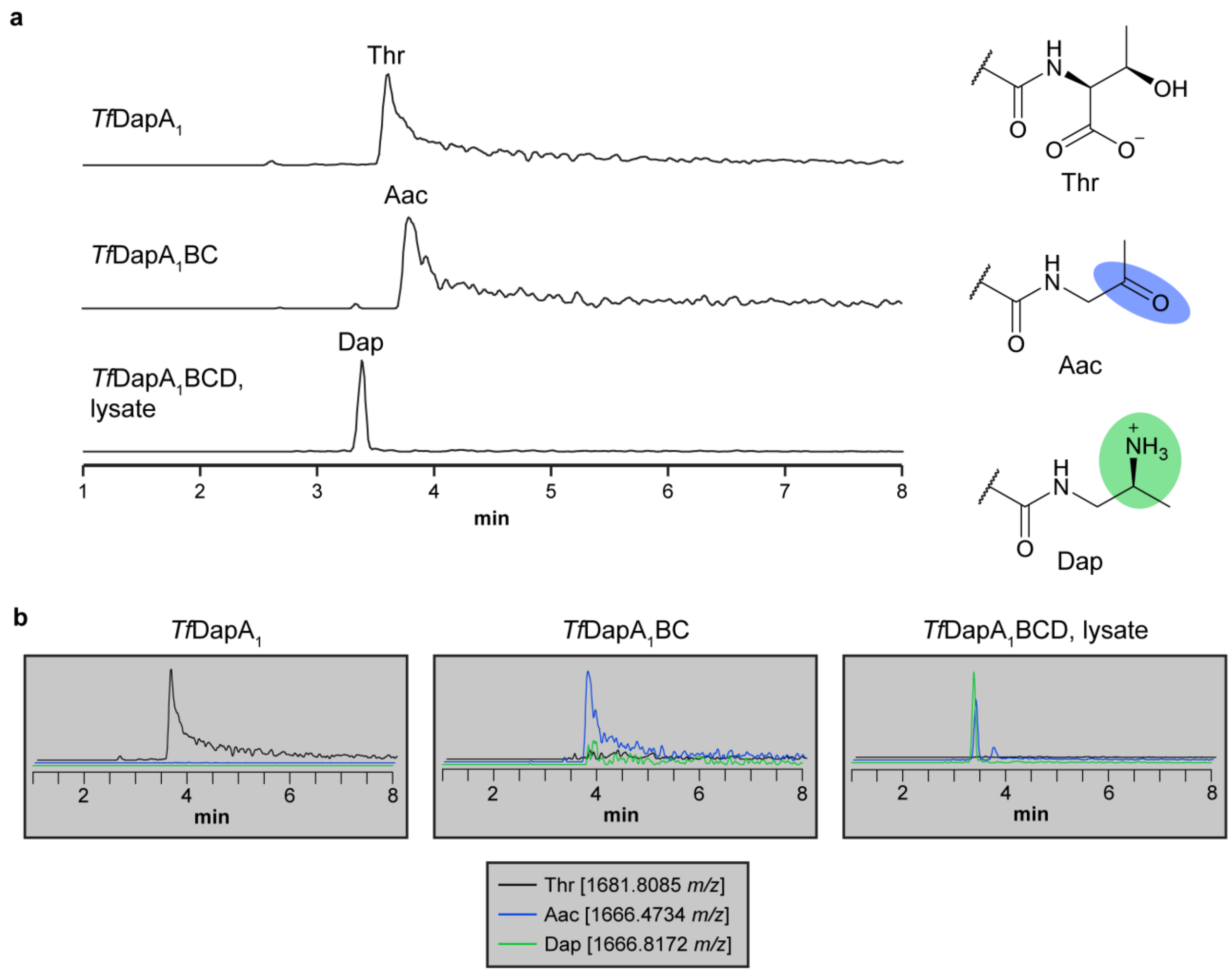

**Figure S13. LC-MS chromatograms for reconstitution of *Tf*DapD.** a, Extracted ion chromatograms are shown for the expected products of each reaction. b, Extracted ion chromatograms corresponding to Thr, Aac, and Dap are shown for each reaction. Extracted ions: Thr, 1681.8085 *m/z*; Aac, 1666.4734 *m/z*, Dap, 1666.8172 *m/z*. Abbreviations: Aac, aminoacetone; Dap, 1,2-diaminopropane.

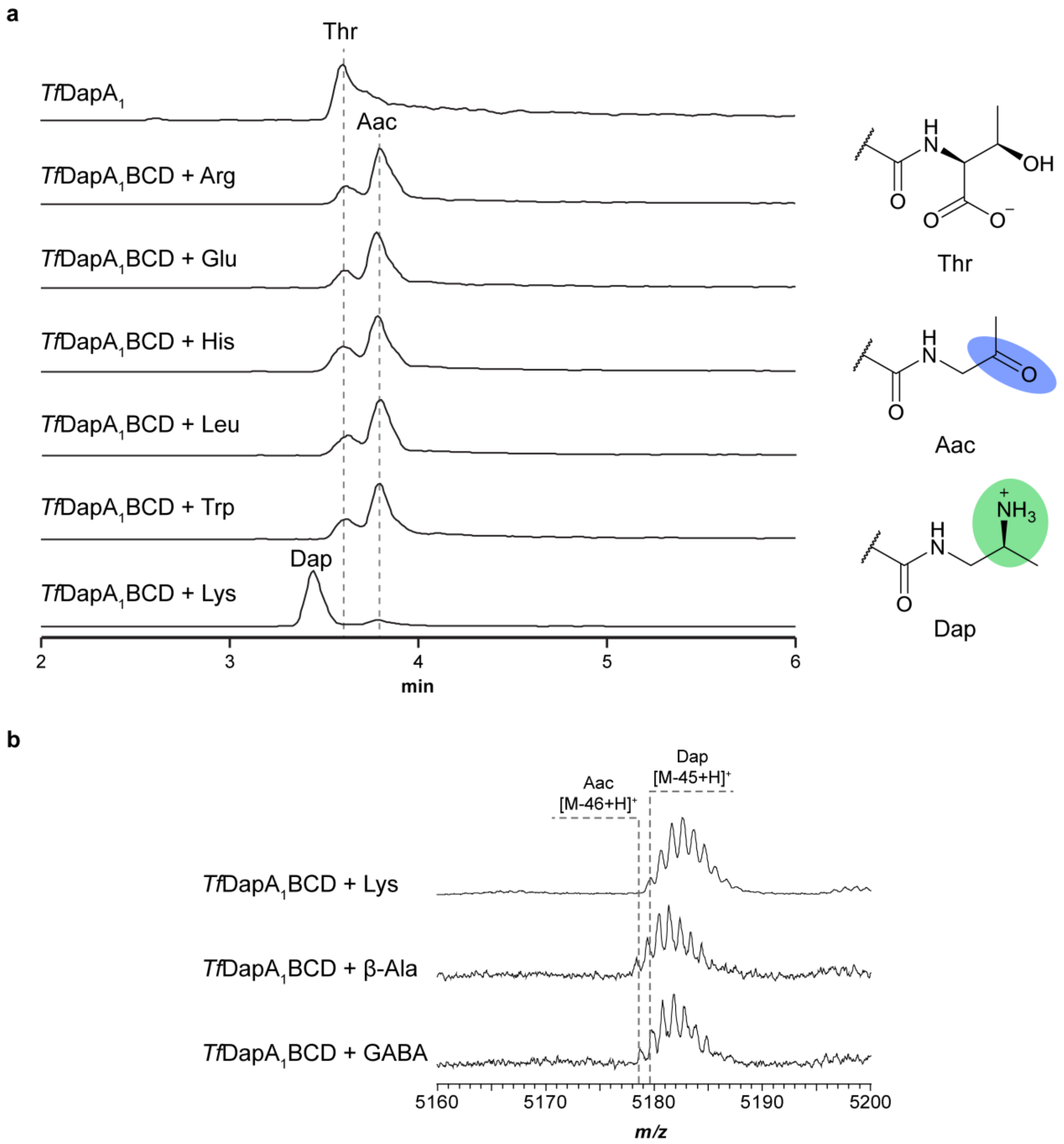

**Figure S14. Co-substrate determination for *Tf*DapD.** a, Extracted ion chromatograms for reactions of *Tf*DapA1 examining different proteinogenic amino acid co-substrates for *Tf*DapD. b, MALDI-TOF mass spectra for examination of additional non-proteinogenic ω-amine co-substrates with (MBP)*Tf*DapA1 as the substrate. MALDI-TOF mass spectra were collected after TEV proteolysis and desalting by C18 ZipTip. Abbreviations: Aac, aminoacetone; Dap, 1,2-diaminopropane.

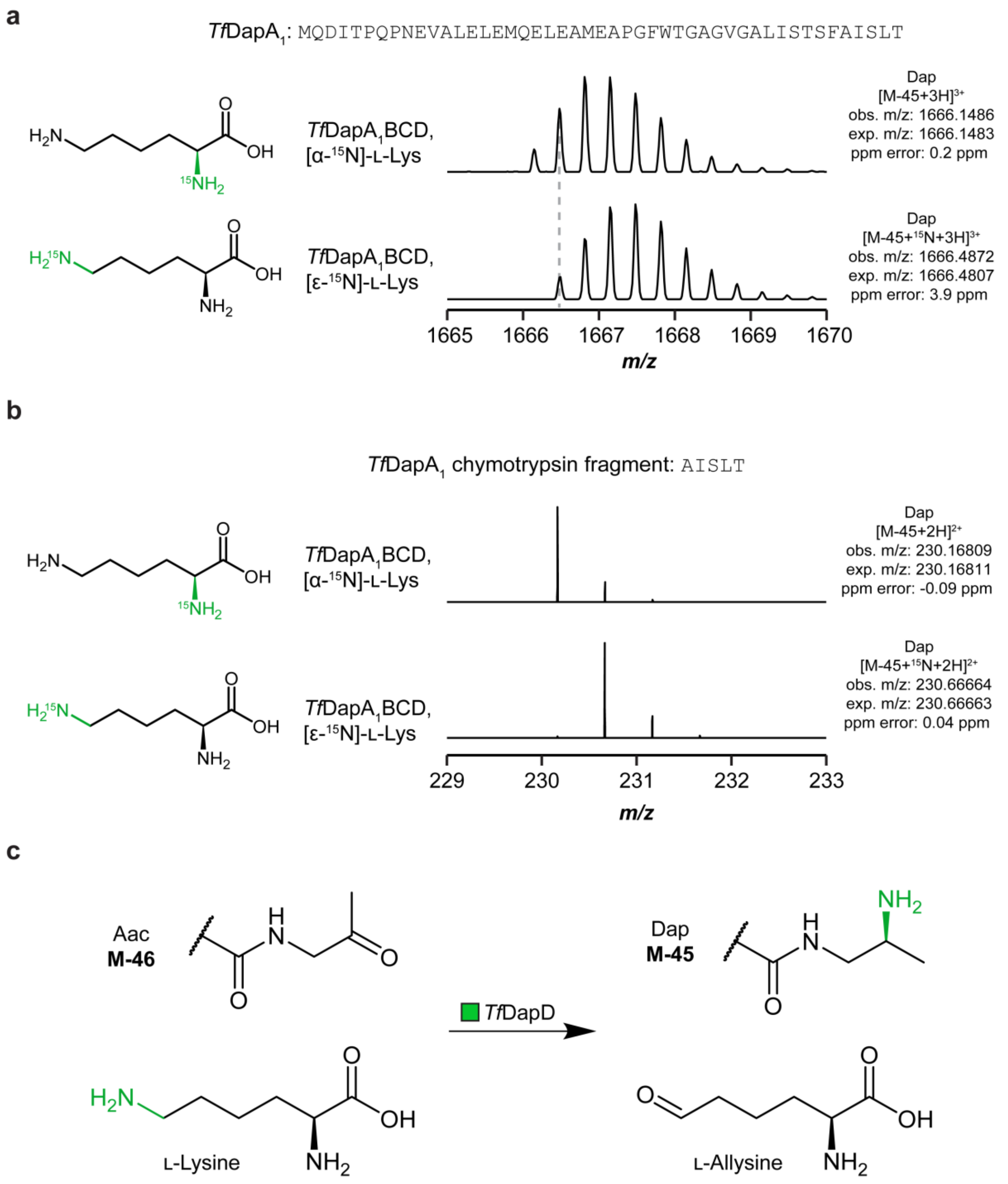

**Figure S15. Isotopic labelling experiments for *Tf*DapD amine selectivity.** a, HR mass spectra are shown for reactions using isotopically labelled L-Lys standards. b, HR mass spectra are shown for fragment obtained by chymotrypsin digestion after reactions using isotopically labelled L-Lys standards. c, Scheme for reaction catalyzed by *Tf*DapD. Abbreviations: Aac, aminoacetone; Dap, 1,2-diaminopropane.

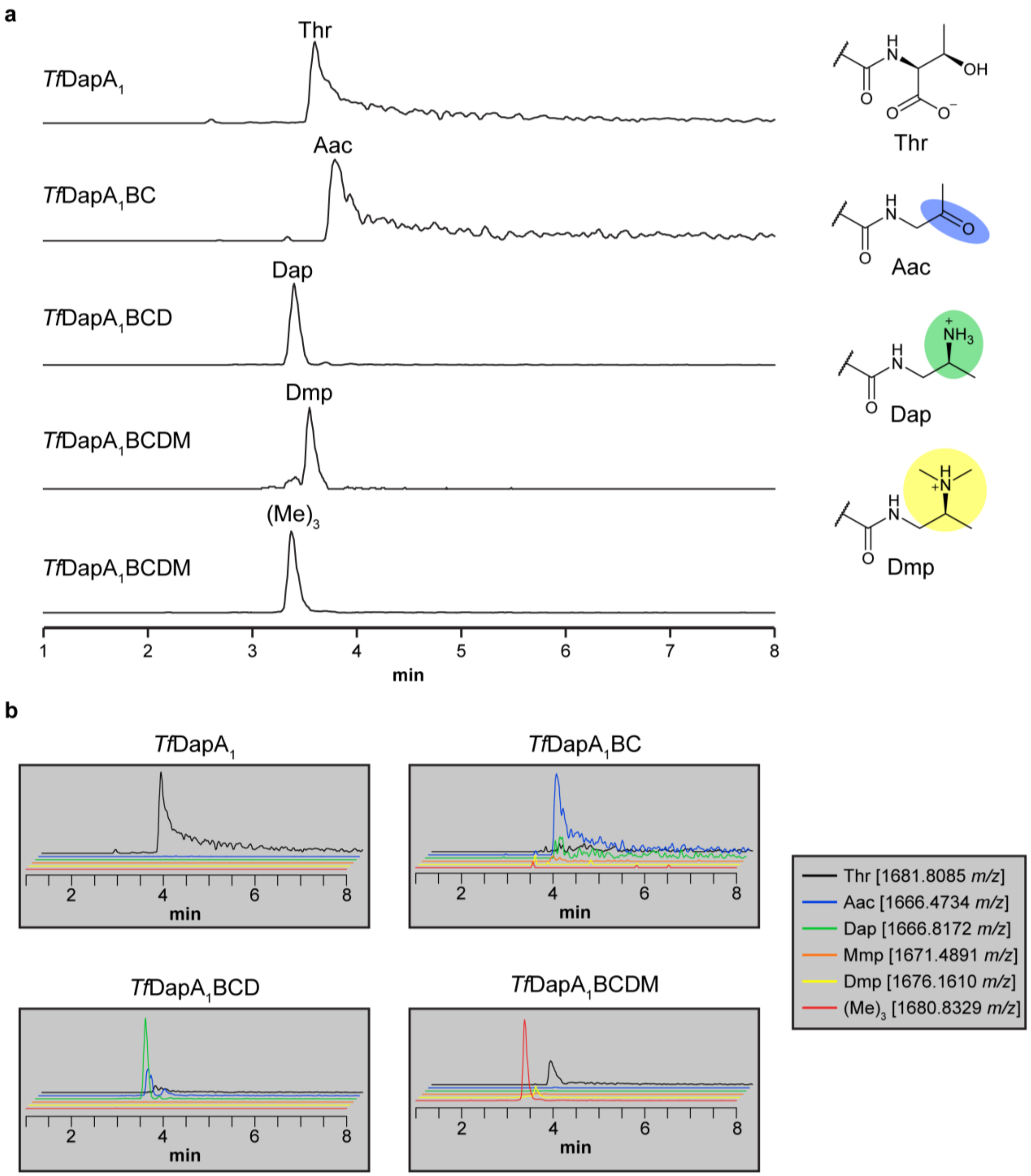

**Figure S16. LC-MS chromatograms for reconstitution of *Tf*DapM.** a, Extracted ion chromatograms are shown for the expected products of each reaction. b, Extracted ion chromatograms corresponding to Thr, Aac, Dap, Mmp, Dmp, and (Me)3 products are shown for each reaction. Extracted ions: Thr, 1681.8085 *m/z*; Aac, 1666.4734 *m/z*; Dap, 1666.8172 *m/z*; Mmp, 1671.4891 *m/z*; Dmp, 1676.1610 *m/z*; (Me)3, 1680.8329 *m/z*. Abbreviations: Aac, aminoacetone; Dap, 1,2-diaminopropane; Mmp, monomethylpropane-1,2-diamine; Dmp, dimethylpropane-1,2-diamine.

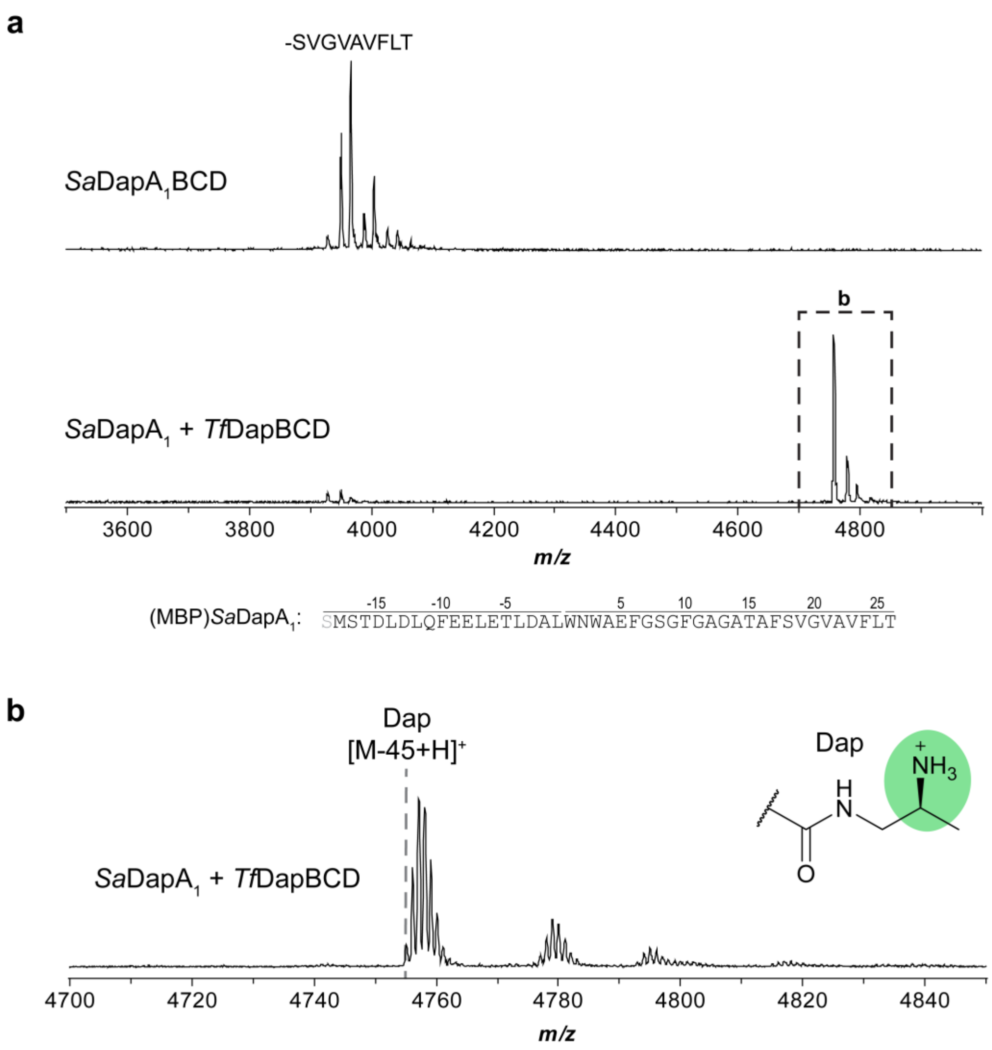

**Figure S17. MALDI-TOF mass spectra for recombinant production of *Sa*DapA$_1$-Dap.** a, Spectra for co-expressions of *Sa*DapA$_1$ with *Sa*DapBCD or *Tf*DapBCD, respectively. b, Zoom-in of *Sa*DapA$_1$-Dap produced by co-expression with *Tf*DapBCD. Spectra were collected after TEV proteolysis and desalting by C18 ZipTip. Abbreviations: Dap, 1,2-diaminopropane. Key expected masses: *Sa*DapA$_1$, 4800 Da; *Sa*DapA$_1$-Dap, 4755 Da.

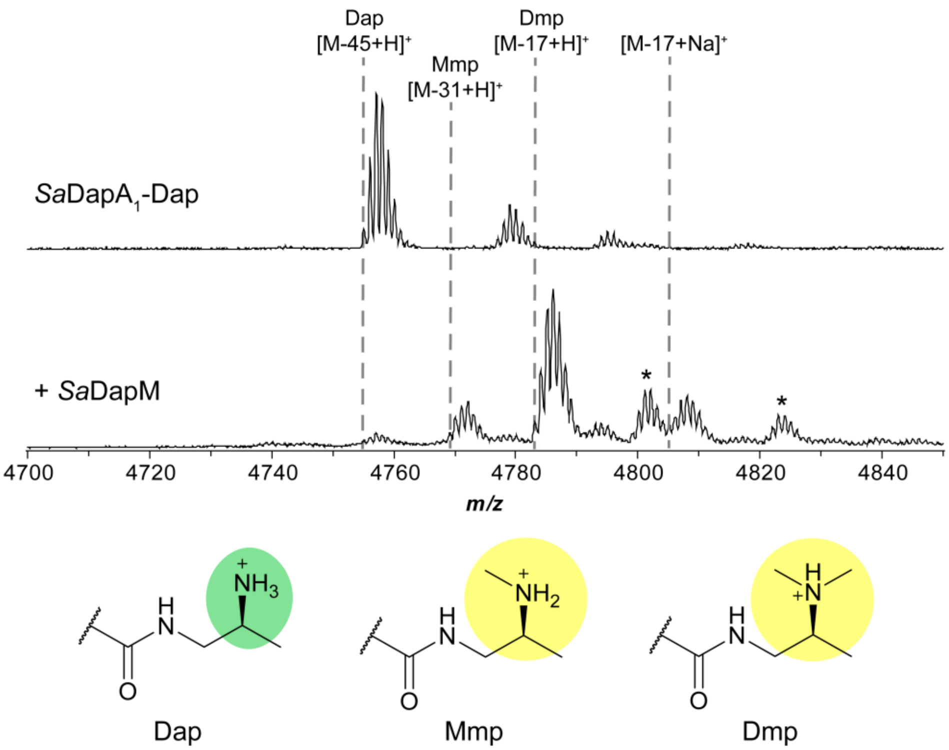

**Figure S18. MALDI-TOF mass spectra for reconstitution of *Sa*DapM.** Reaction with *Sa*DapM was performed using recombinantly expressed (MBP)*Sa*DapA$_1$-Dap as substrate. Asterisks indicate additional oxidation products and/or potassium adducts. Spectra were collected after TEV proteolysis and desalting by C18 ZipTip. Abbreviations: Dap, 1,2-diaminopropane; Mmp, monomethylpropane-1,2-diamine; Dmp, dimethylpropane-1,2-diamine. Key expected masses: *Sa*DapA$_1$-Dap, 4755 Da; *Sa*DapA$_1$-Mmp, 4769 Da; *Sa*DapA$_1$-Dmp, 4783 Da. Sodium adducts result in addition of 22 Da. Potassium adducts result in addition of 38 Da.

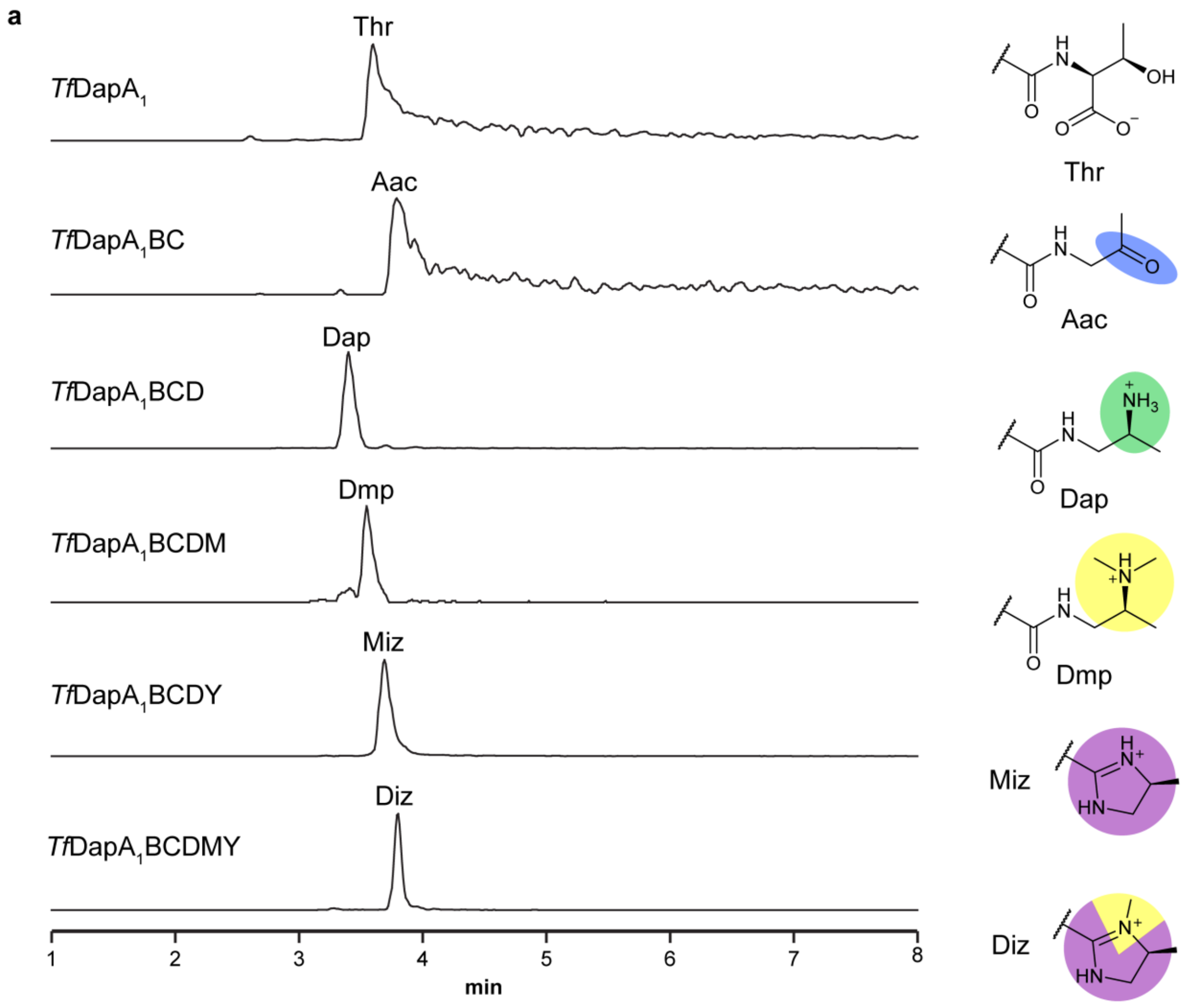

**Figure S19. LC-MS chromatograms for reconstitution of *Tf*DapY.** a, Extracted ion chromatograms are shown for the expected products of each reaction. b, Extracted ion chromatograms corresponding to Thr, Aac, Dap, Mmp, Dmp, $(Me)_3$, Miz, and Diz are shown for each reaction. Extracted ions: Thr, 1681.8085 *m/z*; Aac, 1666.4734 *m/z*; Dap, 1666.8172 *m/z*; Mmp, 1671.4891 *m/z*; Dmp, 1676.1610 *m/z*; $(Me)_3$, 1680.8329 *m/z*; Miz, 1660.8137 *m/z*; Diz, 1665.4856 *m/z*. Abbreviations: Aac, aminoacetone; Dap, 1,2-diaminopropane; Mmp, monomethylpropane-1,2-diamine; Dmp, dimethylpropane-1,2-diamine; Miz, 4-methylimidazoline; Diz, 3,4-dimethylimidazoline.

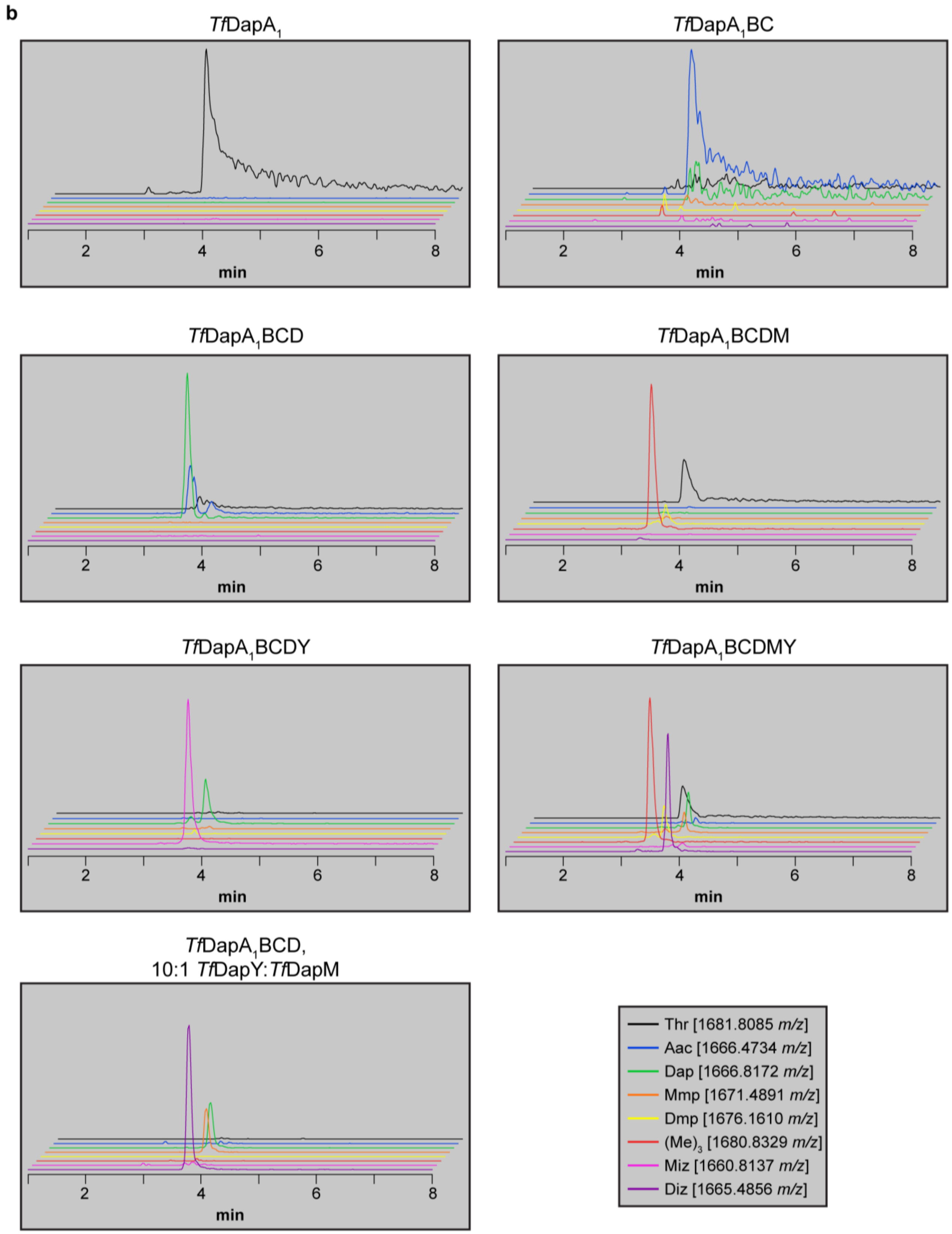

**Figure S19 (cont.). LC-MS chromatograms for initial reconstitution of TfDapY.**

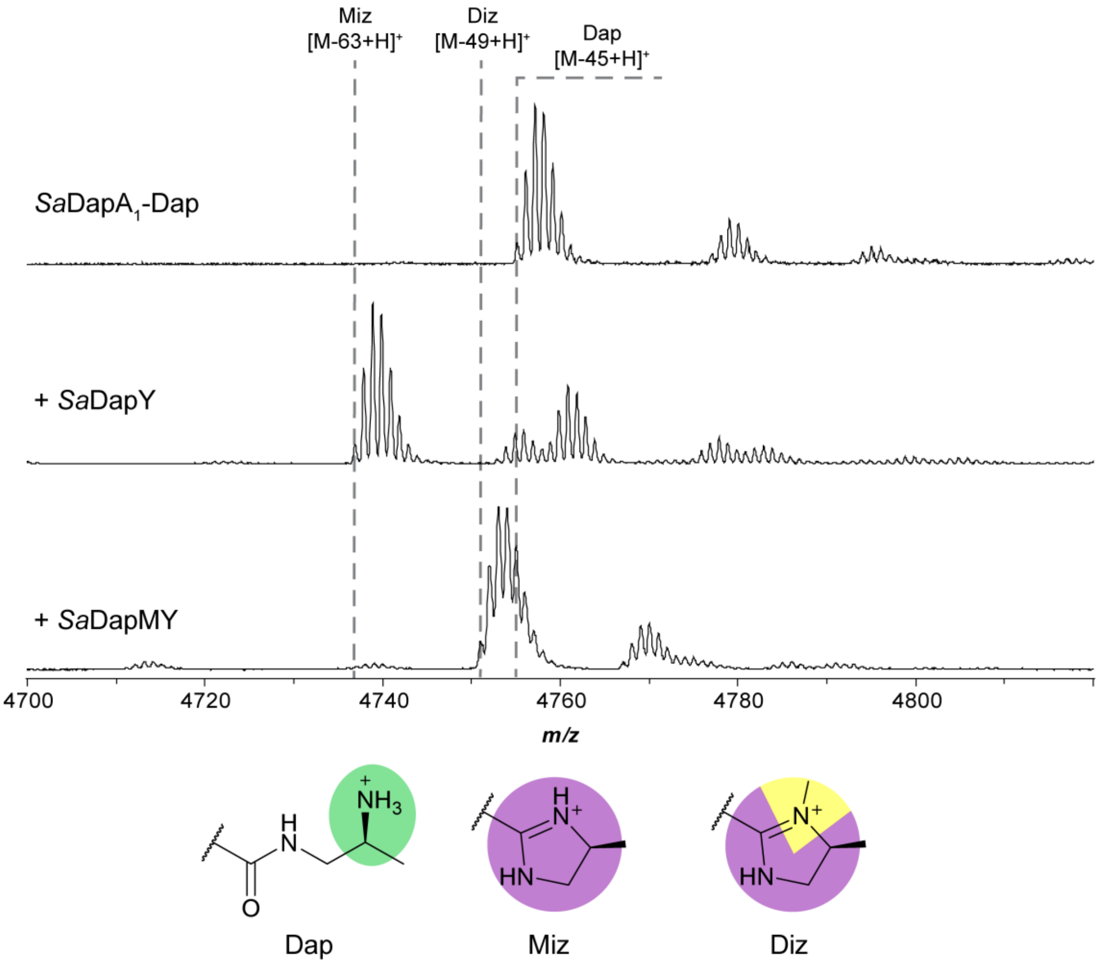

**Figure S20. MALDI-TOF mass spectra for reconstitution of *Sa*DapY activity *in vitro*.** Reactions with *Sa*DapY were performed using recombinantly expressed (MBP)*Sa*DapA1-Dap as substrate. Spectra were collected after TEV proteolysis and desalting by C18 ZipTip. Abbreviations: Dap, 1,2-diaminopropane; Miz, 4-methylimidazoline; Diz, 3,4-dimethylimidazoline. Key expected masses: *Sa*DapA1-Dap, 4755 Da, *Sa*DapA1-Miz, 4737 Da, *Sa*DapA1-Diz, 4751 Da.

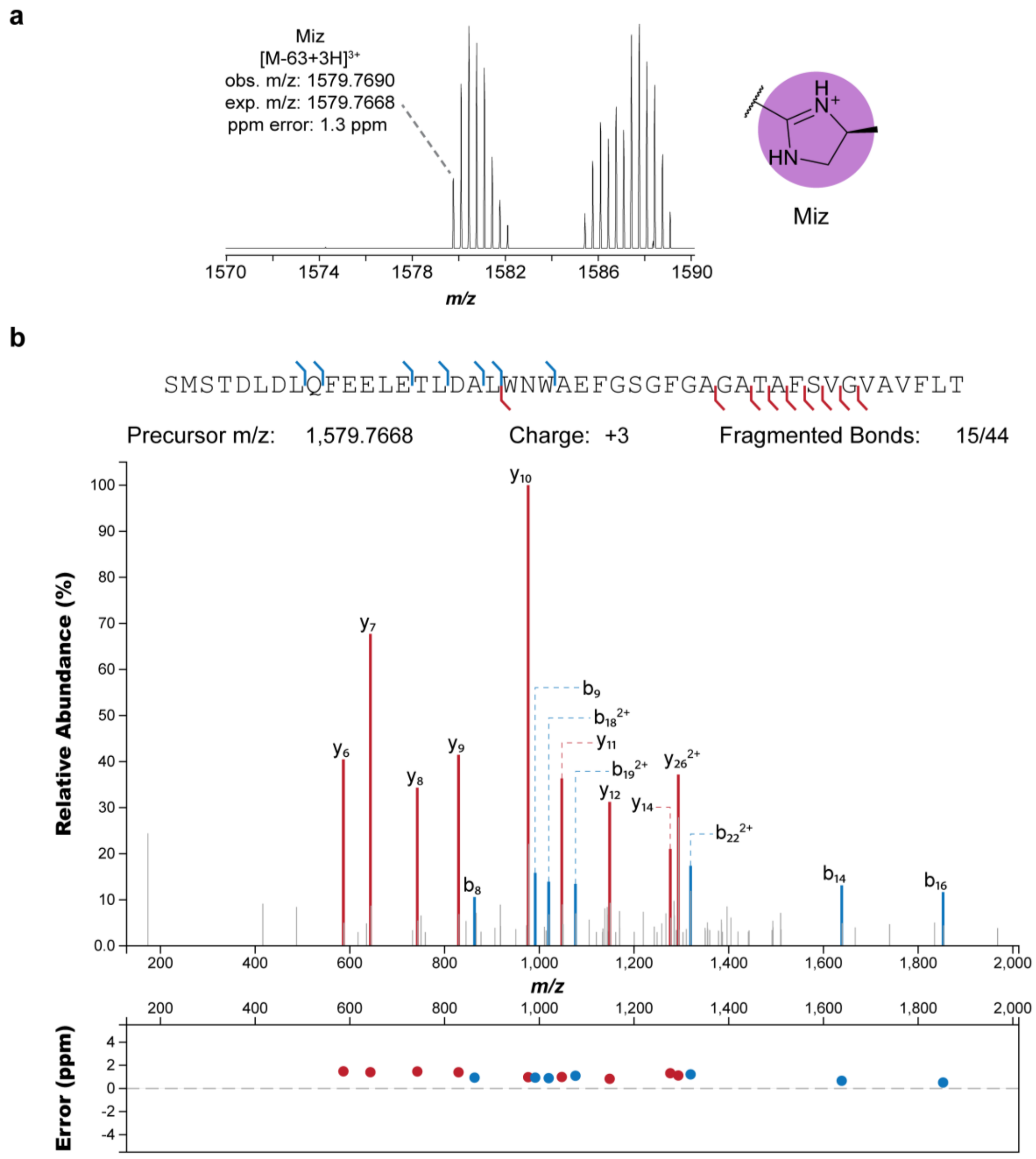

**Figure S21. Localization of modification site for *Sa*DapA$_1$ products modified by *Sa*DapY.** a, HR-MS and b, HR-MS/MS spectra for Miz product detected after reaction of (MBP)*Sa*DapA$_1$-Dap with *Sa*DapY. c, HR-MS, d, HR-MS/MS, and e, MALDI-LIFT-TOF/TOF mass spectra for Diz product detected after reaction of (MBP)*Sa*DapA$_1$-Dap with *Sa*DapMY. HR-MS/MS annotation was performed using the Interactive Peptide Spectral Annotator Tool (IPSA).[18] Annotated fragments are in the +1 charge state unless otherwise indicated. A summary of identified fragments is presented above spectra where appropriate. Calculations using IPSA were made with mass changes applied to the C-terminus.

c

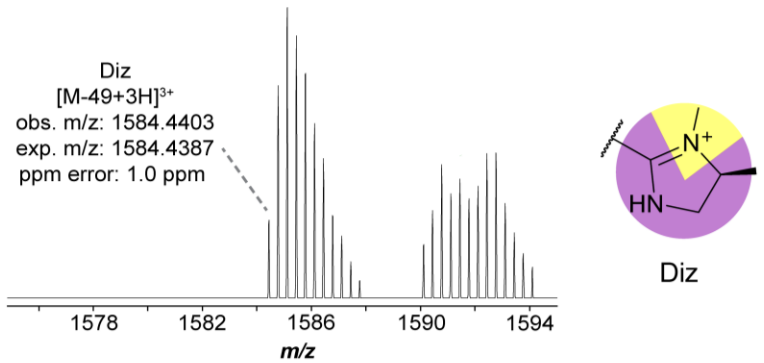

d

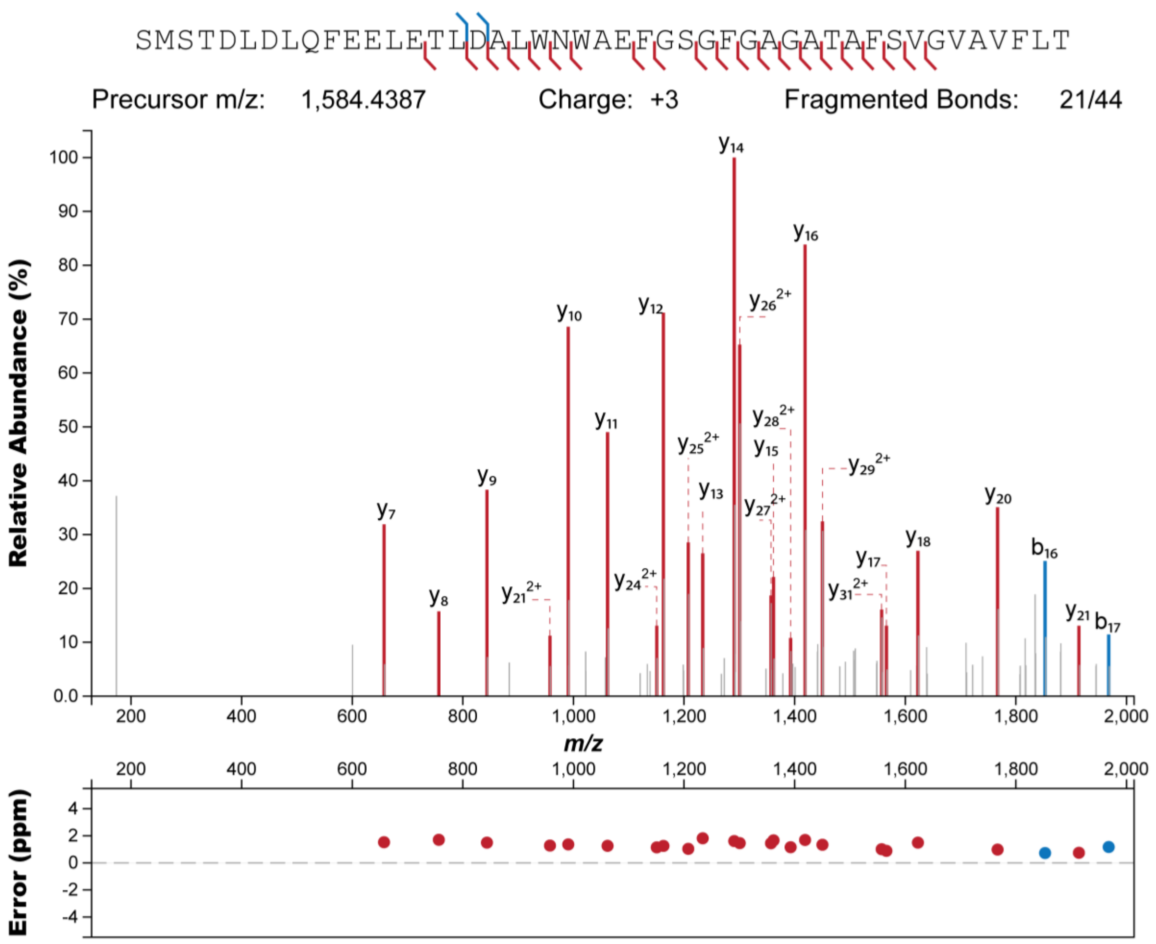

**Figure S21 (cont.). Localization of modification site for *Sa*DapA$_1$ products modified by *Sa*DapY.**

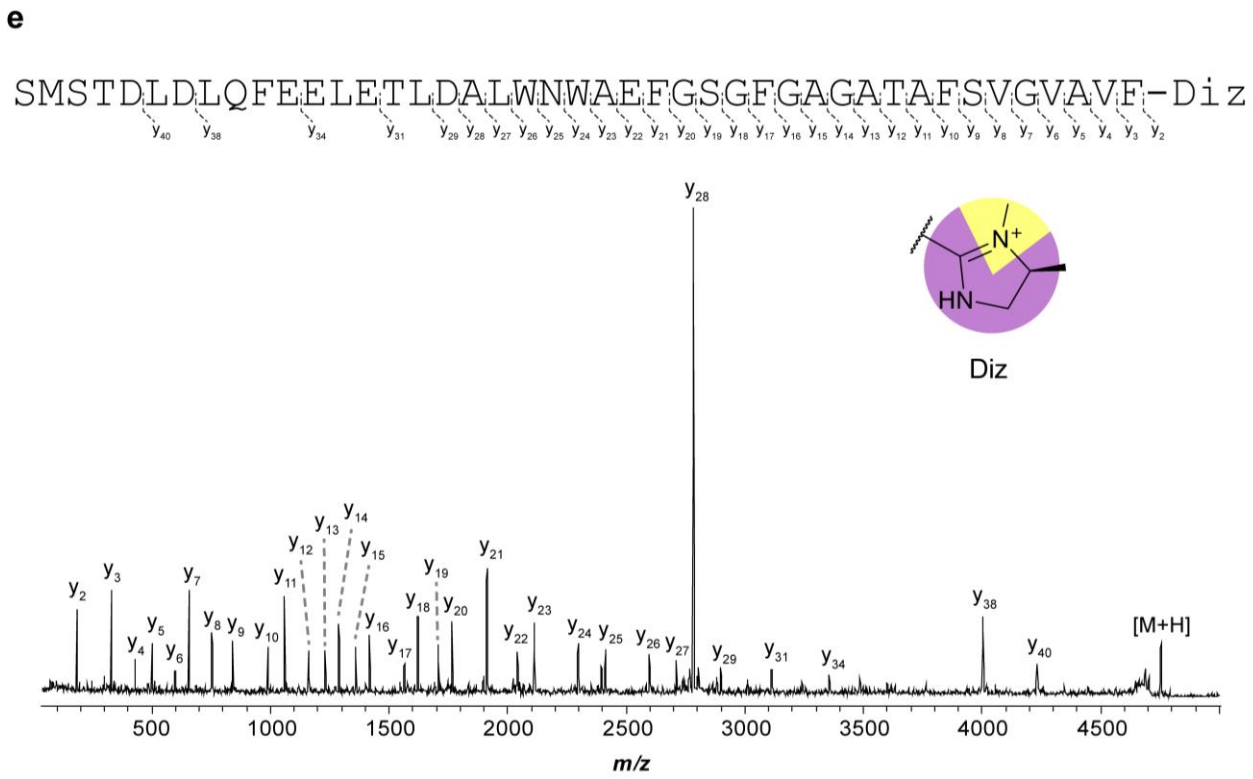

**Figure S21 (cont.). Localization of modification site for *Sa*DapA$_1$ products modified by *Sa*DapY.**

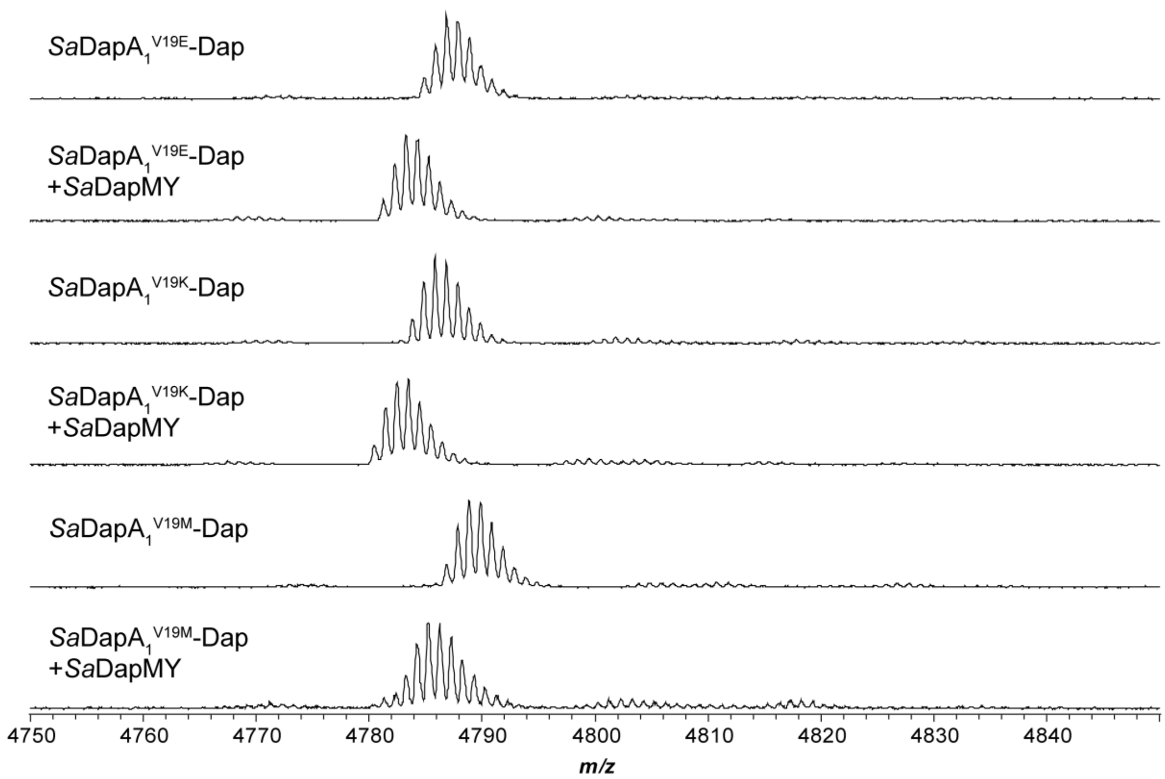

**Figure S22. Modification of single-site variants of *Sa*DapA$_1$.** (MBP)*Sa*DapA$_1$-Dap variants were generated recombinantly in *E. coli*. Further modification by *Sa*DapMY was performed *in vitro*. Spectra were collected following TEV proteolysis and desalting by C18 ZipTip. Key expected masses: *Sa*DapA$_1^{V19E}$-Dap, 4785 Da; *Sa*DapA$_1^{V19E}$-Diz, 4781 Da; *Sa*DapA$_1^{V19K}$-Dap, 4784 Da; *Sa*DapA$_1^{V19K}$-Diz, 4780 Da; *Sa*DapA$_1^{V19M}$-Dap, 4787 Da; *Sa*DapA$_1^{V19M}$-Diz, 4783 Da. Abbreviations: Dap, 1,2-diaminopropane; Diz, 3,4-dimethylimidazoline.

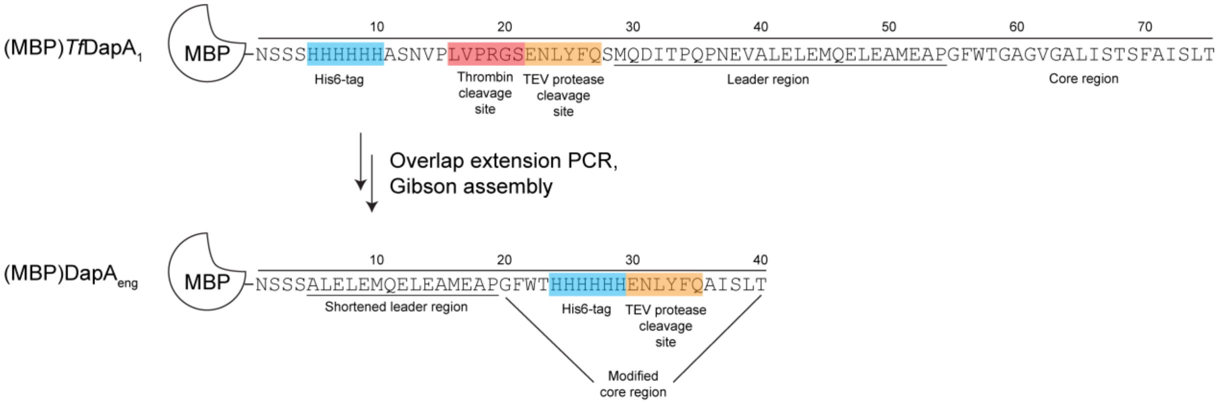

**Figure S23. Comparison of (MBP)*Tf*DapA$_1$ and (MBP)DapA$_{eng}$ construct designs.** Key features of both constructs are annotated on each design.

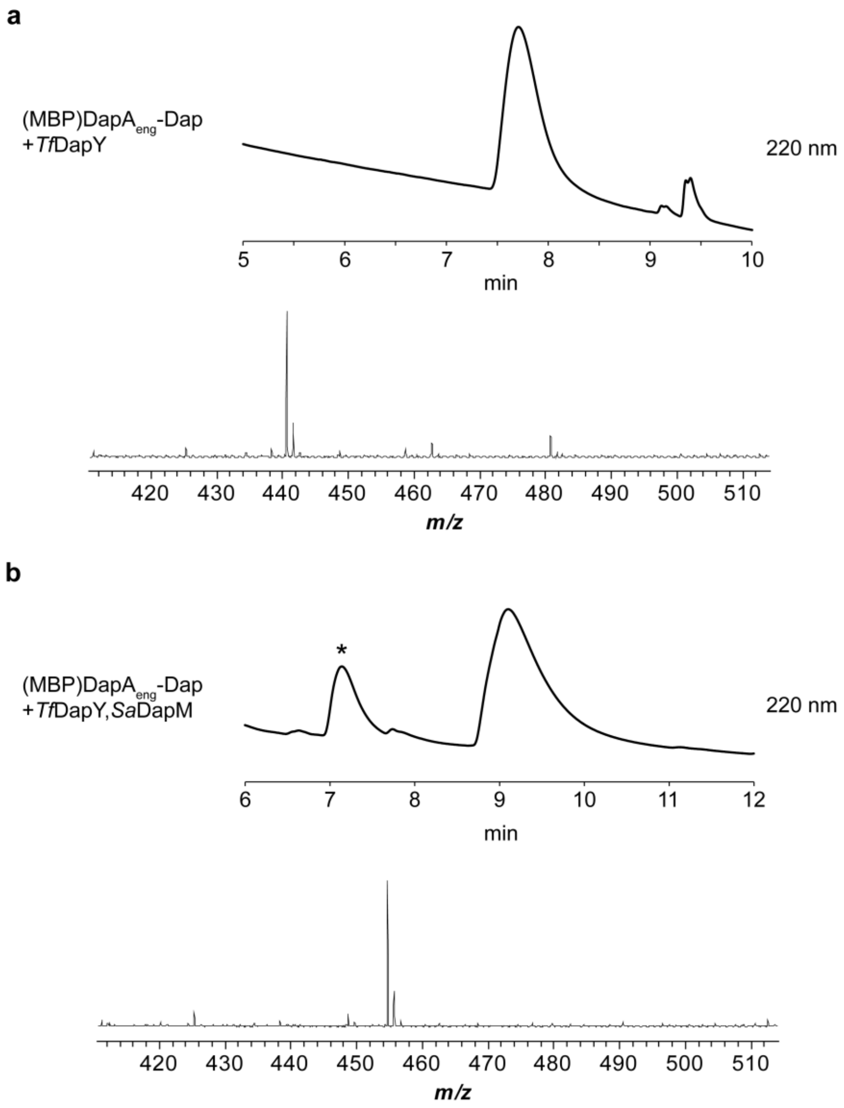

**Figure S24. Purification of modified AISLT peptides by HPLC.** a, Chromatogram for purification of *Tf*DapY-modified peptide released from (MBP)DapA$_{eng}$ with confirmatory MALDI-TOF mass spectrum ($[M\text{-}63+H]^+$ expected mass: 441 Da). b, Chromatogram for purification of *Tf*DapY- and *Sa*DapM-modified peptide released from (MBP)DapA$_{eng}$ with confirmatory MALDI-TOF mass spectrum ($[M\text{-}49+H]^+$ expected mass: 455 Da). The asterisk indicates peptide product which was only modified by *Tf*DapY. Chromatograms were generated by monitoring UV absorbance at 220 nm.

**a**

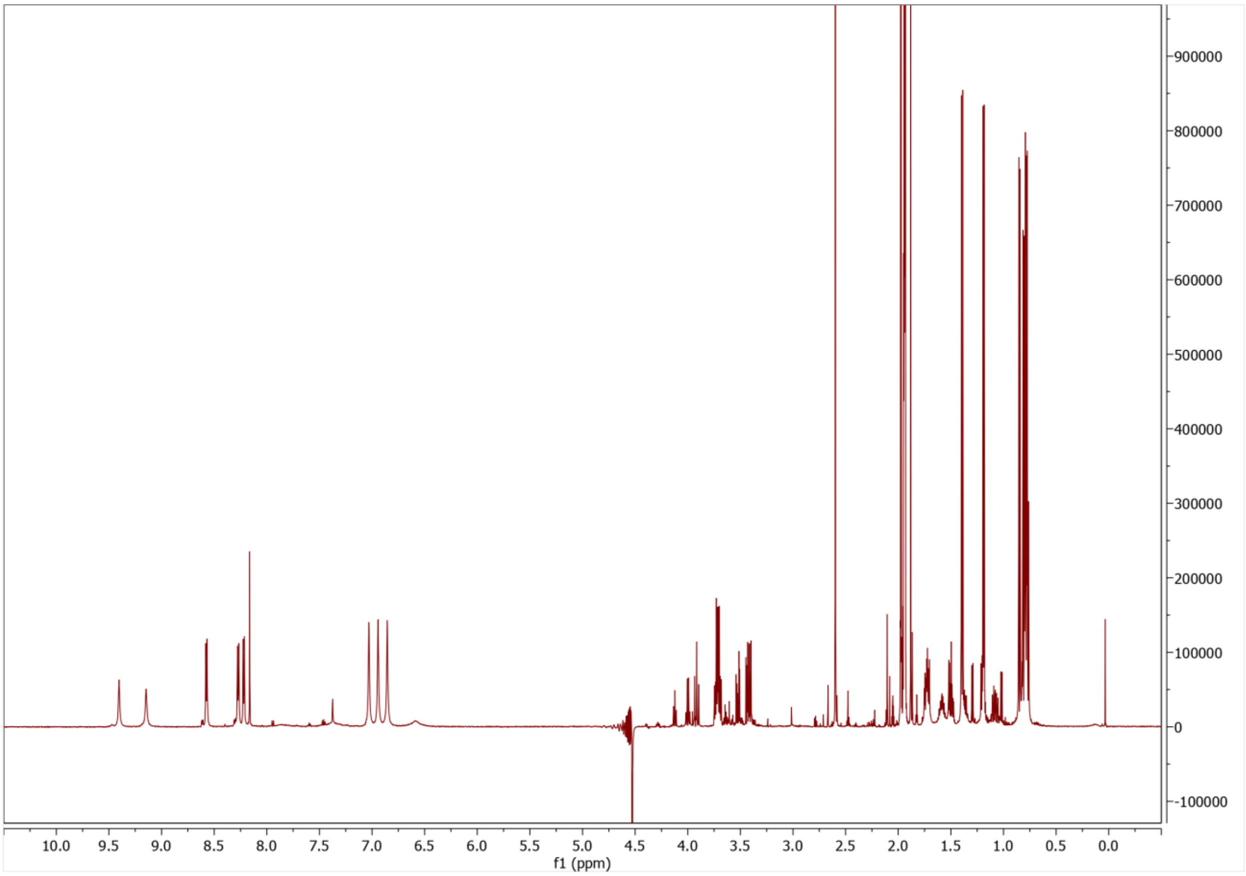

**Figure S25. Collected NMR spectra for Miz-modified peptide.** All spectra were collected in a 79.9:20:0.1 $H_2O$:$CD_3CN$:$CDO_2D$ solvent system using a 600 MHz spectrometer. a, $^1$H NMR of Miz-modified peptide. b, $^1$H-$^1$H COSY of Miz-modified peptide. c, $^1$H-$^1$H TOCSY of Miz-modified peptide. d, $^1$H-$^{13}$C HSQC of Miz-modified peptide. e, $^1$H-$^1$H NOESY of Miz-modified peptide.

**b**

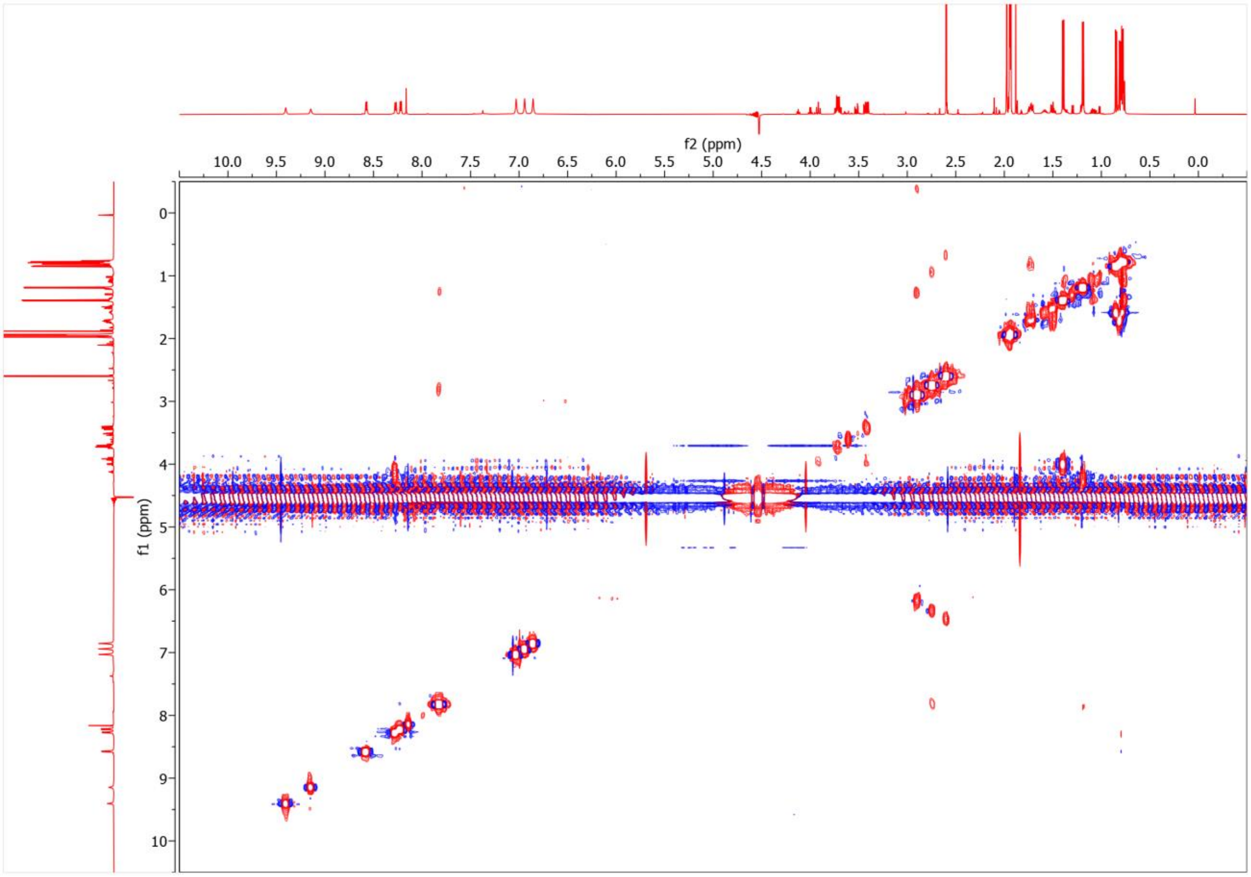

**Figure S25 (cont.). Collected NMR spectra for Miz-modified peptide.** All spectra were collected in a 79.9:20:0.1 $H_2O$:$CD_3CN$:$CDO_2D$ solvent system using a 600 MHz spectrometer. a, $^{1}H$ NMR of Miz-modified peptide. b, $^{1}H$-$^{1}H$ COSY of Miz-modified peptide. c, $^{1}H$-$^{1}H$ TOCSY of Miz-modified peptide. d, $^{1}H$-$^{13}C$ HSQC of Miz-modified peptide. e, $^{1}H$-$^{1}H$ NOESY of Miz-modified peptide.

c

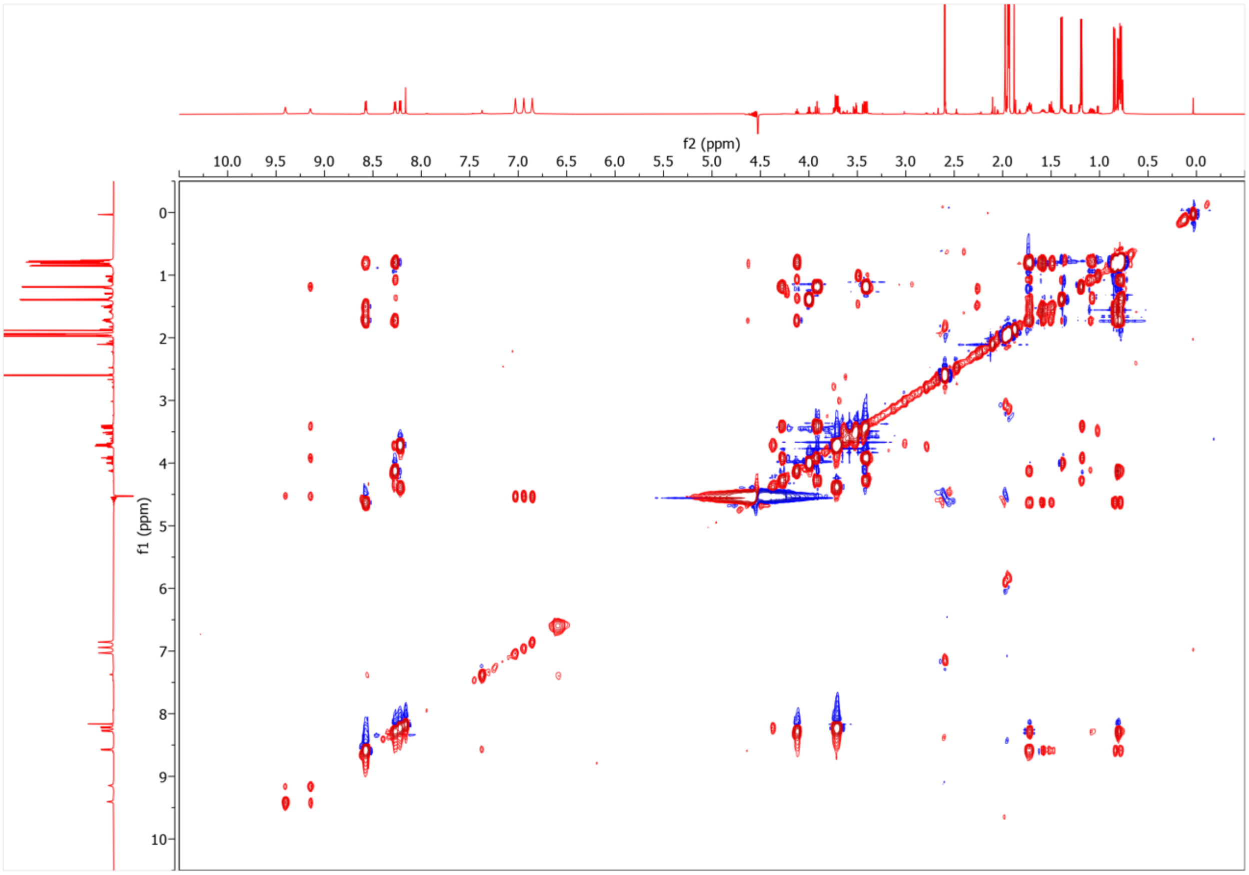

**Figure S25 (cont.). Collected NMR spectra for Miz-modified peptide.** All spectra were collected in a 79.9:20:0.1 $H_2O$:$CD_3CN$:$CDO_2D$ solvent system using a 600 MHz spectrometer. a, $^1H$ NMR of Miz-modified peptide. b, $^1H$-$^1H$ COSY of Miz-modified peptide. c, $^1H$-$^1H$ TOCSY of Miz-modified peptide. d, $^1H$-$^{13}C$ HSQC of Miz-modified peptide. e, $^1H$-$^1H$ NOESY of Miz-modified peptide.

**d**

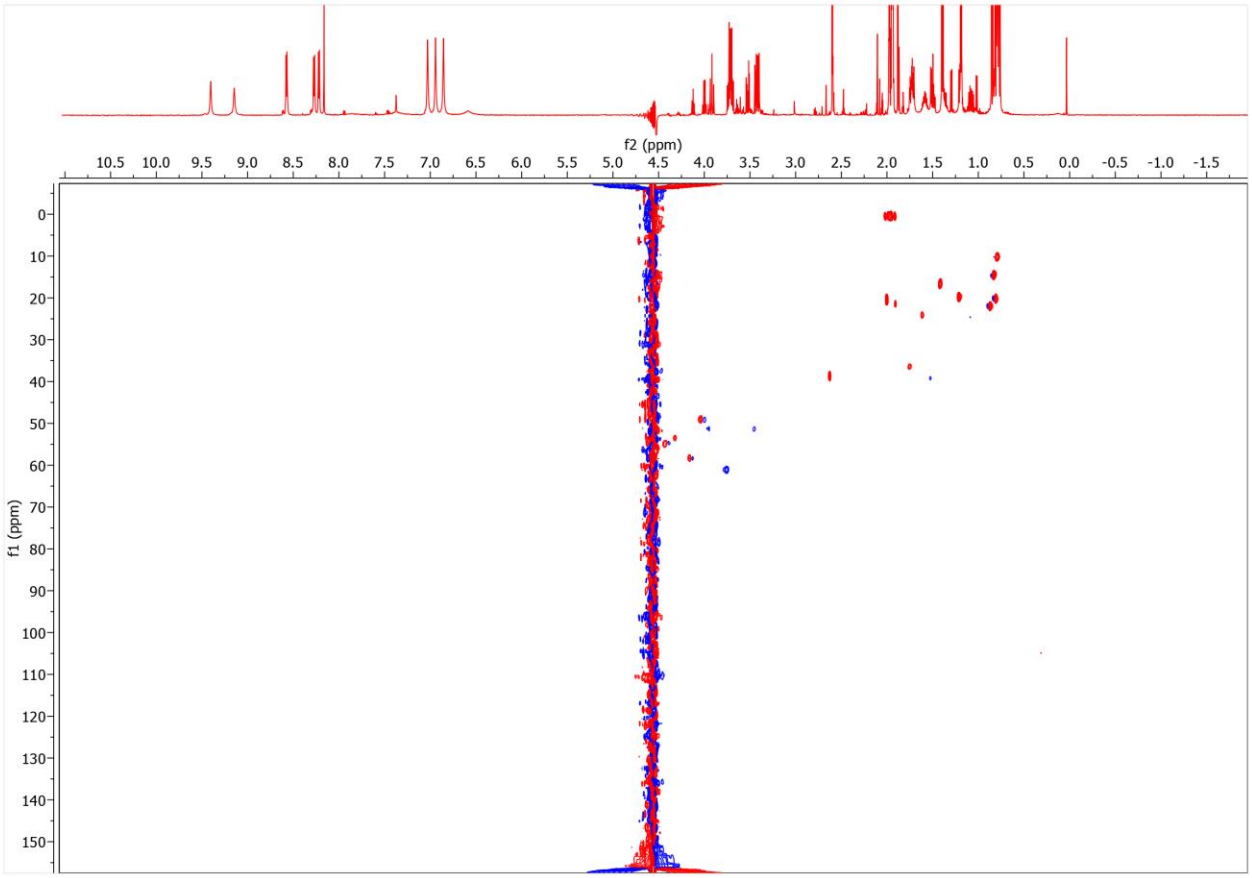

**Figure S25 (cont.). Collected NMR spectra for Miz-modified peptide.** All spectra were collected in a 79.9:20:0.1 $H_2O$:$CD_3CN$:$CDO_2D$ solvent system using a 600 MHz spectrometer. a, $^{1}H$ NMR of Miz-modified peptide. b, $^{1}H$-$^{1}H$ COSY of Miz-modified peptide. c, $^{1}H$-$^{1}H$ TOCSY of Miz-modified peptide. d, $^{1}H$-$^{13}C$ HSQC of Miz-modified peptide. e, $^{1}H$-$^{1}H$ NOESY of Miz-modified peptide.

**e**

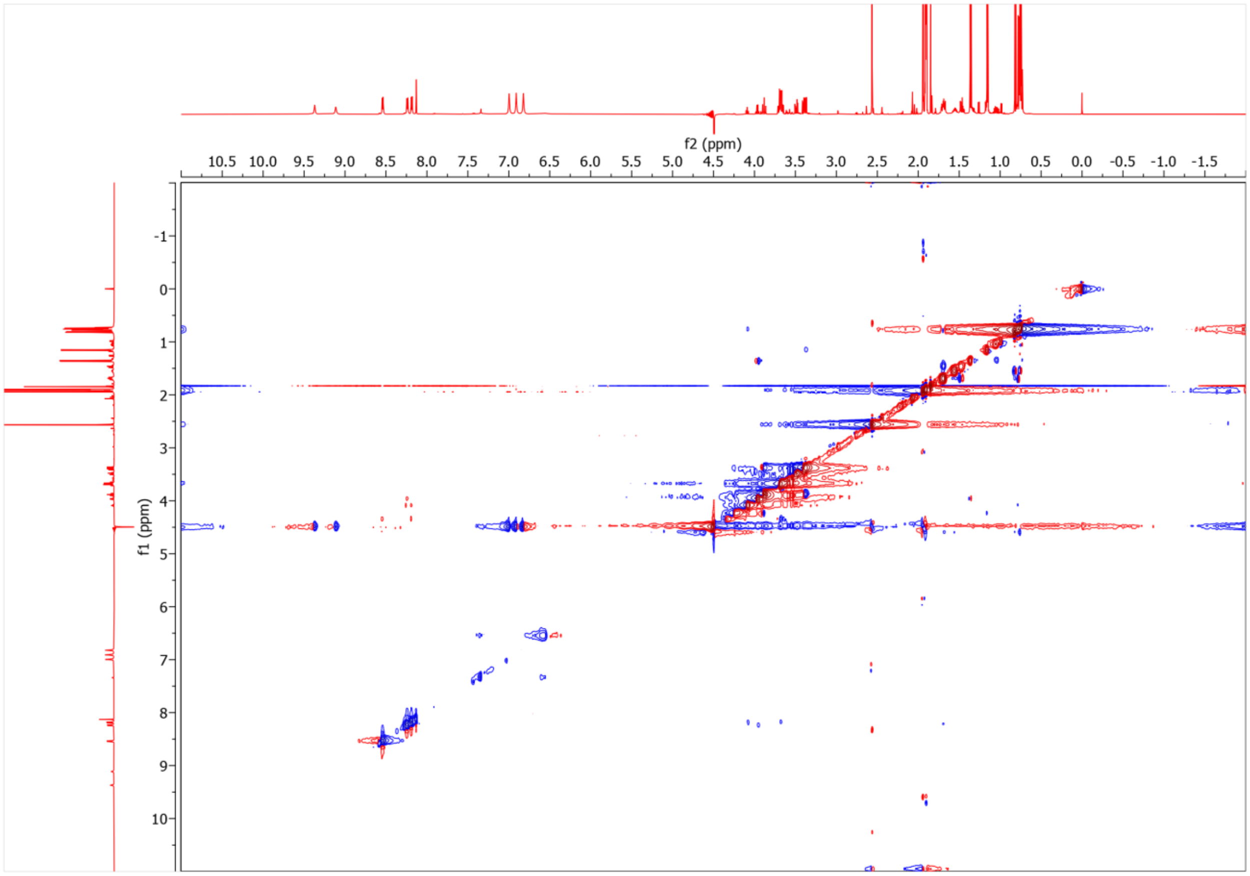

**Figure S25 (cont.). Collected NMR spectra for Miz-modified peptide.** All spectra were collected in a 79.9:20:0.1 $H_2O$:$CD_3CN$:$CDO_2D$ solvent system using a 600 MHz spectrometer. a, $^{1}H$ NMR of Miz-modified peptide. b, $^{1}H$-$^{1}H$ COSY of Miz-modified peptide. c, $^{1}H$-$^{1}H$ TOCSY of Miz-modified peptide. d, $^{1}H$-$^{13}C$ HSQC of Miz-modified peptide. e, $^{1}H$-$^{1}H$ NOESY of Miz-modified peptide.

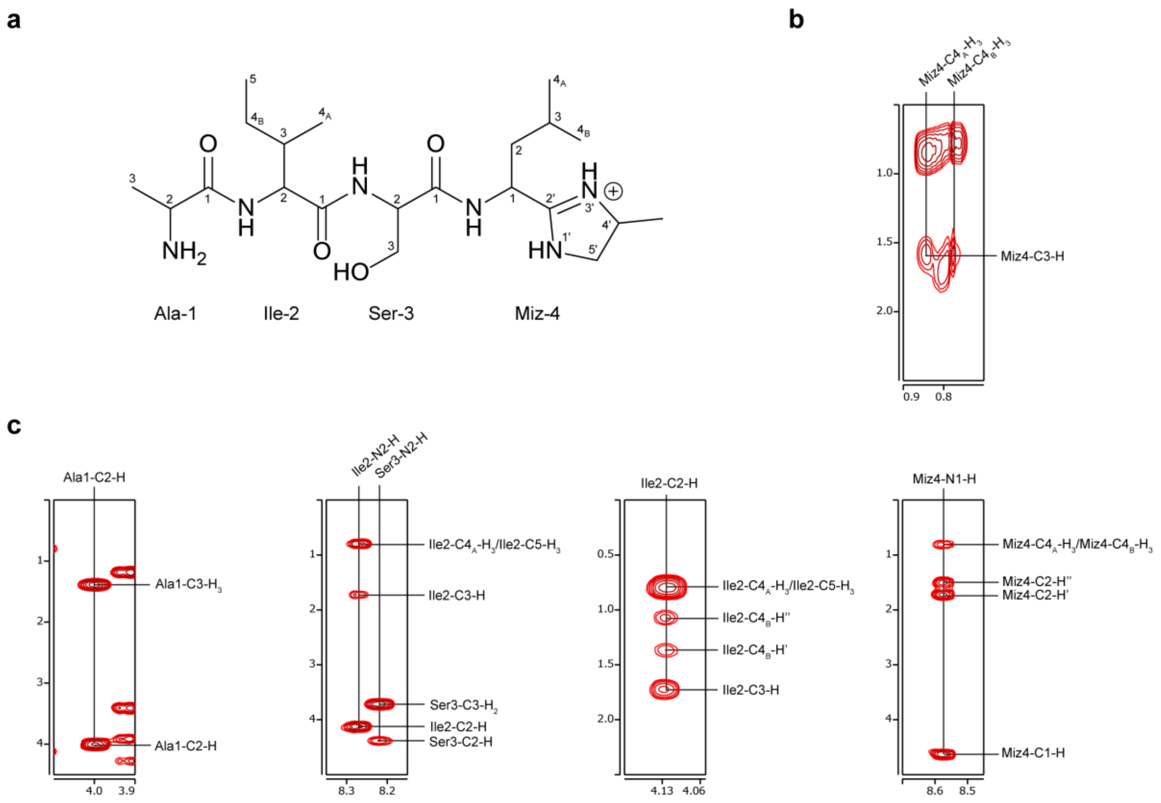

**Figure S26. Key correlations for assignment of Ala, Ile, Ser, and aminoisopentane in Miz-modified peptide.** a, Formal atom numbering for Miz-modified product. b, Key correlations observed in $^{1}H$-$^{1}H$ COSY spectrum. c, Key correlations observed in $^{1}H$-$^{1}H$ TOCSY spectrum.

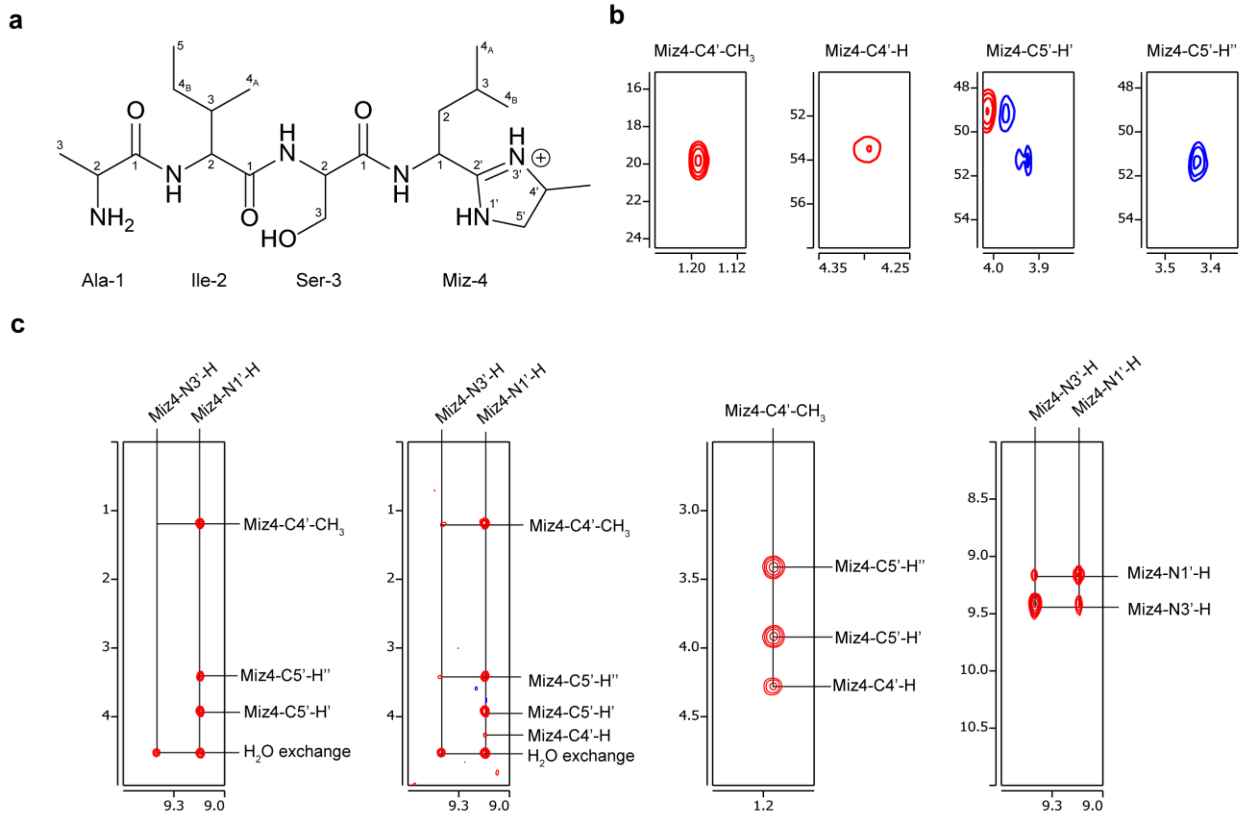

**Figure S27. Key correlations for assignment of methylimidazoline in Miz-modified peptide.** a, Formal atom numbering for Miz-modified product. b, Key correlations observed in the $^{1}H$-$^{13}C$ HSQC spectrum. c, Key correlations observed in the $^{1}H$-$^{1}H$ TOCSY spectrum.

a

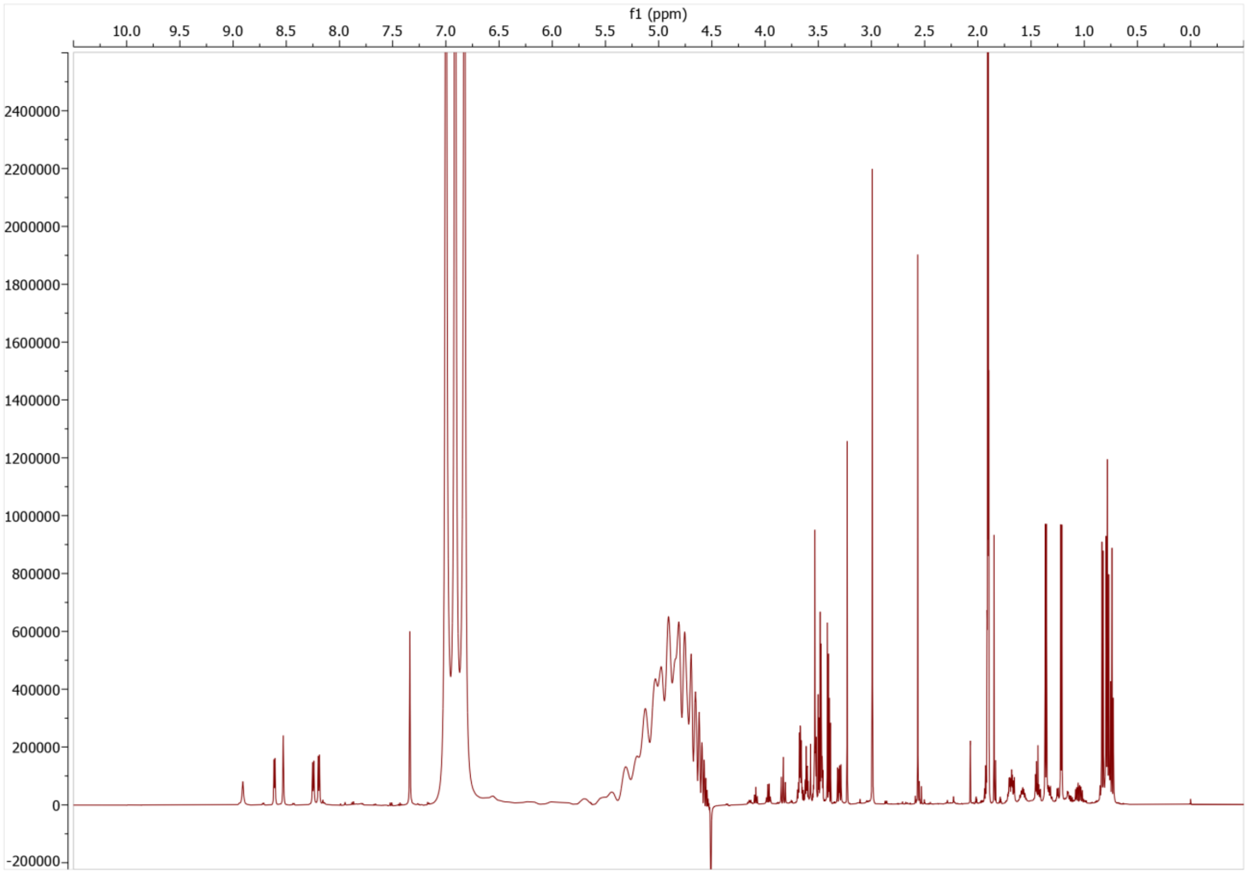

**Figure S28. Collected NMR spectra for Diz-modified peptide.** All spectra were collected in a 79.9:20:0.1 $H_2O$:$CD_3CN$:$CDO_2D$ solvent system using a 600 MHz spectrometer. a, $^{1}H$ NMR of Diz-modified peptide. b, $^{1}H$-$^{1}H$ DQF-COSY of Diz-modified peptide. c, $^{1}H$-$^{1}H$ TOCSY of Diz-modified peptide. d, $^{1}H$-$^{13}C$ HSQC of Diz-modified peptide. e, $^{1}H$-$^{1}H$ NOESY of Diz-modified peptide.

**b**

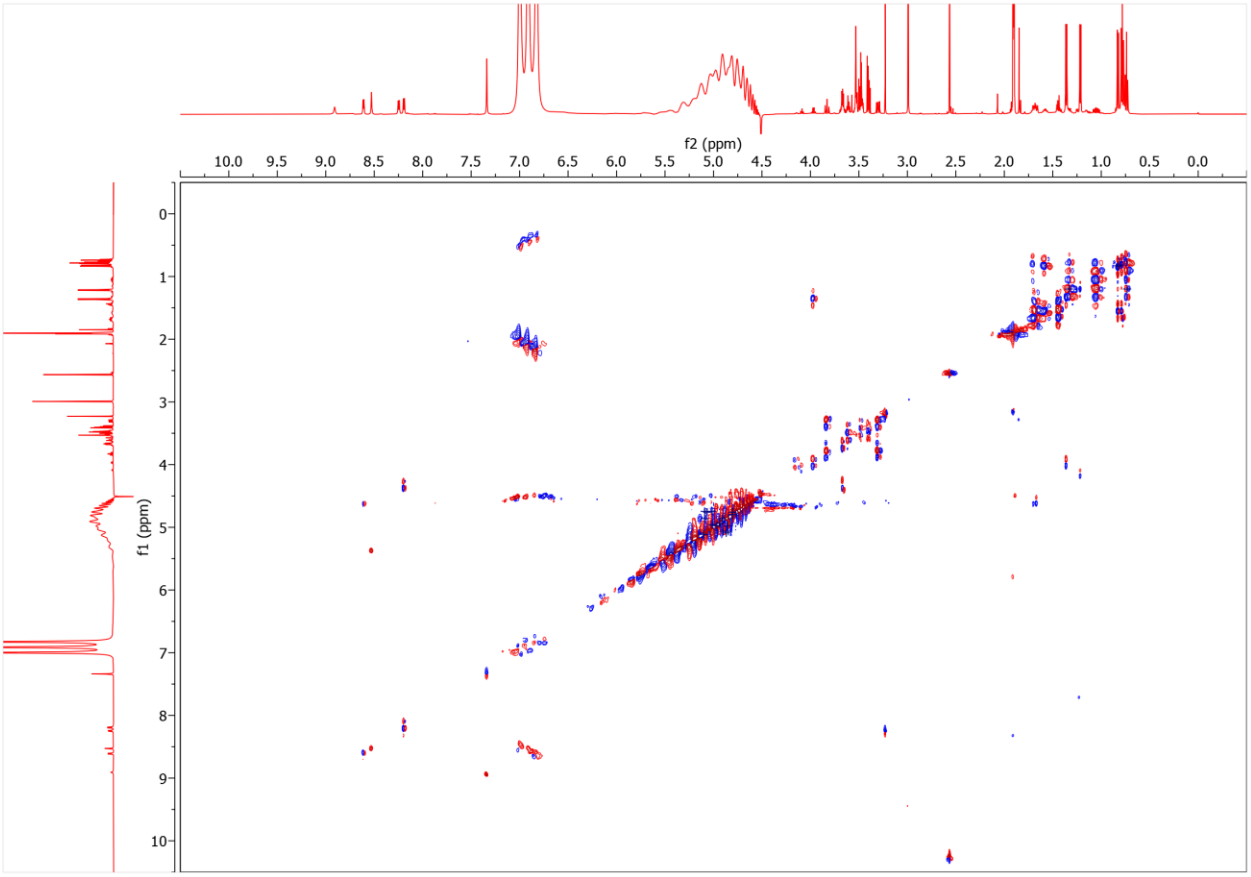

**Figure S28 (cont.). Collected NMR spectra for Diz-modified peptide.** All spectra were collected in a 79.9:20:0.1 $H_2O$:$CD_3CN$:$CDO_2D$ solvent system using a 600 MHz spectrometer. a, $^1H$ NMR of Diz-modified peptide. b, $^1H$-$^1H$ DQF-COSY of Diz-modified peptide. c, $^1H$-$^1H$ TOCSY of Diz-modified peptide. d, $^1H$-$^{13}C$ HSQC of Diz-modified peptide. e, $^1H$-$^1H$ NOESY of Diz-modified peptide.

c

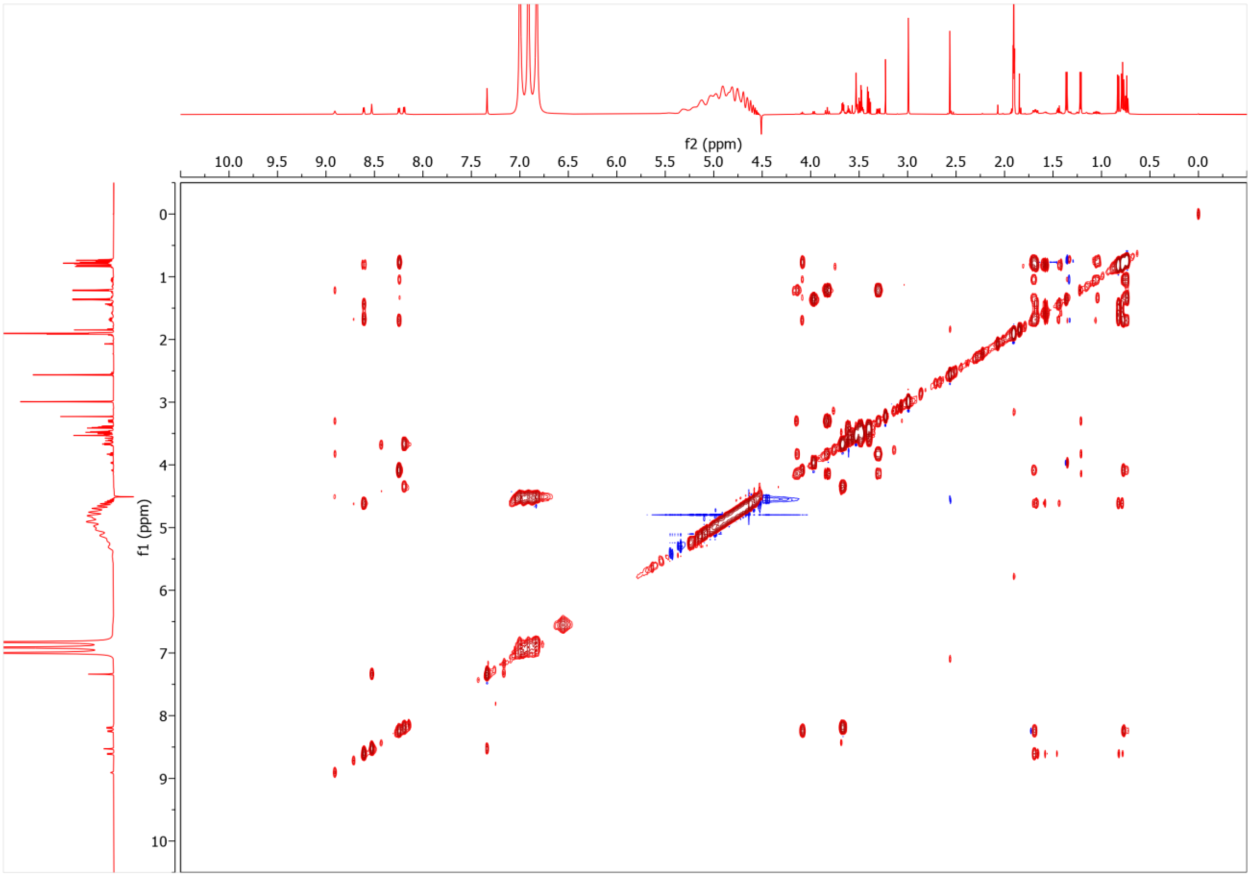

**Figure S28 (cont.). Collected NMR spectra for Diz-modified peptide.** All spectra were collected in a 79.9:20:0.1 $H_2O$:$CD_3CN$:$CDO_2D$ solvent system using a 600 MHz spectrometer. a, $^1$H NMR of Diz-modified peptide. b, $^1$H-$^1$H DQF-COSY of Diz-modified peptide. c, $^1$H-$^1$H TOCSY of Diz-modified peptide. d, $^1$H-$^{13}$C HSQC of Diz-modified peptide. e, $^1$H-$^1$H NOESY of Diz-modified peptide.

**d**

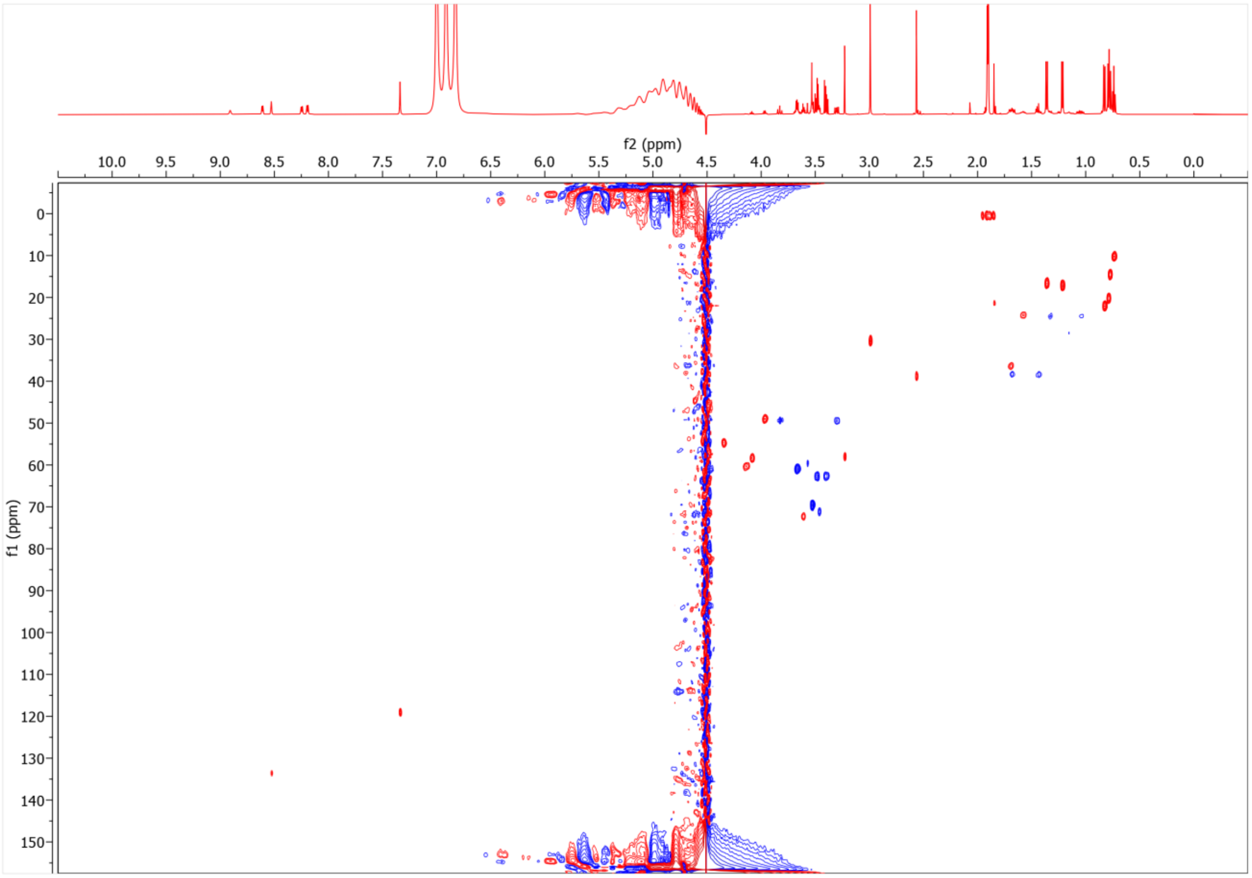

**Figure S28 (cont.). Collected NMR spectra for Diz-modified peptide.** All spectra were collected in a 79.9:20:0.1 $H_2O$:$CD_3CN$:$CDO_2D$ solvent system using a 600 MHz spectrometer. a, $^1$H NMR of Diz-modified peptide. b, $^1$H-$^1$H DQF-COSY of Diz-modified peptide. c, $^1$H-$^1$H TOCSY of Diz-modified peptide. d, $^1$H-$^{13}$C HSQC of Diz-modified peptide. e, $^1$H-$^1$H NOESY of Diz-modified peptide.

**e**

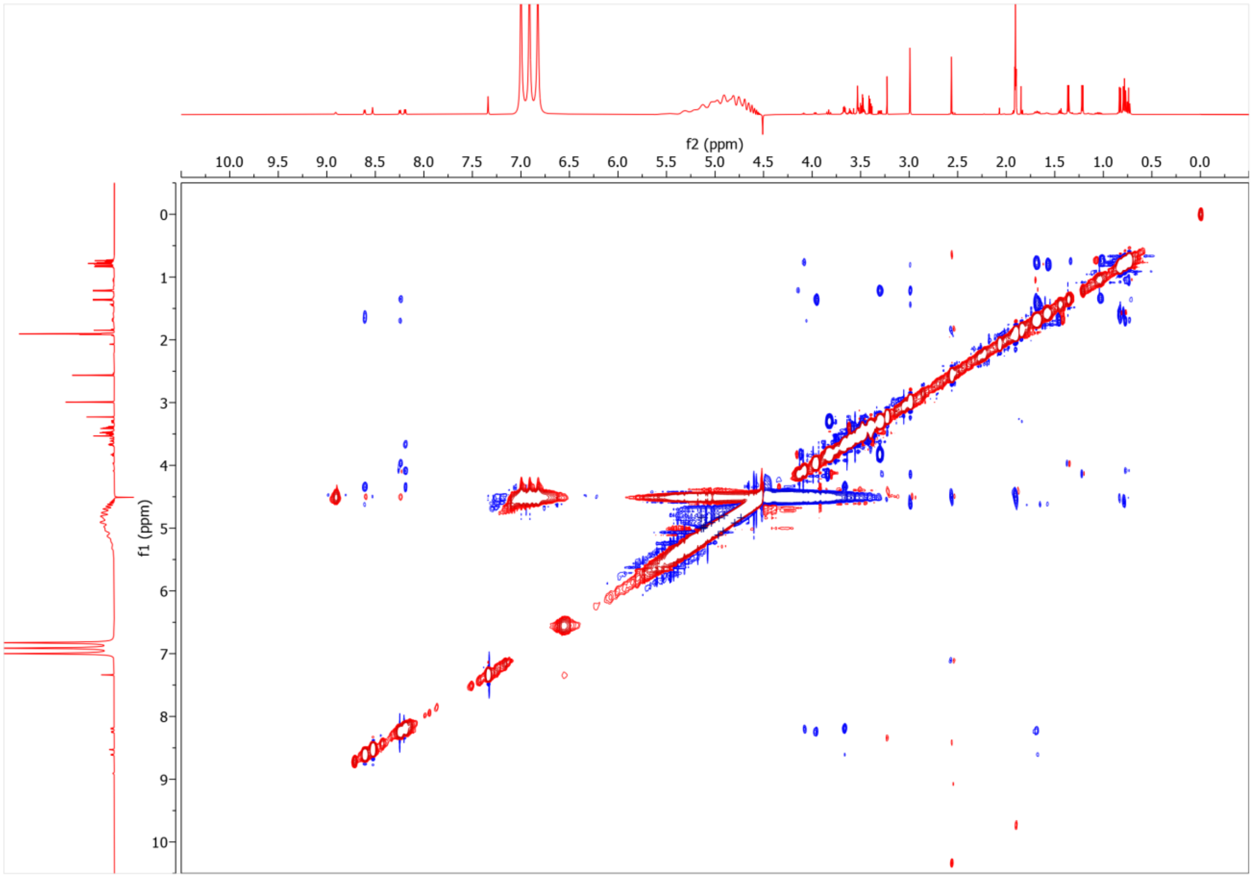

**Figure S28 (cont.). Collected NMR spectra for Diz-modified peptide.** All spectra were collected in a 79.9:20:0.1 $H_2O$:$CD_3CN$:$CDO_2D$ solvent system using a 600 MHz spectrometer. a, $^{1}H$ NMR of Diz-modified peptide. b, $^{1}H$-$^{1}H$ DQF-COSY of Diz-modified peptide. c, $^{1}H$-$^{1}H$ TOCSY of Diz-modified peptide. d, $^{1}H$-$^{13}C$ HSQC of Diz-modified peptide. e, $^{1}H$-$^{1}H$ NOESY of Diz-modified peptide.

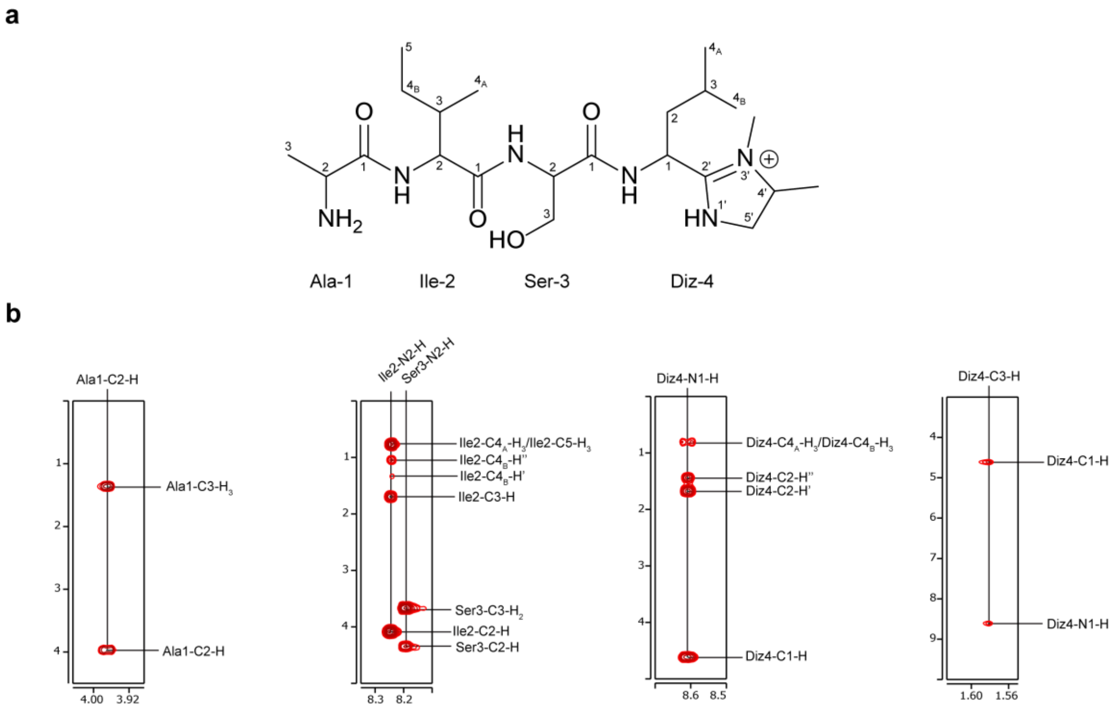

**Figure S29. Key correlations for assignment of Ala, Ile, Ser, and aminoisopentane in Diz-modified peptide.** a, Formal atom numbering for Diz-modified product. b, Key correlations observed in $^{1}H$-$^{1}H$ TOCSY spectrum.

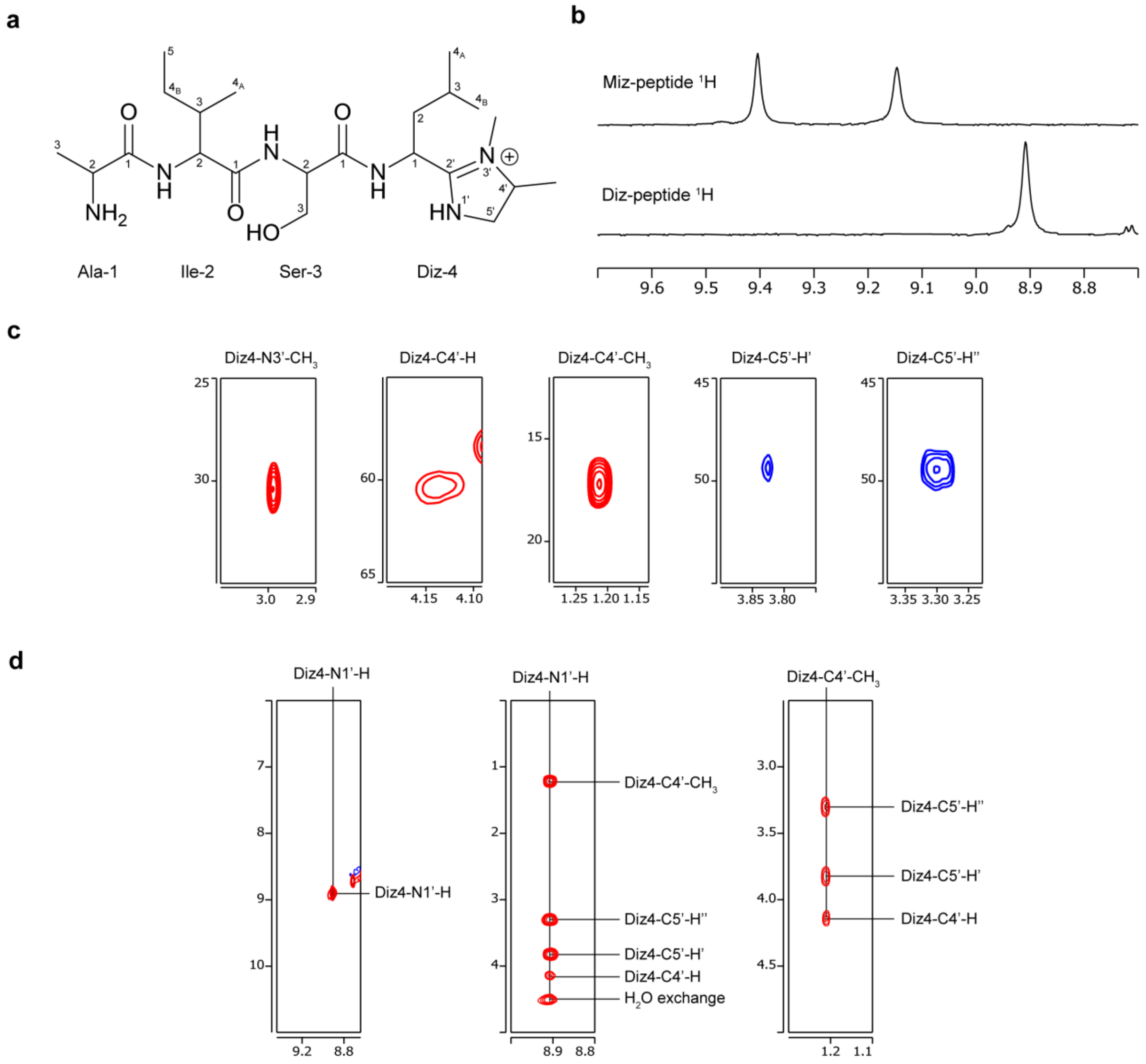

**Figure S30. Key correlations for assignment of dimethylimidazoline in Diz-modified peptide.** a, Formal atom numbering for Diz-modified product. b, Comparison of $^{1}$H NMR for Miz-modified peptide and Diz-modified peptide in the amidine region. c, Key correlations observed in $^{1}$H-$^{13}$C HSQC spectrum. d, Key correlations observed in $^{1}$H-$^{1}$H TOCSY spectrum.

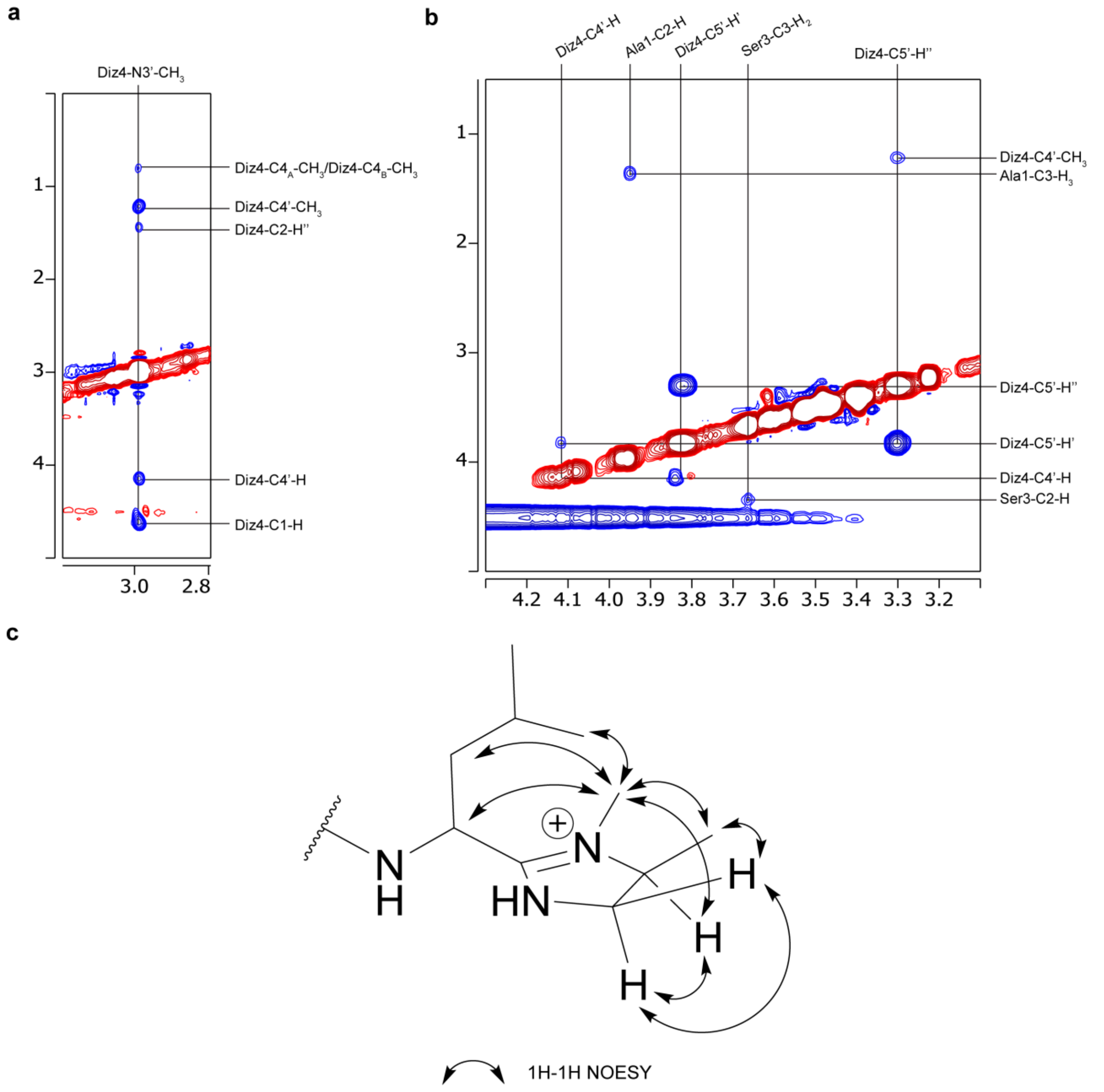

**Figure S31. Key $^1$H-$^1$H NOESY correlations for N-methylation in Diz-modified peptide.** a, $^1$H-$^1$H NOESY correlations for Diz4-N3'-$CH_3$. b, $^1$H-$^1$H NOESY correlations that were used to assist structural assignment of the Diz-modified peptide. c, Annotated $^1$H-$^1$H NOESY correlations for the Diz residue.

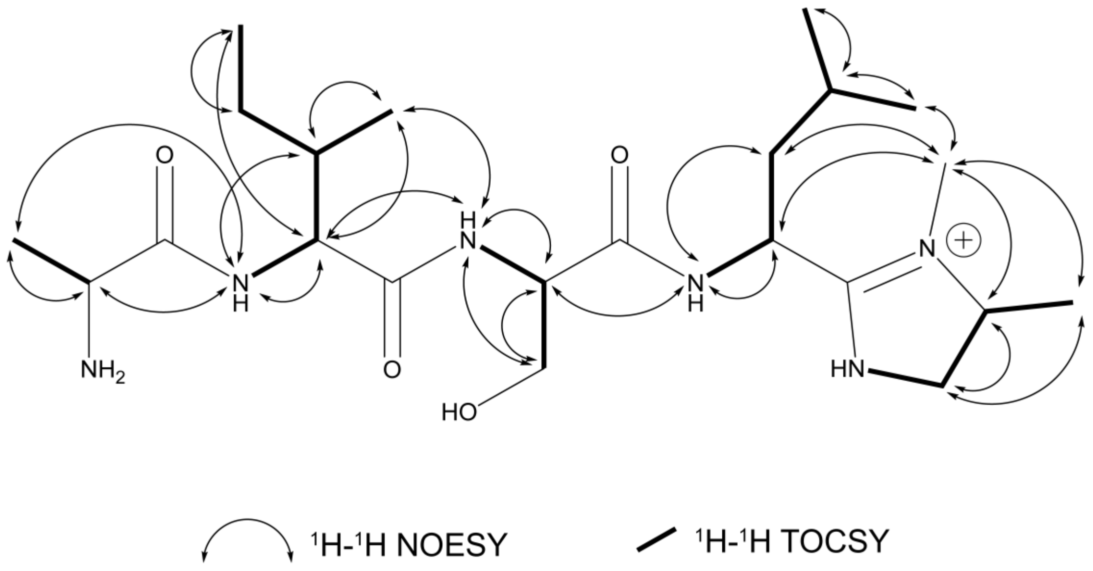

**Figure S32. Annotated structure of Diz-modified peptide.**

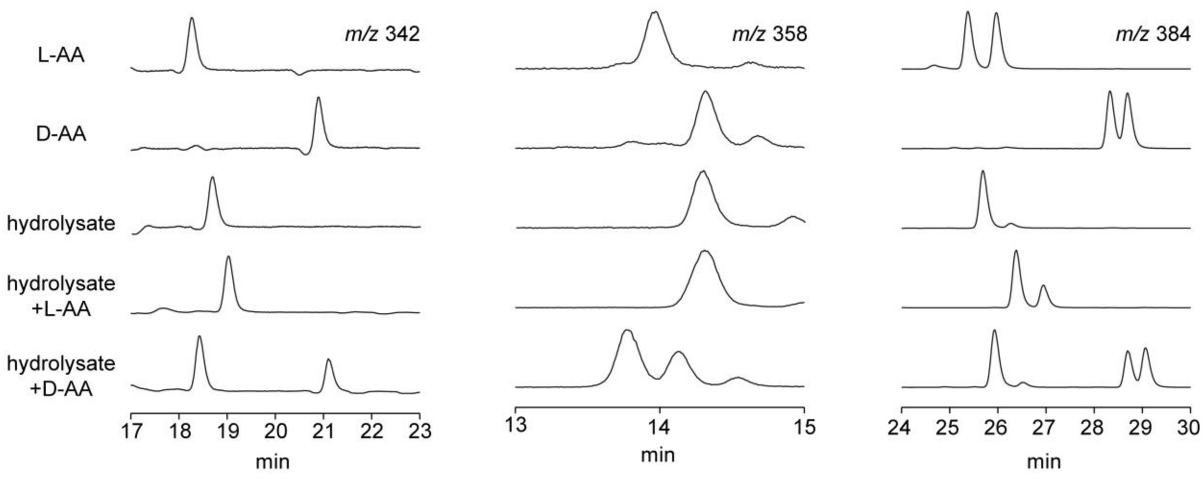

**Figure S33. Determination of stereochemistry in Diz-modified peptide using LC-MS.** Ion-count normalized extracted ion chromatograms (EICs) for Ala (*m/z* 342), Ser (*m/z* 358), and Ile/Leu (*m/z* 384) are shown. For each mass of interest, EICs were generated for the following five samples: L-FDAA-derivatized L-amino acids, D-FDAA-derivatized L-amino acids, L-FDAA-derivatized hydrolysate of Diz-modified peptide, co-injection of L-amino acid and hydrolysate samples, and co-injection of D-amino acid and hydrolysate samples. Note 1: D-FDAA derivatization of L-amino acids produces compounds which are chromatographically identical to L-FDAA derivatization of D-amino acids when analyzed on an achiral stationary phase. This allowed us to use D-FDAA-derivatized L-amino acids as a D-amino acid standard. Note 2: Significant retention time drift was apparent across different injections, but ambiguities in isomer assignment were resolved by co-injections. Note 3: The assigned Ile peak and Leu peak are not equivalent in intensity, despite each occurring in the peptide sequence once. We suspect the decreased intensity of the Leu peak is due to potential degradation and/or incomplete hydrolysis resulting from the increased number of hydrolytic events required to restore Leu from the modified peptide.

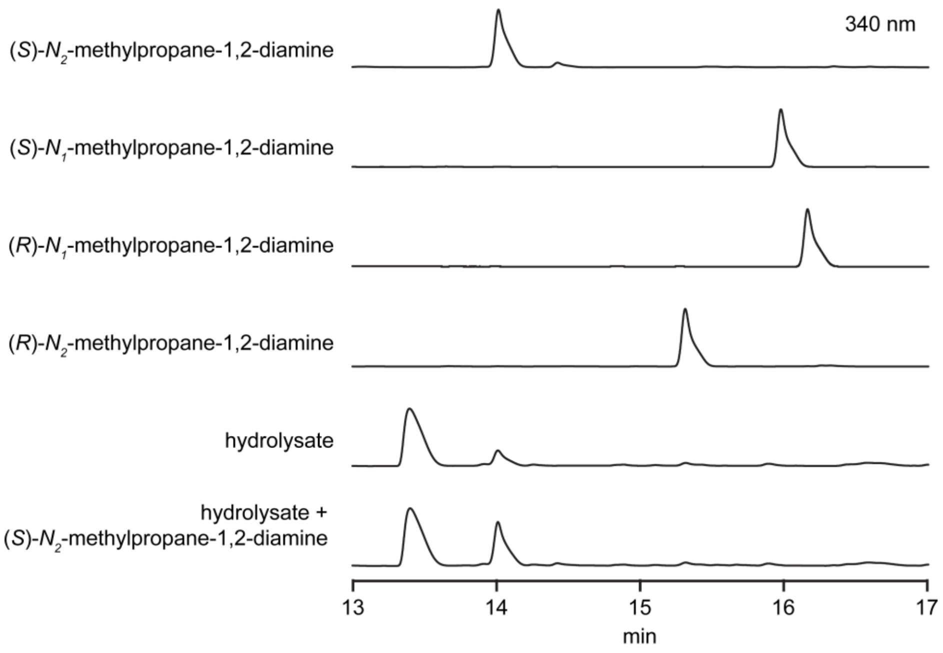

**Figure S34. Marfey's analysis of *N*-methylpropane-1,2-diamine isomers.** Absorbance-normalized chromatograms are shown for injections of L-FDAA-derivatized isomers of *N*-methylpropane-1,2-diamine and Diz-modified peptide hydrolysate. Analytes were monitored by UV absorbance at 340 nm.

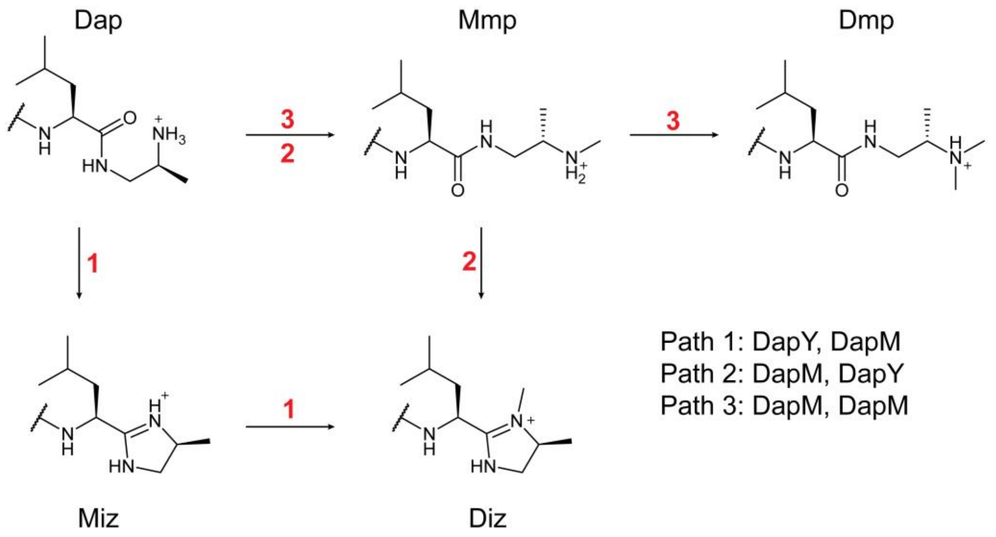

**Figure S35. Potential routes of modification by DapMY.** Abbreviations: Dap, 1,2-diaminopropane; Mmp, monomethylpropane-1,2-diamine; Dmp, dimethylpropane-1,2-diamine; Miz, 4-methylimidazoline; Diz, 3,4-dimethylimidazoline.

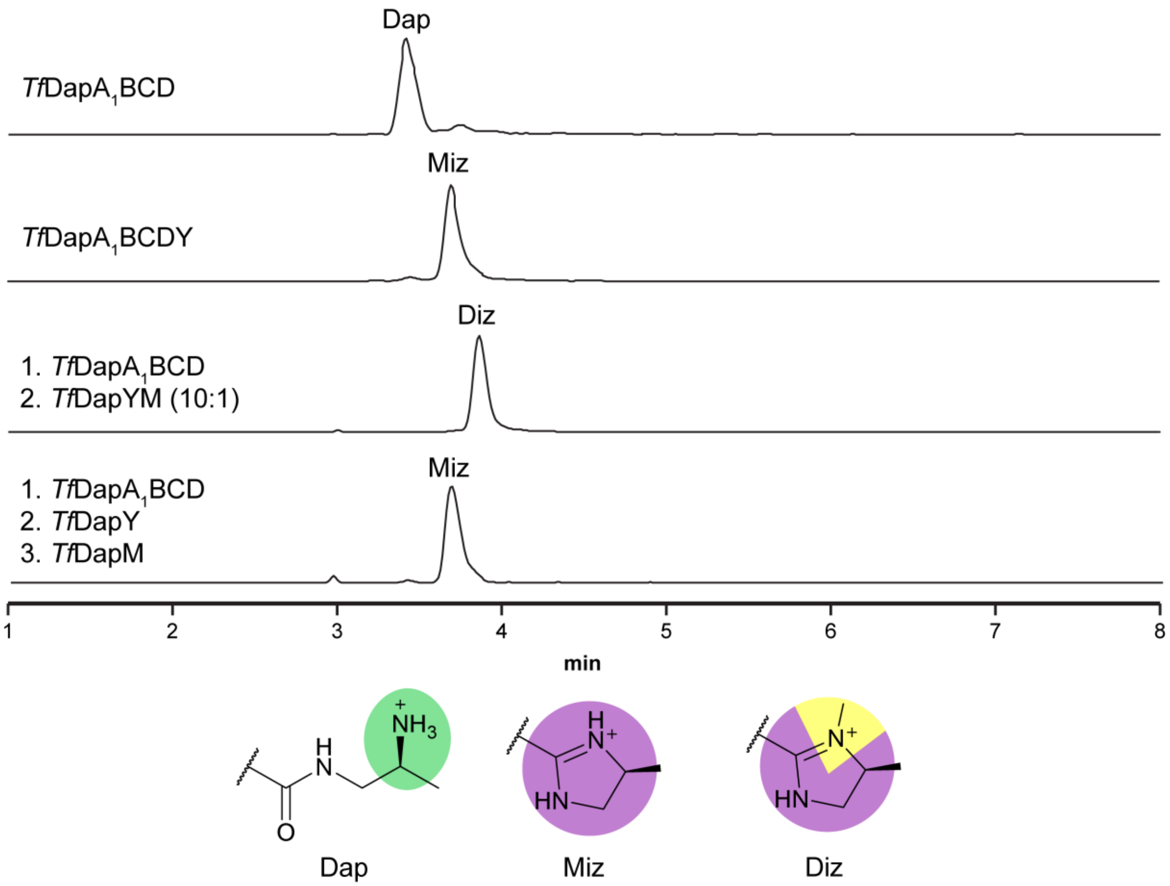

**Figure S36. Sequential reactions with *Tf*DapMY.** a, Extracted ion chromatograms are shown for only the expected products of each reaction. Extracted ions: Dap, 1666.8172 *m/z*; Miz, 1660.8137 *m/z*; Diz, 1665.4856 *m/z*. Abbreviations: Dap, 1,2-diaminopropane; Miz, 4-methylimidazoline; Diz, 3,4-dimethylimidazoline.

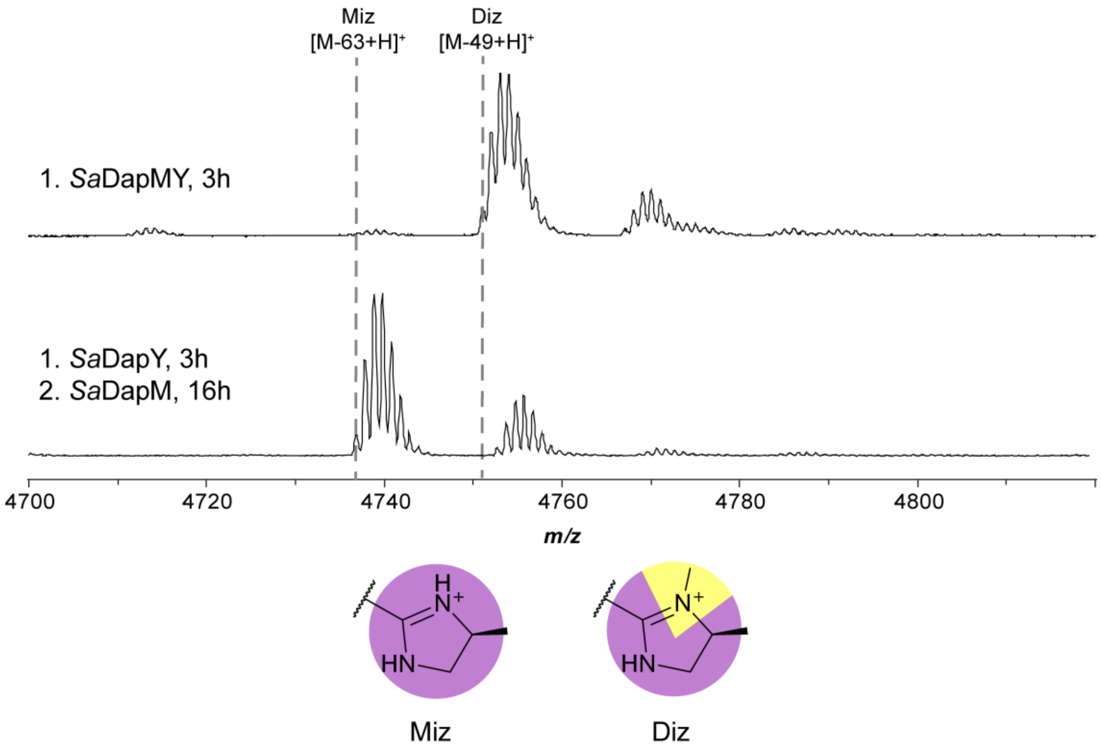

**Figure S37. Sequential reactions with *Sa*DapMY.** Reactions were performed using recombinantly expressed (MBP)*Sa*DapA1-Dap as substrate. Spectra were collected after TEV proteolysis and desalting by C18 ZipTip. Abbreviations: Dap, 1,2-diaminopropane; Miz, 4-methylimidazoline; Diz, 3,4-dimethylimidazoline. Key expected masses: *Sa*DapA1-Miz, 4737 Da; *Sa*DapA1-Diz, 4751 Da.

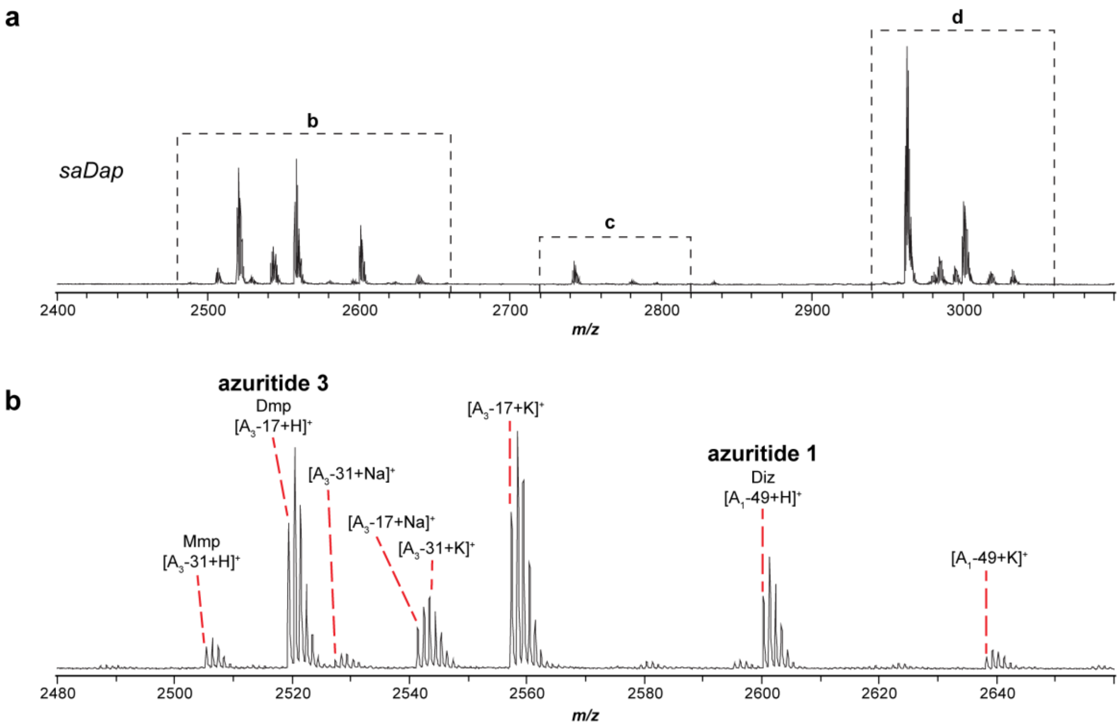

**Figure S38. Heterologous over-expression of azuritides 1-4.** a, MALDI-TOF mass spectrum of methanolic cell extract for heterologous over-expression of *saDap* BGC. b, c, d, Zoom-ins of boxed regions in panel a with annotations. e, f, g, h, MALDI-LIFT-TOF/TOF-MS for azuritides 1, 2, 3, and 4, respectively. Azuritides 1-4 were assigned as the most abundant product detected for each respective precursor peptide encoded by the *saDap* BGC. Abbreviations: Mmp, monomethylpropane-1,2-diamine; Dmp, dimethylpropane-1,2-diamine; Diz, 3,4-dimethylimidazoline. Key expected masses: azuritide 1 (*Sa*Dap$A_1$-Diz), 2600 Da; azuritide 2 (*Sa*Dap$A_2$-Diz), 2741 Da; azuritide 3 (*Sa*Dap$A_3$-Dmp), 2519 Da; azuritide 4 (*Sa*Dap$A_4$-Diz), 2962 Da; *Sa*Dap$A_3$-Mmp, 2505 Da; *Sa*Dap$A_4$-Mmp, 2980 Da; *Sa*Dap$A_4$-Dmp, 2994 Da. Sodium adducts result in addition of 22 Da. Potassium adducts result in addition of 38 Da.

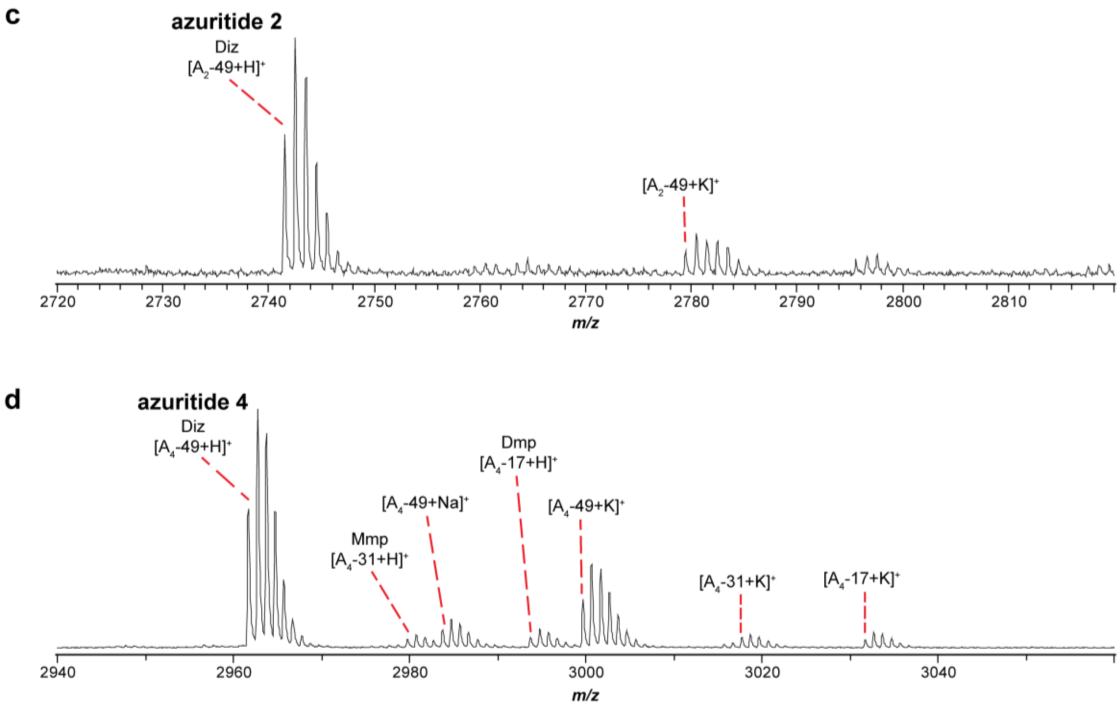

**Figure S38 (cont.). Heterologous over-expression of azuritides 1-4.**

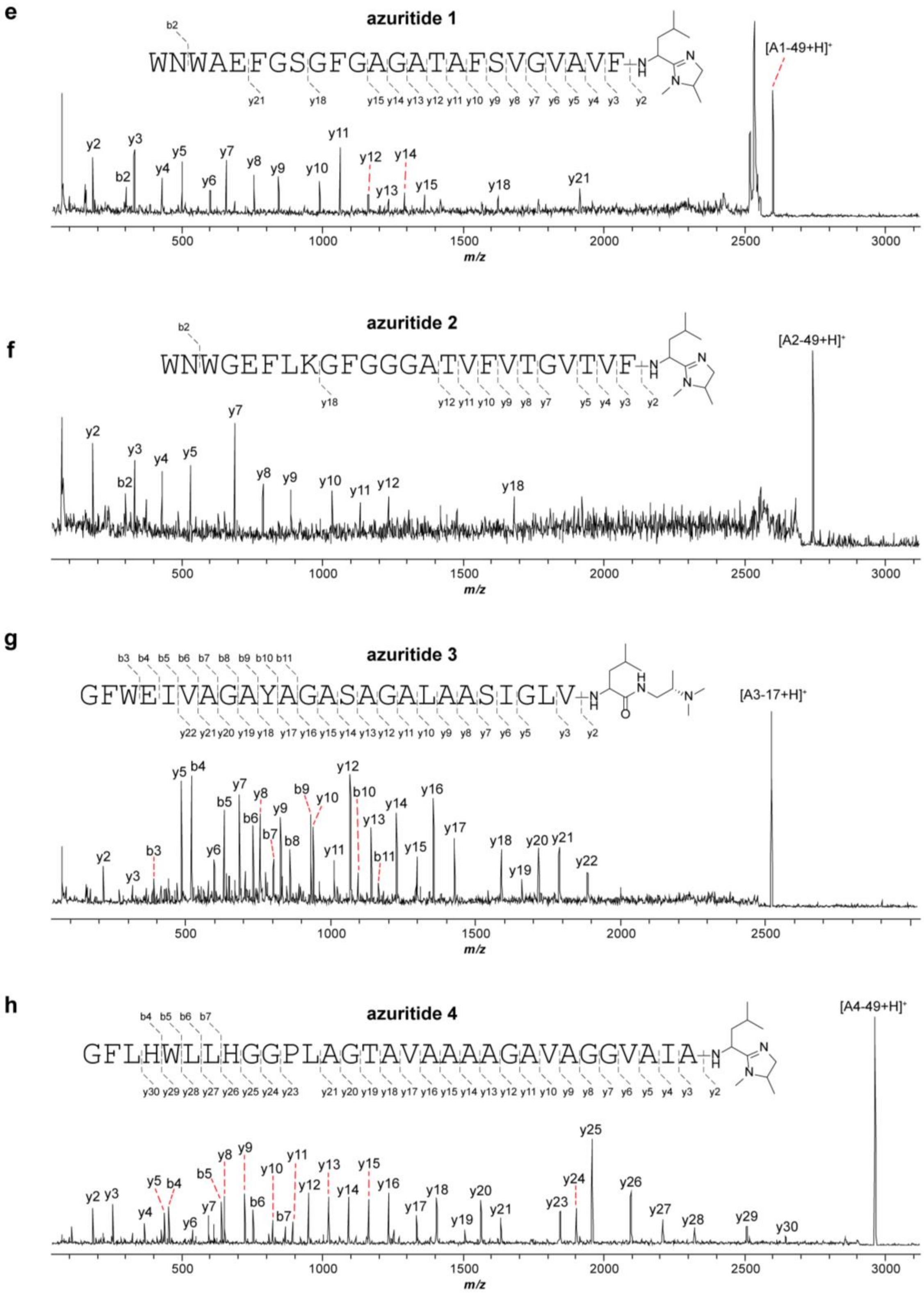

**Figure S38 (cont.). Heterologous over-expression of azuritides 1-4.**

```
CLUSTAL format alignment by MAFFT FFT-NS-i (v7.525)

                        leader                          core
MpaA1           MNE----STEMSIRFQELDPMEAPSWDS---------FYQGVIGALVLIGIGAAAAT
MpaA2           MNA-----ALPALEFTELEAMDAPGDAD-----------DYIRGFAVGVIIGILALT
MpaA3           MQS-------TSLEFLELEQIDTPLEWW-----------EHASYIIAIIGGAAAIAT
ScaA1           MQEIEKIQPVLELG--ELEAMDAPGWWT--AAGVSAGVVVSASAAYGSAALSAAAIT
ScaA2           MQETEKIQPVLELGMQELEAMEAPGFWT---GFSYGVAVSGTAAASAAVSVAASIAT
SaDapA1         ------MSTDLDLQFEELETLDALWNWA-------EFGSGFGAGATAFSVGVAVFLT
SaDapA2         ----------MELQVEELESLEALWNWG-------EFLKGFGGGATVFVTGVTVFLT
SaDapA3         ------MENVTELQIEELEPMQAPGFWE-I-----VAGAYAGASAGALAASIGLVLT
SaDapA4         MNEHVNNPADTAFEVEELEGLHAPGFLHWLLHGGPLAGTAVAAAAGAVAGGVAIALT
                           :   **: :.:                                  *
```

**Figure S39. Comparison of cleavage sites for characterized daptides.**

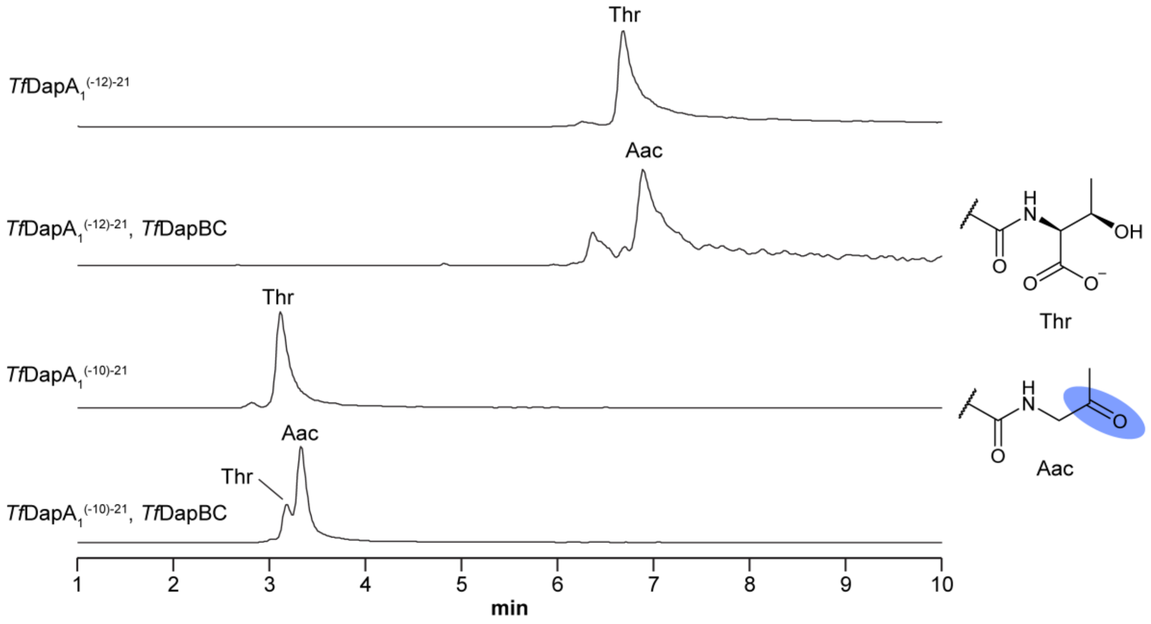

*Tf*DapA$_1^{(-12)-21}$: `KLEXQELEAXEAPKGFWTGAGVGALISTSFAISLT`

*Tf*DapA$_1^{(-10)-21}$: `KXQELEAXEAPKGFWTGAGVGALISTSFAISLT`

**Figure S40. Evaluation of substrates with shortened leader regions.** Extracted ion chromatograms are shown for products of each reaction. Extracted ions: *Tf*DapA$_1^{(-12)—21}$, 1216.9917 *m/z*; *Tf*DapA$_1^{(-12)—21}$-Aac, 1201.6565 *m/z*; *Tf*DapA$_1^{(-10)—21}$, 1136.2828 *m/z*; *Tf*DapA$_1^{(-10)—21}$-Aac, 1120.9476 *m/z*. X = Norleucine. Abbreviations: Aac, aminoacetone.

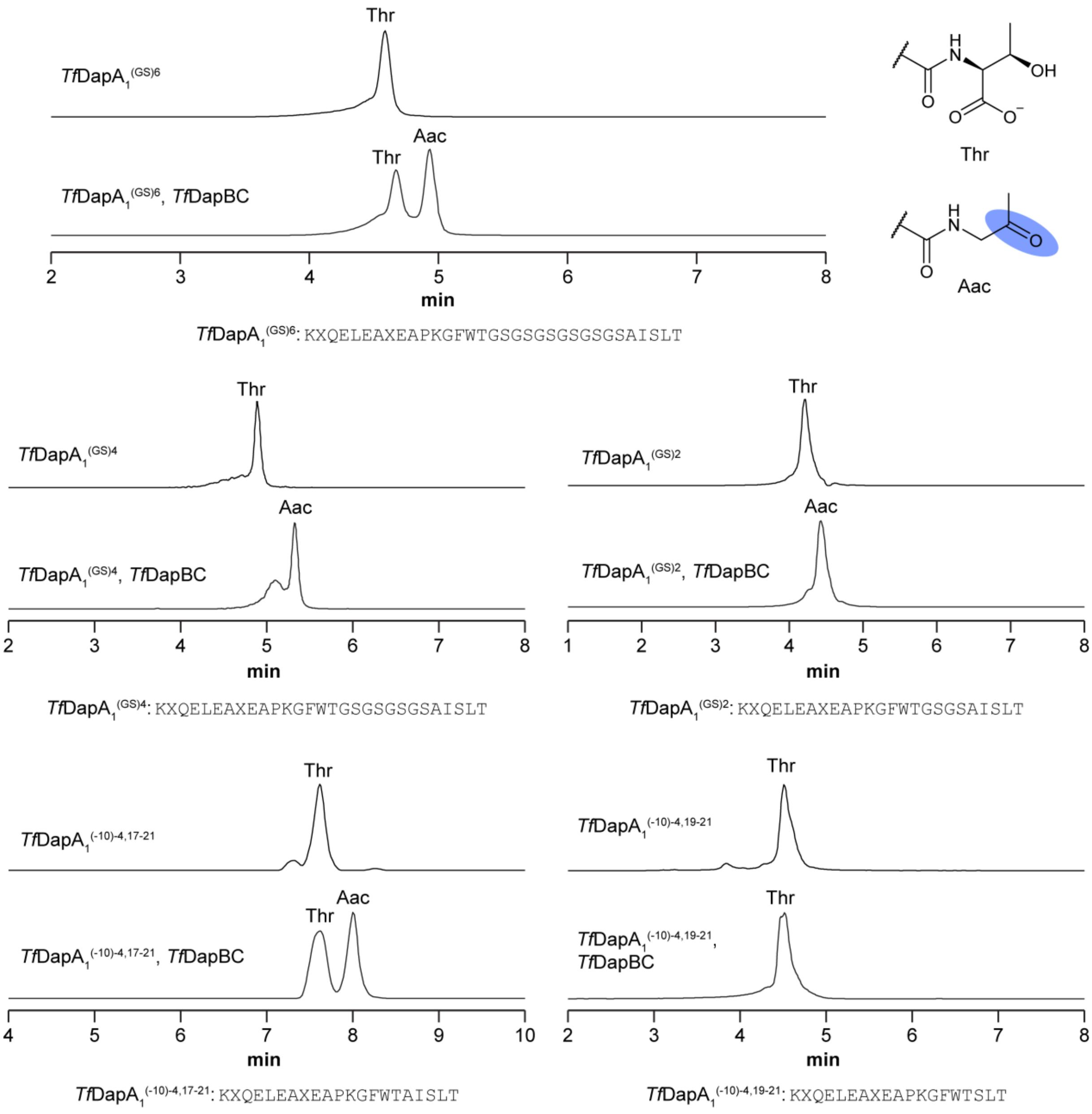

**Figure S41. Evaluation of substrates with shortened core regions.** Extracted ion chromatograms are shown for products of each reaction. Extracted ions: *Tf*DapA$_1^{(GS)6}$, 1070.8713 *m/z*; *Tf*DapA$_1^{(GS)6}$-Aac, 1055.5362 *m/z*; *Tf*DapA$_1^{(GS)4}$, 974.8357 *m/z*; *Tf*DapA$_1^{(GS)4}$-Aac, 959.5005 *m/z*; *Tf*DapA$_1^{(GS)2}$, 1317.6964 *m/z*; *Tf*DapA$_1^{(GS)2}$-Aac, 1294.6936 *m/z*; *Tf*DapA$_1^{(-10)—4,17—21}$, 1173.6429 *m/z*; *Tf*DapA$_1^{(-10)—4,17—21}$-Aac, 1150.6401 *m/z*; *Tf*DapA$_1^{(-10)—4,19—21}$, 1081.5823 *m/z*; *Tf*DapA$_1^{(-10)—4,19—21}$-Aac, 1058.5796 *m/z*. X = Norleucine. Abbreviations: Aac, aminoacetone.

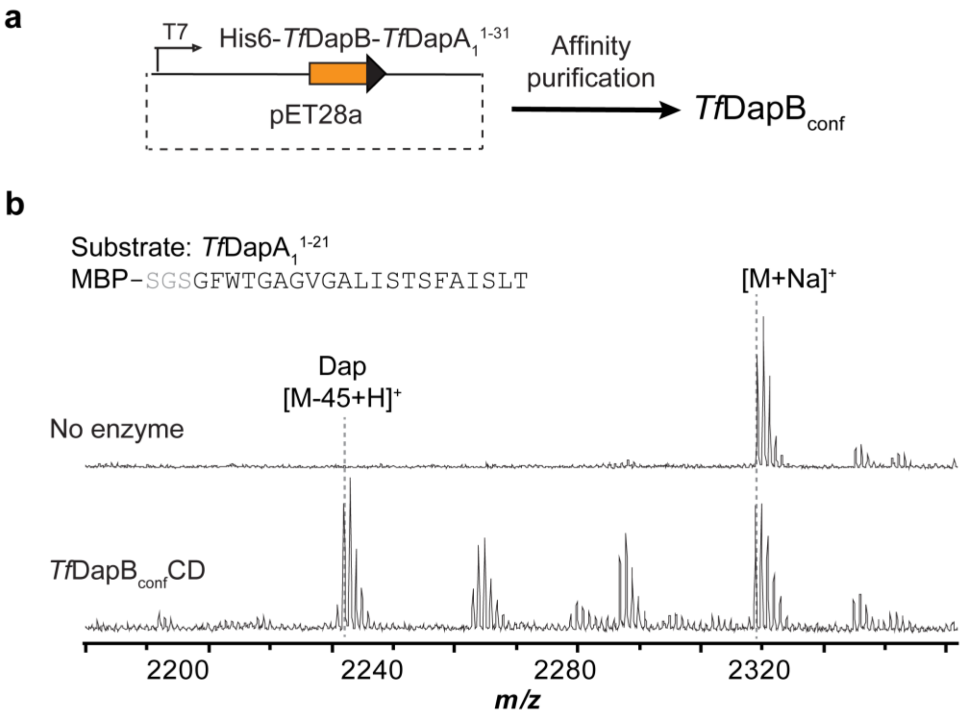

**Figure S42. Modification of core peptide variants using ConFusion enzymes.** MALDI-TOF mass spectra are shown for untreated (MBP)*Tf*DapA$_1$$^{1—21}$, (MBP)*Tf*DapA$_1$$^{7—21}$, and (MBP)*Tf*DapA$_1$$^{12—21}$. Below each untreated sample is the corresponding MALDI-TOF mass spectrum for peptide treated with *Tf*DapB$_{conf}$CD. A sample reaction scheme is provided with the first set of spectra. Spectra were collected following TEV proteolysis and desalting by C18 ZipTip. Abbreviations: Aac, aminoacetone; Dap, 1,2-diaminopropane. Key expected masses: *Tf*DapA$_1$$^{1—21}$, 2287 Da; *Tf*DapA$_1$$^{1—21}$-Aac, 2241 Da; *Tf*DapA$_1$$^{1—21}$-Dap, 2242 Da; *Tf*DapA$_1$$^{7—21}$, 1668 Da; *Tf*DapA$_1$$^{7—21}$-Aac, 1622 Da; *Tf*DapA$_1$$^{7—21}$-Dap, 1623 Da; *Tf*DapA$_1$$^{12—21}$, 1271 Da; *Tf*DapA$_1$$^{12—21}$-Aac, 1225 Da; *Tf*DapA$_1$$^{12—21}$-Dap, 1226 Da. Sodium adducts result in addition of 22 Da.

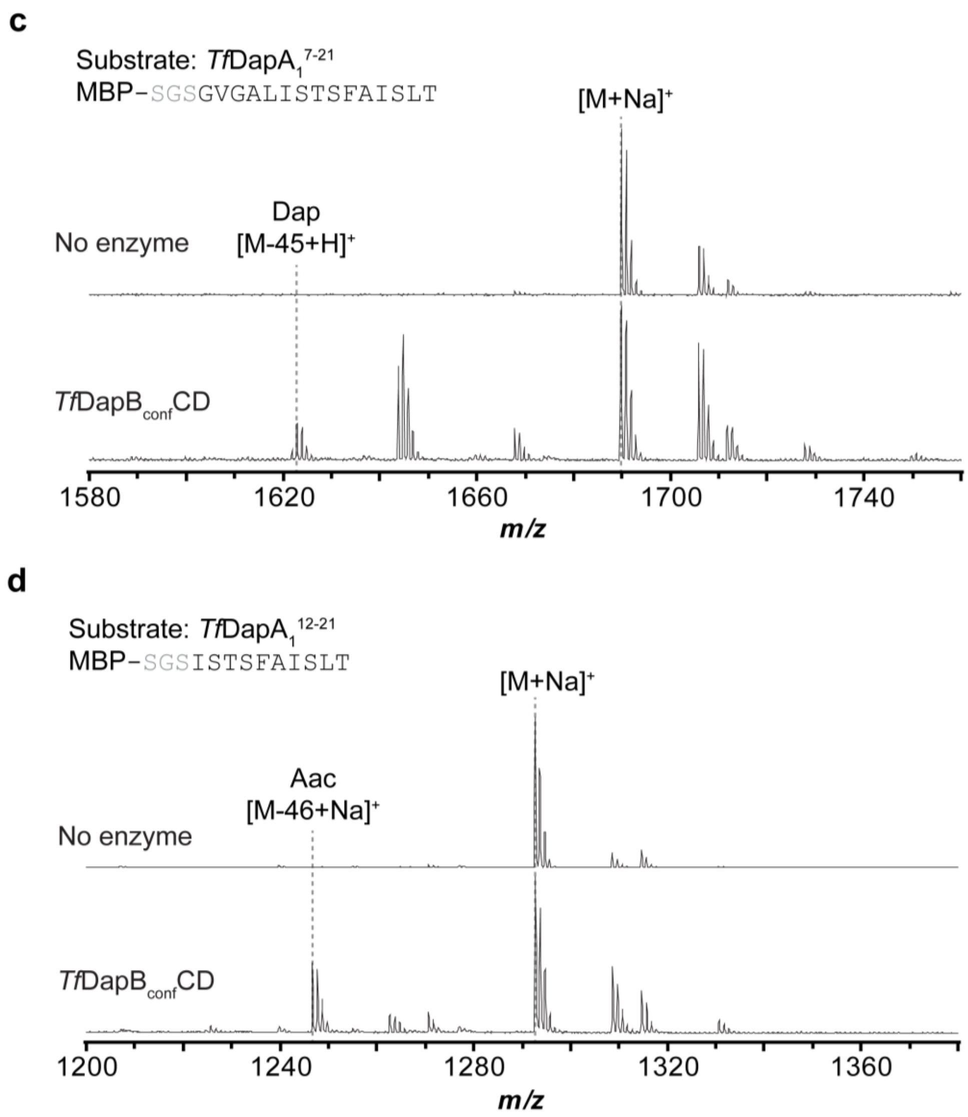

**Figure S42 (cont.). Modification of core peptide variants using ConFusion enzymes.**

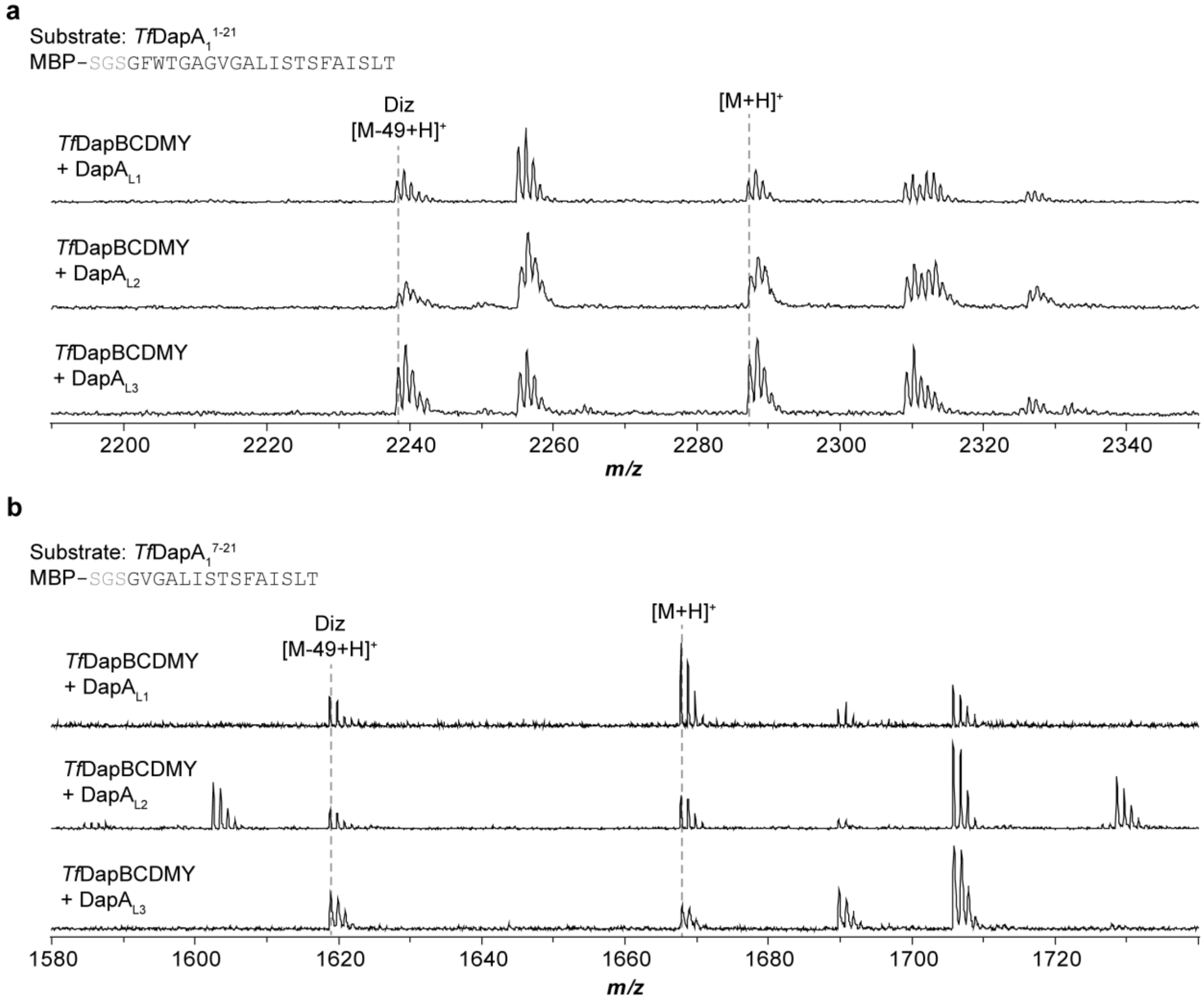

**Figure S43. Modifications installed using leader peptides provided *in trans*.** MALDI-TOF mass spectra are shown for reactions with a, (MBP)*Tf*DapA$_1^{1—21}$; b, (MBP)*Tf*DapA$_1^{7—21}$; and c, (MBP)*Tf*DapA$_1^{12—21}$. Each core variant was reacted separately with three different leader variants provided *in trans*. d, MALDI-TOF mass spectrum is shown for reaction of *Tf*DapBC with DapA$_{L1}$. Sequences for leader variants: DapA$_{L1}$, RXQELEAXEAPGFWT (X = norleucine); DapA$_{L2}$, MQELEAMEAPGFWTG; DapA$_{L3}$, MQELEAMEAPG. Spectra were collected following TEV proteolysis and desalting by C18 ZipTip. Key expected masses: *Tf*DapA$_1^{1—21}$, 2287 Da; *Tf*DapA$_1^{1—21}$-Aac, 2241 Da; *Tf*DapA$_1^{1—21}$-Dap, 2242 Da; *Tf*DapA$_1^{7—21}$, 1668 Da; *Tf*DapA$_1^{7—21}$-Aac, 1622 Da; *Tf*DapA$_1^{7—21}$-Dap, 1623 Da; *Tf*DapA$_1^{12—21}$, 1271 Da; *Tf*DapA$_1^{12—21}$-Aac, 1225 Da; *Tf*DapA$_1^{12—21}$-Dap, 1226 Da; DapA$_{L1}$, 1760 Da; DapA$_{L1}$-Aac, 1714. Sodium adducts result in addition of 22 Da.

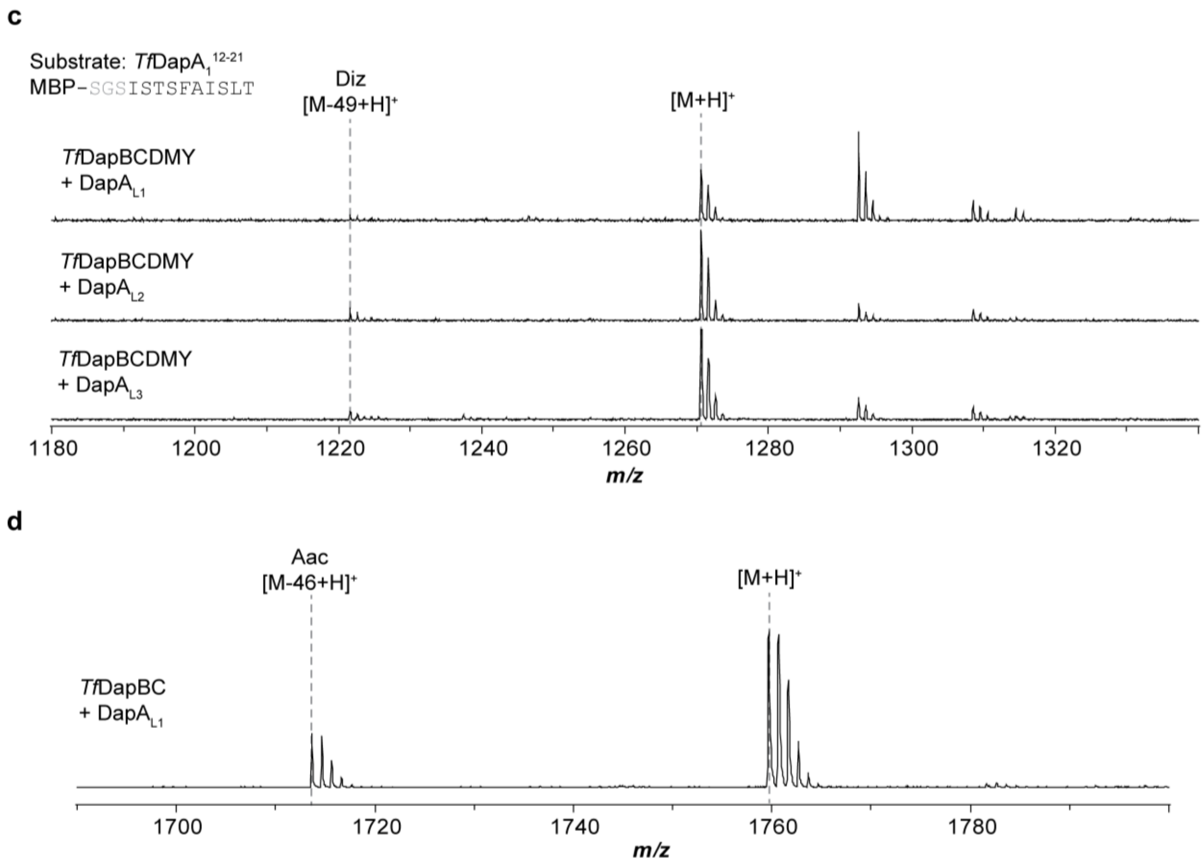

**Figure S43 (cont.). Modifications installed using leader peptides provided in trans.**

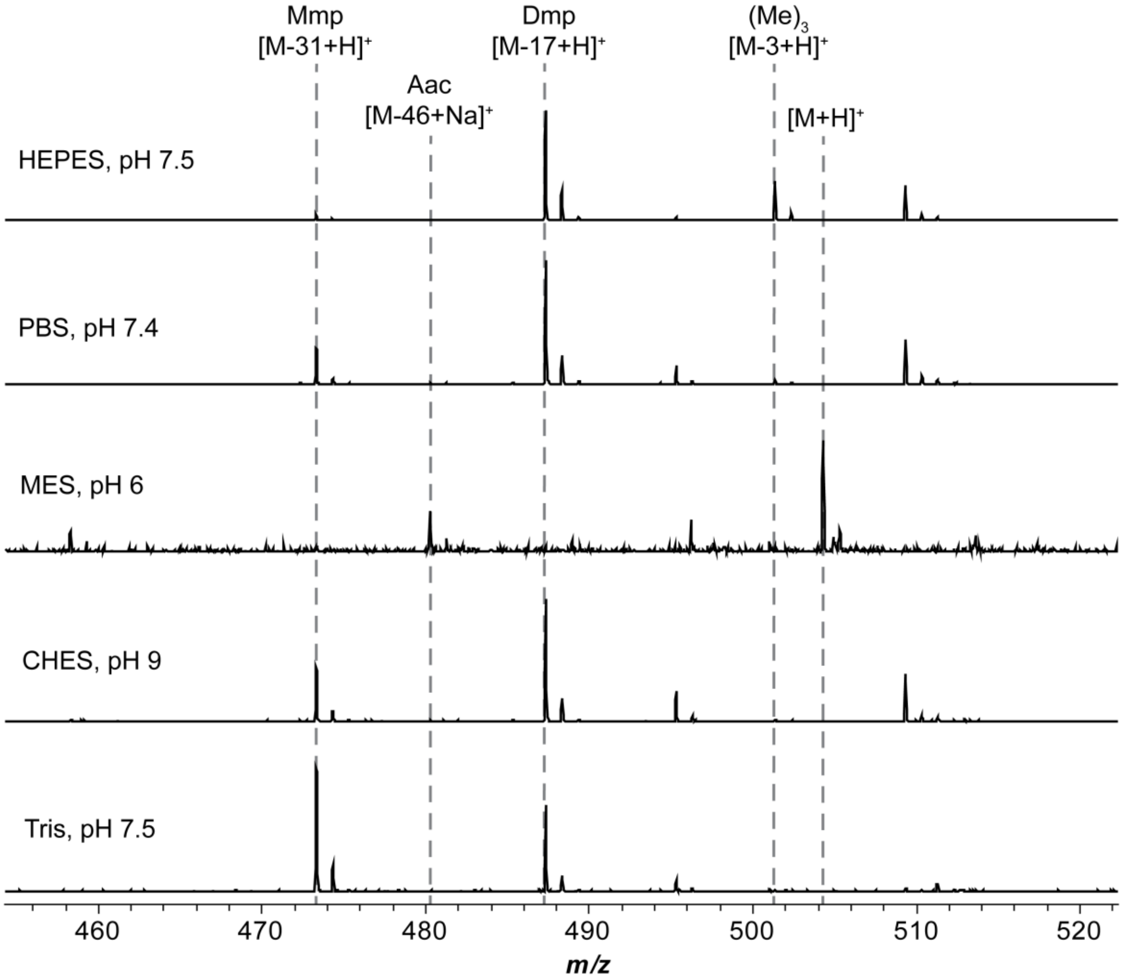

**Figure S44. Examination of different buffer systems.** MALDI-TOF mass spectra are shown for reactions of DapA$_{eng}$ with *Tf*DapBCDM using a panel of different buffer compositions. Spectra were collected following TEV proteolysis and desalting by C18 ZipTip. Abbreviations: Aac, aminoacetone; Mmp, monomethylpropane-1,2-diamine; Dmp, dimethylpropane-1,2-diamine. Key expected masses: AISLT, 504 Da; AISL-Aac, 458 Da; AISL-Mmp, 473 Da; AISL-Dmp, 487 Da.

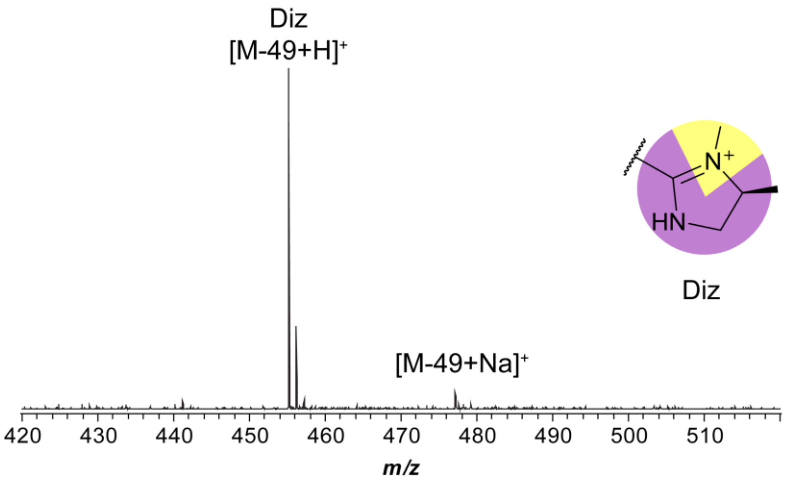

**Figure S45. Modification of DapA$_{eng}$ after lyophilization of *Tf*DapBCDMY.** MALDI-TOF mass spectrum is shown for the reaction of DapA$_{eng}$ with *Tf*DapBCDMY after the enzymes were subjected to lyophilization. Spectra were collected following TEV proteolysis and desalting by C18 ZipTip. Abbreviations: Diz, 3,4-dimethylimidazoline. Key expected masses: AIS-Diz, 455 Da. Sodium adducts result in an increase of 22 Da.

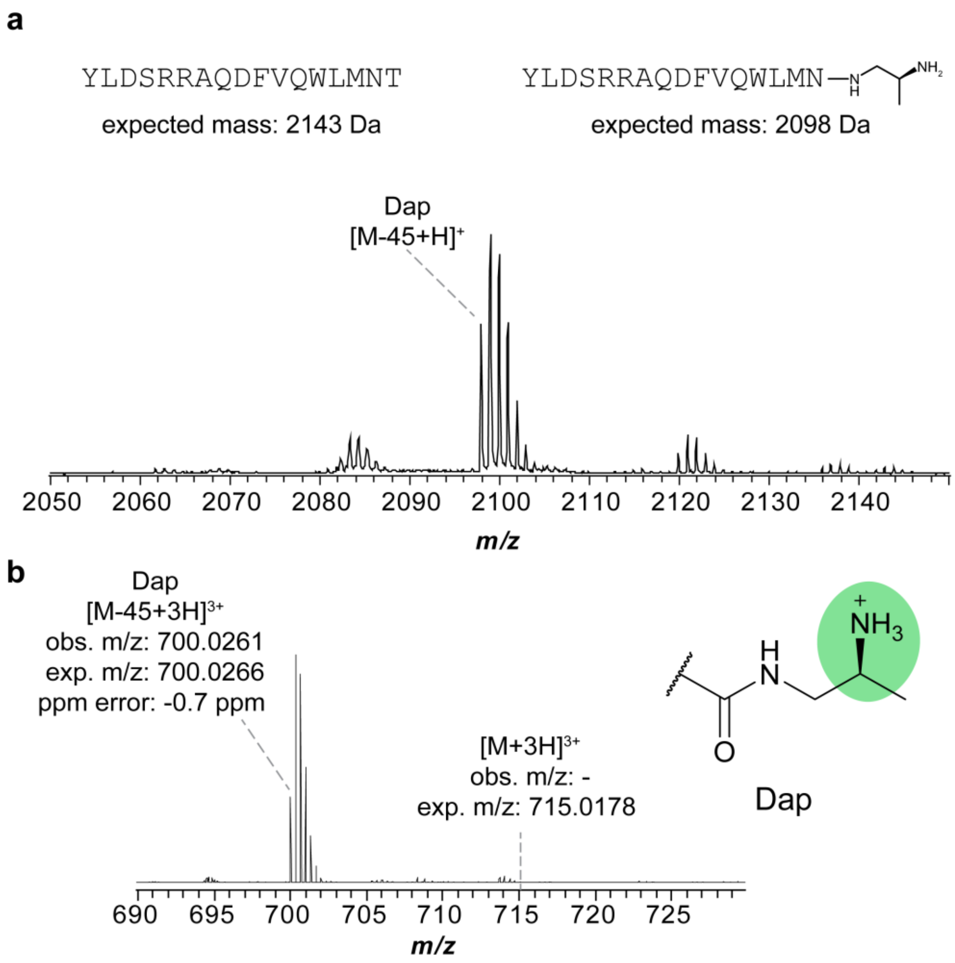

**Figure S46. LysC digestion of DapA$_{glucagon}$.** a, MALDI-TOF mass spectrum of LysC digested (MBP)DapA$_{glucagon}$. Abbreviations: Dap, 1,2-diaminopropane. Key expected masses: glucagon$^{13—29}$-Dap, 2098 Da. b, HR mass spectrum for LysC digested (MBP)DapA$_{glucagon}$.

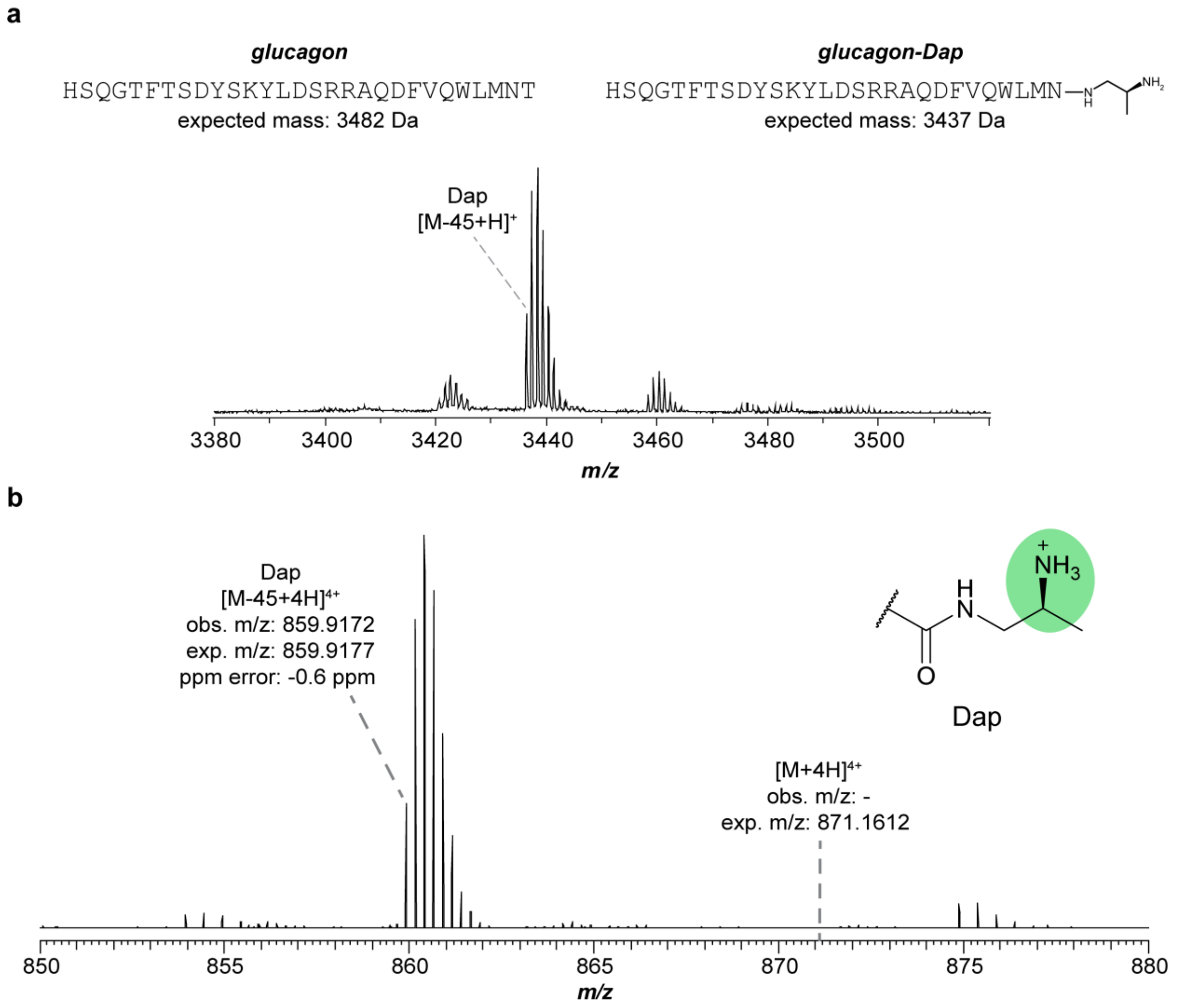

**Figure S47. GluC digestion of DapA$_{\mathbf{glucagon}}$.** a, MALDI-TOF mass spectrum of GluC digested (MBP)DapA$_{glucagon}$. Abbreviations: Dap, 1,2-diaminopropane. Key expected masses: glucagon-Dap, 3437 Da. b, HR mass spectrum for GluC digested (MBP)DapA$_{glucagon}$.

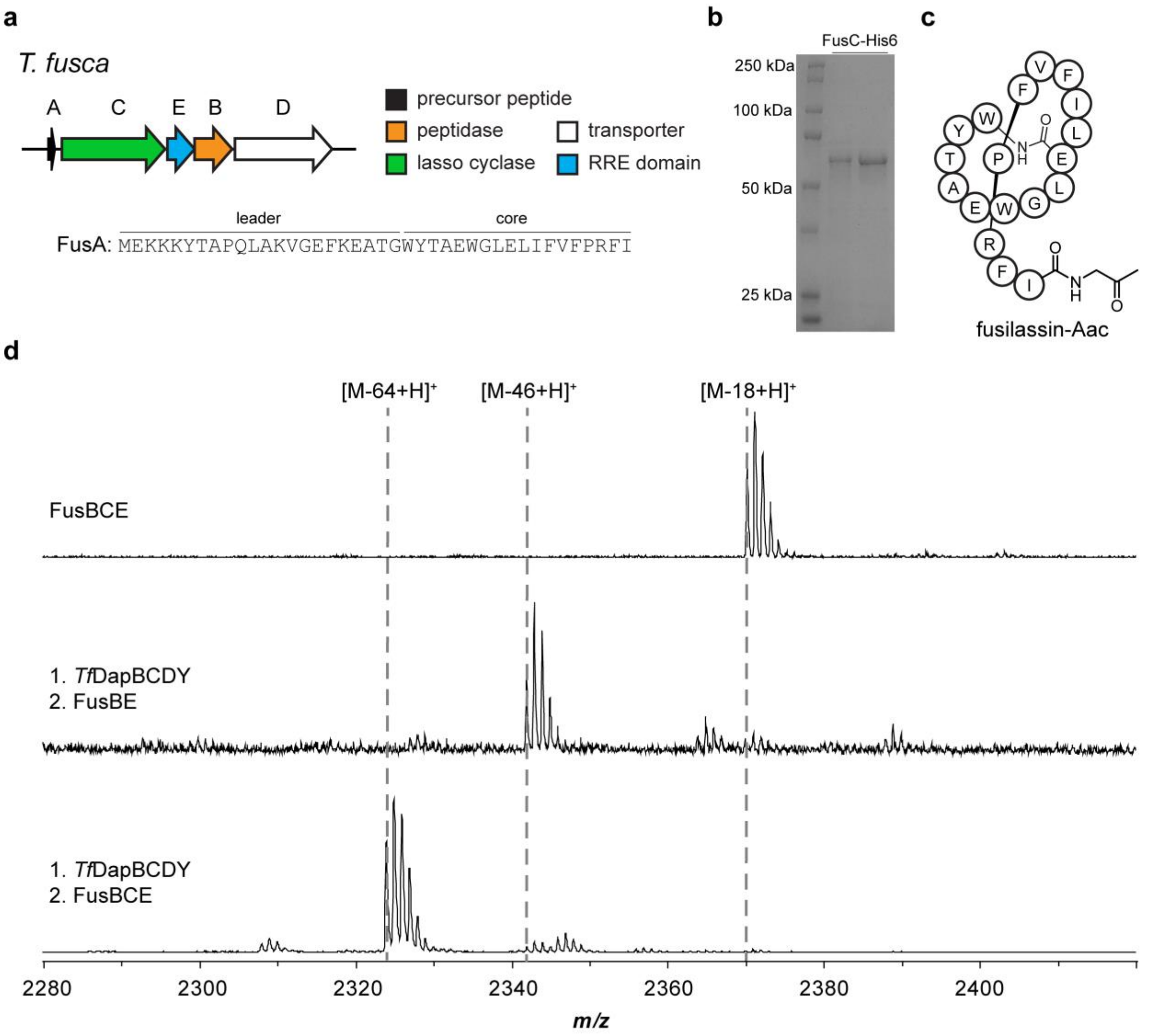

**Figure S48. Production of fusilassin-Aac.** a, Biosynthetic gene cluster diagram for fusilassin. b, SDS-PAGE gel for expressed and purified FusC-His6 protein. c, Fusilassin-Aac bubble diagram. d, MALDI-TOF mass spectra for modification of the chimeric daptide-fusilassin precursor peptide by either fusilassin enzymes, daptide enzymes, or both. Spectra were collected after desalting by C18 ZipTip. Key expected masses: fusilassin-Thr, 2370 Da; linear fusilassin-Aac, 2342 Da; fusilassin-Aac, 2324 Da. Abbreviations: Aac, aminoacetone.

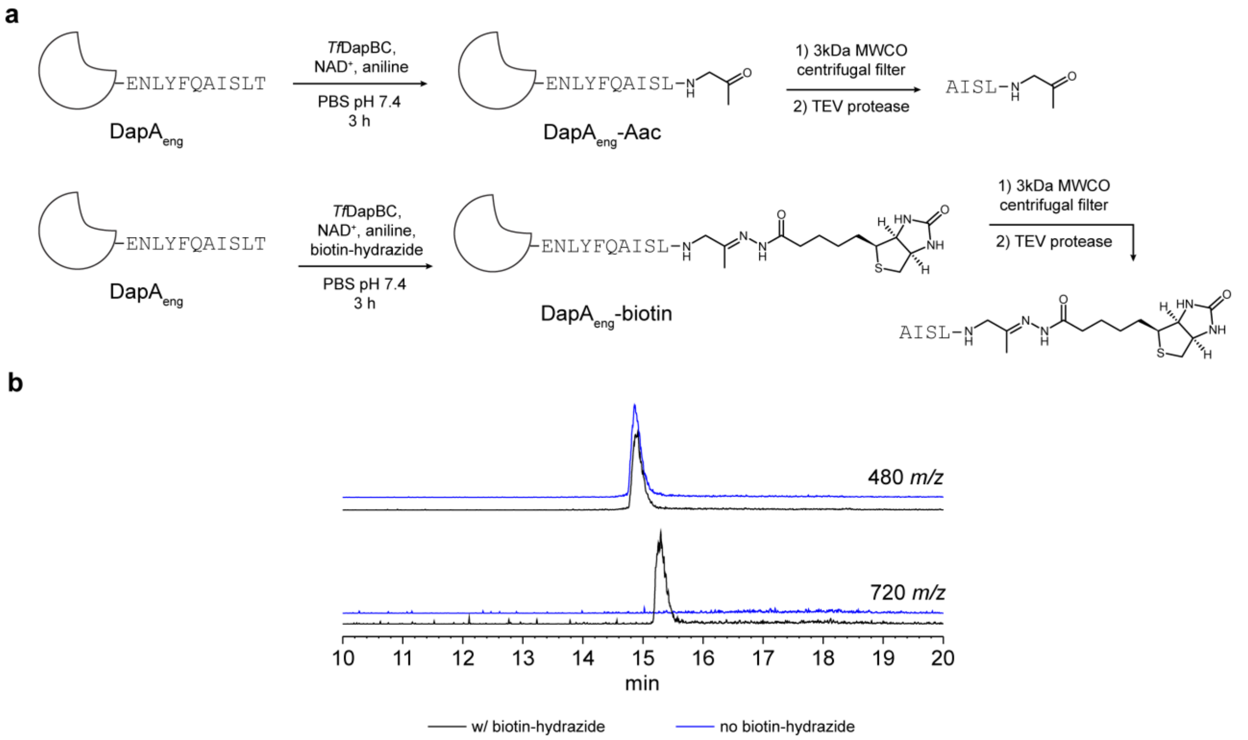

**Figure S49. Biotinylation of (MBP)DapA$_{eng}$.** a, The reaction scheme is shown for *in vitro* generation of the C-terminal Aac group and subsequent biotinylation. b, Extracted ion chromatograms are shown for Aac-modified peptide ([M+Na]$^+$: 480 *m/z*) and biotinylated peptide ([M+Na]$^+$: 720 *m/z*). Unmodified peptide was not detected. Abbreviations: Aac, aminoacetone.

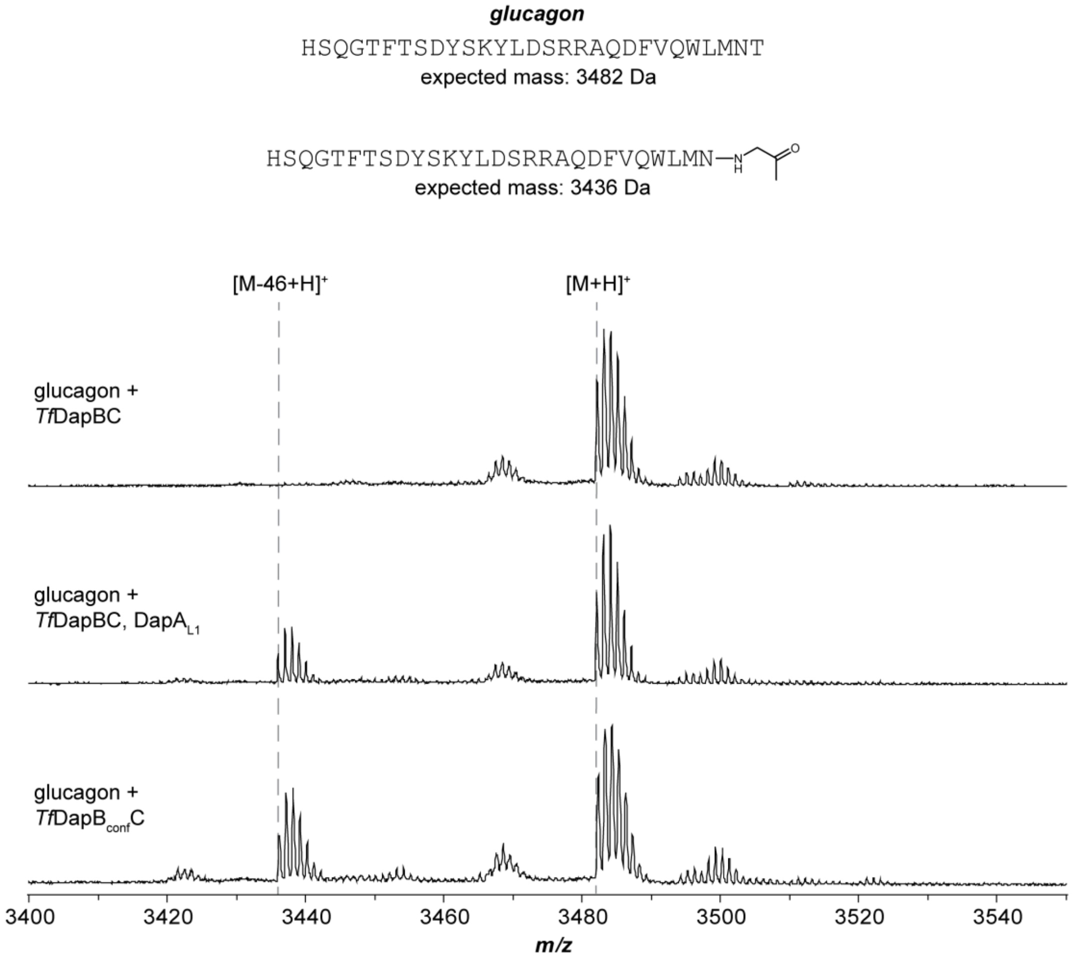

**Figure S50. Production of aminoacetone-modified glucagon.** MALDI-TOF mass spectra were collected after desalting by C18 ZipTip.

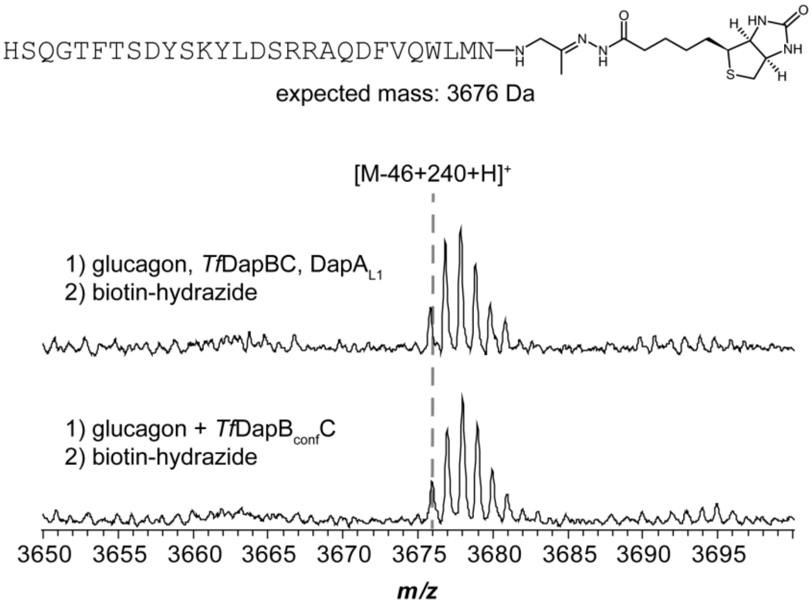

**Figure S51. Biotinylation of glucagon peptide.** MALDI-TOF mass spectra were collected after desalting by C18 ZipTip.

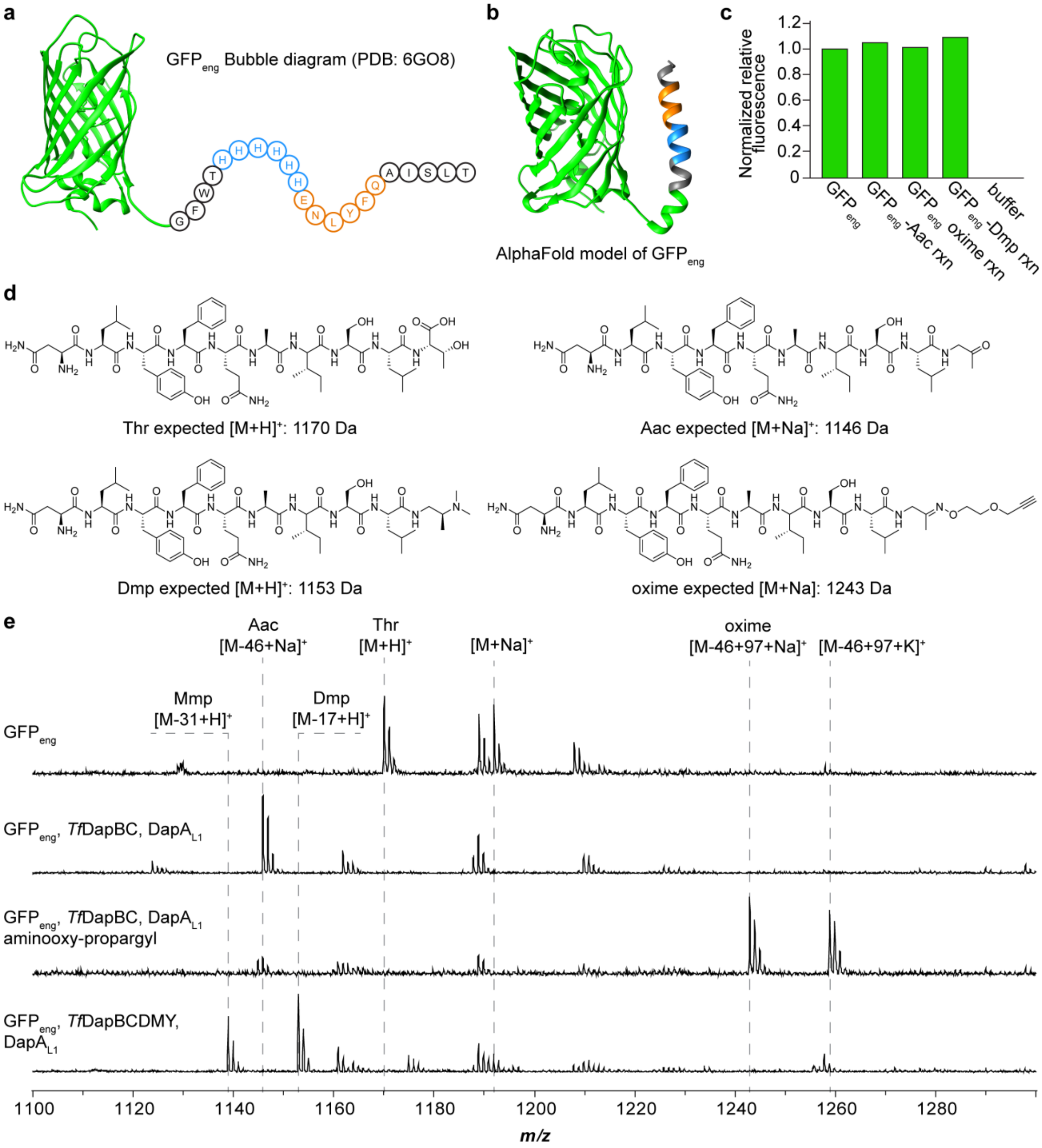

**Figure S52. GFP$_{eng}$ substrate evaluation and bioconjugation reactions**. a, GFP$_{eng}$ bubble diagram (PDB: 6GO8).[19] b, AlphaFold model of GFP$_{eng}$. Blue, His$_6$-tag; Orange, TEV protease site. c, Relative fluorescence of GFP$_{eng}$ reactions measured after 24 h. Samples were diluted ten-fold and results are normalized to the control sample containing only GFP$_{eng}$. d, GFP$_{eng}$ peptide fragments after GluC digestion. e, MALDI-TOF mass spectra for GluC digests of reactions containing GFP$_{eng}$. Mass spectra were collected after desalting by C18 ZipTip. Abbreviations: Aac, aminoacetone; Mmp, monomethylpropane-1,2-diamine; Dmp, dimethylpropane-1,2-diamine.

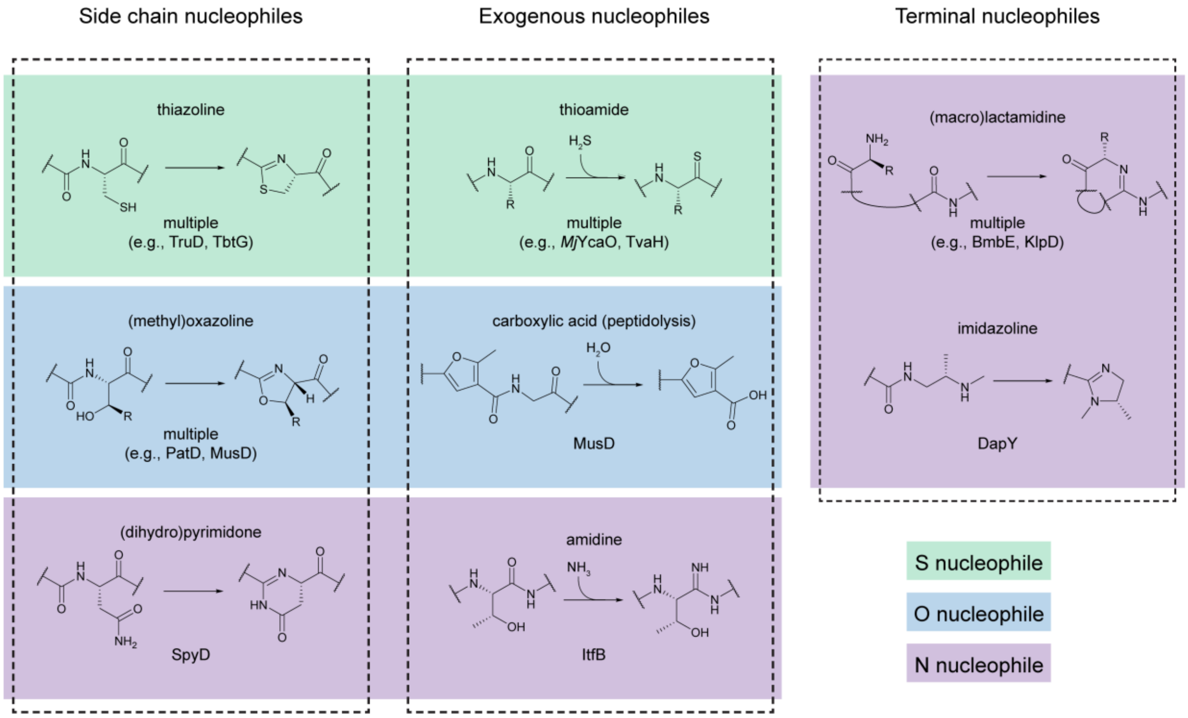

**Figure S53. Characterized YcaO-catalyzed reactions.**[9–16] Reaction schemes are shown for prototypic YcaO-catalyzed post-translational modifications. Reactions are sorted by nucleophile source (*i.e.*, side chain, exogenous, or terminal nucleophile) and colored by atomic identity of the nucleophilic atom.

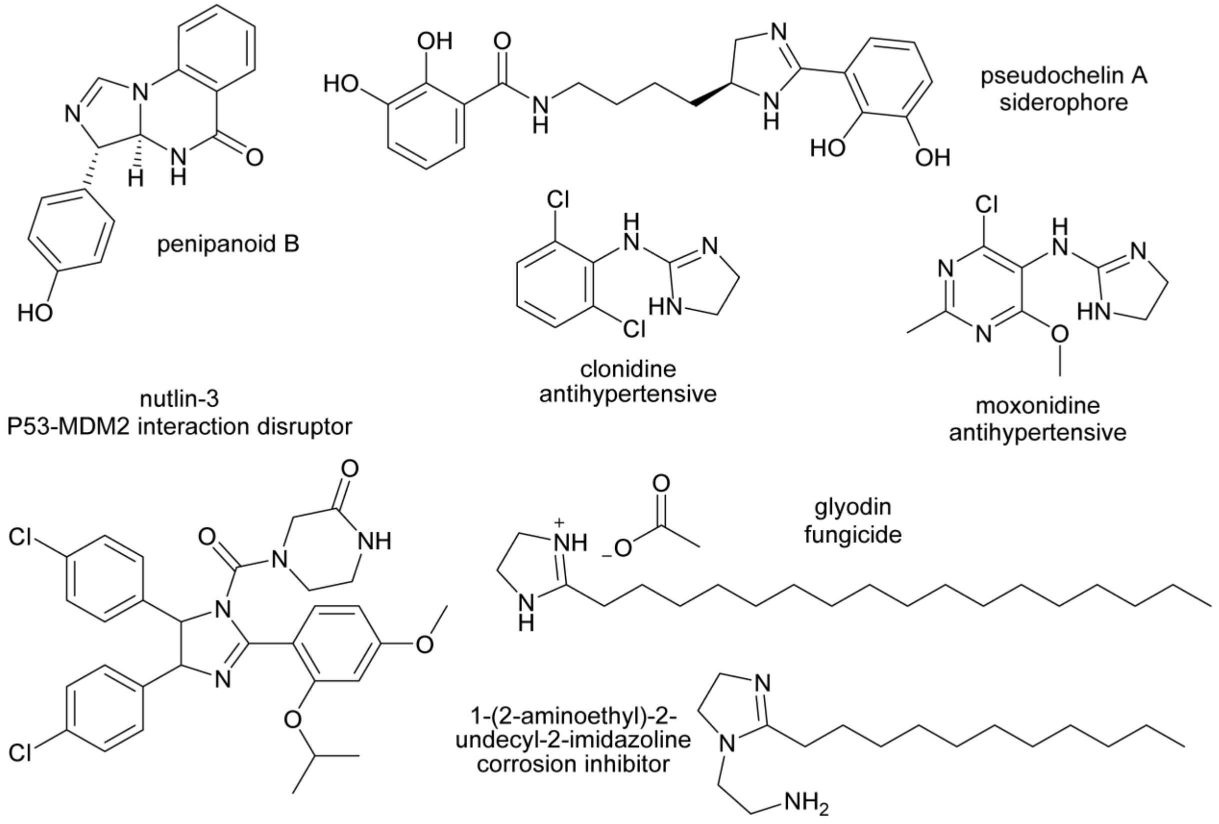

**Figure S54. Representative imidazoline compounds.**[20–25]
